# Structure-based design of stabilized recombinant influenza neuraminidase tetramers

**DOI:** 10.1101/2021.05.17.444468

**Authors:** Daniel Ellis, Julia Lederhofer, Oliver J. Acton, Yaroslav Tsybovsky, Sally Kephart, Christina Yap, Rebecca A. Gillespie, Adrian Creanga, Tyler Stephens, Deleah Pettie, Michael Murphy, Andrew J. Borst, Young-Jun Park, Kelly K. Lee, Barney S. Graham, David Veesler, Neil P. King, Masaru Kanekiyo

## Abstract

Influenza virus neuraminidase (NA) is a major antiviral drug target and has recently reemerged as a key target of antibody-mediated protective immunity. Here we show that recombinant NAs across all non-bat subtypes adopt various tetrameric conformations, including a previously unreported “open” state that may help explain poorly understood variations in NA stability across viral strains and subtypes. We used homology-directed protein design to uncover the structural principles underlying these distinct tetrameric conformations and stabilize multiple recombinant NAs in the “closed” state. In addition to improving thermal stability, conformational stabilization improved affinity to protective antibodies elicited by viral infection, including antibodies targeting a quaternary epitope and the broadly conserved catalytic site. The stabilized NA proteins can also be integrated into viruses without affecting fitness. Our findings provide a deeper understanding of NA structure, stability, and antigenicity, as well as a roadmap towards structure-based discovery of NA-directed therapeutics and vaccines.

## Introduction

Influenza viruses are negative-sense RNA viruses in the family *Orthomyxoviridae*. The two major viral glycoproteins, hemagglutinin (HA) and neuraminidase (NA), facilitate viral entry and egress from host cells, respectively. NA is an enzyme that binds and cleaves sialosides from glycans on the host cell surface to facilitate the release of nascent viral particles from infected cells, and is also proposed to assist other stages in the virus replication cycle^1,2^. Like HA, influenza A NAs are divided into subtypes that are clustered into the phylogenetically defined groups 1 (N1, N4, N5, and N8) and 2 (N2, N3, N6, N7, and N9), whereas influenza B NAs are clustered in a separate branch^3^. NA is a homotetrameric, type II integral membrane protein with a short cytoplasmic N-terminal domain. The C-terminal catalytic domain folds into a disulfide-stabilized and glycosylated six-bladed beta propeller and is supported by a hypervariable stalk domain^4,5^. Published crystal structures of NA catalytic domains, either proteolytically released from virions or produced recombinantly, have consistently shown that four identical subunits interact non-covalently to form a globular “head” with four-fold symmetry^2,6^. The head features deep pockets on the exterior face of each protomer comprising the catalytic residues. NA contains several Ca^2+^ binding sites, one of which is highly conserved and supports the periphery of the catalytic pocket on each protomer, while another site at the four-fold axis is frequently, but not universally, observed across various subtypes^7^. Ca^2+^ ions have been shown to assist catalytic activity^2^ and have been resolved to varying degrees in crystal structures, with reports and predictions of increased structural flexibility in their absence^8,9^. These observations, combined with remote epistatic effects that impact sialoside binding and resistance to catalytic inhibitors, highlight the relatively plastic conformational landscape of NA^10–14^.

While multiple studies have shown that antibodies against NA correlate with immunity against influenza viruses^15–20^, substantial gaps remain in our knowledge of NA-directed immunity, particularly in terms of the relationships between NA structure, conformational stability, and antigenicity. The antigenicity of NA has long been detailed using monoclonal antibodies (mAbs) isolated from immunized and/or infected mice, which provided an initial understanding of its distinct antigenic regions—some of which span adjacent protomers—as well as mechanisms of both enzymatic inhibition and protection^21–25^. Although an NA-based vaccine, extracted from viral membranes and inactivated by formalin, improved humoral NA-directed inhibition compared to commercial vaccines in humans in a Phase I clinical trial more than 25 years ago^26,27^, only preclinical studies of NA-based vaccines have been reported since^28–32^. More recently, studies comparing antibodies elicited by infection or immunization with split virus vaccines in humans as well as mice have clearly demonstrated that infection results in substantially more robust NA-directed humoral responses^33^. Protective human mAbs have therefore most frequently been isolated from infected individuals, with the recent exception of mAbs isolated from a minority of subjects receiving H7N9 vaccines that generated detectable anti-NA responses^33–36^. These studies have further defined NA epitopes targeted by protective antibodies, most notably leading to the isolation of broadly cross-reactive and protective antibodies from an H3N2-infected individual that bind the catalytic site^36^. The disconnect between infection- and vaccine-elicited responses appears to be due to existing vaccine manufacturing processes causing NA to unfold and lose its native structure^33^, suggesting that there is room to improve the composition and stability of NA-based immunogens.

Despite the atomic structure of NA having been known for decades, its conformational stability in solution has not been fully detailed. In agreement with crystal structures, tomographic models of NA on the surface of virions have shown a compact, “closed” four-fold symmetric arrangement of the catalytic domains^37,38^. In contrast to other sialidases that share the same six-bladed propeller fold and can function as monomers^39^, it has frequently been reported that tetramer formation is a prerequisite for the function of influenza NA^40–44^. Consequently, catalytic activity has often been considered an indicator of the biologically relevant conformation^45^. Recent attempts to understand and improve NA stability have shown some promise, and have suggested connections between exposure to Ca^2+^ ions, antigenicity, oligomerization, and general stability^44,46,47^.

Here we characterize a previously unreported “open” tetrameric conformation commonly adopted by recombinant NAs and provide a general structure-based design strategy for stabilizing the closed tetrameric state. Closed versions of recombinant N1 NAs have improved stability and affinity to infection-elicited protective antibodies relative to their wild-type counterparts, providing new opportunities to design and discover NA-based vaccines and therapeutics.

### Tetrameric Conformations of Recombinant NA Proteins Vary Across Influenza Strains and Subtypes

We expressed and purified recombinant soluble NA proteins from every non-bat influenza A subtype and two influenza B viruses using a previously described configuration^48^ in which the NA head domains are genetically fused to the hVASP tetramerization domain^49^ (Fig. 1A–B). The proteins were secreted from mammalian cells and purified from clarified supernatants, and yielded homogeneous size exclusion chromatography (SEC) profiles featuring a major peak corresponding to the estimated elution volume of a tetramer, with minimal aggregation (Extended Data Fig. 1A). However, two-dimensional (2D) class averages obtained by electron microscopy analysis of negatively stained samples revealed substantial structural heterogeneity (Fig. 1C). Two tested NAs each from the N2 subtype and influenza B viruses, as well as representative NAs from N3 and N5, formed “closed” tetrameric structures in which the head domains packed tightly and resembled the four-fold symmetric structure classically observed by X-ray crystallography. In contrast, representative NAs from the N1 and N6 subtypes formed “open” tetramers in which the head domains did not form a single, compact structure. These NAs instead exhibited a clover-like appearance characterized by a thin stalk tethered to independent densities corresponding to the four globular head domains. Representative NAs from the N4, N7, N8, and N9 subtypes, in addition to a third tested N2 strain, formed mixtures of open and closed tetramers. We also produced and analyzed a previously reported N9 NA construct containing the stabilizing mutation Y169aH^23,47^, which, despite showing an improved proportion of the closed state compared to its WT counterpart, still formed a significant population of open tetramers (Extended Data Fig. 1A). Furthermore, we obtained two N1 and one N2 recombinant NAs from the BEI Resources Repository at the NIH, which were also genetically fused to hVASP. The N2 NA was found to adopt the closed conformation and further form rosettes through the hVASP domains, while both N1 NAs appeared open or disordered (Extended Data Fig. 1B).

**Fig. 1.**
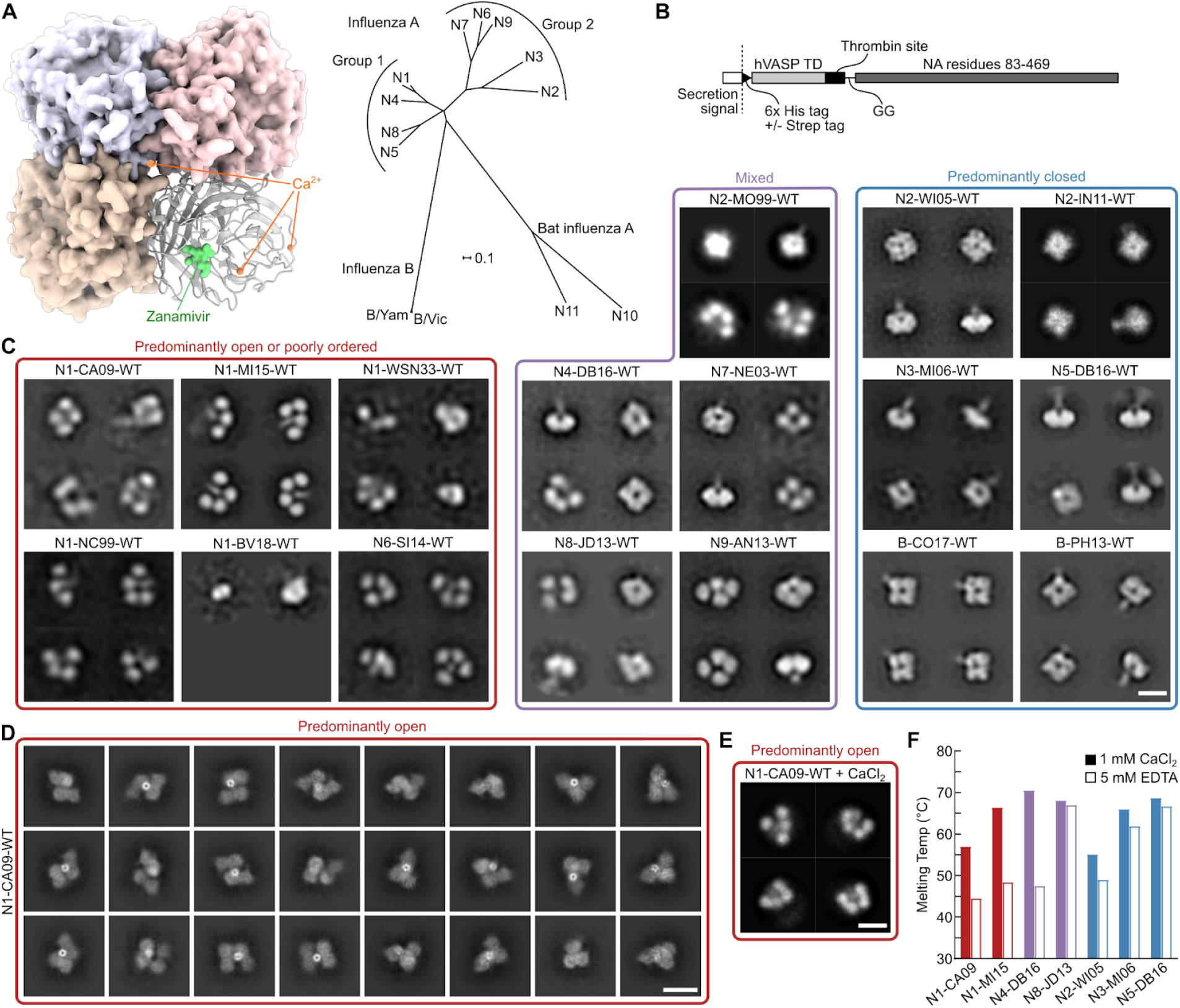
Tetrameric Conformations of Recombinant NA Proteins Varies Across Influenza Strains and Subtypes. (A) NA structure and phylogeny. (Left) Model of NA structure as observed crystallographically, featuring a bound NA inhibitor (Zanamivir, green) and Ca^2+^ ions (orange). Model from PDB ID 4B7Q with the Ca^2+^ ion near the four-fold axis placed using an alignment with PDB ID 3NSS. (Right) Phylogenetic tree of NA of influenza A and B viruses. Scale bar, 0.1 amino acid substitutions per site. (B) Construct diagram for recombinant NA proteins used in this work. The globular head domain of NA (residues 83–469, N1 numbering) was genetically fused to the hVASP tetramerization domain (TD) by a flexible GG linker and a thrombin protease cleavage site. All constructs contained an N-terminal hexahistidine tag, and some constructs additionally contained a Strep Tag. See Supplementary Item 1 for amino acid sequences. (C) NS-EM 2D class averages of NA (scale bar, 10 nm). (Left) NA preparations from strains that show predominantly open or poorly ordered tetramers. (Middle) NA preparations from strains that show mixtures of closed and open tetramers. (Right) NA preparations from strains that show predominantly closed tetramers similar to those classically observed in NA crystal structures. (D) Cryo-EM 2D class averages of recombinant N1-CA09-WT (scale bar, 10 nm). (E) NS-EM 2D class averages of recombinant N1-CA09-WT in the presence of 1 mM CaCl_2_ (scale bar, 10 nm). (F) Melting temperatures of representative open, mixed, and closed recombinant NAs (see also Extended Data Fig. 1).

To determine whether destabilization of the closed tetrameric conformation was due to the conditions the proteins experienced during negative stain electron microscopy (NS-EM), the WT NA from the H1N1 strain A/California/07/2009 (abbreviated as N1-CA09-WT) was vitrified and imaged by cryoelectron microscopy (cryo-EM). 2D classification of the data revealed heterogeneity consistent with the NS-EM data, with N1-CA09-WT forming a variety of poorly ordered open tetramers (Fig. 1D). However, several 2D classes indicated that adjacent head domains formed dimers reminiscent of conformations observed in distantly related proteins with the same six-bladed propeller fold, such as the hemagglutinin (H), hemagglutinin-neuraminidase (HN), and G proteins of paramyxoviruses^50,51^. We found no clear correlation between SEC profiles (Extended Data Fig. 1C) or catalytic activity (Extended Data Fig. 1D) and the conformations observed by NS-EM: open tetramers remained catalytically active. Although NA contains an interprotomeric Ca^2+^ binding site that thermally stabilizes the protein^46^, the addition of 1 mM CaCl_2_ to purified N1-CA09-WT prior to NS-EM did not result in tetramer closure, indicating that potential Ca^2+^ deficiency does not explain the conformational differences between different recombinant NAs (Fig. 1E). We also determined the thermal stability of multiple NAs using nano differential scanning fluorimetry (nanoDSF) by monitoring intrinsic tryptophan fluorescence in buffers containing either 1 mM CaCl_2_ or 5 mM ethylenediaminetetraacetic acid (EDTA) (Fig. 1F and Extended Data Fig. 1E). NAs from representative N3 and N5 (identified as closed by NS-EM) and N8 (mixed) strains were found to have the highest melting temperatures (T_m_), all above 60°C regardless of buffer additives (Fig. 1F). N8-JD13-WT (A/Jiangxi Donghu/346/2013) had an incomplete yet clear transition between 46 and 52°C, depending on the buffer additive, suggesting a partial unfolding event or the unfolding of a minor population of less stable species. Another closed NA, N2-WI05-WT (A/Wisconsin/67/2005), was less stable but also relatively insensitive to Ca^2+^, with T_m_s of 55.1°C in the presence of Ca^2+^ and 49.0°C in 5 mM EDTA. In contrast, the open N1-CA09-WT and N1-MI15-WT (A/Michigan/45/2015), and particularly the mixed N4-DB16-WT (A/Redknot/Delaware Bay/310/2016), showed much greater sensitivity to Ca^2+^, with T_m_s below 50°C in the presence of EDTA becoming 12.5–23°C higher in the presence of Ca^2+^. These data indicate that the thermal stability of recombinant NAs can vary over a wide range, and suggest that Ca^2+^ contributes less to the stability of NAs that tend to form closed tetramers, possibly because head tetramerization has an independent stabilizing effect.

### Stabilization of the Closed Tetrameric State of N1 CA09 NA

Although high-resolution structures for NA in its native context in the viral membrane are not yet available, cryo-electron tomographic reconstructions at modest resolution suggest that NA on the surface of H1N1 CA09 virions forms closed tetramers^38^. This observation is further supported by the protective mAb CD6, isolated from a mouse infected with H1N1 virus, which binds a quaternary epitope spanning two subunits of the NA tetramer^52^. Inspired by these observations and the potential utility of recombinant NA antigens with improved stability and homogeneity, we set out to stabilize N1 NA in the closed tetrameric state.

The inter-protomer interface of N1-CA09-WT (PDB ID 4B7Q) has 3,172 Å^2^ of buried surface area between each protomer, but features many aqueous cavities and non-ideal polar contacts. We attempted to design closed N1-CA09 tetramers by stabilizing the inter-protomeric interface using homology-directed mutations inspired by NAs we identified as closed, as well as novel mutations identified purely by structural modeling (Fig. 2A). Based on our data indicating that recombinant N2, N3, and N5 NAs form predominantly closed tetramers, we analyzed mutational differences at the interface between representative NAs from each of these subtypes and N1-CA09-WT. Due to the higher sequence similarity of N4 to N1 NAs, an N4 NA structure was also included in this analysis even though it adopted a mixture of the open and closed states. To organize our design strategy, we identified residues involved in interprotomer interactions by visual inspection and categorized them into four “spaces” (A, B, C, and D) that spanned the entire interface (Fig. 2B). We developed a computational pipeline for homology-directed design using the Rosetta macromolecular modeling software^53,54^, in which potential interface-stabilizing mutations derived from closed NAs were symmetrically modeled both individually and in combination within each space. In parallel, design trajectories that allowed Rosetta to identify novel mutations without restriction to naturally observed substitutions were performed. Thirty-two designs were selected for experimental characterization, comprising designs with mutations restricted to each individual space as well as combined across multiple spaces.

**Fig. 2.**
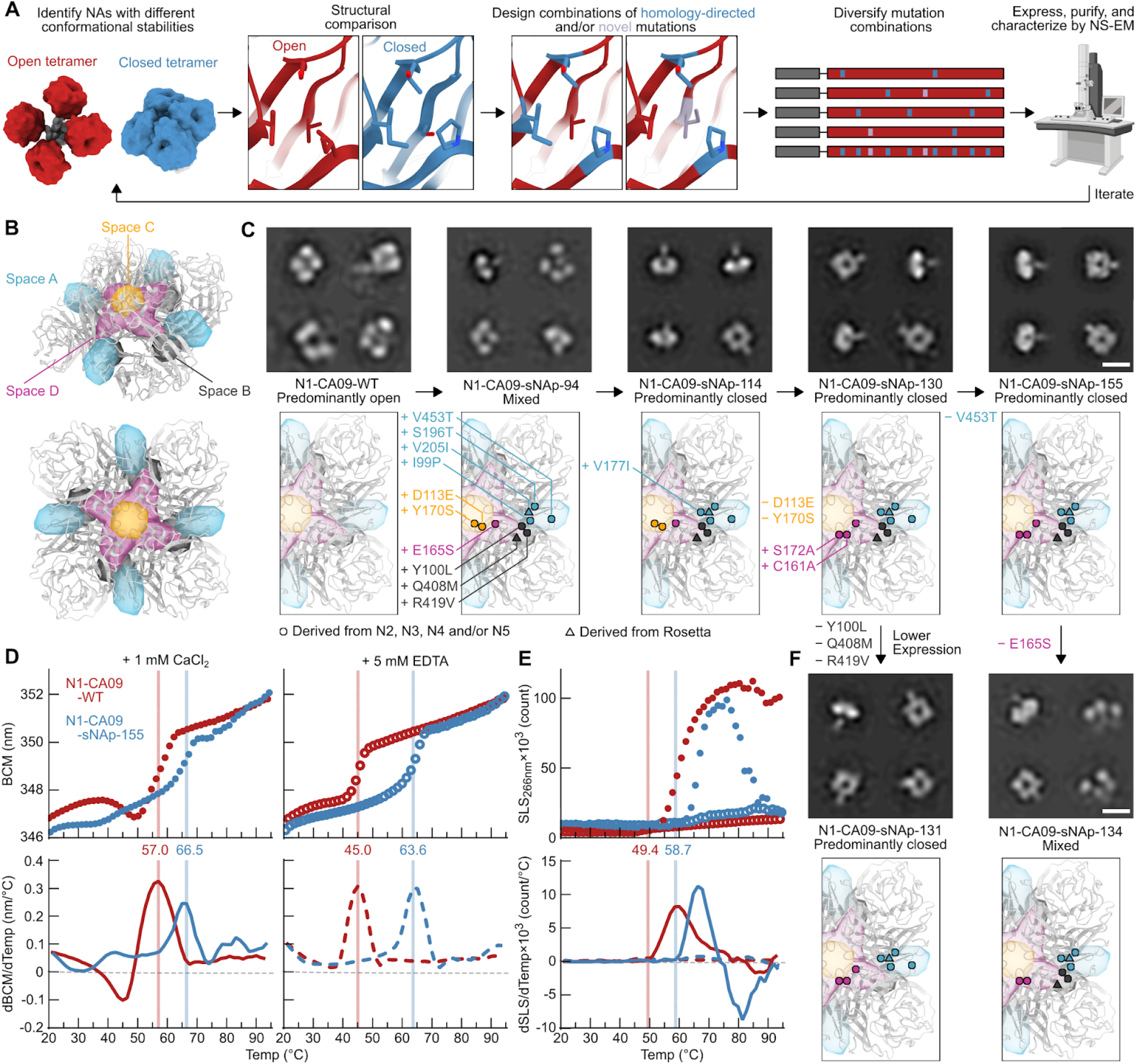
Stabilization of the Closed Tetrameric State of N1 CA09 NA. (A) Design and analysis pipeline for stabilization of closed tetrameric NAs using homology-directed mutations and computational protein design. Models constructed using PDB-IDs 4B7Q, 6BR5, and 1USE. (B) Structural depiction of the four different spaces in NA targeted for design. Two views of the CA09 NA tetramer are shown. (C) NS-EM 2D class averages (top) and designed mutations (bottom) in a series of CA09 NA mutants that exhibit varying degrees of tetramer closure (scale bar, 10 nm). (D) Thermal denaturation of N1-CA09-WT and N1-CA09-sNAp-155 in the presence of 1 mM CaCl_2_ (closed circles and solid lines) or 5 mM EDTA (open circles and dashed lines), monitored by intrinsic tryptophan fluorescence. Top panels show raw data, while lower panels show smoothed first derivatives used to calculate melting temperatures. The barycentric mean (BCM) of the fluorescence emission spectra is plotted. (E) SLS during thermal denaturation of N1-CA09-WT and N1-CA09-sNAp-155 in the presence of 1 mM CaCl_2_ or 5 mM EDTA, with aggregation temperatures shown for CaCl_2_-treated samples. Data are plotted as in panel D. (F) NS-EM 2D class averages (top) and mutations (bottom) in two CA09 NA mutants derived from N1-CA09-sNAp-130 and N1-CA09-sNAp-155, respectively, that explore the roles of spaces A, B, and D in tetramer closure and expression (scale bar, 10 nm).

Genes for each stabilized NA protein (sNAp) were synthesized as genetic fusions to the hVASP tetramerization domain (Fig. 1B) and transfected into mammalian cells to assess expression. Nine designs that were detectable in cell culture supernatants by SDS-PAGE were purified by affinity chromatography and analyzed by NS-EM (see Supplementary Item 1 for amino acid sequences). Of these nine designs, only one (N1-CA09-sNAp-94) contained a substantial population of closed structures, with approximately half of the particles classified as closed based on the 2D class averages (Fig. 2C). To improve this design, one additional N2- and N3-inspired mutation was added that appended an individual methyl group to a valine in space A (V177I; N1-CA09-sNAp-114), which is distal to the C4 axis and near the CD6 binding site, resulting in predominantly closed particles by NS-EM (Fig. 2C), with only a minor fraction of particles in the open state still detectable.

Two additional rounds of design were performed to revert unproductive mutations from N1-CA09-sNAp-114 while maintaining closure and maximizing expression levels. Space C, which includes residues immediately surrounding the C4 axis, featured mutations derived from N4 NA that could be reverted without deleterious effects. Further, the N5-inspired combination of C161V and S172A was added to remove an unpaired cysteine and slightly repack space D, adjacent to space C on the “underside” of NA, which greatly improved expression (N1-CA09-sNAp-130). N1-CA09-sNAp-155, which reverted an extra mutation at the surface-exposed position 453 of N1-CA09-sNAp-130, maintained closure with similar expression to N1-CA09-WT and had its stabilizing mutations maximally buried to maintain proper antigenicity (Fig. 2C). Relative to N1-CA09-WT, N1-CA09-sNAp-155 features four mutations in space A, three in space B (on the underside of NA distal to the C4 axis), and three in space D. sNAp-130 and sNAp-155 were used interchangeably as representative sNAps for N1 CA09.

We found that the tetramer-stabilizing mutations in N1-CA09-sNAp-155 increased the T_m_ by 9.5 and 19.1°C in the presence of CaCl_2_ and EDTA, respectively, when compared to N1-CA09-WT (Fig. 2D). Sensitivity to Ca^2+^ was substantially decreased, with only a 2.9°C difference between Ca^2+^- and EDTA-containing conditions. This finding is consistent with our earlier observation that the thermal stability of closed tetramers tended to be less dependent on Ca^2+^ (Fig. 1F). Monitoring static light scattering (SLS) also revealed an increase in aggregation temperature for N1-CA09-sNAp-155 of 9.7°C in the presence of Ca^2+^ (Fig. 2E). Interestingly, neither protein aggregated in the presence of EDTA. To assess stability over time, conformational stability of the closed state was monitored by NS-EM for 15 days after storage at 4°C and showed retention of the closed state during this period (Extended Data Fig. 2A). Hydrogen-deuterium exchange mass spectrometry (HDX-MS) was used to collect data on a small set of deuterium-exposed peptides from both N1-CA09-WT and N1-CA09-sNAp-130, which indicated multiple regions of differential stability (Extended Data Fig. 2B). Several peptides were identified near the inter-protomer interface and the catalytic pocket that showed decreased deuterium uptake for the sNAp, including one peptide that includes the main catalytic residue (Y402) and another peptide containing the substrate-binding residue R368. One peptide (residues 215–224) was more dynamic in the sNAp, which includes a solvent-exposed loop and part of a beta-strand that is adjacent to two stabilizing mutations (S196T and V205I). Overall, most of the changes observed by HDX-MS showed a reduction in solvent accessibility or an increase in local ordering consistent with the sNAp adopting a more “closed” conformation compared to N1-CA09-WT. Finally, N1-CA09-sNAp-130 was catalytically active, exhibiting activity that was similar to N1-CA09-WT (Extended Data Fig. 2C).

While the V177I mutation suggested that improved packing in space A assists closure, the specific contributions of other mutations were less clear since they were introduced simultaneously in N1-CA09-sNAp-94. Reversion of the three space B mutations in N1-CA09-sNAp-130 did not decrease tetrameric closure (N1-CA09-sNAp-131), although it did reveal a role for these mutations in increasing sNAp expression (Fig. 2F). Strikingly, reversion of the single N5-derived space D mutation E165S from N1-CA09-sNAp-155 resulted in a roughly equal mixture of open and closed tetramers (N1-CA09-sNAp-134). Together, these data demonstrate that combining mutations in spaces A and D can drive complete closure of CA09 NA.

### Structural and Bioinformatic Analysis of Closed State Stabilizing Mutations

To better understand the roles of our stabilizing mutations, we determined a cryo-EM structure of N1-CA09-sNAp-155 at 3.2 Å resolution (Fig. 3 and Extended Data Fig. 3). 2D class averages obtained from vitrified samples exclusively showed closed NA tetramers resembling those observed in crystal structures, and the 3D reconstruction confirmed that N1-CA09-sNAp-155 forms a closed tetramer in solution (Fig. 3A). The structure is highly similar to N1 CA09 crystal structures (1.18 and 1.83 Å backbone root-mean-square deviation (RMSD) to PDB 4B7Q over one and all four chains of the tetramer, respectively). The main differences were in a few solvent-exposed loops that were poorly ordered in N1-CA09-sNAp-155, including the C-terminal region from residues 458–469, the ‘150-loop’ consisting residues 145-150, and the ‘430-loop’ consisting residues 429-437. The apparent flexibility of the 150- and 430-loops is consistent with previous structural observations and computational modeling of these regions^55–57^. Additionally, a backbone segment adjacent to the space A mutations was slightly rearranged such that W455 packed closely against the introduced mutations and W457 moved away from its native position at the interface (Fig. 3B). Despite this rearrangement, none of the mutated positions in spaces A, B, and D showed substantial changes in their main chain coordinates and, other than their interactions with W455 and W457, the intended interactions were formed. For example, P99 and T196 (T195 in N2 numbering) in space A form a central hydrophobic contact across the interface that closely matches the same interaction seen in the NA structures used for homology-directed design (Fig. 3B and 4A). Likewise, I177 (I176 in N2 numbering) provides additional hydrophobic packing in both N1-CA09-sNAp-155 and the NAs of other subtypes relative to N1 CA09, while I205 adds further hydrophobic packing that was favored by Rosetta. In space B, the Rosetta-derived Q408M and the homology-inspired Y100L and R419V simultaneously provide improved hydrophobic packing and decrease the number of partially buried polar groups at the interface.

**Fig. 3.**
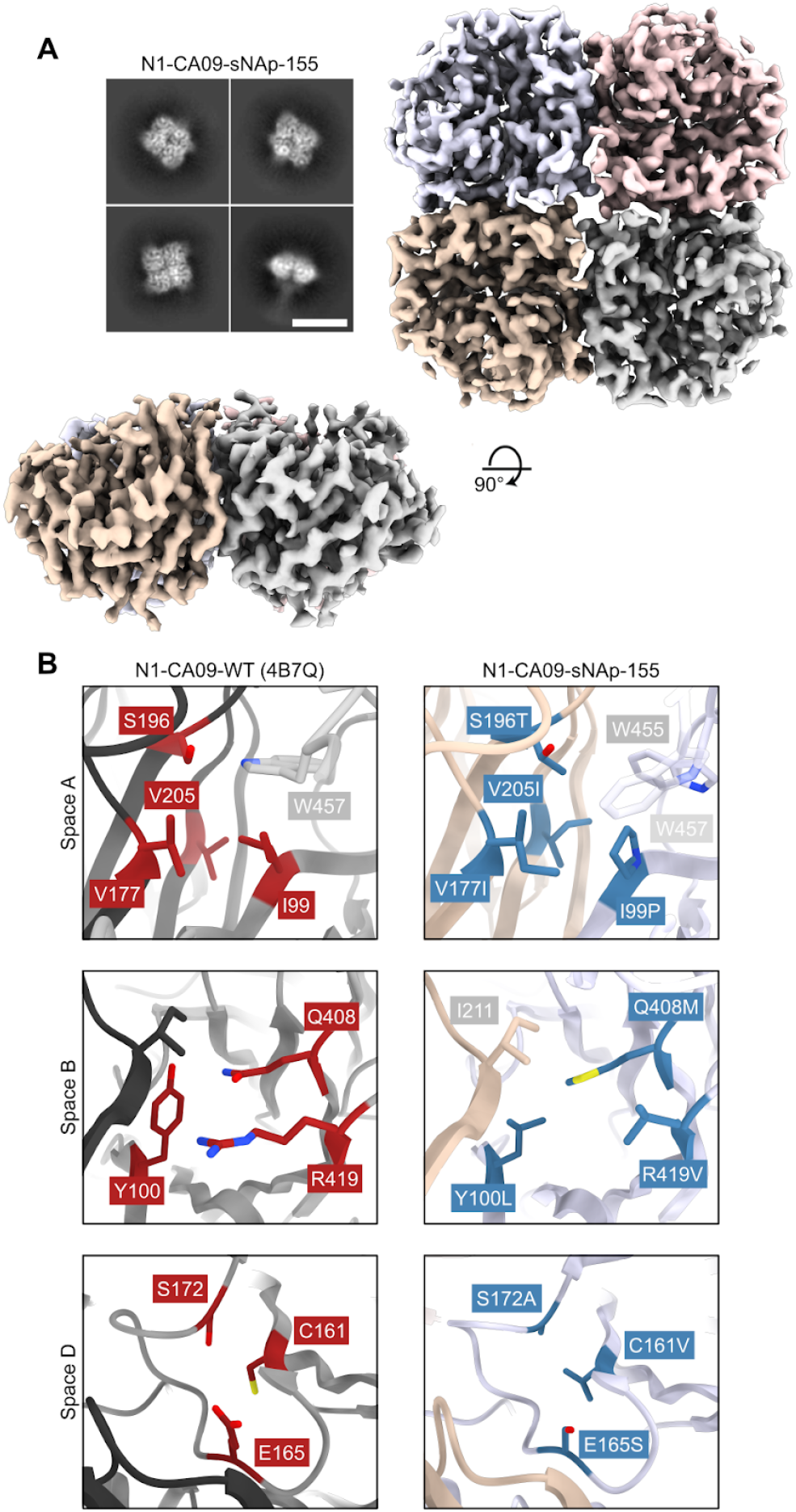
Cryo-EM structure of N1-CA09-sNAp-155. (A) Cryo-EM 2D class averages for N1-CA09-sNAp-155 showing fully closed tetramers (scale bar, 10nm) (top left). Cryo-EM map, colored by individual head domain, viewed down the four-fold symmetry axis (top right) and an orthogonal view (bottom left). (B) Snapshots of the three major regions targeted for mutation: WT N1-CA09 (PDB 4B7Q, left), with residues targeted for mutation indicated (red). Space A (top), Space B (middle), and space D (bottom). Corresponding regions in the N1-CA09-sNAp-155 single particle reconstruction (right), with mutated residues indicated (blue) and non-mutated residues involved in packing colored by chain as in (A).

In contrast to intuitive stabilization mechanisms such as the cavity-filling mutations employed in space A, the N5-inspired space D mutation E165S does not have an obvious structural explanation. Crystal structures of N2, N3, N4, and N5 NAs all show less bulky residues at position 165, including S, T, and V (Fig. 4A). Interestingly, in all known NA structures a substantial cavity surrounds D103, adjacent to the 160-loop and near the inter-protomeric interface, raising the possibility that the E165S mutation may function by helping pre-organize this region during tetramer assembly.

**Fig. 4.**
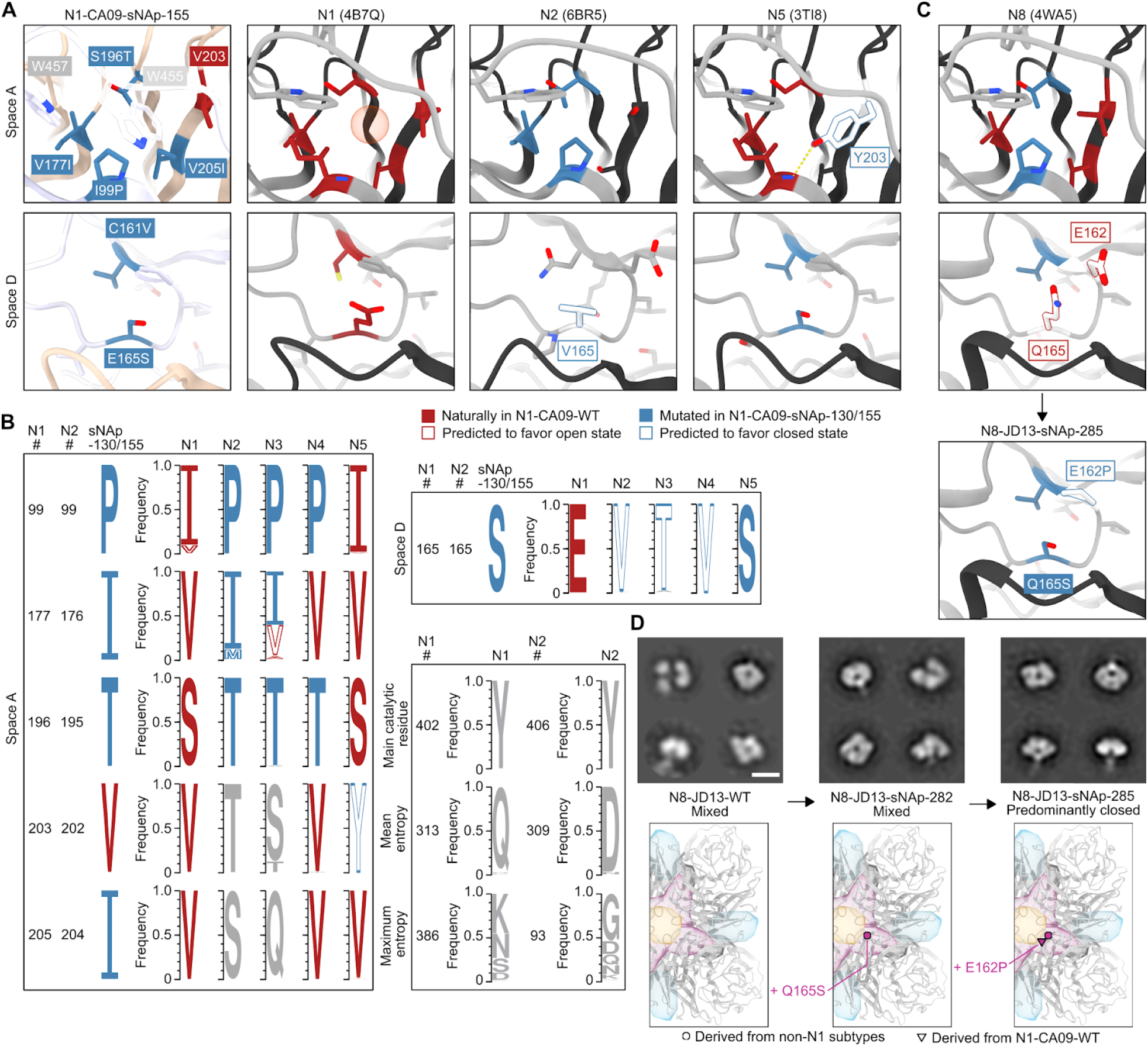
Structural and Bioinformatic Analysis of Closed State-Stabilizing Mutations. (A) Zoomed-in views of space A- and D-stabilizing mutations in the N1-CA09-sNAp-155 cryo-EM structure and corresponding regions in predominantly open N1 (PDB ID 4B7Q) and predominantly closed N2 (PDB ID 6BR5) and N5 (PDB ID 3TI8) NAs. The red sphere in N1 highlights the position of a structured water molecule in the space A cavity. (B) Sequence logos for N1-CA09-sNAp-155 and naturally occurring NAs at positions that can alter NA tetrameric conformation, derived from a compiled database of all unique sequences from each subtype prior to December 31st, 2019. Sequence logos for the main catalytic residue, a representative position of mean entropy, and the position of maximum entropy within both N1 and N2 subtypes are also shown for comparison. (C) (Top) Spaces A and D in a representative crystal structure of an N8 NA (PDB ID 4WA5) and (Bottom) design model for space D of N8-JD13-sNAp-285. (D) Mutation schematics and NS-EM comparing N8-JD13-WT, N8-JD13-sNAp-282, and N8-JD13-sNAp-285 (scale bar, 10 nm).

To better understand NA sequence-structure relationships, a comprehensive dataset of unique NA sequences from all available influenza A viruses isolated before December 31, 2019 was compiled, and the mutational frequencies of key positions in spaces A and D were evaluated within each subtype (Fig. 4B). As expected, some variation was observed in the amount of sequence diversity at each position, ranging from universal conservation of the tyrosine nucleophile in the catalytic site (Y402 and Y406 in N1 and N2 numbering, respectively) to high diversity at several surface-exposed positions (e.g., position 386 in N1 NAs). However, sequence diversity within each subtype was generally low: positions of mean entropy in N1 and N2 NAs, for which far greater numbers of unique sequences were available compared to the animal-derived N3, N4, and N5 subtypes, showed ∼97% conservation of the predominant amino acid.

Fewer than 0.05% of the 4,422 animal and 13,735 human N1 NA sequences included in our dataset contained any of the five stabilizing mutations at positions 99, 165, 177, 196, and 205 (Fig. 4B). The vast majority of sequences instead possessed the less stable identities observed in N1-CA09-WT. Position 99 was the most variable, with valine present in ∼9.8% of the sequences. We predict that both isoleucine and valine at position 99 favor the open state, and 99.9% of N1 NA sequences contain one of these two residues. NAs from N2 (dataset of 8,888 animal and 23,121 human sequences), N3 (942 animal sequences), and N4 (242 animal sequences) are also highly conserved at most of these positions and share multiple features that are distinct from N1 NAs, despite the close phylogenetic relationship between N1 and N4 (Fig. 1A). For example, P99 and T196 (195 in N2 numbering) are nearly completely conserved in all three subtypes. These residues form a consistent hydrophobic contact and play key roles in the sNAp designs. In contrast, position 177 (176 in N2 numbering) in space A is more variable in N2 and N3 NAs, with multiple identities appearing frequently, while N4 has a universally conserved valine at this position. N2-IN11-WT (A/Indiana/10/2011), which we found to be predominantly closed (Fig. 1C), contains a methionine at position 176, showing that this substitution can also favor the closed state similarly to isoleucine. N5 NAs have converged on a different solution in space A, maintaining complete conservation of the distinctive Y203, which both fills the cavity and accepts a hydrogen bond from the backbone amide of residue 99, thereby achieving the same outcome as P99 and T196 (Fig. 4A–B). In space D, further similarities are observed among the more stable subtypes (N2, N3, N4, and N5) at position 165, with less flexible smaller or beta-branched amino acids (A/S/T/V/I) found in >99.9% of all sequences in these subtypes, with valine, serine, and threonine most prevalent by far.

These structural and bioinformatic analyses provide sequence-based signatures that appear to explain the conformational differences we observed in recombinant NAs by NS- and cryo-EM. Specifically, the combination of a small side chain at position 165 and a well-packed space A correlates with the stability of the closed tetrameric conformation.

### Design of Stabilized Closed Tetrameric N8 NA

Although our finding that recombinant N8-JD13-WT forms a mixture of open and closed tetramers (Fig. 1C) is consistent with a previous report showing that N8 NAs can form monomers and dimers, a number of infection-elicited N8 NA-specific antibodies recognized a quaternary epitope^23^. The latter observation provides an incentive for tetramer stabilization. We applied the lessons learned from stabilizing N1 CA09 to generate closed N8 NA tetramers. Analysis of existing N8 NA crystal structures showed hydrophobic packing in space A to be similar to that of N4 NAs, featuring P99 and T196. However, space D contained the large, polar Q165 and a notable difference from other subtypes with a glutamic acid at position 162 instead of a proline (Fig. 4C). This combination of a well-packed space A and a more “open-like” space D appeared to explain its heterogeneous conformational phenotype. Although Q165S alone (N8-JD13-sNAp-282) was insufficient, the combination of E162P and Q165S (N8-JD13-sNAp-285) provided complete stabilization of the closed state of N8 as assessed by NS-EM, further emphasizing the cooperative effects of spaces A and D on stabilization (Fig. 4D). While N8-JD13-sNAp-285 showed only small improvements in its primary T_m_ in the presence of either Ca^2+^ or EDTA relative to N8-JD13-WT, the stabilizing mutations minimized an earlier partial unfolding transition at 45–55°C, consistent with its more homogeneous adoption of the closed state (Extended Data Fig. 4A). A similar effect on T_agg_ was observed (Extended Data Fig. 4B). However, stability assessment over time by NS-EM showed that, in contrast to N1-CA09-sNAp-155, tetramer closure of N8-JD13-sNAp-285 decreased over the course of a week at 4°C, indicating that further stabilization should be possible (Extended Data Fig. 4C). Finally, N8-JD13-sNAp-285 was still catalytically active, although slightly less so than N8-JD-WT (Extended Data Fig. 4D).

### Mutation of Stabilizing Residues in N2 NA Prevents Closed Tetramer Formation

Given the clear stabilizing effects of key positions in spaces A and D, we hypothesized that mutating them to amino acids observed in more open subtypes would destabilize the closed N2-WI05-WT tetramer. Mutations were made separately in each space to make space A more strongly resemble WT N1 NAs and space D resemble N1 or N8 NAs, while avoiding steric clashes (Extended Data Fig. 4E). The V165E mutation in space D alone was sufficient to induce global structural changes including dissociation of the heads, but also led to the formation of soluble aggregates. V165Q resulted in a more subtle effect, causing about half of the tetrameric particles to adopt the open state while the rest remained closed. In space A, an I176V/T195S double mutant remained closed, but the addition of P99I led to formation of unassembled monomers and aggregates. We next combined V165Q with I176V and T195S and found that the protein formed either open tetramers or lower-order oligomers (Extended Data Fig. 4E). Thermal denaturation of this triple mutant destabilized NA protein (desNAp), N2-WI05-desNAp-255, showed a decreased T_m_, a strongly redshifted baseline in its intrinsic fluorescence emission spectrum, and no apparent stabilization by Ca^2+^ (Extended Data Fig. 4F). Enzymatic activity was also abolished in N2-WI05-desNAp-255 (Extended Data Fig. 4G). Together, these data indicate that the same design rules that enabled stabilization of the closed tetrameric state of N1 CA09 and N8 JD13 could be applied in reverse to destabilize the closed state of N2 WI05.

### Design of Additional Closed Recombinant N1 NA Tetramers

To assess the portability of the closed state-stabilizing mutations, mutations from N1-CA09-sNAp-155 were introduced into the NA of N1 MI15, from the same post-2009 human lineage as CA09. We found that the sNAp-155 mutations clearly assisted stabilization of the closed state but were insufficient for complete closure, as N1-MI15-sNAp-155 adopted a mixture of the open and closed states (Fig. 5A).

**Fig. 5.**
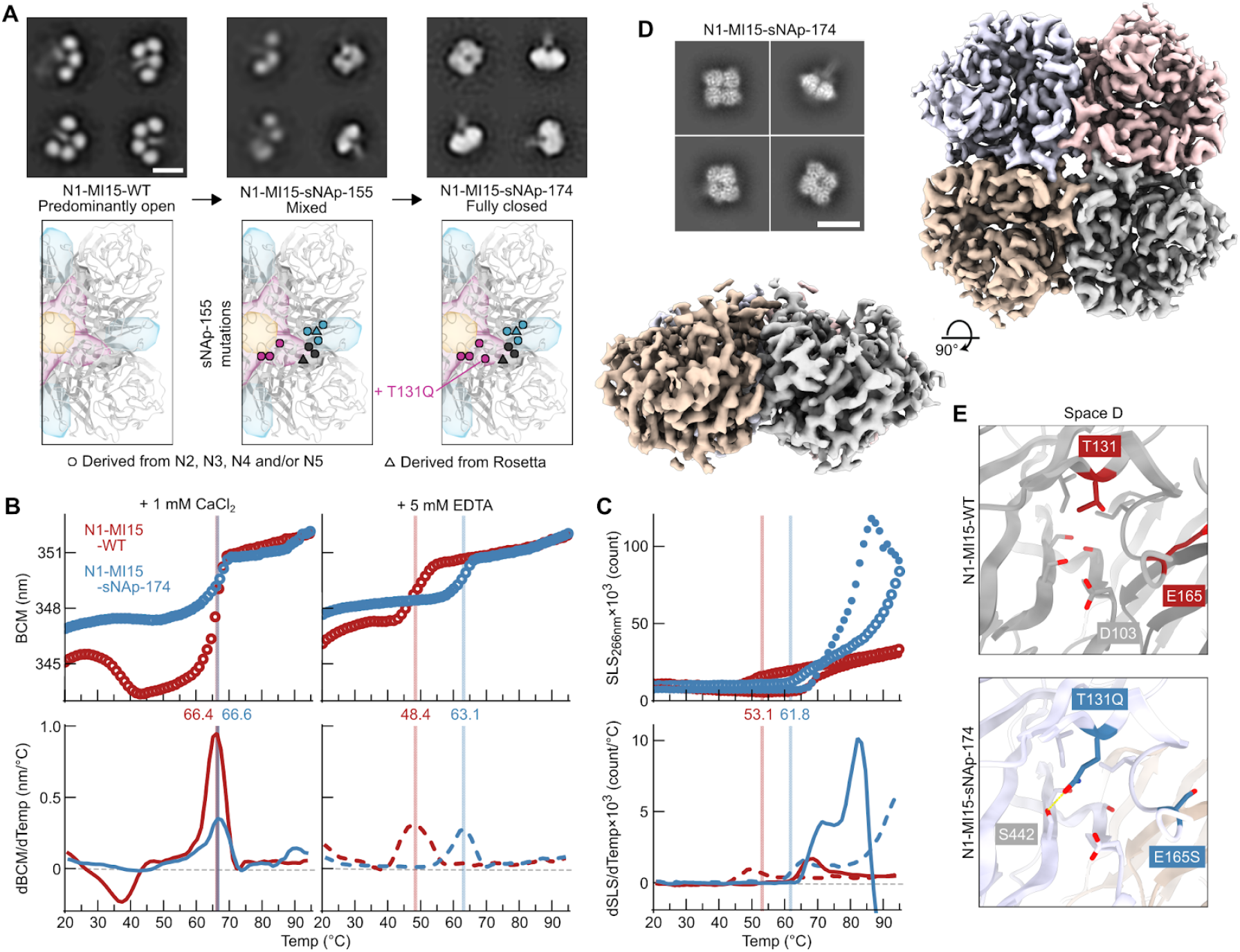
Design and Structural Characterization of N1-MI15-sNAp-174. (A) Application of sNAp-155 mutations to N1 MI15 and addition of the space D stabilizing mutation T131Q resulted in fully closed N1-MI15-sNAp-174 tetramers (scale bar, 10 nm). (B) Thermal denaturation of N1-MI15-WT and N1-MI15-sNAp-174 in the presence of 1 mM CaCl_2_ (closed circles and solid lines) or 5 mM EDTA (open circles and dashed lines), monitored by intrinsic tryptophan fluorescence. Top panels show raw data, while lower panels show smoothed first derivatives used to calculate melting temperatures. The barycentric mean (BCM) of the fluorescence emission spectra is plotted. (C) SLS during thermal denaturation of N1-MI15-WT and N1-MI15-sNAp-174 in the presence of 1 mM CaCl_2_ or 5 mM EDTA, with aggregation temperatures shown for CaCl_2_-treated samples. Data are plotted as in panel B. (D) Representative 2D class averages (scale bar, 10 nm) (top left) and final 3D density (bottom left and right) from cryo-EM of N1-MI15-sNAp-174. (E) Space D cavity in N1-MI15-WT (top; from PDB ID 4B7Q) and N1-MI15-sNAp-174 (bottom; cryo-EM reconstruction). All residues shown as sticks in the WT model are conserved between N1-CA09-WT and N1-MI15-WT.

Intrigued by the possible requirement of an ordered space D for closure, we focused on redesigning the aqueous cavity surrounding D103 to stabilize N1 MI15. This cavity is adjacent to, but not immediately at, the inter-protomer interface. After generating a homology model of N1-MI15 using PDB ID 4B7Q as a template, we applied our Rosetta-based homology-directed design strategy to this space. Among 20 designs building on N1-MI15-sNAp-155, four were found to predominantly favor the closed state, with three showing exclusively closed particles in NS-EM 2D class averages (Fig. 5A and Extended Data Fig. 5A). These designs utilized N2-inspired proline mutations (N1-MI15-sNAp-165 and -176), an N2-inspired T131Q mutation that provides both improved packing and hydrogen bonding in the cavity (N1-MI15-sNAp-174 and -176), and homology-directed hydrophobic filling of the cavity (N1-MI15-sNAp-183). Mirroring the results obtained with other sNAps, the stability of N1-MI15-sNAp-174 (which added only a single amino acid from the sNAp-155 mutations) was relatively insensitive to Ca^2+^ and the T_m_ of the sNAp was 14.7°C higher than N1-MI15-WT in the presence of EDTA (Fig. 5B). The T_agg_ was also higher in both in the presence of EDTA and Ca^2+^ (Fig. 5C). N1-MI15-sNAp-174 also maintained the closed tetrameric state over a period of 15 days (Extended Data Fig. 5B). It remained catalytically active, although at reduced levels compared to its open WT counterpart (Extended Data Fig. 5C). The success of these designs established multiple sets of mutations that stabilize closed N1 NA tetramers—including several that do not make direct contact across the interface—and highlight the cavity in space D as a key determinant of closed tetramer stability.

We determined the structure of the N1-MI15-sNAp-174 tetramer at 3.2 Å resolution using cryo-EM (Fig. 5D and Extended Data Fig. 6). The structure, which strongly resembled the overall structure of N1-CA09-sNAp-155 (backbone RMSD of 0.77 and 0.80 Å over one and all four subunits of the tetramer, respectively), confirmed that N1-MI15-sNAp-174 adopts the closed tetrameric state in solution. The backbone rearrangement in space A involving W455 and W457 observed in the N1-CA09-sNAp-155 structure was also found in N1-MI15-sNAp-174, although W457 was not resolved. The new T131Q mutation helps fill the space D cavity as intended while forming a hydrogen bond to the side chain of S442 (Fig. 5E).

We next applied the sNAp-155 mutations to more distantly related N1 NAs from the avian H5N1 strain A/Vietnam/1203/2004 (VN04) and a pre-pandemic lineage of human H1N1, A/WSN/1933 (WSN33). The resultant N1-VN04-sNAp-155 and N1-WSN33-sNAp-155 formed predominantly open tetramers (Extended Data Fig. 5D,E). We reasoned that making the sequences of these sNAps more like N1-CA09-sNAp-155 and N1-MI15-sNAp-174 overall should stabilize them in the closed tetrameric state. We therefore made substitutions at other buried positions in N1 VN04 and N1 WSN33 that differed from N1 CA09 and N1 MI15, an approach similar to the Repair-and-Stabilize strategy recently developed for stabilization of HIV-1 Env trimers^58^, and combined these with mutations used to stabilize N1 CA09 and N1 MI15. N1-VN04-sNAp-354, which combined the T131Q substitution in space D that stabilized N1-MI15-sNAps with mutation of three nearby residues to the amino acids in N1 CA09 and N1 MI15 (I106V, V163I, and A166V), formed exclusively closed tetramers as determined by NS-EM (Extended Data Fig. 5D).

The sequence of N1 WSN33 NA differs from N1 CA09 and N1 MI15 at many positions in both the space D cavity and the rest of the tetrameric interface. In addition to the sNAp-155 mutation set, substitution of six residues across spaces A, B, and D to the identities of N1 CA09 and N1 MI15 (G105S, I106V, A157T, V163I, A166V, and R210G; N1-WSN33-sNAp-366) substantially improved tetramer closure (Extended Data Fig. 5E), demonstrating that there are inherent differences in interface stability between N1 WSN33 and N1 CA09/N1 MI15 that can be altered by focusing purely on the interface and the space D cavity. The addition of T131Q to this set of mutations resulted in a sNAp (N1-WSN33-sNAp-367) that further improved the tetramer closure, while hydrophobic packing mutations in the cavity (N1-WSN33-sNAp-375) were less effective. To determine if sequence differences distant from the interface contribute to closed tetramer formation, eight buried residues distributed throughout the globular head of N1-WSN-sNAp-375 were further changed to the amino acids in N1 CA09, resulting in the predominantly closed N1-WSN-sNAp-378 (Extended Data Fig. 5E). These data suggest that while spaces A and D are key for closure, packing interactions distant from the interface can allosterically influence interface-stabilizing mutations. Transfer of general packing residues from one strain to another thus offers an additional strategy for stabilization while maintaining native surface antigenicity. Together, the strategies we provide should be widely applicable across many N1 NAs as well as NAs from other subtypes.

### Effects of N1 NA Tetramer Stabilization on Antigenicity and Virus Fitness

To determine whether stabilizing NA tetramers in the closed conformation alters the antigenicity of recombinant NA, we assessed the binding kinetics of mAbs that recognize two antigenic sites on N1 NAs (Fig. 6A). The murine mAb CD6, isolated from a mouse infected with a sublethal dose of A/California/07/2009 (H1N1) virus, recognizes a quaternary epitope that spans two neighboring NA protomers^52^. CD6 is highly protective and potently inhibits viral replication *in vitro*. The second mAb, 1G01, isolated from a human immediately after H3N2 virus infection, is the broadest anti-NA antibody described to date, providing protection against many influenza A and B viruses by targeting the highly conserved catalytic site^36^. We measured antibody binding affinities to the open N1-CA09-WT and closed N1-CA09-sNAp-130 by biolayer interferometry. While both proteins were recognized by 1G01 IgG with similar affinity in the presence of CaCl_2_, the antibody bound to N1-CA09-WT with much lower affinity than to N1-CA09-sNAp-130 in the presence of EDTA (Fig. 6B). Strikingly, the quaternary epitope-specific CD6 IgG minimally bound N1-CA09-WT regardless of the presence of Ca^2+^ ions, indicating that the CD6 epitope was not properly formed in the open N1-CA09-WT. In contrast, the same antibody readily and avidly bound to N1-CA09-sNAp-130 in the presence or absence of Ca^2+^ (Fig. 6B). These data further illuminate the difference in tetrameric conformation between open NAs and sNAps in solution, and demonstrate that tetramer closure better preserves antigenic sites—including a quaternary epitope—targeted by highly protective, infection-elicited antibodies.

**Fig. 6.**
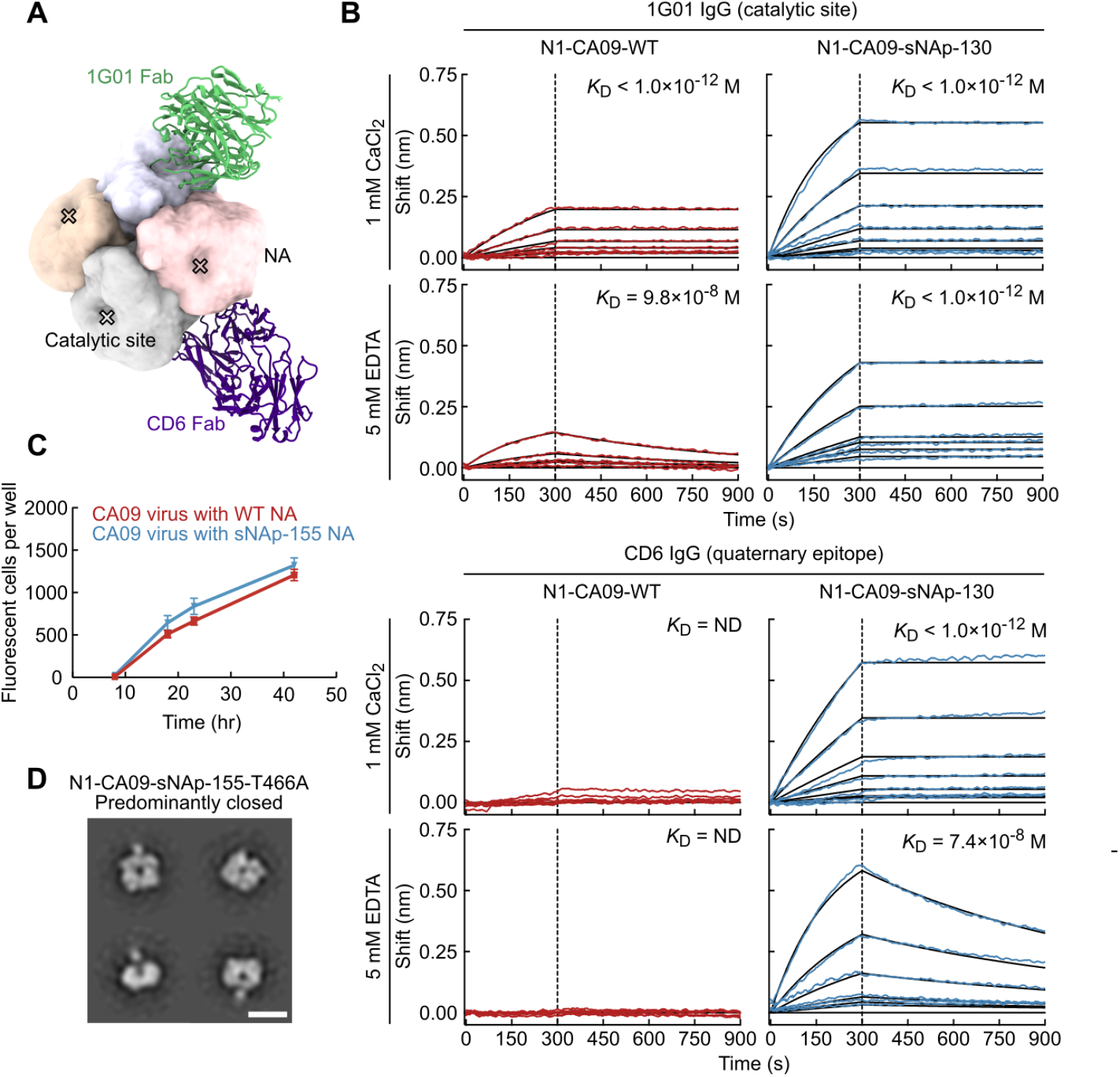
Effects of N1 NA Tetramer Stabilization on Antigenicity and Virus Fitness. (A) Models of Fabs of murine CD6 (purple) and human 1G01 (green) bound to CA09 NA. CD6 binds a quaternary epitope that spans two protomers, while 1G01 binds within and around the catalytic pocket. Models were generated by aligning crystal structures of CD6 (PDB ID 4QNP) and 1G01 (PDB ID 6Q23) Fabs to a single tetramer of CA09 NA (PDB ID 4B7Q). (B) Binding kinetics of anti-NA mAbs to N1-CA09-WT and N1-CA09-sNAp-130. Biolayer interferometry sensorgrams of 1G01 IgG (left) and CD6 IgG (right) binding to N1-CA09-WT (red) and N1-CA09-sNAp-130 (blue). Upper and lower sensorgrams were measured in buffers containing 1 mM CaCl_2_ and 5 mM EDTA, respectively. Experimental data (colored traces) were fitted (black lines) with the binding equations describing a 2:1 interaction. (C) Growth curve of A/California 07/2009 H1N1 influenza reporter viruses with WT NA and NA with sNAP-155 mutations on MDCK-SIAT1-PB1 cells. Error bars denote the standard deviation of measurements from 64 wells at each time point. (D) NS-EM of a recombinant soluble N1-CA09-sNAp-155 with the T466A mutation acquired during growth of the R3ΔPB1 A/California 07/2009 H1N1 sNAp-155 virus (scale bar, 10 nm).

We then tested whether mutations that stabilize the closed tetrameric state impacted viral fitness. We generated CA09 H1N1 reporter influenza viruses^59^ with and without the N1-CA09-sNAp-155 mutations in the NA gene by reverse genetics, and observed similar growth kinetics over 42 hours post infection (Fig. 6C). Sequencing revealed that the virus carrying N1-CA09-sNAp-155 acquired a T466A mutation in the stabilized NA, and both viruses acquired mutations in HA (Extended Data Table 1). We found that recombinant N1-CA09-sNAp-155 with the T466A mutation adopted the closed tetrameric state, confirming that this mutation does not affect the tetramer closure (Fig. 6D). These results demonstrate that tetramer-stabilizing mutations are compatible with virus growth, suggesting that it may be possible to improve the conformational and antigenic stability of NA in inactivated and live-attenuated vaccines produced using traditional manufacturing processes.

## Discussion

Influenza NA has been an attractive yet temperamental target for universal influenza countermeasures. We have shown that recombinant NA tetramers from several different influenza strains and subtypes adopt a previously unreported open tetrameric conformation, while others form the crystallographically observed closed tetramer. These data may help explain reports of differing stabilities among NAs from various subtypes^45^ and are consistent with observations that NA protomers have a tendency to dissociate^23,47^. The open conformation we observed is similar to those found in distantly related tetrameric viral glycoproteins, leaving open the possibility that this conformation has biological relevance^50,51^. Nevertheless, several lines of evidence suggest that NA forms closed tetramers on the surface of influenza virions, and our work highlights the importance of the closed state for proper stability and binding to protective infection-elicited antibodies. Such differences between native transmembrane and recombinant soluble forms of glycoproteins are not uncommon. For example, many class I fusion proteins require stabilizing mutations to maintain their native—and often metastable—prefusion conformations when produced recombinantly^60^. Further study will be required to more precisely define the native tetrameric conformation(s) of NA on the surface of diverse influenza virus strains and subtypes.

Inspired by knowledge of the sequences and structural phenotypes of homologous NAs, we stabilized the closed tetrameric state using a homology-directed design strategy similar to the recently reported Repair-and-Stabilize approach applied to HIV envelope trimers^58,61^. We identified several structural features that work together to stabilize the closed NA tetramer: direct interactions between subunits at the tetrameric interface, pre-organization of the tetrameric interface within each subunit by second-shell interactions, and general stabilization of the structure of each subunit by packing interactions distant from the interface. Two regions of the NA, which we refer to as spaces A and D, appear to have a particularly strong influence on NA stability and conformation. Multiple naturally occuring stabilizing and destabilizing mutations in space D have been described previously^10,46^, and comparisons to homologous NAs led to a single stabilizing mutation in space C in an N9 NA which facilitated structure determination of NA–antibody complexes^47^. Whereas mutations in space A and the space D cavity stabilize the closed tetrameric state by improving atomic packing in these interface or interface-proximal regions, the mechanism of other mutations is less clear. For example, there is no obvious structural rationale for the stabilizing effect of the E/Q165S mutation, yet this position—which stood out when sequence alignments of several NA subtypes were compared—is a critical determinant of tetramer closure. The success of homology-based mutations reflects the sensitivity and complexity of the determinants of NA stability and provides a roadmap towards the design of additional stabilizing mutations to NAs from diverse influenza strains and subtypes.

Our findings have several implications for previous and future studies of influenza NA. We found that several commonly used assays—SEC, enzymatic activity, and thermal stability—were generally unable to distinguish between the open and closed forms of the enzyme due to inherent differences between NAs from different strains. Structural characterization by EM was required to establish whether each NA adopted the open or closed tetrameric state, or a mixture of the two. Antigenic characterization using protective mAbs that bind a quaternary epitope or the catalytic site also distinguished the open and closed forms. We anticipate that with the identification of additional NA-specific mAbs with the desired properties, antigenic characterization could become a rapid, widely accessible, and powerful tool for assessing NA tetrameric conformation, which could be vital for vaccine manufacturing.

Furthermore, our data suggest that the recombinant NA proteins used in many previous studies, while presumed to be in the crystallographically observed closed tetrameric state based on biochemical assays, may have been partially or wholly in the open conformation. This disconnect could lead to erroneous interpretations of experimental outcomes, most particularly for serological measurements, and could limit discovery of valuable mAbs. If so, our understanding of NA antigenicity and immunogenicity may be incomplete or compromised, particularly for NAs from the highly relevant but less stable N1 subtype.

The sNAps and design methods described here should enable new approaches to structure-based design of NA antigens to include in future influenza vaccines. As highlighted by the transformational effects of the development of prefusion-stabilized class I viral fusion proteins^60,63–65^, the availability of stable, native-like recombinant glycoproteins can broadly catalyze research and development efforts. First, sNAp-based probes may improve serological assays for NA-specific antibodies elicited by infection or immunization, as well as the isolation of NA-specific B cells and mAbs. Their enhanced stability and monodispersity could expedite rapid structure determination of antibody-NA complexes, which will provide valuable information on mechanisms of antibody-mediated protection. More directly, sNAps may themselves become superior NA-based immunogens. We note that recombinant wild-type N1 NA heads fused to hVASP tetramerization domains, which we predict adopt the open conformation, have previously been shown to elicit protective immunity against lethal virus challenge in mice, suggesting that the less stable open state can be useful as an immunogen^28,44^. The increased affinity of an N1 sNAp towards the quaternary epitope-specific mAb CD6 and the broadly protective mAb 1G01 suggests that sNAp-based immunogens may be more likely to elicit antibody responses targeting these sites of vulnerability and other quaternary epitopes. Finally, the increased physical stability of sNAps and their compatibility with live influenza virus application may help overcome product stability issues that have historically plagued both virus-derived and recombinant NAs, enabling the design and evaluation of next-generation NA-based influenza vaccines.

## Supporting information

Supplemental Item 1

Supplemental Item 2

Supplemental Item 3

## Acknowledgments

We would like to thank J. Boyington and G. Stewart-Jones (VRC) for initial design insights; J. Mascola (VRC) for advisory support; and H. Kleanthous (Bill & Melinda Gates Foundation), members of the King laboratory, and members of the Influenza Program at the VRC for comments on the manuscript. This study was supported by the intramural research program of the Vaccine Research Center, National Institute of Allergy and Infectious Diseases, National Institutes of Health (M.K., B.S.G.); a generous gift from the Open Philanthropy Project (N.P.K.); a generous gift from the Audacious Project (N.P.K.); the National Institute of General Medical Sciences (R01GM120553, D.V.); the National Institute of Allergy and Infectious Diseases (DP1AI158186 and HHSN272201700059C, D.V.); a Pew Biomedical Scholars Award (D.V.); an Investigators in the Pathogenesis of Infectious Disease Award from the Burroughs Wellcome Fund (D.V.); and the University of Washington Arnold and Mabel Beckman cryo-EM center. This study has been funded in part with Federal funds from the Frederick National Laboratory for Cancer Research, National Institutes of Health, under Contract No. HHSN261200800001E (Y.T., T.S.). Molecular graphics and analyses were performed with UCSF ChimeraX, developed by the Resource for Biocomputing, Visualization, and Informatics at the University of California, San Francisco, with support from National Institutes of Health R01GM129325 and the Office of Cyber Infrastructure and Computational Biology, National Institute of Allergy and Infectious Diseases.

## Author contributions

Conceptualization: M.K.; Modeling and design: D.E., J.L.; Formal Analysis: D.E., J.L., O.A., Y.T., S.K., C.Y., R.A.G., A.C., T.S., D.P., M.M., A.J.B., K.K.L., D.V., N.P.K., M.K.; Investigation: D.E., J.L., O.A., Y.T., S.K., C.Y., R.A.G., A.C., T.S., D.P., M.M., A.J.B., K.K.L., D.V., N.P.K., M.K.; Resources: C.Y., D.P., M.M.; Writing – Original Draft: D.E., J.L., O.A., S.K., N.P.K., M.K.; Writing – Review & Editing: All authors; Visualization: D.E., J.L., O.A., Y.T., S.K., T.S., A.B., N.P.K., M.K.; Supervision: K.K.L., B.S.G., D.V., N.P.K., M.K.; Funding Acquisition: B.S.G., D.V., N.P.K.

## Competing interests

D.E., J.L., B.S.G., N.P.K., and M.K. are named as inventors on patent applications filed by the University of Washington based on the studies presented in this paper. N.P.K. is a co-founder, shareholder, paid consultant, and chair of the scientific advisory board of Icosavax, Inc., and has received unrelated sponsored research agreements from Pfizer. D.V. is a consultant for and has received an unrelated sponsored research agreement from Vir Biotechnology, Inc.

**Correspondence and requests for materials** should be addressed to N.P.K., and M.K.

## Methods

### Cell Lines

Expi293F cells, a female human embryonic kidney cell line adapted to grow in suspension, were obtained from Thermo Fisher Scientific.

### Protein expression and purification

Sequences for all NA proteins are provided in Supplementary Item 1. The cytoplasmic, transmembrane, and stalk domains of the wild-type (WT) NA (residues 1–82, N1 numbering) were replaced by affinity purification tags, a tetramerization domain derived from human vasodilator-stimulated phosphoprotein (hVASP)^49^, a thrombin protease cleavage site, and a two-residue GG linker. NA constructs were expressed by transient transfection in Expi293F cells (ThermoFisher Scientific) at a density of 2.5×10^6^ cells/ml using the ExpiFectamine™ 293 Transfection Kit (ThermoFisher Scientific). The supernatants were harvested 5 days post-transfection and centrifuged at 4000 rpm to remove cell debris. Proteins were purified from clarified supernatants by immobilized metal affinity chromatography (IMAC) using either Ni^2+^- or Co^2+^-containing resin. For Ni^2+^-based IMAC, clarified supernatants were incubated for 2 h at room temperature with Ni^2+^ Sepharose High Performance histidine-tagged protein purification resin (Cytiva) and bound protein eluted using 50 mM Tris pH 8.0, 0.5 M NaCl, 300 mM imidazole. For Co^2+^-based resin, clarified supernatants were flowed over Talon resin (Takara), and bound protein eluted in 20 mM Tris pH 8.0, 300 mM NaCl, 300 mM imidazole. Eluted proteins were further purified by SEC into phosphate-buffered saline (PBS) or 25 mM Tris pH 8.0, 150 mM NaCl, 5% glycerol using a Superdex 200 Increase 10/300 column (Cytiva). Recombinant NA proteins derived from A/California/7/2009 (H1N1) (Catalog #NR-19234), A/New Caledonia/20/1999 (Catalog #NR-43779) and A/Wisconsin/67/2005 (Catalog #NR19237) were obtained from BEI Resources

### Negative stain EM sample preparation and analysis

NS-EM and particle image averaging was used to assess whether the head domains of recombinant NA proteins adopted the open or closed tetrameric structure. Proteins were diluted to between 0.1–0.2 mg/mL using either 10 mM HEPES pH 7.0, 150 mM NaCl or 10 mM Tris pH 7.5, 150 mM NaCl. Samples were adsorbed to glow-discharged carbon-coated copper grids. The grids were either washed with a drop of the same buffer three times and stained with 0.75% uranyl formate, or blotted and stained directly with 0.75% uranyl formate. Images were recorded with sampling ranging between 1.9 Å/pixel and 2.2 Å/pixel, depending on the microscope. Data were collected on either an FEI Tecnai T20 electron microscope equipped with an FEI Eagle CCD camera and operated at 200 kV using SerialEM^66^, or on an FEI Tecnai 12 Spirit 120kV electron microscope equipped with a Gatan Ultrascan 4000 CCD camera. Particles were selected from the micrographs automatically using either in-house software (Yaroslav Tsybovsky, unpublished) or were picked in a reference-free manner in cisTEM^67^. For the latter datasets, particles were extracted after correcting for the effect of the CTF for each micrograph with cisTEM^67^. Depending on user, dataset, and microscope, particles were either extracted into 120×120-pixel boxes with a final pixel size of 2.2 Å/pixel, or extracted into 230×230-pixel boxes with a final pixel size of 2.2 Å/pixel, or extracted with a box size of 176×176 pixels and binned to a final box size of 44×44 pixels (to a pixel size of 6.4 Å/pixel). Reference-free 2D classification was performed using either Relion 1.4 or Relion 3.1 (refs. 68,69). Representative 2D class averages for all datasets were chosen based on the unambiguous identification of NA particle quality and conformational state, with qualitative descriptions based on overall class average appearance and/or semi-quantitative measurements of particles assigned to each class when distinct populations could be reliably measured. A designation of “fully closed” was assigned to samples where only closed particles were resolved into interpretable 2D class averages. “Predominantly closed” NA variants were assigned to samples with 2D class averages of the closed state comprising >80% of well-resolved NA tetramers, with open or poorly resolvable classes containing the remaining <20% of particles. “Mostly closed” was assigned to NA variants where roughly three quarters of particles were classified into the closed conformation. “Mostly open” was assigned when roughly one quarter of particles were observed as closed. “Predominantly open” was assigned for samples with clearly >80% of resolvable particles being classified in an open conformation. “Mixed” was assigned to any sample in between “mostly closed” and “mostly open”. Representative electron micrographs and complete sets of 2D class averages are provided in Supplementary Item 2 for each NA.

### Thermal denaturation and static light scattering

Non-equilibrium melting temperatures were determined using an UNcle (UNchained Labs) based on the barycentric mean of intrinsic tryptophan fluorescence emission spectra collected from 20–95°C using a thermal ramp of 1°C per minute in buffers of 25 mM Tris pH 8.0, 150 mM NaCl and 5% glycerol supplemented with either 1 mM CaCl_2_ or 5 mM EDTA. Static light scattering (SLS) data was simultaneously collected. Melting temperatures were defined as the maximum point of the first derivative of the melting curve, with first derivatives calculated using GraphPad Prism software after smoothing with four neighboring points using 2nd order polynomial settings. Aggregation temperatures were defined as the first positive data point above baseline of the first derivative of SLS, with the first derivative similarly calculated using GraphPad Prism.

### Enzymatic activity

Neuraminidase activity was measured with the NA-Fluor™ Influenza Neuraminidase Assay Kit (ThermoFisher Scientific) according to the manufacturer’s protocol. Briefly, two-fold serial dilutions of each NA protein were made in a black 96-well, flat bottom plate, starting from protein stocks at concentrations of 25–50 µg/mL. The wells in column 12 were left empty for background controls. NA-Fluor Substrate was prepared according to the manufacturer’s protocol and added to each well. Plates were incubated for 1 h at 37°C and reactions were stopped with NA-Fluor Stop Solution. Plates were read using an excitation wavelength range of 350–365 nm and an emission wavelength range of 440–460 nm. Background control wells were subtracted for each protein serial dilution. Protein concentrations were plotted versus relative fluorescence unit (RFU) values.

### Structure-based design using Rosetta

All calculations in Rosetta were made using versions v2017.18-dev59451, v2019.21-dev60746, or v2019.45-dev61026. For design of N1 CA09 sNAps, residue positions were manually sorted into spaces A, B, C, and D based on perceived local interactions in the inter-protomeric interface of N1-CA09-WT (PDB ID 4B7Q). The sequence of PDB ID 4B7Q contains an additional Y351F mutation compared to the sequence used for N1-CA09-WT, which was maintained in all designs. Based on either amino acid appearing at this position in multiple closed and open constructs described in this manuscript, we do not expect that this mutation has an impact on closure. Representative structures of N2 (PDB ID 6BR5), N3 (PDB ID 4HZV), N4 (PDB ID 2HTV), and N5 (PDB ID 3TI8) NAs were analyzed and residue identities at each of the positions were recorded. The four-fold symmetry axis of N1-CA09-WT was aligned with [0,0,1] and a single protomer was saved in .pdb format. A design protocol was written using RosettaScripts^53^ that takes the aligned protomer and a custom resfile as inputs, with the resfile dictating the side chain identities and conformations sampled during design (Supplementary Item 3). Briefly, the protocol applies two rounds of design based on the input resfile, with side chain and backbone energy minimization applied after each design step. Both design and minimization steps were allowed to repack or minimize residues within 5 Å of all mutable or packable residues listed in the resfile. Multiple resfiles were set up for each space. One set of resfiles was designed to place specific substitutions from N2 (PDB ID 4H52), N3, N4, or N5 NA structures into N1-CA09-WT using the ‘PIKAA’ option. Other sets used the ‘PIKAA’ option to allow Rosetta’s packer to choose between residue identities from the N1, N2, N3, N4, or N5 NAs. A final set used the ‘PIKAA’ option to add further residue identities that were not observed in the N1, N2, N3, N4, or N5 structures to identify novel mutations. Design models and scores were manually inspected to identify interactions across the interface that appeared structurally feasible. Mutations were discarded if they buried polar groups that were natively solvent-exposed or involved in hydrogen bonds. To prevent undesired alterations to antigenicity, mutations to surface-exposed residues were not frequently considered. Favorable interactions were iteratively retested in resfiles and manually refined to finalize a diverse set of designs for each space. Twenty-six designs targeted individual spaces, while six designs consisted of combinations of designs to all four spaces that were manually picked and combined.

A slightly modified protocol was used to design N1 MI15 sNAps that contained additional mutations in space D. Due to the lack of a crystal structure for N1 MI15, N1 CA09 (PDB-ID 4B7Q) was used as a template for a homology model. All design trajectories were performed while adding sNAp-155 mutations (I99P, Y100L, C161V, E165S, S172A, V177I, S196T, V205I, Q408M, R419V) in addition to mutational differences between N1 CA09 and N1 MI15 (N200S, V241I, N248D, V264I, N270K, I314M, I321V, N369K, N386K, K432E). Beyond those mutations, the design process focused entirely on residues within space D with resfile inputs drawn from either crystal structures of other non-bat NA subtypes or other identities not naturally observed in all subtypes. Disulfide mutations were designed outside of Rosetta based on distances between pairs of residues.

### Hydrogen-deuterium exchange mass spectrometry

For each time point, 40 pmol of N1-CA09-WT and N1-CA09-sNAp-130 were incubated in deuterated buffer (85% D_2_O, pH* 7.4) for 3, 60, 1,800, or 72,000 s at room temperature and subsequently mixed with an equal volume of ice-cold quench buffer (4 M urea, 200 mM tris(2-chlorethyl) phosphate (TCEP), 0.2% formic acid) to a final pH* of 2.5. Samples were immediately frozen in liquid nitrogen and stored at 80°C until analysis. Fully deuterated samples were prepared by digesting 40 pmol of undeuterated sample over a pepsin column, followed by concentration under vacuum, resuspension in deuterated buffer at 65°C for 1 hour, and then quenching/freezing. Zero time point samples were prepared as previously described^70^. A 3 s exchange was performed at the beginning and end of the longest exchange reaction to monitor protein stability under the experiment conditions. Online pepsin digestion was performed and analyzed by LC-MS-IMS utilizing a Waters Synapt G2-Si Q-TOF mass spectrometer as previously described^70^. Deuterium uptake analysis was performed using HX-Express v2 (refs. 71,72). The relative deuterium exchange was corrected for in-exchange using the zero time point. Internal exchange standards (Pro-Pro-Pro-Ile [PPPI] and Pro-Pro-Pro-Phe [PPPF]) were included in each reaction to ensure that conditions were consistent throughout all of the labeling reactions.

### Cryo-electron microscopy sample preparation, data collection, and image processing

sNAp proteins were diluted to 1–1.5 μM in buffer (10 mM Tris, pH 7.5, 150 mM NaCl) and 3 μL sample loaded onto a freshly glow-discharged 1.2/1.2 UltrAuFoil grid (300 mesh) prior to plunge freezing using a vitrobot Mark IV (ThermoFisher Scientific) with a blot force of -1 and 3.5–4.5 s blot time at 100% humidity and 4°C.

Data were acquired on an FEI Glacios transmission electron microscope operated at 200 kV and equipped with a Gatan K2 Summit direct detector. Automated data collection was carried out using Leginon^73^ at a nominal magnification of 36,000× with a pixel size of 1.16 Å. The dose rate was adjusted to 8 counts/pixel/s, and each movie was acquired in counting mode fractionated in 50 frames of 200 ms. For the N1-CA09-sNAp-c155 and N1-MI15-sNAp-174 complexes, 1,034 and 1,099 micrographs were collected, respectively, with a defocus range between -0.5 and -2.5 µm. For the N1-CA09-WT complex, 605 micrographs were collected with a defocus range between -1.0 and -3.0 µm. Movie frame alignment, estimation of microscope CTF parameters, and automatic particle picking and extraction were carried out using Warp^74^. Particles were extracted unbinned into a box size of 204 pixels.

For the N1-CA09-WT dataset, three rounds of reference-free 2D classification were performed using cryoSPARC^75^. After each round, well-defined images corresponding to NA tetramers were selected and re-classified until 2D classification stabilized. For the N1-CA09-sNAp-c155 and N1-MI15-sNAp-c174 datasets, two rounds of reference-free 2D classification were performed in cryoSPARC, again selecting for well defined images. Particles selected post-2D classification were subjected to two rounds of 3D classification in Relion^76^ without imposing symmetry (angular sampling 7.5° for 25 iterations followed by 1.8° with local searches for a further 25 iterations). For each dataset, ab initio models were generated in cryoSPARC as reference maps for 3D classification^75^. Refinements of 3D maps were carried out using non-uniform refinement along with per-particle defocus refinement as implemented in cryoSPARC^77^. Particles were transferred back into Relion to perform Bayesian polishing^68,78^ before an additional round of non-uniform refinement, per-particle defocus refinement, and finally one more round of non-uniform refinement imposing four-fold symmetry. Local resolution estimation, filtering, and sharpening were carried out in cryoSPARC^79^. All reported resolutions were determined using gold-standard Fourier shell correlation (FSC) calculations with a cutoff criterion of 0.143 (ref. 80) and FSC curves were corrected for the effects of soft masking by high-resolution noise substitution^81^.

### Cryo-EM model building and analysis

UCSF Chimera^82^ and Coot^83^ were used to fit atomic models (PDB ID 4B7Q) into the cryoEM maps and point mutations were made manually in Coot. Models were refined and relaxed using Rosetta using both sharpened and unsharpened maps^84,85^ and validated using Molprobity^86^, Phenix^87^, and EMRinger^88^. Figures were generated using UCSF ChimeraX^89^.

### Collection of NA sequences and entropy calculations

Comprehensive sequence datasets for each NA subtype were downloaded from GISAID (www.gisaid.org). Each dataset included animal and human sequences longer than 1,350 nucleotides deposited before January 14, 2020. Sequences were aligned using Mafft v7 for large numbers of short sequences^90^. Sequences with more than 135 ambiguous bases or large gaps were eliminated. To calculate amino acid frequencies, we truncated NA sequences to retain only the globular head of NA ectodomain. Unique amino acid sequences of the NA head domain were selected using CD-HIT with the “Sequence identity cut-off” set at 1.0 (ref. 91). Sequence logos were generated using WebLogo 3 (ref. 92), http://weblogo.threeplusone.com/). From aligned sequences containing gaps, entropy calculations were measured using the dms_tools2 python library (https://jbloomlab.github.io/dms_tools2/). Representative positions for maximum and mean entropies were the individual positions with entropy values that were highest or closest to the mean calculated using all positions in a given sequence, respectively.

### Biolayer interferometry

All biosensors were hydrated in PBS prior to use. Recombinant NA tetramers were immobilized on HIS2 biosensors (fortéBio) through their hexahistidine tags. After briefly dipping in assay buffer (25 mM Tris pH 8, 150 mM NaCl, 5% glycerol, 1% BSA) supplemented with either 1 mM CaCl_2_ or 5 mM EDTA, the biosensors were dipped in a two-fold dilution series of IgG for 5 min. Biosensors were then dipped in the assay buffer (with either CaCl_2_ or EDTA) to allow IgG to dissociate from NA for 10 min. All assay steps were performed at 30°C with agitation set at 1,000 rpm in the Octet HTX instrument (fortéBio). Correction to subtract non-specific baseline drift was carried out by subtracting the measurements recorded for a sensor loaded with the NA in the same buffer with no antibody. Data analysis and curve fitting were carried out using Octet analysis software (version 11). Experimental data were fitted with the binding equations describing a 2:1 (bivalent binding) interaction. Global analyses of the complete data sets assuming binding was reversible (full dissociation) were carried out using nonlinear least-squares fitting allowing a single set of binding parameters to be obtained simultaneously for all concentrations used in each experiment.

### Virus generation

A/California 07/2009 H1N1 influenza reporter virus was rescued by reverse genetics of eight-plasmid system with HA and NA segments of CA09 and internal gene segments of A/WSN/1933 except for PB1 segment which was replaced with reporter gene (tdKatushka2). To generate the virus with stabilized NA, NA segment was modified to incorporate sNAP-155 mutations and used the modified NA segment for virus rescue. Both viruses with WT NA and with stabilized NA were propagated in MDCK-SIAT1 cells constitutively expressing PB1 as described^59^. Both viruses were passaged ten times prior to assessing growth kinetics. For virus growth kinetics, pretitrated viruses were added to MDCK-SIAT1-PB1 cells (preseeded at 1 × 10^5^ cells/ml) in OptiMEM with 1 µg/ml of TPCK-treated trypsin in a 384-well black plate (final 50 µl/well) with transparent bottom (Greiner). Viral growth was measured at 8, 18, 21, 23, and 42 hours post infection by counting fluorescent foci using Celigo Image Cytometer (Nexcelom) with a customized red channel for optimal detection of the tdKatushka2 reporter. In Celigo operation and analysis software v4.1, Target 1 protocol was used to detect and count fluorescent foci.

### Data availability

All images and data were generated and analyzed by the authors, and will be made available by the corresponding authors (N.P.K., and M.K.) upon reasonable request. Structural models and density maps wili be deposited in the Protein Data Bank and Electron Microscopy Data Bank under accession numbers PDB xxxx and EMD-xxxxx.

