## Extended Data for "Structure-based design of stabilized recombinant influenza neuraminidase tetramers"

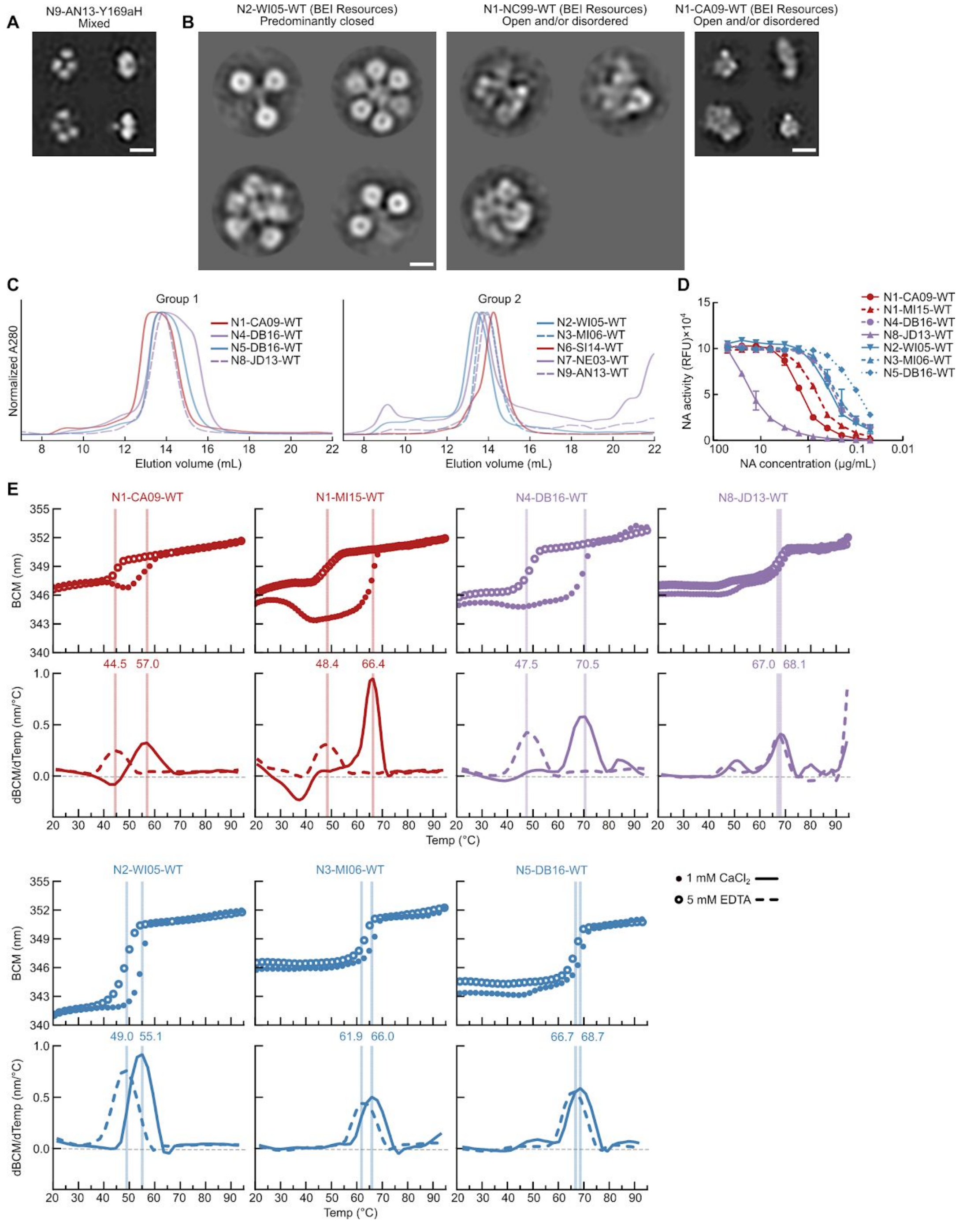


**Extended Data Fig. 1 | NS-EM, Preparative SEC, Enzymatic Activity, and nanoDSF of Recombinant WT and Y169aH-containing NA Tetramers**

(A) NS-EM 2D class averages of recombinant N9-AN13 NA containing the previously identified Y169aH stabilizing mutation (scale bar, 10 nm). (B) NS-EM 2D class averages of recombinant N2-WI05-WT, N1-NC99-WT, and N1-CA09-WT NAs obtained from BEI Resources (scale bar, 10 nm). (C) Preparative SEC of recombinant NAs from all nine non-bat influenza A subtypes on a Superdex 200 Increase 10/300 GL column. (D) Enzymatic activity of purified recombinant NAs. RFU, relative fluorescence units. Error bars denote the standard deviation for each dilution, performed in duplicate. (E) Thermal denaturation of WT NA tetramers in the presence of 1 mM $CaCl_2$ (closed circles and solid lines) or 5 mM EDTA (open circles and dashed lines), monitored by intrinsic tryptophan fluorescence. The barycentric mean (BCM) of the fluorescence emission spectra is plotted. Top panels show raw data, while lower panels show smoothed first derivatives used to calculate melting temperatures, which are indicated by vertical lines.

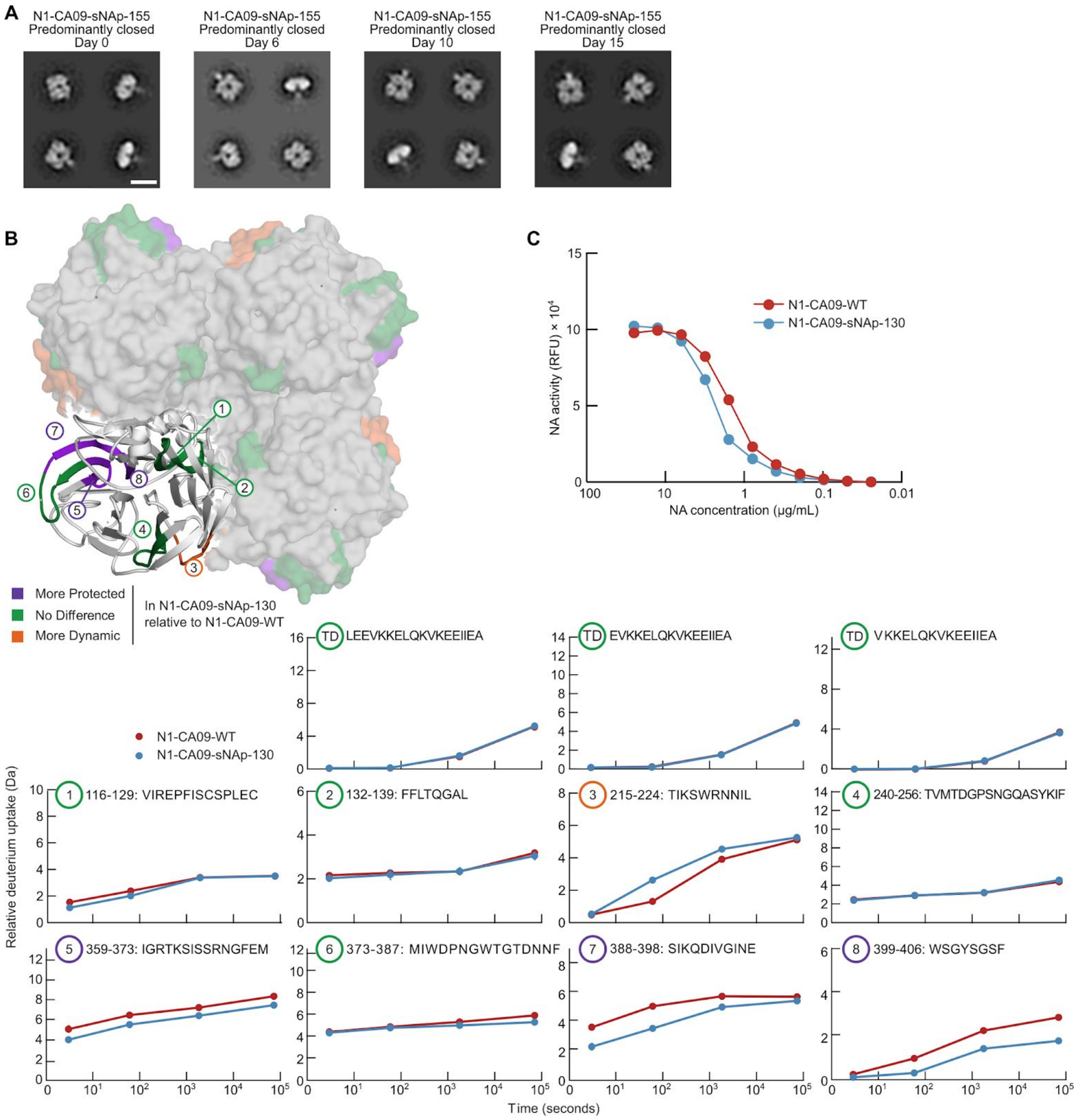


**Extended Data Fig. 2 | Shelf-life Stability, Hydrogen-Deuterium Exchange, and Catalytic Activity of Unmodified and Stabilized N1 CA09 NA tetramers**

(A) Shelf-life stability of N1-CA09-sNAp-155 over 15 days at 4°C as assessed by NS-EM (scale bar, 10 nm). (B) (Top left) Peptide segments, numbered 1 to 8, mapped onto the structure of N1 NA (PDB ID 4B7Q). Colors indicate whether the peptide is more protected (purple) or more dynamic (orange) in N1-CA09-sNAp-130 compared to N1-CA09-WT. Green indicates peptides where there is no difference in deuterium exchange between variants. (Bottom) Kinetics of hydrogen-deuterium exchange for numbered peptides and peptides from the tetramerization domain (TD) at multiple timepoints up to 20 h. Each point is an average of duplicate measurements (N1-CA09-WT) and triplicate measurements (N1-CA09-sNAp-130), except for a limited number of replicates that were discarded due to low signal to noise. Standard deviations for all measurements were smaller than the points plotted. (C) Enzymatic activity of N1-CA09-WT and N1-CA09-sNAp-130. RFU, relative fluorescence units. Performed in duplicate, with error bars smaller than the plotted points and not shown.

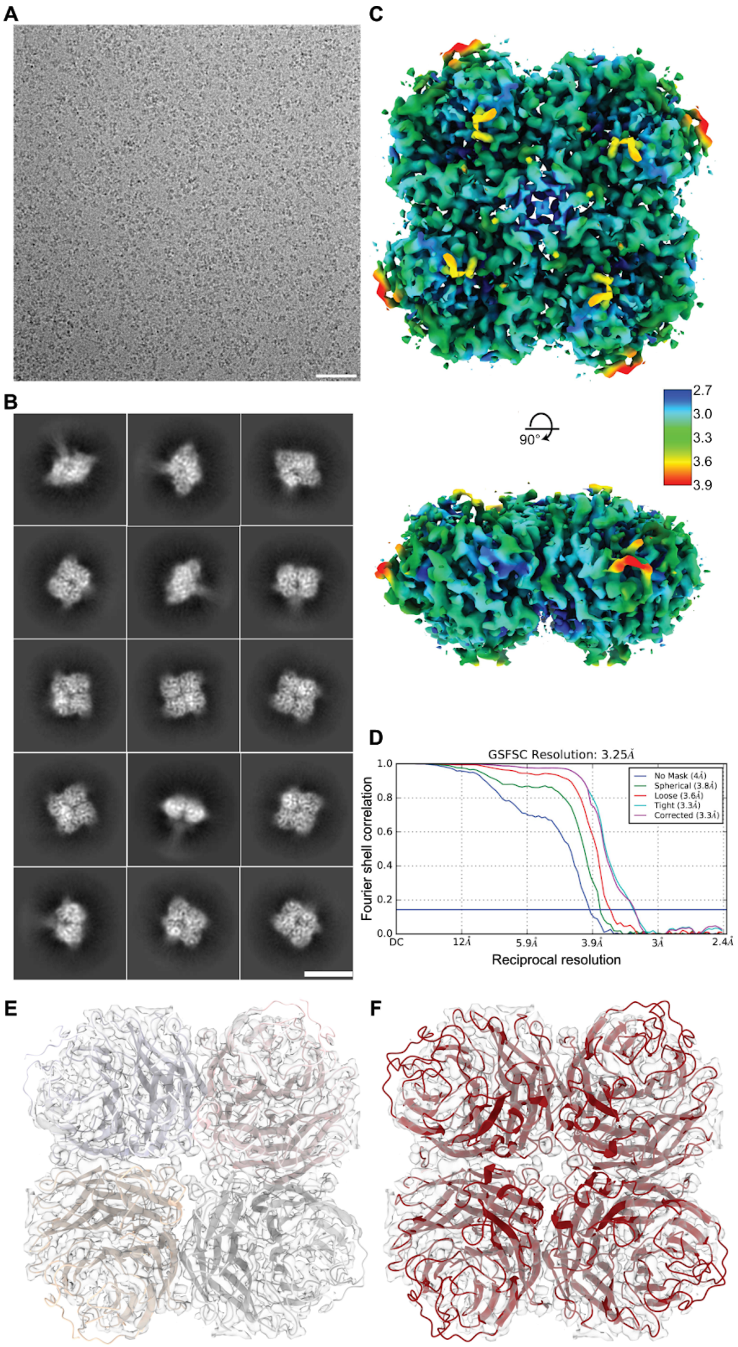


**Extended Data Fig. 3 | Cryo-EM analysis of N1-CA09-sNAp-155**

(A) Representative micrograph of N1-CA09-sNAp-155. Scale bar, 50 nm. (B) 2D class averages of N1-CA09-sNAp-155. Scale bar, 10 nm. (C) Orthogonal views of the cryo-EM reconstruction of N1-CA09-sNAp-155. Map is colored according to local resolution as indicated by color scale (in Å). (D) Gold standard FSC for cryo-EM reconstruction. (E) N1-CA09-sNAp-155 model coloured by chain shown in transparent density showing overall tetramer conformation of map. (F) N1-CA09-WT (4B7Q, red) rigid-body docked into the N1-CA09-sNAp-155 map, showing overall agreement with tetramers observed by X-ray crystallography.

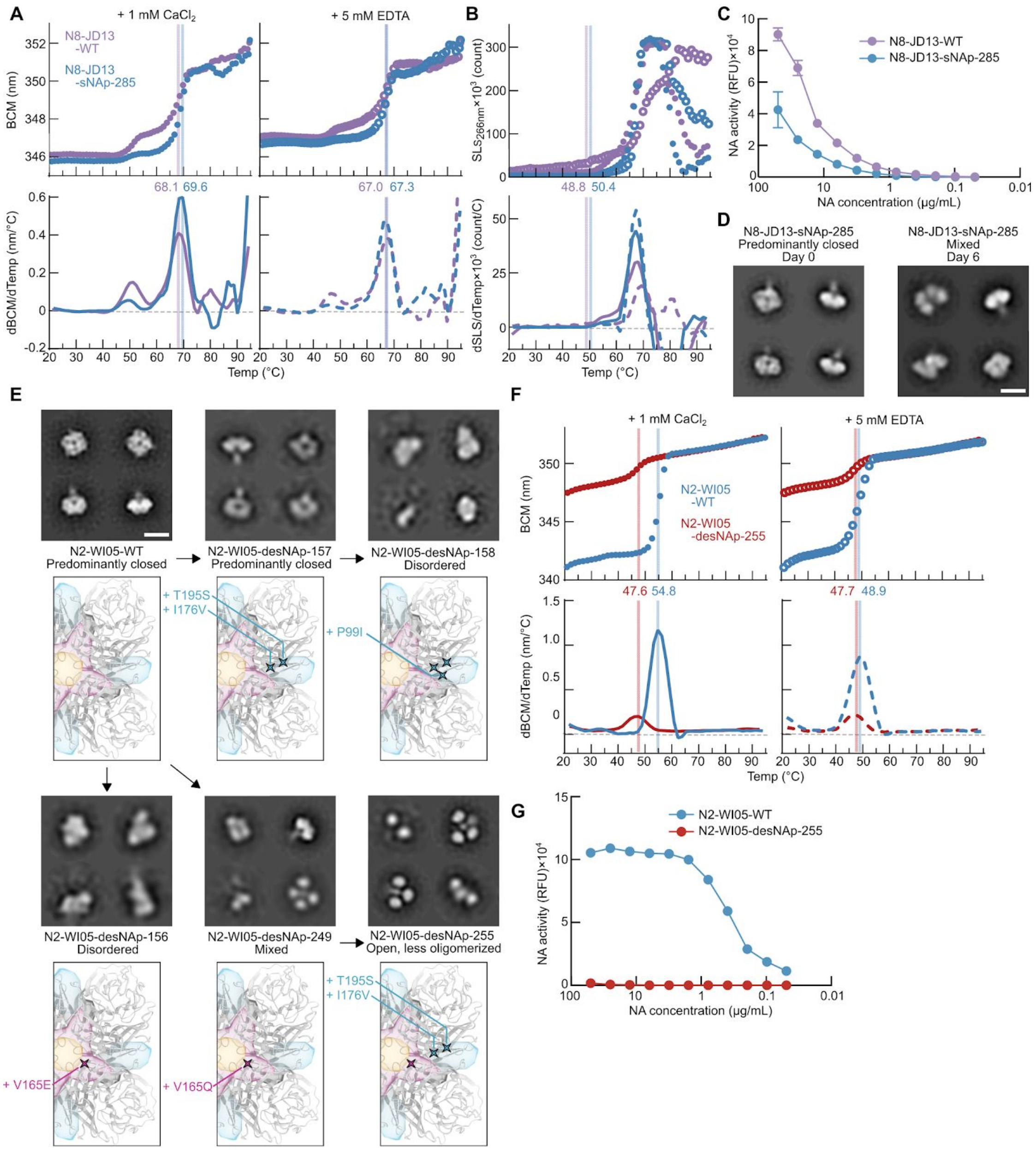


**Extended Data Fig. 4 | Structural, Biophysical, and Biochemical Characterization of a Stabilized N8 NA Tetramer and Destabilized N2 NA Tetramers**

(A) Thermal denaturation of N8-JD18-WT and N8-JD13-sNAp-285 in the presence of 1 mM $CaCl_2$ (closed circles and solid lines) or 5 mM EDTA (open circles and dashed lines), monitored by intrinsic tryptophan fluorescence. Data are presented as in Fig. 2D. (B) SLS during thermal denaturation of N8-JD18-WT and N8-JD13-sNAp-285 in the presence of 1 mM $CaCl_2$ or 5 mM EDTA. Data are presented as in Fig. 2E. (C) Enzymatic activity of N8-JD13-WT and N8-JD13-sNAp-285. RFU, relative fluorescence units. Error bars denote the standard deviation for each dilution, performed in duplicate. (D) Shelf-life stability of N8-JD13-sNAp-285 over 6 days at 4°C as assessed by NS-EM (scale bar, 10 nm). (E) Mutation of predicted stabilizing residues in N2 NA to less stable counterparts observed in N1 or N8 and corresponding NS-EM 2D class averages. All numbers are listed in N2 numbering in which positions 99, 165, 176 and 195 are equivalent to positions 99, 165, 177 and 196 in N1 numbering, respectively (scale bar, 10 nm). (F) Thermal denaturation of N2-WI05-WT and N2-WI05-desNAp-255 in the presence of 1 mM $CaCl_2$ or 5 mM EDTA, monitored by intrinsic tryptophan fluorescence. Data are presented as in Fig. 2D. (G) Enzymatic activity of N2-WI05-WT and N2-WI05-desNAp-255. RFU, relative fluorescence units. Error bars are smaller than the plotted points and not shown.

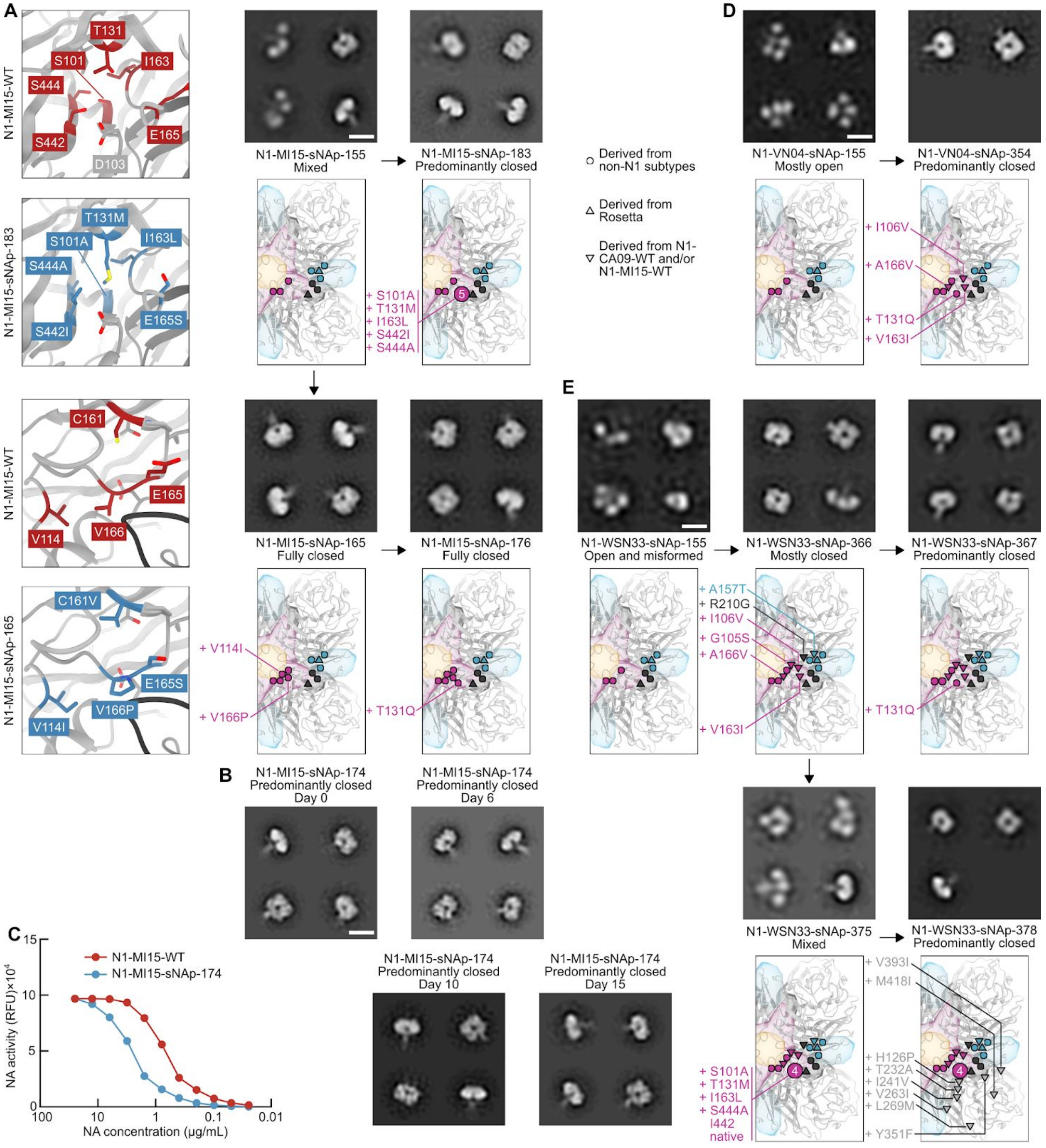

**Extended Data Fig. 5 | Design of Additional Closed Recombinant N1 NA Tetramers**

(A) Multiple mutations used to successfully design sNAps for N1 MI15. Structural models were derived from a Rosetta-generated homology model of N1 MI15 NA based on PDB ID 47BQ (scale bar, 10nm). (B) Shelf-life stability of N1-MI15-sNAp-174 over 15 days at 4°C as assessed by NS-EM (scale bar, 10 nm). (C) Enzymatic activity of N1-MI15-WT compared to N1-MI15-sNAp-174, performed in duplicate. RFU, relative fluorescence units. Error bars are smaller than the plotted points and not shown. (D) Design of a predominantly closed N1 VN04 sNAp using sNAp-155 mutations in addition to T131Q and three other mutations in space D that are derived from N1 CA09 and N1 MI15 (scale bar, 10 nm). (E) Design of multiple N1 WSN33 sNAps using homology-directed mutations in combination with strain-specific mutations derived from N1 CA09 and N1 MI15. The mutations added to finalize N1-WSN33-sNAp-375 utilize the same mutation set featured in N1-MI15-sNAp-183. I442 was already native to N1 WSN33 (scale bar, 10 nm).

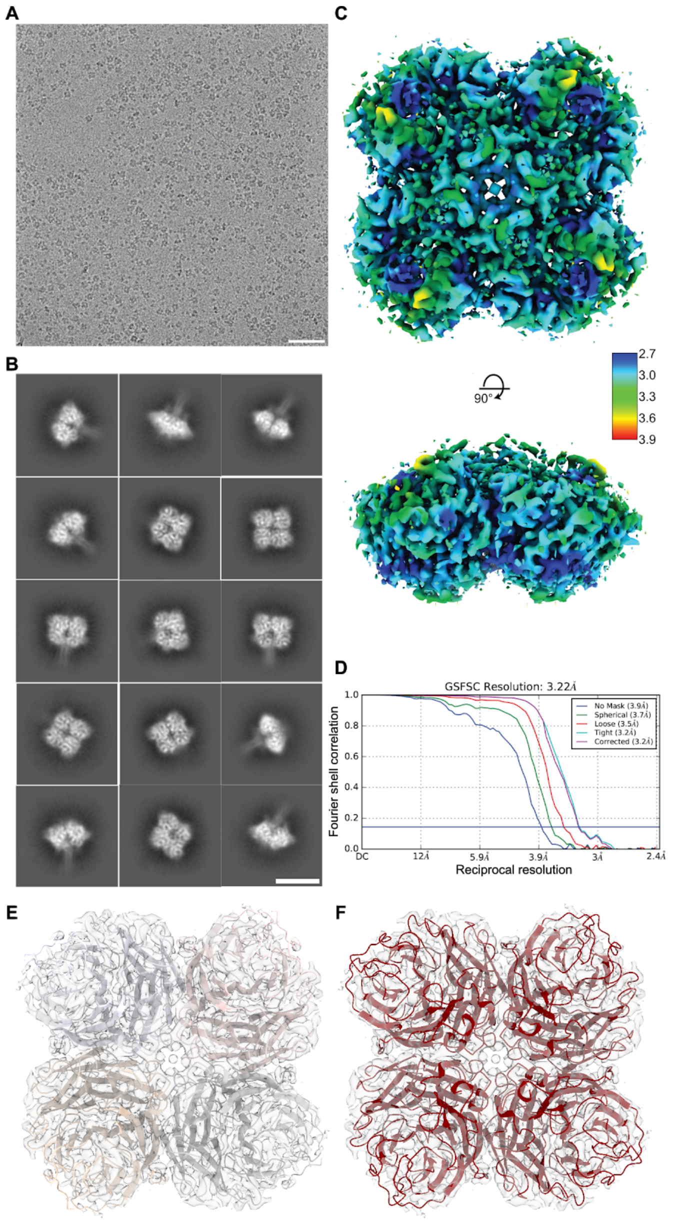


**Extended Data Fig. 6 | Cryo-EM analysis of N1-MI15-sNAp-174**

(A) Representative micrograph of N1-MI15-sNAp-174. Scale bar, 50 nm. (B) 2D class averages of N1-MI15-sNAp-174. Scale bar, 10 nm. (C) Orthogonal views of the cryo-EM reconstruction of N1-CA09-sNAp-155. Map is colored according to local resolution as indicated by color scale (in Å). (D) Gold standard FSC for cryo-EM reconstruction. (E) N1-CA09-sNAp-174 model coloured by chain shown in transparent density showing overall tetramer conformation of map (F) N1-CA09-WT (4B7Q, red) rigid-body docked into the N1-MI15-sNAp-174 map, showing overall agreement with tetramers observed by X-ray crystallography.

**Extended Data Table 1 | Mutations to Rescued H1N1 CA09 Viruses with and without N1-CA09-sNAp-155 Mutations, Related to Fig. 6**

| | CA09 with WT NA | CA09 with sNAp-155 NA |
|---|---|---|
| **HA substitutions*** | V41I, G148S | K171N |
| **NA substitution*** | None | T466A |

*Non-synonymous mutations in HA and NA genes of R3ΔPB1 A/California/07/2009 H1N1 virus and R3ΔPB1 A/California/07/2009 H1N1 sNAp-155 virus on MDCK-SIAT1 PB1 cells. Both HA and NA numbering start from the methionine at the beginning of each open reading frame.
