## Supplemental Item 1 for "Structure-based design of stabilized recombinant influenza neuraminidase tetramers"

All NA head sequences shown below were preceded with either of the following sequences containing secretion signal, affinity tags, hVASP tetramerization domain, thrombin cleavage site and GG linker:

Only Hexa-His tag

MEFGLSWIFLAAILKGVQCADPHHHHHHSSSDYSDLQRVKQELLEEVKKELQKVKEEIIEAFVQ
ELRKRGSLVPRGSGG

Both Hexa-His tag and Strep tag

MEFGLSWIFLAAILKGVQCADPHHHHHHGSAWSHPQFEKGGSSSDYSDLQRVKQELLEEVKKEL
QKVKEEIIEAFVQELRKRGSLVPRGSGG

N1-CA09-WT

H1N1 A/California/07/2009

VKLAGNSSLCPVSGWAIYSKDNSVRIGSKGDVFVIREPFISCSPLECRTFFLTQGALLNDKHSN
GTIKDRSPYRTLMSCPIGEVPSPYNSRFESVAWSASACHDGINWLTIGISGPDNGAVAVLKYNG
IITDTIKSWRNNILRTQESECACVNGSCFTVMTDGPSNGQASYKIFRIEKGKIVKSVEMNAPNY
HYEECSCYPDSSEITCVCRDNWHGSNRPWVSFNQNLEYQIGYICSGIFGDNPRPNDKTGSCGPV
SSNGANGVKGFSYKYGNGVWIGRTKSISSRNGFEMIWDPNGWTGTDNNFSIKQDIVGINEWSGY
SGSFVQHPELTGLDCIRPCFWVELIRGRPKENTIWTSGSSISFCGVNSDTVGWSWPDGAELPFT
IDK

N1-MI15-WT

H1N1 A/Michigan/45/2015

VKLAGNSSLCPVSGWAIYSKDNSVRIGSKGDVFVIREPFISCSPLECRTFFLTQGALLNDKHSN
GTIKDRSPYRTLMSCPIGEVPSPYNSRFESVAWSASACHDGINWLTIGISGPDSGAVAVLKYNG
IITDTIKSWRNNILRTQESECACVNGSCFTIMTDGPSDGQASYKIFRIEKGKIIKSVEMKAPNY
HYEECSCYPDSSEITCVCRDNWHGSNRPWVSFNQNLEYQMGYICSGVFGDNPRPNDKTGSCGPV
SSNGANGVKGFSFKYGNGVWIGRTKSISSRKGFEMIWDPNGWTGTDNKFSIKQDIVGINEWSGY
SGSFVQHPELTGLDCIRPCFWVELIRGRPEENTIWTSGSSISFCGVNSDTVGWSWPDGAELPFT
IDK

N1-NC99-WT

H1N1 A/New_Caledonia/20/1999

VTLAGNSSLCSISGWAIYTKDNSIRIGSKGDVFVIREPFISCSHLECRTFFLTQGALLNDKHSN
GTVKDRSPYRALMSCPLGEAPSPYNSKFESVAWSASACHDGMGWLTIGISGPDNGAVAVLKYNG
IITETIKSWKKRILRTQESECVCVNGSCFTIMTDGPSNGAASYKIFKIEKGKVTKSIELNAPNF
HYEECSCYPDTGTVMCVCRDNWHGSNRPWVSFNQNLDYQIGYICSGVFGDNPRPKDGEGSCNPV
TVDGADGVKGFSYKYGNGVWIGRTKSNRLRKGFEMIWDPNGWTDTDSDFSVKQDVVAITDWSGY
SGSFVQHPELTGLDCIRPCFWVELVRGLPRENTTIWTSGSSISFCGVNSDTANWSWPDGAELPF
TIDK

N1-WSN33-WT

H1N1 A/WSN/1933

VILTGNSSLCPIRGWAIHSKDNGIRIGSKGDVFVIREPFISCSHLECRTFFLTQGALLNDKHSR
GTFKDRSPYRALMSCPVGEAPSPYNSRFESVAWSASACHDGMGWLTIGISGPDDGAVAVLKYNR
IITETIKSWRKNILRTQESECTCVNGSCFTIMTDGPSDGLASYKIFKIEKGKVTKSIELNAPNS
HYEECSCYPDTGKVMCVCRDNWHGSNRPWVSFDQNLDYKIGYICSGVFGDNPRPKDGTGSCGPV
SADGANGVKGFSYKYGNGVWIGRTKSDSSRHGFEMIWDPNGWTETDSRFSMRQDVVAITNRSGY

SGSFVQHPELTGLDCMRPCFWVELIRGLPEEDAIWTSGSIISFCGVNSDTVDWSWPDGAELPFT
IDK

N1-BV18-WT
H1N1 A/Brevig_Mission/1/1918
VILTGNSSLCPISGWAIYSKDNGIRIGSKGDVFVIREPFISCSHLECRTFFLTQGALLNDKHSN
GTVKDRSPYRTLMSCPVGEAPSPYNSRFESVAWSASACHDGMGWLTIGISGPDNGAVAVLKYNG
IITDTIKSWRNNILRTQESECACVNGSCFTIMTDGPSNGQASYKILKIEKGKVTKSIELNAPNY
HYEECSCYPDTGKVMCVCRDNWHGSNRPWVSFDQNLDYQIGYICSGVFGDNPRPNDGTGSCGPV
SSNGANGIKGFSFRYDNGVWIGRTKSTSSRSGFEMIWDPNGWTETDSSFSVRQDIVAITDWSGY
SGSFVQHPELTGLDCMRPCFWVELIRGQPKENTIWTSGSSISFCGVNSDTVGWSWPDGAELPFS
IDK

N6-SI14-WT
H5N6 A/Sichuan/26221/2014
HLLNLTKPLCEVNSWHILSKDNAIRIGEDAHIIVTREPYLSCDPQGCRMFALSQGTTLRGKHAN
GTIHDRSPFRALVSWEMGQAPSPYNTRVECIGWSSTSCHDGISRMSICISGPNNNASAVVWYGG
RPVTEIPSWAGNILRTQESECVCHGGICPVVMTDGPANNRAETKIIYFKEGKIKKIEELKGDAQ
HIEECSCYGASEMIKCICRDNWKGANRPVITIDPEMMTHTSKYLCSKILTDTSRPNDPTNGKCE
APITGGSPDPGVKGFAFLDGENSWLGRTISKDSRSGYEMLKVPNAETDTQSGAISHQIIVNNQN
WSGYSGAFIDYWANKECFNPCFYVELIRGRPKESSVLWTSNSIVALCGSKERLGSWSWHDGAEI
IYFK

N2-MO99-WT
H3N2 A/Moscow/10/1999
EYRNWSKPQCNITGFAPFSKDNSIRLSAGGDIWVTREPYVSCDPDKCYQFALGQGTTLNNGHSN
DTVHDRTPYRTLLMNELGVPFHLGTKQVCIAWSSSSCHDGKAWLHVCVTGDDENATASFIYNGR
LVDSIGSWSKKILRTQESECVCINGTCTVVMTDGSASGKADTKILFIEEGKIVHTSPLSGSAQH
VEECSCYPRYPGVRCVCRDNWKGSNRPIVDINVKDYSIVSSYVCSGLVGDTPRKNDSSSSSHCL
DPNNEEGGHGVKGWAFDDGNDVWMGRTISEKLRSGYETFKVIEGWSKPNSKLQINRQVIVDRGN
RSGYSGIFSVEGKSCINRCFYVELIRGRKQETEVLWTSNSIVVFCGTSGTYGTGSWPDGADINL
MPI

N4-DB16-WT
H10N4 A/Red_knot/Delaware_Bay/310/2016
VHYSSGRDLCPIRGWAPLSKDNGIRIGSRGEVFVIREPFISCSISECRTFFLTQGALLNDKHSN
GTVKDRSPFRTLMSCPIGVAPSPSNSRFESVAWSATACSDGPGWLTLGITGPDSTAVAVLKYNG
IITDTLKSWKGNIMRTQESECVCQDEFCYTLVTDGPSDAQAFYKILKIRKGKIVSMKDVDATGF
HFEECSCYPSGTEIECVCRDNWRGSNRPWIRFNSDLDYQIGYVCSGIFGDNPRPVDGTGSCNGP
VNNGKGRYGVKGFSFRYGDGVWIGRTKSLESRSGFEMVWDANGWVSTDKDSNGVQDIIDNDNWS
GYSGSFSIRGETTGKNCTVPCFWVEMIRGQPKEKTIWTSGSSIAFCGVNSDTTGWSWPDGALLP
FDIDK

N7-NE03-WT
H7N7 A/Netherlands/219/03
SYLLLNKSLCNVEGWVVIAKDNAVRFGESEQIIVTREPYVSCDPTGCKMYALHQGTTIRNKHSN
GTIHDRTAFRGLISTPLGTPPTVSNSDFMCVGWSSTTCHDGIARMTICIQGNNDNATATVYYNR
RLTTTIKTWARNILRTQESECVCHNGTCAVVMTDGSASSQAYTKVMYFHKGLVVKEEELRGSAR

HIEECSCYGHNQKVTCVCRDNWQGANRPIIEIDMSTLEHTSRYVCTGILTDTSRPGDKSSGDCS
NPITGSPGVPGVKGFGFLNGDNTWLGRTISPRSRSGFEMLKIPNAGTDPNSRIAERQEIVDNNN
WSGYSGSFIDYWNDNSECYNPCFYVELIRGRPEEAKYVWWASNSLIALCGSPFPVGSGSFPDGA
QIQYFS

N8-JD13-WT
H10N8 A/Jiangxi-Donghu/346-2/2013
HFMNNTEALCDAKGFAPFSKDNGIRIGSRGHVFVIREPFVSCSPTECRTFFLTQGSLLNDKHSN
GTVKDRSPYRTLMSVEIGQSPNVYQARFEAVAWSATACHDGKKWMTIGVTGPDAKAVAVVHYGG
IPTDVINSWAGDILRTQESSCTCIQGECFWVMTDGPANRQAQYRAFKAKQGKIVGQAEISFNGG
HIEECSCYPNEGKVECVCKDNWTGTNRPVLVISPDLSYRVGYLCAGLPSDTPRGEDSQFTGSCT
SPMGNQGYGVKGFGFRQGNDVWMGRTISRTSRSGFEILKVRNGWVQNSKEQIKRQVVVDNLNWS
GYSGSFTLPAELTKRNCLVPCFWVEMIRGNPEEKTIWTSSSSIVMCGVDHEIADWSWHDGAILP
FDIDKM

N9-AN13-WT
H7N9 A/Anhui/1/2013
NFNNLTKGLCTINSWHIYGKDNAVRIGESSDVLVTREPYVSCDPDECRFYALSQGTTIRGKHSN
GTIHDRSQYRALISWPLSSPPTVYNSRVECIGWSSTSCHDGKSRMSICISGPNNNASAVVWYNR
RPVAEINTWARNILRTQESECVCHNGVCPVVFTDGSATGPADTRIYYFKEGKILKWESLTGTAK
HIEECSCYGERTGITCTCRDNWQGSNRPVIQIDPVAMTHTSQYICSPVLTDNPRPNDPNIGKCN
DPYPGNNNNGVKGFSYLDGANTWLGRTISTASRSGYEMLKVPNALTDDRSKPIQGQTIVLNADW
SGYSGSFMDYWAEGDCYRACFYVELIRGRPKEDKVWWTSNSIVSMCSSTEFLGQWNWPDGAKIE
YFL

N2-WI05-WT
H3N2 A/Wisconsin/67/2005
EYRNWSKPQCNITGFAPFSKDNSIRLSAGGDIWVTREPYVSCDPDKCYQFALGQGTTLNNVHSN
DTVHDRTPYRTLLMNELGVPFHLGTKQVCIAWSSSSCHDGKAWLHVCVTGDDKNATASFIYNGR
LVDSIVSWSKEILRTQESECVCINGTCTVVMTDGSASGKADTKILFIEEGKIVHTSTLSGSAQH
VEECSCYPRYLGVRCVCRDNWKGSNRPIVDINIKDYSIVSSYVCSGLVGDTPRKNDSSSSSHCL
DPNNEEGGHGVKGWAFDDGNDVWMGRTISEKLRSGYETFKVIEGWSNPNSKLQINRQVIVDRGN
RSGYSGIFSVEGKSCINRCFYVELIRGRKEETEVLWTSNSIVVFCGTSGTYGTGSWPDGADINL
MPI

N2-IN11-WT
H3N2 A/Indiana/10/2011
EYRNWSKPQCNITGFAPFSKDNSIRLSAGGDIWVTREPYVSCDPDKCYQFALGQGTTLNNGHSN
NTVHDRTPYRTLLMNELGVPFHLGTRQVCMAWSSSSCHDGKAWLHVCITGNDNNATASFIYNGR
LVDSIGSWSKNILRTQESECVCINGTCTVVMTDGSASGKADTKILFVEEGKIVHISTLSGSAQH
VEECSCYPRFPGVRCVCRDNWKGSNRPIVDINVKNYSIVSSYVCSGLVGDTPRKSDSVSSSYCL
DPNNEKGGHGVKGWAFDDGNDVWMGRTINETLRLGYETFKVIEGWSKANSKLQTNRQVIVEKGD
RSGYSGIFSVEGKSCINRCFYVELIRGRKEETKVWWTSNSIVVFCGTSGTYGTGSWPDGADINL
MPI

N3-MI06-WT
H2N3 A/Swine/Missouri/2124514/2006

EEERPFKSPLPLCPFRGFFPFHKDNAIRLGENKDVIVTREPYVSCDNDNCWSFALAQGALLGTK
HSNGTIKDRTPYRSLIRFPIGTAPVLGNYKEICIAWSSSSCFDGKEWMHVCMTGNDNDASAQII
YGGRMTDSIKSWRKDILRTQESECQCIDGTCVVAVTDGPAANSADYRVYWIREGKIIKYENVPK
TKIQHLEECSCYVDIDVYCICRDNWKGSNRPWMRINNETILETGYVCSKFHSDTPRPADPSTMS
CDSPSNVNGGPGVKGFGFKAGDDVWLGRTVSTSGRSGFEIIKVTEGWINSPNHVKSITQTLVSN
NDWSGYSGSFIVKAKDCFQPCFYVELIRGRPNKNDDVSWTSNSIVTFCGLDNEPGSGNWPDGSN
IGFMPK

N5-DB16-WT
H10N5 A/Shorebird/Delaware_Bay/309/2016
EFLNNTEPLCNVSGFAIVSKDNGIRIGSRGHVFVIREPFVACGPTECRTFFLTQGALLNDKHSN
NTVKDRSPYRALMSVPLGSSPNAYQAKFESVAWSATACHDGKRWLAVGISGADDDAYAVIHYGG
MPTDVVRSWRKQILRTQESSCVCMKGNCYWVMTDGPANSQASYKIFKSHKGMVTNEREVSFQGG
HIEECSCYPNLGKVECVCRDNWNGMNRPVLTFDEDLNYEVGYLCAGIPTDTPRVQDNSFIGSCT
NAVGGSGTNNYGVKGFGFRQGNSVWAGRTVSISSRSGFEILLVEDGWVKTSKNVVKKVEVLNNK
NWSGYSGAFTIPITMTSKQCLVPCFWLEMIRGKPEERTSIWTSSSSTVFCGVSSEVPGWSWDDG
AILPFDIDKM

B-CO17-WT
B-Victoria B/Colorado/06/2017
PEPEWTYPRLSCPGSTFQKALLISPHRFGETKGNSAPLIIREPFVACGPNECKHFALTHYAAQP
GGYYNGTRGDRNKLRHLISVKLGKIPTVENSIFHMAAWSGSACHDGKEWTYIGVDGPDNNALLK
VKYGEAYTDTYHSYANNILRTQESACNCIGGNCYLMITDGSASGVSECRFLKIREGRIIKEIFP
TGRVKHTEECTCGFASNKTIECACRDNRYTAKRPFVKLNVETDTAEIRLMCTDTYLDTPRPNDG
SITGPCESDGDKGSGGIKGGFVHQRMKSKIGRWYSRTMSQTERMGMGLYVKYGGDPWADSDALA
FSGVMVSMKEPGWYSFGFEIKDKKCDVPCIGIEMVHDGGKETWHSAATAIYCLMGSGQLLWDTV
TGVDMAL

B-PH13-WT
B-Yamagata B/Phuket/3073/2013
PEPEWTYPRLSCPGSTFQKALLISPHRFGETKGNSAPLIIREPFIACGPKECKHFALTHYAAQP
GGYYNGTREDRNKLRHLISVKLGKIPTVENSIFHMAAWSGSACHDGREWTYIGVDGPDSNALLK
IKYGEAYTDTYHSYAKNILRTQESACNCIGGDCYLMITDGPASGISECRFLKIREGRIIKEIFP
TGRVKHTEECTCGFASNKTIECACRDNSYTAKRPFVKLNVETDTAEIRLMCTKTYLDTPRPNDG
SITGPCESDGDEGSGGIKGGFVHQRMASKIGRWYSRTMSKTKRMGMGLYVKYDGDPWTDSEALA
LSGVMVSMEEPGWYSFGFEIKDKKCDVPCIGIEMVHDGGKTTWHSAATAIYCLMGSGQLLWDTV
TGVNMTL

N9-AN13-Y169aH
H7N9 A/Anhui/1/2013
NFNNLTKGLCTINSWHIYGKDNAVRIGESSDVLVTREPYVSCDPDECRFYALSQGTTIRGKHSN
GTIHDRSQYRALISWPLSSPPTVHNSRVECIGWSSTSCHDGKSRMSICISGPNNNASAVVWYNR
RPVAEINTWARNILRTQESECVCHNGVCPVVFTDGSATGPADTRIYYFKEGKILKWESLTGTAK
HIEECSCYGERTGITCTCRDNWQGSNRPVIQIDPVAMTHTSQYICSPVLTDNPRPNDPNIGKCN
DPYPGNNNNGVKGFSYLDGANTWLGRTISTASRSGYEMLKVPNALTDDRSKPIQGQTIVLNADW
SGYSGSFMDYWAEGDCYRACFYVELIRGRPKEDKVWWTSNSIVSMCSSTEFLGQWNWPDGAKIE
YFL

N1-CA09-sNAp-94
H1N1 A/California/07/2009
VKLAGNSSLCPVSGWAPLSKDNSVRIGSKGEVFVIREPFISCSPLECRTFFLTQGALLNDKHSN
GTIKDRSPYRTLMSCPIGSVPSPSNSRFESVAWSASACHDGINWLTIGITGPDNGAVAILKYNG
IITDTIKSWRNNILRTQESECACVNGSCFTVMTDGPSNGQASYKIFRIEKGKIVKSVEMNAPNY
HYEECSCYPDSSEITCVCRDNWHGSNRPWVSFNQNLEYQIGYICSGIFGDNPRPNDKTGSCGPV
SSNGANGVKGFSFKYGNGVWIGRTKSISSRNGFEMIWDPNGWTGTDNNFSIKQDIVGINEWSGY
SGSFVMHPELTGLDCIVPCFWVELIRGRPKENTIWTSGSSISFCGVNSDTTGWSWPDGAELPFT
IDK

N1-CA09-sNAp-114
H1N1 A/California/07/2009
VKLAGNSSLCPVSGWAPLSKDNSVRIGSKGEVFVIREPFISCSPLECRTFFLTQGALLNDKHSN
GTIKDRSPYRTLMSCPIGSVPSPSNSRFESIAWSASACHDGINWLTIGITGPDNGAVAILKYNG
IITDTIKSWRNNILRTQESECACVNGSCFTVMTDGPSNGQASYKIFRIEKGKIVKSVEMNAPNY
HYEECSCYPDSSEITCVCRDNWHGSNRPWVSFNQNLEYQIGYICSGIFGDNPRPNDKTGSCGPV
SSNGANGVKGFSFKYGNGVWIGRTKSISSRNGFEMIWDPNGWTGTDNNFSIKQDIVGINEWSGY
SGSFVMHPELTGLDCIVPCFWVELIRGRPKENTIWTSGSSISFCGVNSDTTGWSWPDGAELPFT
IDK

N1-CA09-sNAp-130
H1N1 A/California/07/2009
VKLAGNSSLCPVSGWAPLSKDNSVRIGSKGDVFVIREPFISCSPLECRTFFLTQGALLNDKHSN
GTIKDRSPYRTLMSVPIGSVPSPYNARFESIAWSASACHDGINWLTIGITGPDNGAVAILKYNG
IITDTIKSWRNNILRTQESECACVNGSCFTVMTDGPSNGQASYKIFRIEKGKIVKSVEMNAPNY
HYEECSCYPDSSEITCVCRDNWHGSNRPWVSFNQNLEYQIGYICSGIFGDNPRPNDKTGSCGPV
SSNGANGVKGFSFKYGNGVWIGRTKSISSRNGFEMIWDPNGWTGTDNNFSIKQDIVGINEWSGY
SGSFVMHPELTGLDCIVPCFWVELIRGRPKENTIWTSGSSISFCGVNSDTTGWSWPDGAELPFT
IDK

N1-CA09-sNAp-155
H1N1 A/California/07/2009
VKLAGNSSLCPVSGWAPLSKDNSVRIGSKGDVFVIREPFISCSPLECRTFFLTQGALLNDKHSN
GTIKDRSPYRTLMSVPIGSVPSPYNARFESIAWSASACHDGINWLTIGITGPDNGAVAILKYNG
IITDTIKSWRNNILRTQESECACVNGSCFTVMTDGPSNGQASYKIFRIEKGKIVKSVEMNAPNY
HYEECSCYPDSSEITCVCRDNWHGSNRPWVSFNQNLEYQIGYICSGIFGDNPRPNDKTGSCGPV
SSNGANGVKGFSFKYGNGVWIGRTKSISSRNGFEMIWDPNGWTGTDNNFSIKQDIVGINEWSGY
SGSFVMHPELTGLDCIVPCFWVELIRGRPKENTIWTSGSSISFCGVNSDTVGWSWPDGAELPFT
IDK

N1-CA09-sNAp-131
H1N1 A/California/07/2009
VKLAGNSSLCPVSGWAPYSKDNSVRIGSKGDVFVIREPFISCSPLECRTFFLTQGALLNDKHSN
GTIKDRSPYRTLMSVPIGSVPSPYNARFESIAWSASACHDGINWLTIGITGPDNGAVAILKYNG
IITDTIKSWRNNILRTQESECACVNGSCFTVMTDGPSNGQASYKIFRIEKGKIVKSVEMNAPNY
HYEECSCYPDSSEITCVCRDNWHGSNRPWVSFNQNLEYQIGYICSGIFGDNPRPNDKTGSCGPV
SSNGANGVKGFSFKYGNGVWIGRTKSISSRNGFEMIWDPNGWTGTDNNFSIKQDIVGINEWSGY

SGSFVQHPELTGLDCIRPCFWVELIRGRPKENTIWTSGSSISFCGVNSDTTGWSWPDGAELPFT
IDK

N1-CA09-sNAp-134

H1N1 A/California/07/2009

VKLAGNSSLCPVSGWAPLSKDNSVRIGSKGDVFVIREPFISCSPLECRTFFLTQGALLNDKHSN
GTIKDRSPYRTLMSVPIGEVPSPYNARFESIAWSASACHDGINWLTIGITGPDNGAVAILKYNG
IITDTIKSWRNNILRTQESECACVNGSCFTVMTDGPSNGQASYKIFRIEKGKIVKSVEMNAPNY
HYEECSCYPDSSEITCVCRDNWHGSNRPWVSFNQNLEYQIGYICSGIFGDNPRPNDKTGSCGPV
SSNGANGVKGFSFKYGNGVWIGRTKSISSRNGFEMIWDPNGWTGTDNNFSIKQDIVGINEWSGY
SGSFVMHPELTGLDCIVPCFWVELIRGRPKENTIWTSGSSISFCGVNSDTVGWSWPDGAELPFT
IDK

N8-JD13-sNAp-282

H10N8 A/Jiangxi-Donghu/346-2/2013

HFMNNTEALCDAKGFAPFSKDNGIRIGSRGHVFVIREPFVSCSPTECRTFFLTQGSLLNDKHSN
GTVKDRSPYRTLMSVEIGSSPNVYQARFEAVAWSATACHDGKKWMTIGVTGPDAKAVAVVHYGG
IPTDVINSWAGDILRTQESSCTCIQGECFWVMTDGPANRQAQYRAFKAKQGKIVGQAEISFNGG
HIEECSCYPNEGKVECVCKDNWTGTNRPVLVISPDLSYRVGYLCAGLPSDTPRGEDSQFTGSCT
SPMGNQGYGVKGFGFRQGNDVWMGRTISRTSRSGFEILKVRNGWVQNSKEQIKRQVVVDNLNWS
GYSGSFTLPAELTKRNCLVPCFWVEMIRGNPEEKTIWTSSSSIVMCGVDHEIADWSWHDGAILP
FDIDKM

N8-JD13-sNAp-285

H10N8 A/Jiangxi-Donghu/346-2/2013

HFMNNTEALCDAKGFAPFSKDNGIRIGSRGHVFVIREPFVSCSPTECRTFFLTQGSLLNDKHSN
GTVKDRSPYRTLMSVPIGSSPNVYQARFEAVAWSATACHDGKKWMTIGVTGPDAKAVAVVHYGG
IPTDVINSWAGDILRTQESSCTCIQGECFWVMTDGPANRQAQYRAFKAKQGKIVGQAEISFNGG
HIEECSCYPNEGKVECVCKDNWTGTNRPVLVISPDLSYRVGYLCAGLPSDTPRGEDSQFTGSCT
SPMGNQGYGVKGFGFRQGNDVWMGRTISRTSRSGFEILKVRNGWVQNSKEQIKRQVVVDNLNWS
GYSGSFTLPAELTKRNCLVPCFWVEMIRGNPEEKTIWTSSSSIVMCGVDHEIADWSWHDGAILP
FDIDKM

N2-WI05-desNAp-156

H3N2 A/Wisconsin/67/2005

EYRNWSKPQCNITGFAPFSKDNSIRLSAGGDIWVTREPYVSCDPDKCYQFALGQGTTLNNVHSN
DTVHDRTPYRTLLMNELGEPFHLGTKQVCIAWSSSSCHDGKAWLHVCVTGDDKNATASFIYNGR
LVDSIVSWSKEILRTQESECVCINGTCTVVMTDGSASGKADTKILFIEEGKIVHTSTLSGSAQH
VEECSCYPRYLGVRCVCRDNWKGSNRPIVDINIKDYSIVSSYVCSGLVGDTPRKNDSSSSSHCL
DPNNEEGGHGVKGWAFDDGNDVWMGRTISEKLRSGYETFKVIEGWSNPNSKLQINRQVIVDRGN
RSGYSGIFSVEGKSCINRCFYVELIRGRKEETEVLWTSNSIVVFCGTSGTYGTGSWPDGADINL
MPI

N2-WI05-desNAp-157

H3N2 A/Wisconsin/67/2005

EYRNWSKPQCNITGFAPFSKDNSIRLSAGGDIWVTREPYVSCDPDKCYQFALGQGTTLNNVHSN
DTVHDRTPYRTLLMNELGVPFHLGTKQVCVAWSSSSCHDGKAWLHVCVSGDDKNATASFIYNGR
LVDSIVSWSKEILRTQESECVCINGTCTVVMTDGSASGKADTKILFIEEGKIVHTSTLSGSAQH

VEECSCYPRYLGVRCVCRDNWKGSNRPIVDINIKDYSIVSSYVCSGLVGDTPRKNDSSSSSHCL
DPNNEEGGHGVKGWAFDDGNDVWMGRTISEKLRSGYETFKVIEGWSNPNSKLQINRQVIVDRGN
RSGYSGIFSVEGKSCINRCFYVELIRGRKEETEVLWTSNSIVVFCGTSGTYGTGSWPDGADINL
MPI

N2-WI05-desNAp-158
H3N2 A/Wisconsin/67/2005
EYRNWSKPQCNITGFAIFSKDNSIRLSAGGDIWVTREPYVSCDPDKCYQFALGQGTTLNNVHSN
DTVHDRTPYRTLLMNELGVPFHLGTKQVCVAWSSSSCHDGKAWLHVCVSGDDKNATASFIYNGR
LVDSIVSWSKEILRTQESECVCINGTCTVVMTDGSASGKADTKILFIEEGKIVHTSTLSGSAQH
VEECSCYPRYLGVRCVCRDNWKGSNRPIVDINIKDYSIVSSYVCSGLVGDTPRKNDSSSSSHCL
DPNNEEGGHGVKGWAFDDGNDVWMGRTISEKLRSGYETFKVIEGWSNPNSKLQINRQVIVDRGN
RSGYSGIFSVEGKSCINRCFYVELIRGRKEETEVLWTSNSIVVFCGTSGTYGTGSWPDGADINL
MPI

N2-WI05-desNAp-249
H3N2 A/Wisconsin/67/2005
EYRNWSKPQCNITGFAPFSKDNSIRLSAGGDIWVTREPYVSCDPDKCYQFALGQGTTLNNVHSN
DTVHDRTPYRTLLMNELGQPFHLGTKQVCIAWSSSSCHDGKAWLHVCVTGDDKNATASFIYNGR
LVDSIVSWSKEILRTQESECVCINGTCTVVMTDGSASGKADTKILFIEEGKIVHTSTLSGSAQH
VEECSCYPRYLGVRCVCRDNWKGSNRPIVDINIKDYSIVSSYVCSGLVGDTPRKNDSSSSSHCL
DPNNEEGGHGVKGWAFDDGNDVWMGRTISEKLRSGYETFKVIEGWSNPNSKLQINRQVIVDRGN
RSGYSGIFSVEGKSCINRCFYVELIRGRKEETEVLWTSNSIVVFCGTSGTYGTGSWPDGADINL
MP

N2-WI05-desNAp-255
H3N2 A/Wisconsin/67/2005
EYRNWSKPQCNITGFAPFSKDNSIRLSAGGDIWVTREPYVSCDPDKCYQFALGQGTTLNNVHSN
DTVHDRTPYRTLLMNELGQPFHLGTKQVCVAWSSSSCHDGKAWLHVCVSGDDKNATASFIYNGR
LVDSIVSWSKEILRTQESECVCINGTCTVVMTDGSASGKADTKILFIEEGKIVHTSTLSGSAQH
VEECSCYPRYLGVRCVCRDNWKGSNRPIVDINIKDYSIVSSYVCSGLVGDTPRKNDSSSSSHCL
DPNNEEGGHGVKGWAFDDGNDVWMGRTISEKLRSGYETFKVIEGWSNPNSKLQINRQVIVDRGN
RSGYSGIFSVEGKSCINRCFYVELIRGRKEETEVLWTSNSIVVFCGTSGTYGTGSWPDGADINL
MPI

N1-MI15-sNAp-155
VKLAGNSSLCPVSGWAPLSKDNSVRIGSKGDVFVIREPFISCSPLECRTFFLTQGALLNDKHSN
GTIKDRSPYRTLMSVPIGSVPSPYNARFESIAWSASACHDGINWLTIGITGPDSGAVAILKYNG
IITDTIKSWRNNILRTQESECACVNGSCFTIMTDGPSDGQASYKIFRIEKGKIIKSVEMKAPNY
HYEECSCYPDSSEITCVCRDNWHGSNRPWVSFNQNLEYQMGYICSGVFGDNPRPNDKTGSCGPV
SSNGANGVKGFSFKYGNGVWIGRTKSISSRKGFEMIWDPNGWTGTDNKFSIKQDIVGINEWSGY
SGSFVMHPELTGLDCIVPCFWVELIRGRPEENTIWTSGSSISFCGVNSDTVGWSWPDGAELPFT
IDK

N1-MI15-sNAp-174
H1N1 A/Michigan/45/2015
VKLAGNSSLCPVSGWAPLSKDNSVRIGSKGDVFVIREPFISCSPLECRQFFLTQGALLNDKHSN
GTIKDRSPYRTLMSVPIGSVPSPYNARFESIAWSASACHDGINWLTIGITGPDSGAVAILKYNG

IITDTIKSWRNNILRTQESECACVNGSCFTIMTDGPSDGQASYKIFRIEKGKIIKSVEMKAPNY
HYEECSCYPDSSEITCVCRDNWHGSNRPWVSFNQNLEYQMGYICSGVFGDNPRPNDKTGSCGPV
SSNGANGVKGFSFKYGNGVWIGRTKSISSRKGFEMIWDPNGWTGTDNKFSIKQDIVGINEWSGY
SGSFVMHPELTGLDCIVPCFWVELIRGRPEENTIWTSGSSISFCGVNSDTVGWSWPDGAELPFT
IDK

N1-MI15-sNAp-165

H1N1 A/Michigan/45/2015

VKLAGNSSLCPVSGWAPLSKDNSVRIGSKGDIFVIREPFISCSPLECRTFFLTQGALLNDKHSN
GTIKDRSPYRTLMSVPIGSPPSPYNARFESIAWSASACHDGINWLTIGITGPDSGAVAILKYNG
IITDTIKSWRNNILRTQESECACVNGSCFTIMTDGPSDGQASYKIFRIEKGKIIKSVEMKAPNY
HYEECSCYPDSSEITCVCRDNWHGSNRPWVSFNQNLEYQMGYICSGVFGDNPRPNDKTGSCGPV
SSNGANGVKGFSFKYGNGVWIGRTKSISSRKGFEMIWDPNGWTGTDNKFSIKQDIVGINEWSGY
SGSFVMHPELTGLDCIVPCFWVELIRGRPEENTIWTSGSSISFCGVNSDTVGWSWPDGAELPFT
IDK

N1-MI15-sNAp-176

H1N1 A/Michigan/45/2015

VKLAGNSSLCPVSGWAPLSKDNSVRIGSKGDIFVIREPFISCSPLECRQFFLTQGALLNDKHSN
GTIKDRSPYRTLMSVPIGSPPSPYNARFESIAWSASACHDGINWLTIGITGPDSGAVAILKYNG
IITDTIKSWRNNILRTQESECACVNGSCFTIMTDGPSDGQASYKIFRIEKGKIIKSVEMKAPNY
HYEECSCYPDSSEITCVCRDNWHGSNRPWVSFNQNLEYQMGYICSGVFGDNPRPNDKTGSCGPV
SSNGANGVKGFSFKYGNGVWIGRTKSISSRKGFEMIWDPNGWTGTDNKFSIKQDIVGINEWSGY
SGSFVMHPELTGLDCIVPCFWVELIRGRPEENTIWTSGSSISFCGVNSDTVGWSWPDGAELPFT
IDK

N1-MI15-sNAp-183

H1N1 A/Michigan/45/2015

VKLAGNSSLCPVSGWAPLAKDNSVRIGSKGDVFVIREPFISCSPLECRMFFLTQGALLNDKHSN
GTIKDRSPYRTLMSVPLGSVPSPYNARFESIAWSASACHDGINWLTIGITGPDSGAVAILKYNG
IITDTIKSWRNNILRTQESECACVNGSCFTIMTDGPSDGQASYKIFRIEKGKIIKSVEMKAPNY
HYEECSCYPDSSEITCVCRDNWHGSNRPWVSFNQNLEYQMGYICSGVFGDNPRPNDKTGSCGPV
SSNGANGVKGFSFKYGNGVWIGRTKSISSRKGFEMIWDPNGWTGTDNKFSIKQDIVGINEWSGY
SGSFVMHPELTGLDCIVPCFWVELIRGRPEENTIWTSGSIIAFCGVNSDTVGWSWPDGAELPFT
IDK

N1-VN04-sNAp-155

H5N1 A/Vietnam/1203/2004

VKLAGNSSLCPINGWAPLSKDNSIRIGSKGDVFVIREPFISCSHLECRTFFLTQGALLNDKHSN
GTVKDRSPHRTLMSVPVGSAPSPYNARFESIAWSASACHDGTSWLTIGITGPDNGAVAILKYNG
IITDTIKSWRNNILRTQESECACVNGSCFTVMTDGPSNGQASYKIFKMEKGKVVKSVELDAPNY
HYEECSCYPNAGEITCVCRDNWHGSNRPWVSFNQNLEYQIGYICSGVFGDNPRPNDGTGSCGPV
SSNGAYGVKGFSFKYGNGVWIGRTKSTNSRSGFEMIWDPNGWTETDSSFSVKQDIVAITDWSGY
SGSFVMHPELTGLDCIVPCFWVELIRGRPKESTIWTSGSSISFCGVNSDTVGWSWPDGAELPFT
IDK

N1-VN04-sNAp-354

H5N1 A/Vietnam/1203/2004
VKLAGNSSLCPINGWAPLSKDNSVRIGSKGDVFVIREPFISCSHLECRQFFLTQGALLNDKHSN
GTVKDRSPHRTLMSVPIGSVPSPYNARFESIAWSASACHDGTSWLTIGITGPDNGAVAILKYNG
IITDTIKSWRNNILRTQESECACVNGSCFTVMTDGPSNGQASYKIFKMEKGKVVKSVELDAPNY
HYEECSCYPNAGEITCVCRDNWHGSNRPWVSFNQNLEYQIGYICSGVFGDNPRPNDGTGSCGPV
SSNGAYGVKGFSFKYGNGVWIGRTKSTNSRSGFEMIWDPNGWTETDSSFSVKQDIVAITDWSGY
SGSFVMHPELTGLDCIVPCFWVELIRGRPKESTIWTSGSSISFCGVNSDTVGWSWPDGAELPFT
IDK

<u>N1-WSN33-sNAp-155</u>
H1N1 A/WSN/1933
VILTGNSSLCPIRGWAPLSKDNGIRIGSKGDVFVIREPFISCSHLECRTFFLTQGALLNDKHSR
GTFKDRSPYRALMSVPVGSAPSPYNARFESIAWSASACHDGMGWLTIGITGPDDGAVAILKYNR
IITETIKSWRKNILRTQESECTCVNGSCFTIMTDGPSDGLASYKIFKIEKGKVTKSIELNAPNS
HYEECSCYPDTGKVMCVCRDNWHGSNRPWVSFDQNLDYKIGYICSGVFGDNPRPKDGTGSCGPV
SADGANGVKGFSYKYGNGVWIGRTKSDSSRHGFEMIWDPNGWTETDSRFSMRQDVVAITNRSGY
SGSFVMHPELTGLDCMVPCFWVELIRGLPEEDAIWTSGSIISFCGVNSDTVDWSWPDGAELPFT
IDK

<u>N1-WSN33-sNAp-366</u>
H1N1 A/WSN/1933
VILTGNSSLCPIRGWAPLSKDNSVRIGSKGDVFVIREPFISCSHLECRTFFLTQGALLNDKHSR
GTFKDRSPYRTLMSVPIGSVPSPYNARFESIAWSASACHDGMGWLTIGITGPDDGAVAILKYNG
IITETIKSWRKNILRTQESECTCVNGSCFTIMTDGPSDGLASYKIFKIEKGKVTKSIELNAPNS
HYEECSCYPDTGKVMCVCRDNWHGSNRPWVSFDQNLDYKIGYICSGVFGDNPRPKDGTGSCGPV
SADGANGVKGFSYKYGNGVWIGRTKSDSSRHGFEMIWDPNGWTETDSRFSMRQDVVAITNRSGY
SGSFVMHPELTGLDCMVPCFWVELIRGLPEEDAIWTSGSIISFCGVNSDTVDWSWPDGAELPFT
IDK

<u>N1-WSN33-sNAp-367</u>
H1N1 A/WSN/1933
VILTGNSSLCPIRGWAPLSKDNSVRIGSKGDVFVIREPFISCSHLECRQFFLTQGALLNDKHSR
GTFKDRSPYRTLMSVPIGSVPSPYNARFESIAWSASACHDGMGWLTIGITGPDDGAVAILKYNG
IITETIKSWRKNILRTQESECTCVNGSCFTIMTDGPSDGLASYKIFKIEKGKVTKSIELNAPNS
HYEECSCYPDTGKVMCVCRDNWHGSNRPWVSFDQNLDYKIGYICSGVFGDNPRPKDGTGSCGPV
SADGANGVKGFSYKYGNGVWIGRTKSDSSRHGFEMIWDPNGWTETDSRFSMRQDVVAITNRSGY
SGSFVMHPELTGLDCMVPCFWVELIRGLPEEDAIWTSGSIISFCGVNSDTVDWSWPDGAELPFT
IDK

<u>N1-WSN33-sNAp-375</u>
H1N1 A/WSN/1933
VILTGNSSLCPIRGWAPLAKDNSVRIGSKGDVFVIREPFISCSHLECRMFFLTQGALLNDKHSR
GTFKDRSPYRTLMSVPLGSVPSPYNARFESIAWSASACHDGMGWLTIGITGPDDGAVAILKYNG
IITETIKSWRKNILRTQESECTCVNGSCFTIMTDGPSDGLASYKIFKIEKGKVTKSIELNAPNS
HYEECSCYPDTGKVMCVCRDNWHGSNRPWVSFDQNLDYKIGYICSGVFGDNPRPKDGTGSCGPV
SADGANGVKGFSYKYGNGVWIGRTKSDSSRHGFEMIWDPNGWTETDSRFSMRQDVVAITNRSGY
SGSFVMHPELTGLDCMVPCFWVELIRGLPEEDAIWTSGSIIAFCGVNSDTVDWSWPDGAELPFT
IDK

N1-WSN33-sNAp-378

H1N1 A/WSN/1933

```
VILTGNSSLCPIRGWAPLAKDNSVRIGSKGDVFVIREPFISCSPLECRMFFLTQGALLNDKHSR
GTFKDRSPYRTLMSVPLGSVPSPYNARFESIAWSASACHDGMGWLTIGITGPDDGAVAILKYNG
IITETIKSWRKNILRTQESECACVNGSCFTVMTDGPSDGLASYKIFKIEKGKITKSIEMNAPNS
HYEECSCYPDTGKVMCVCRDNWHGSNRPWVSFDQNLDYKIGYICSGVFGDNPRPKDGTGSCGPV
SADGANGVKGFSFKYGNGVWIGRTKSDSSRHGFEMIWDPNGWTETDSRFSMRQDIVAITNRSGY
SGSFVMHPELTGLDCIVPCFWVELIRGLPEEDAIWTSGSIIAFCGVNSDTVDWSWPDGAELPFT
IDK
```

N1-CA09-sNAp-155-T466A

H1N1 A/California/07/2009

```
VKLAGNSSLCPVSGWAPLSKDNSVRIGSKGDVFVIREPFISCSPLECRTFFLTQGALLNDKHSN
GTIKDRSPYRTLMSVPIGSVPSPYNARFESIAWSASACHDGINWLTIGITGPDNGAVAILKYNG
IITDTIKSWRNNILRTQESECACVNGSCFTVMTDGPSNGQASYKIFRIEKGKIVKSVEMNAPNY
HYEECSCYPDSSEITCVCRDNWHGSNRPWVSFNQNLEYQIGYICSGIFGDNPRPNDKTGSCGPV
SSNGANGVKGFSFKYGNGVWIGRTKSISSRNGFEMIWDPNGWTGTDNNFSIKQDIVGINEWSGY
SGSFVMHPELTGLDCIVPCFWVELIRGRPKENTIWTSGSSISFCGVNSDTVGWSWPDGAELPFA
IDK
```
