## Supplemental Item 2 for "Structure-based design of stabilized recombinant influenza neuraminidase tetramers"

# N1-CA09-WT

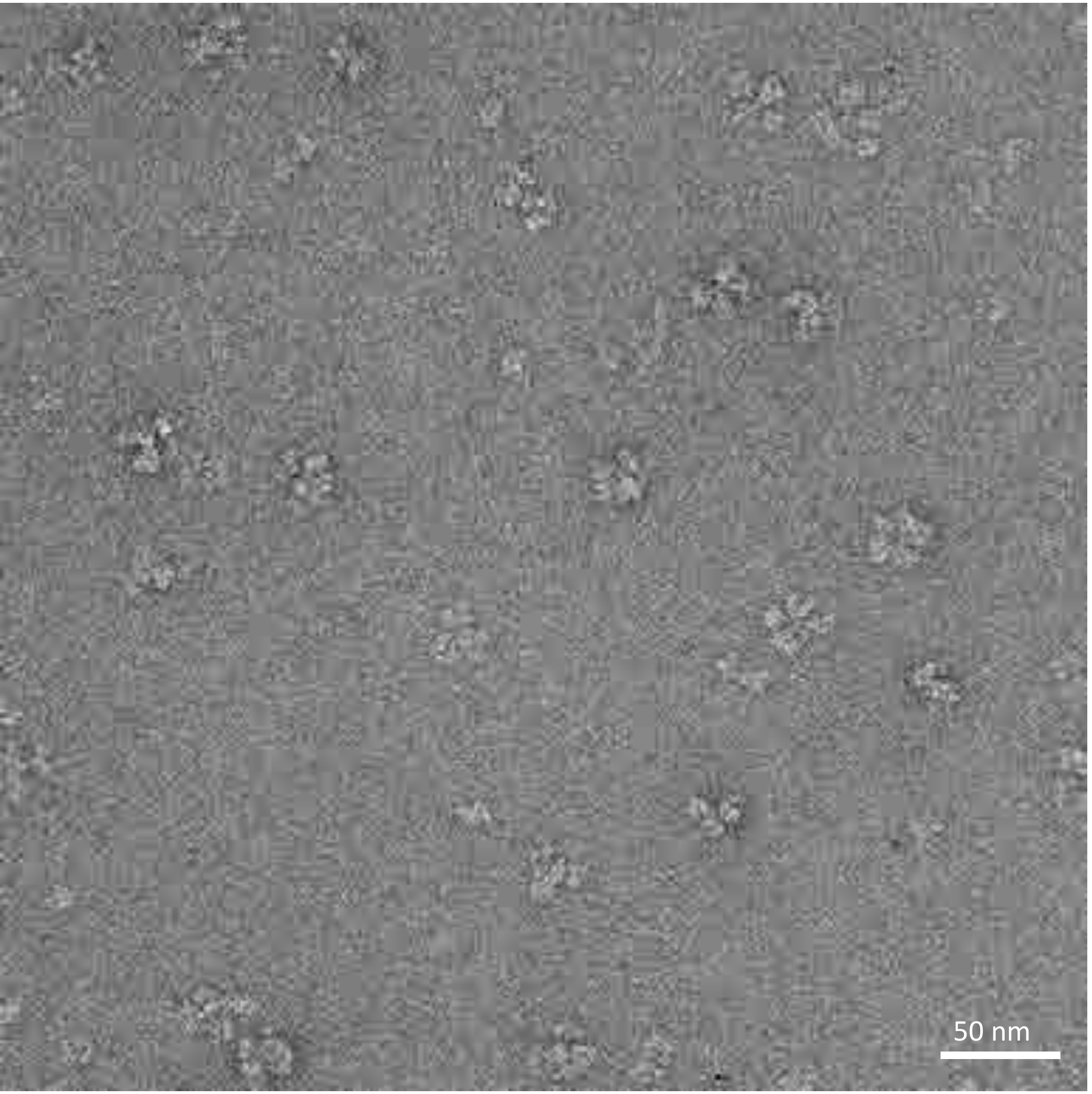


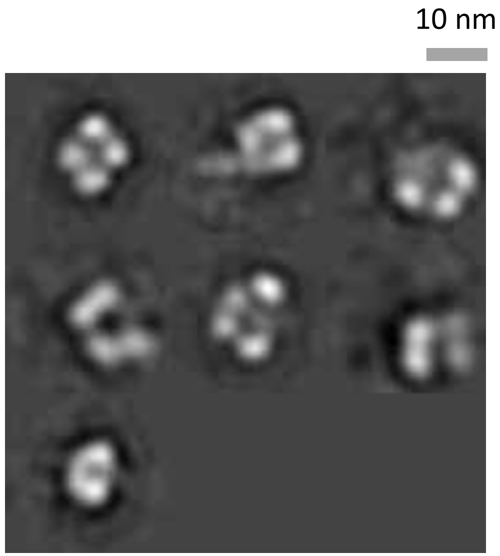

# N1-MI15-WT

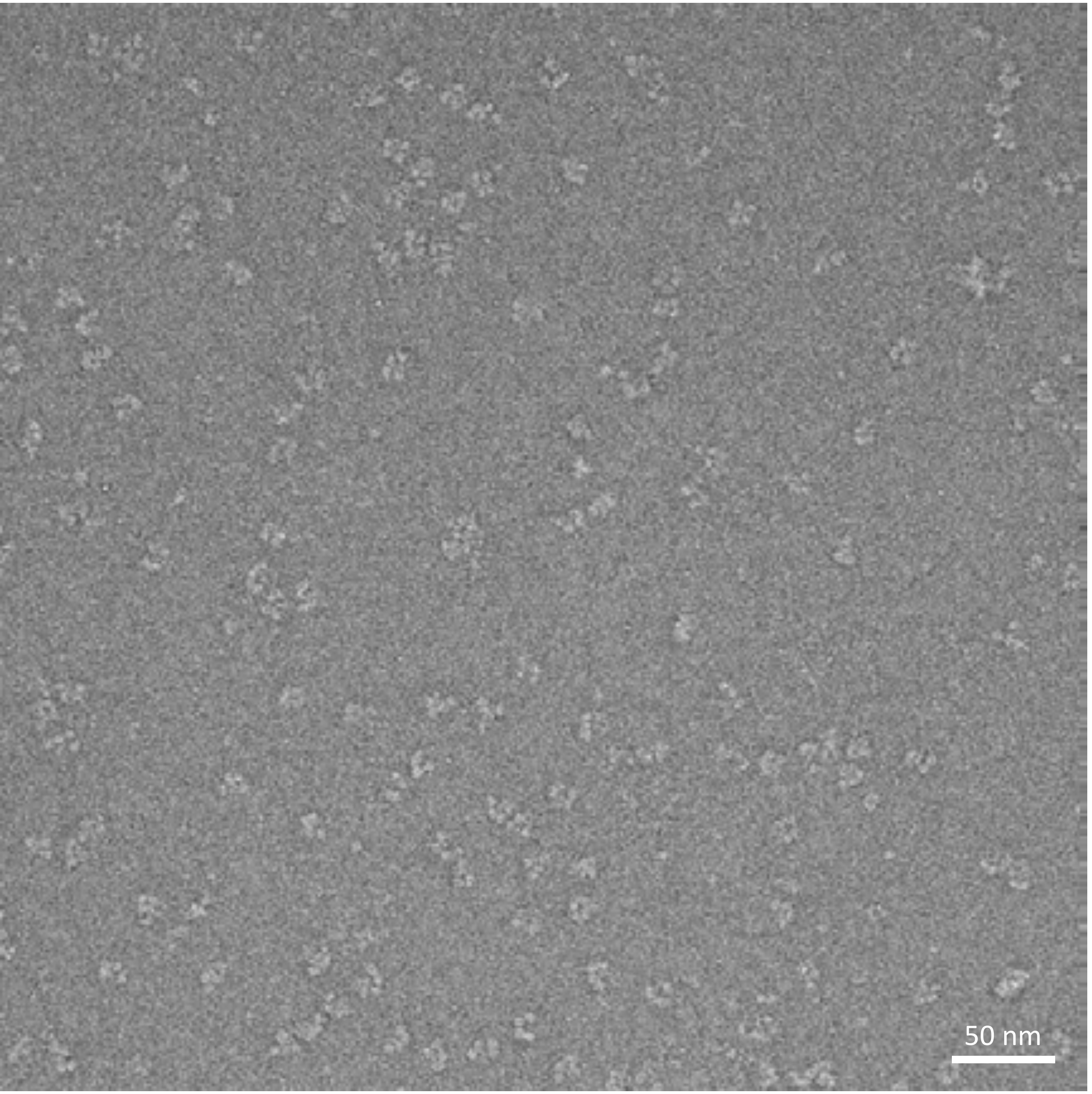


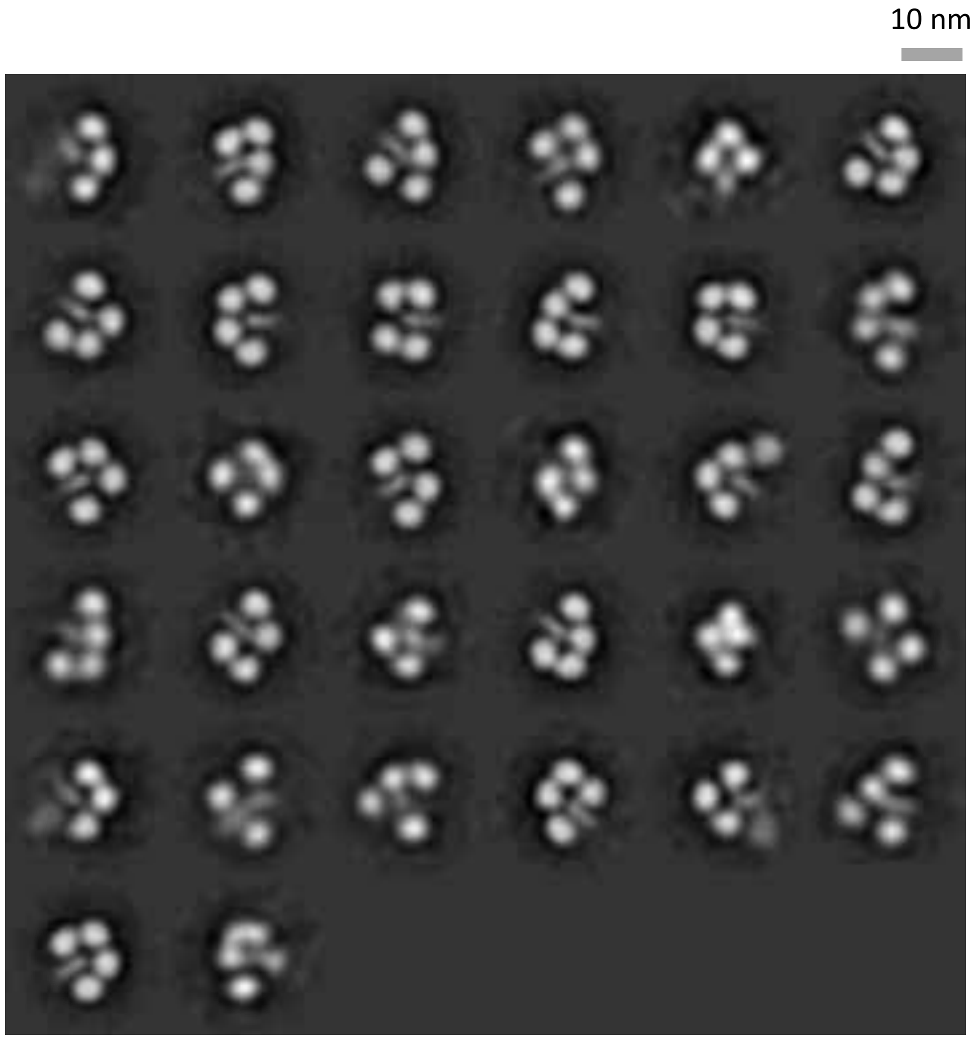

# N1-NC99-WT

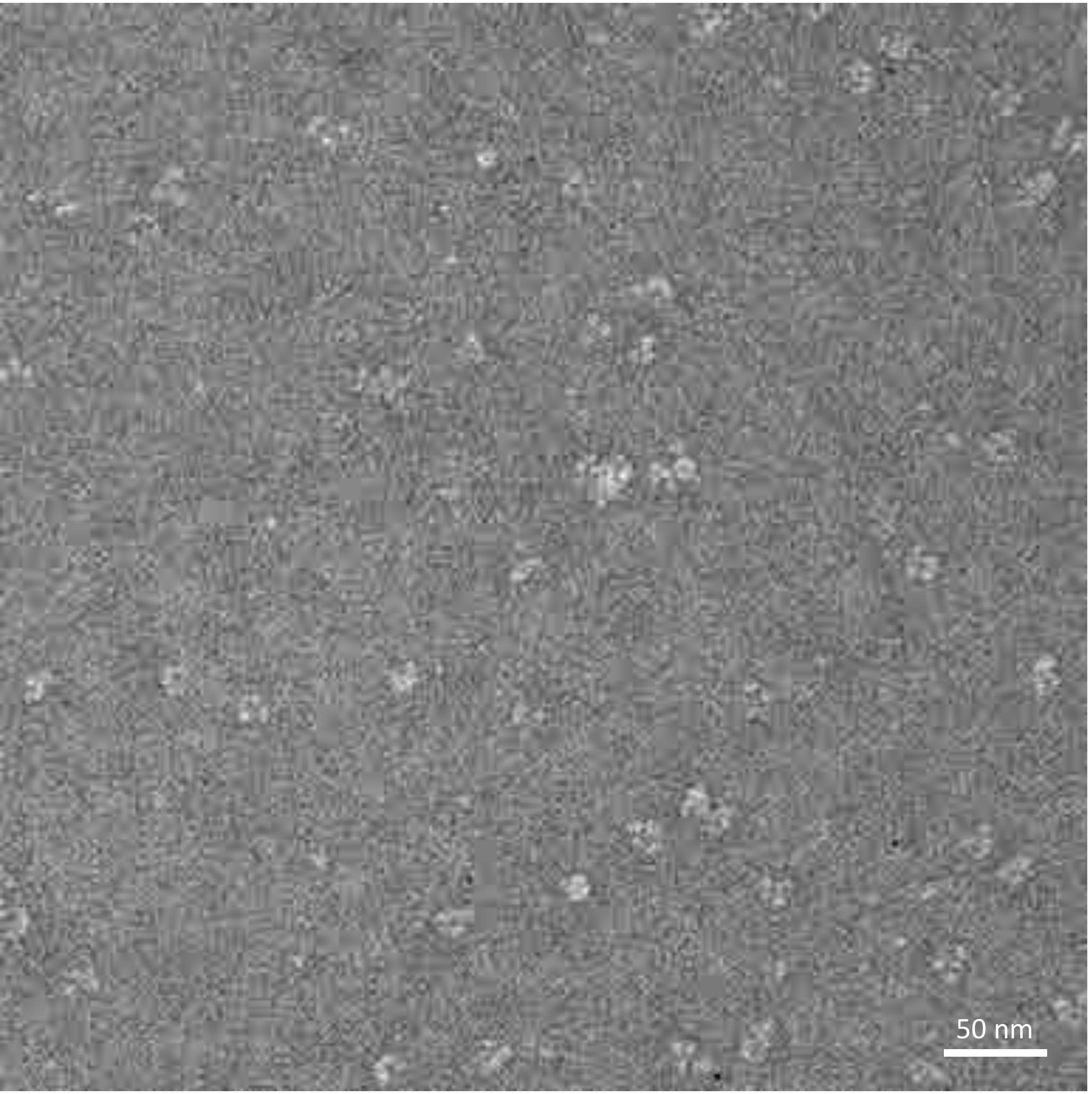


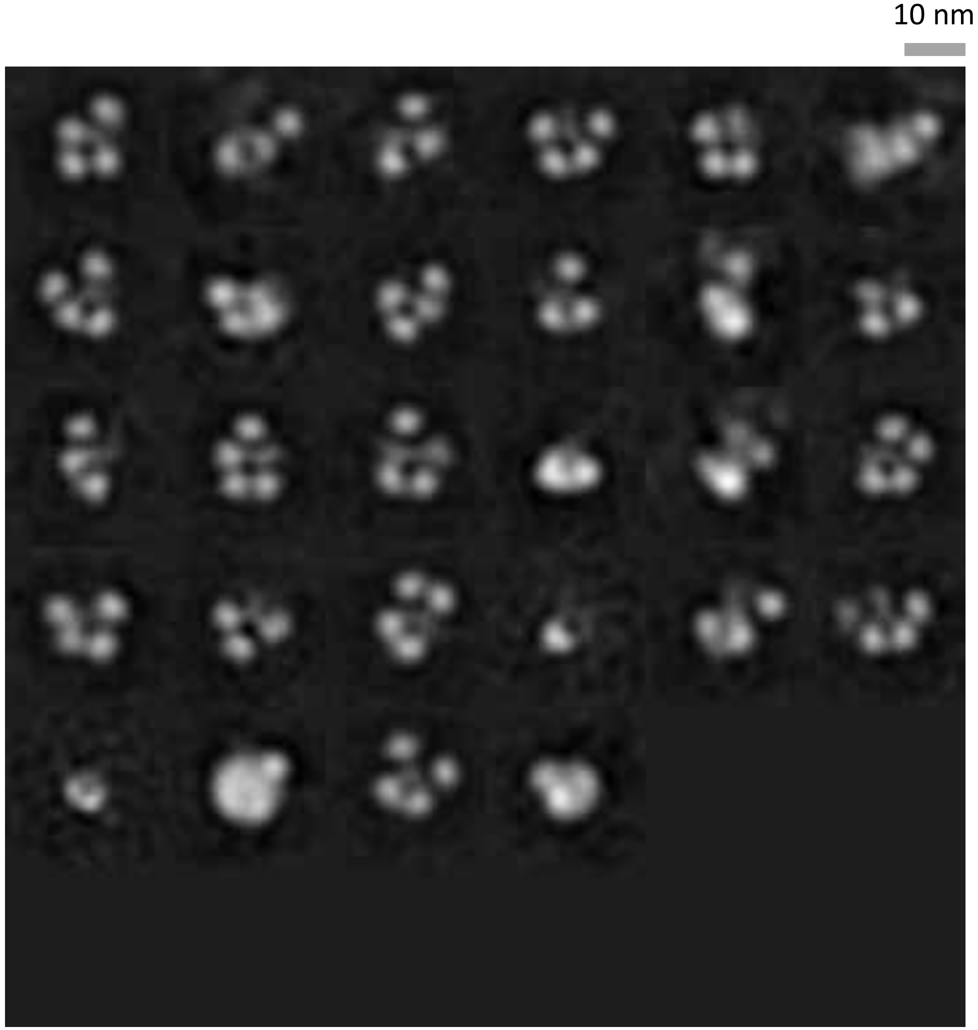

### N1-WSN33-WT

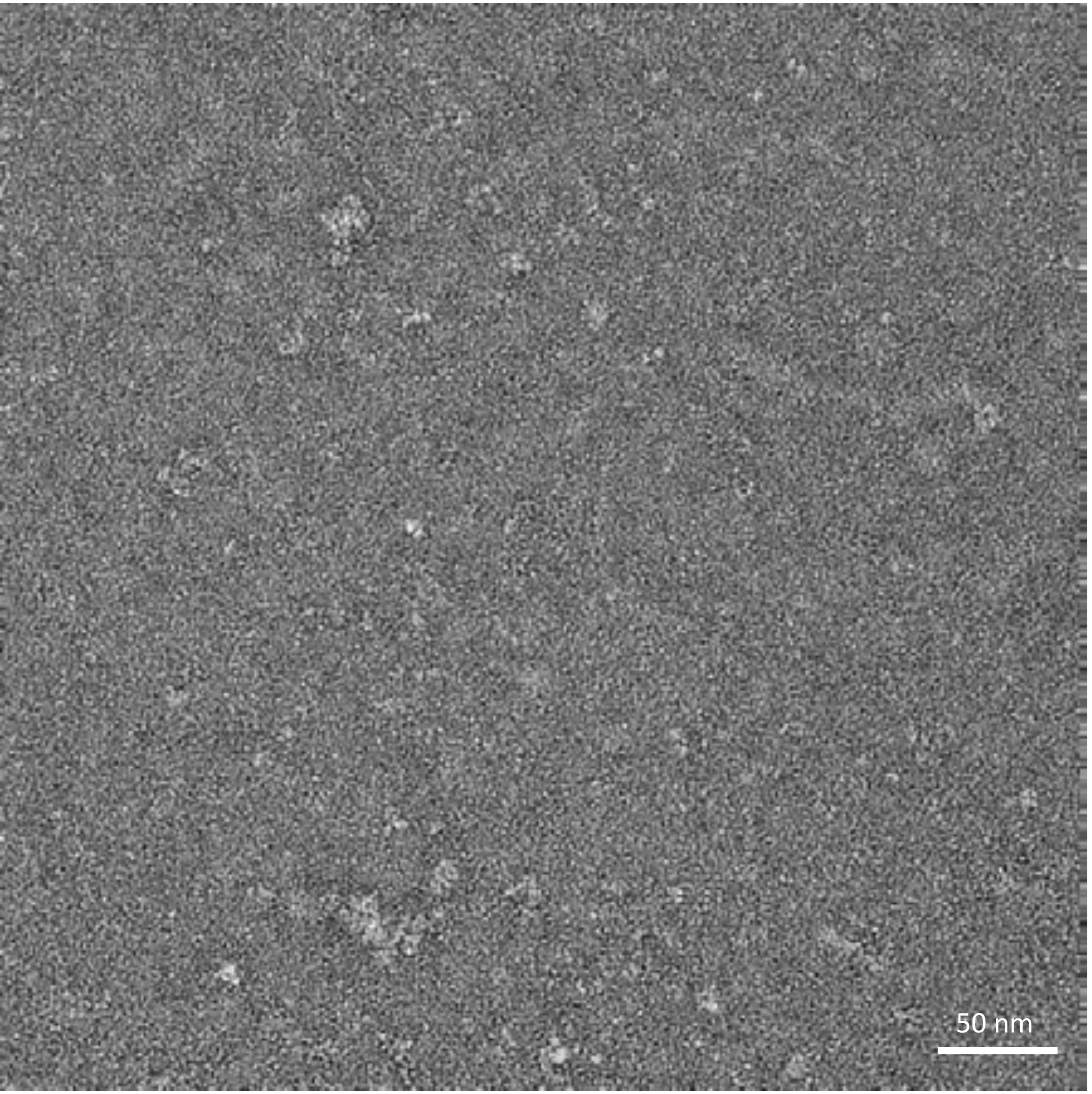


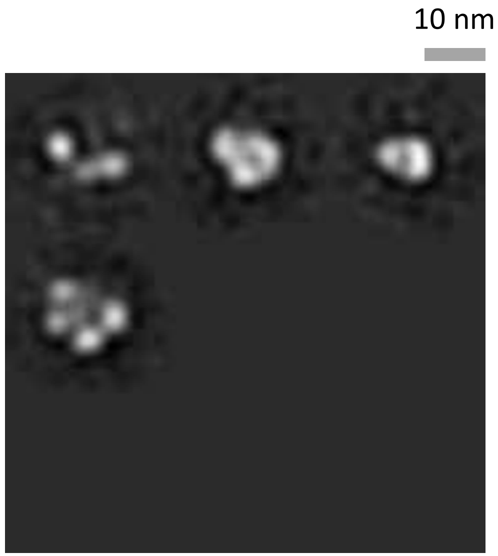

# N1-BV18-WT

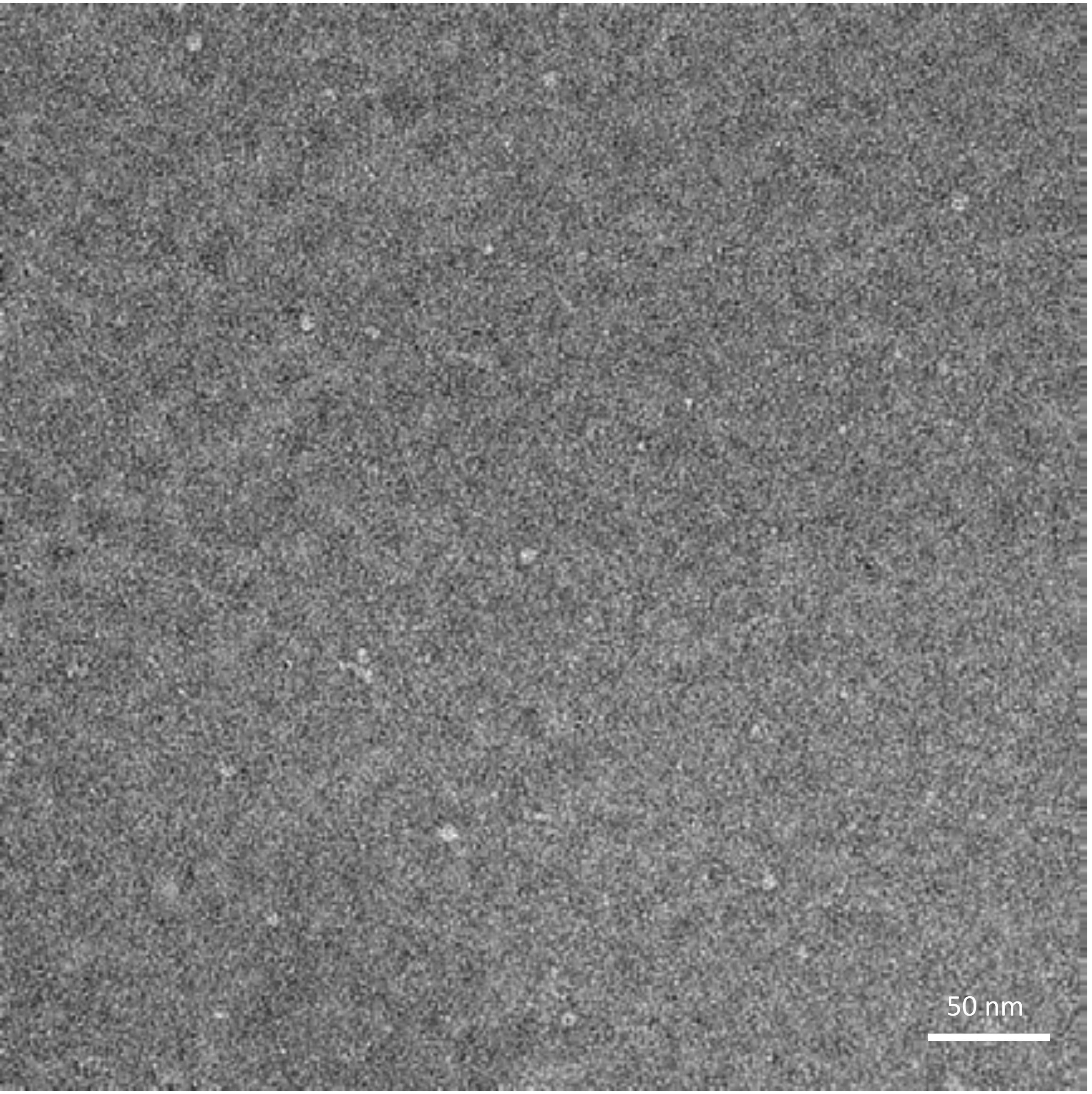


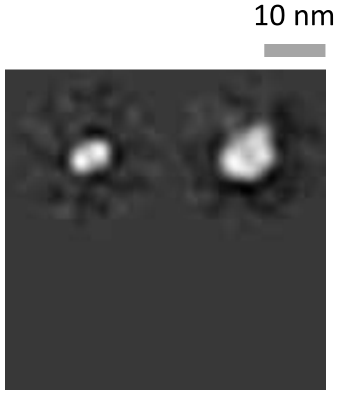

# N6-SI14-WT

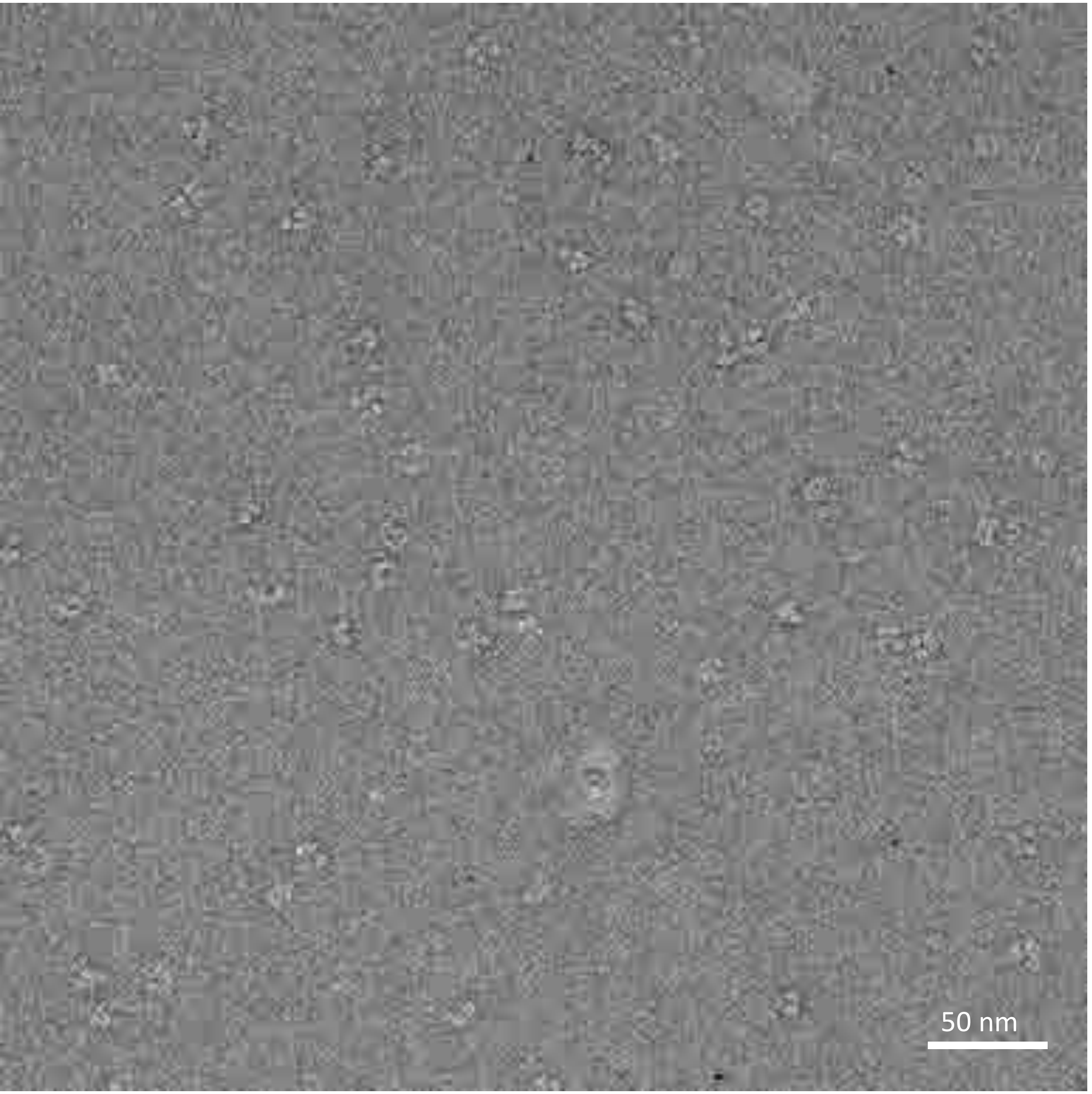


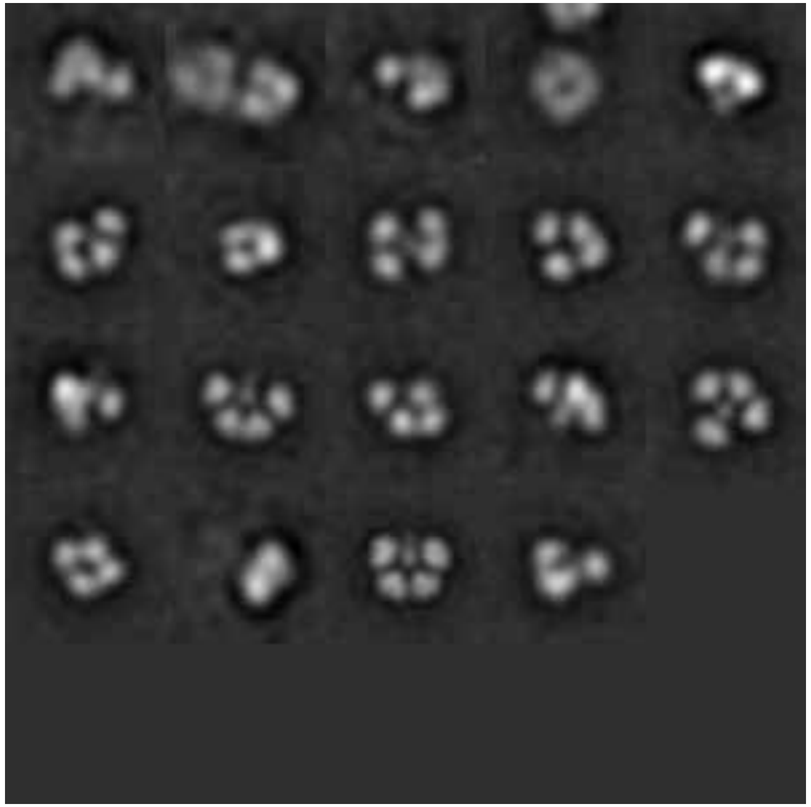

# N2-MO99-WT

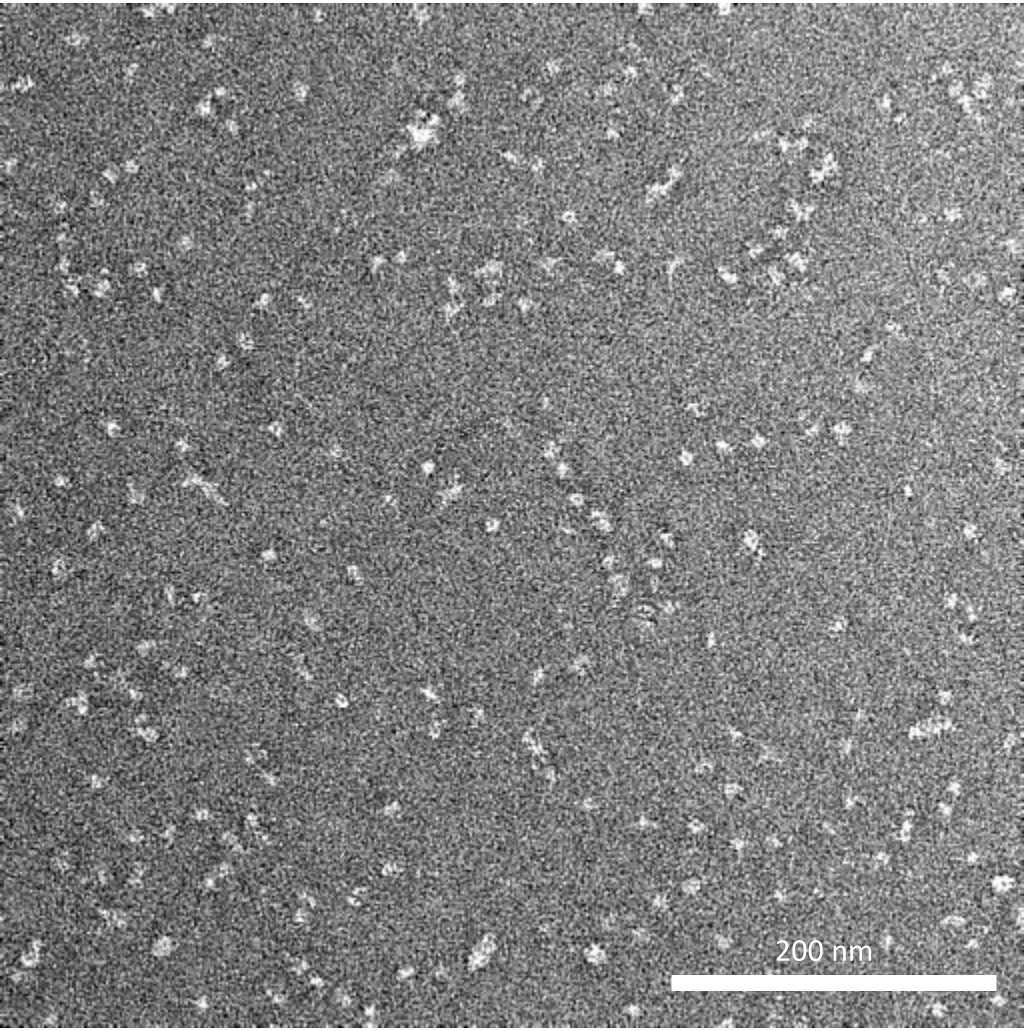


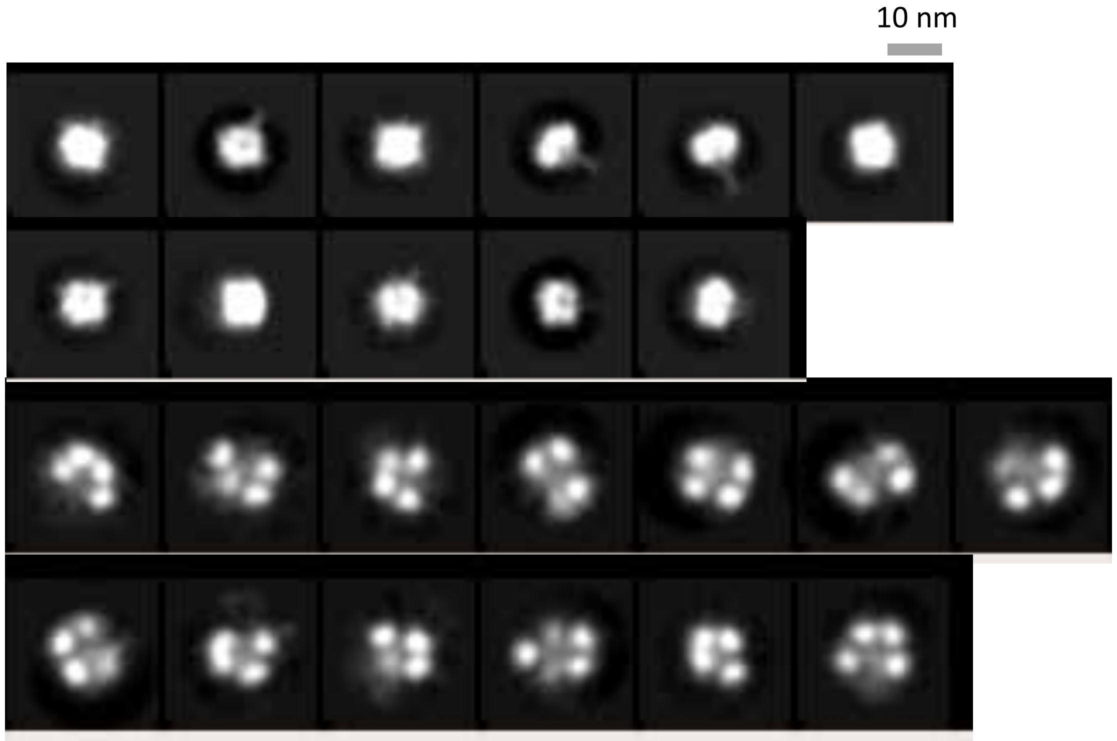

# N4-DB16-WT

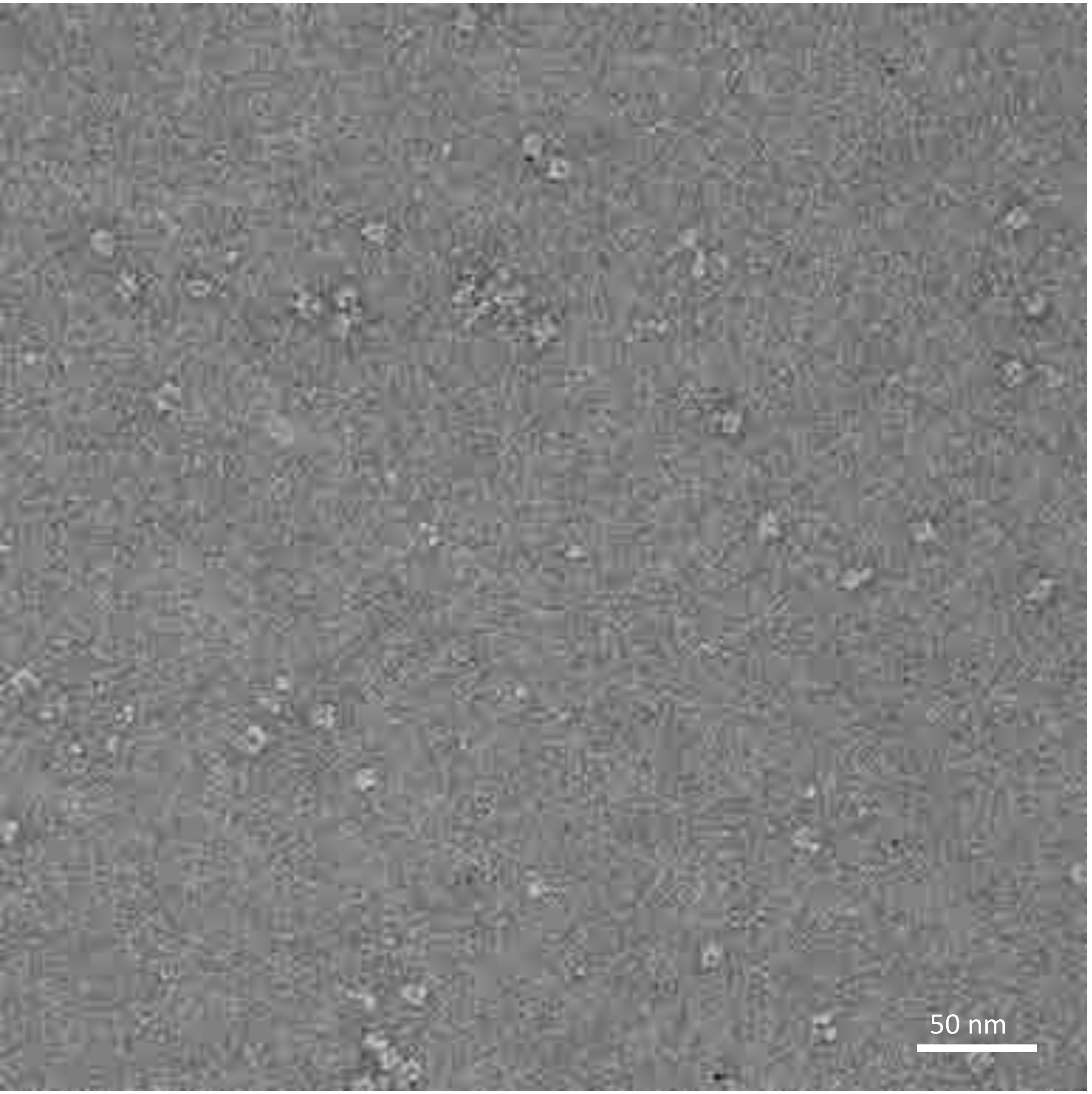


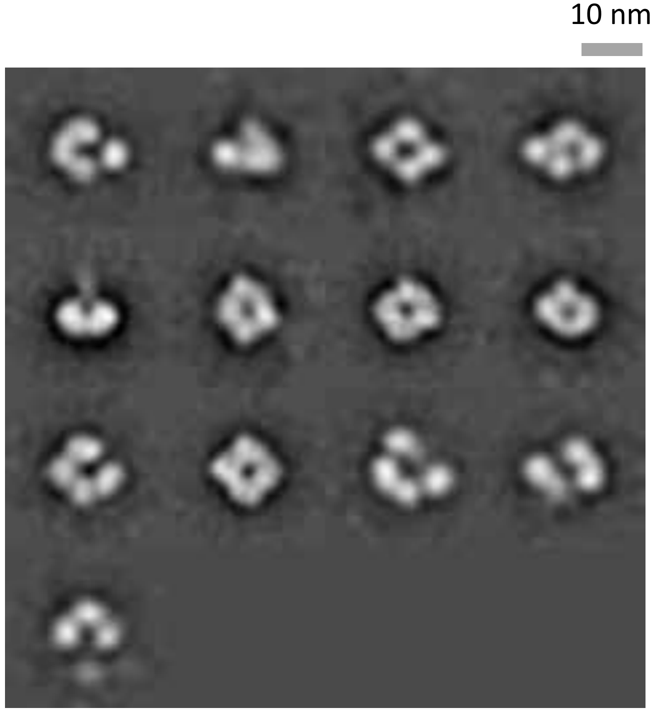

# N7-NE03-WT

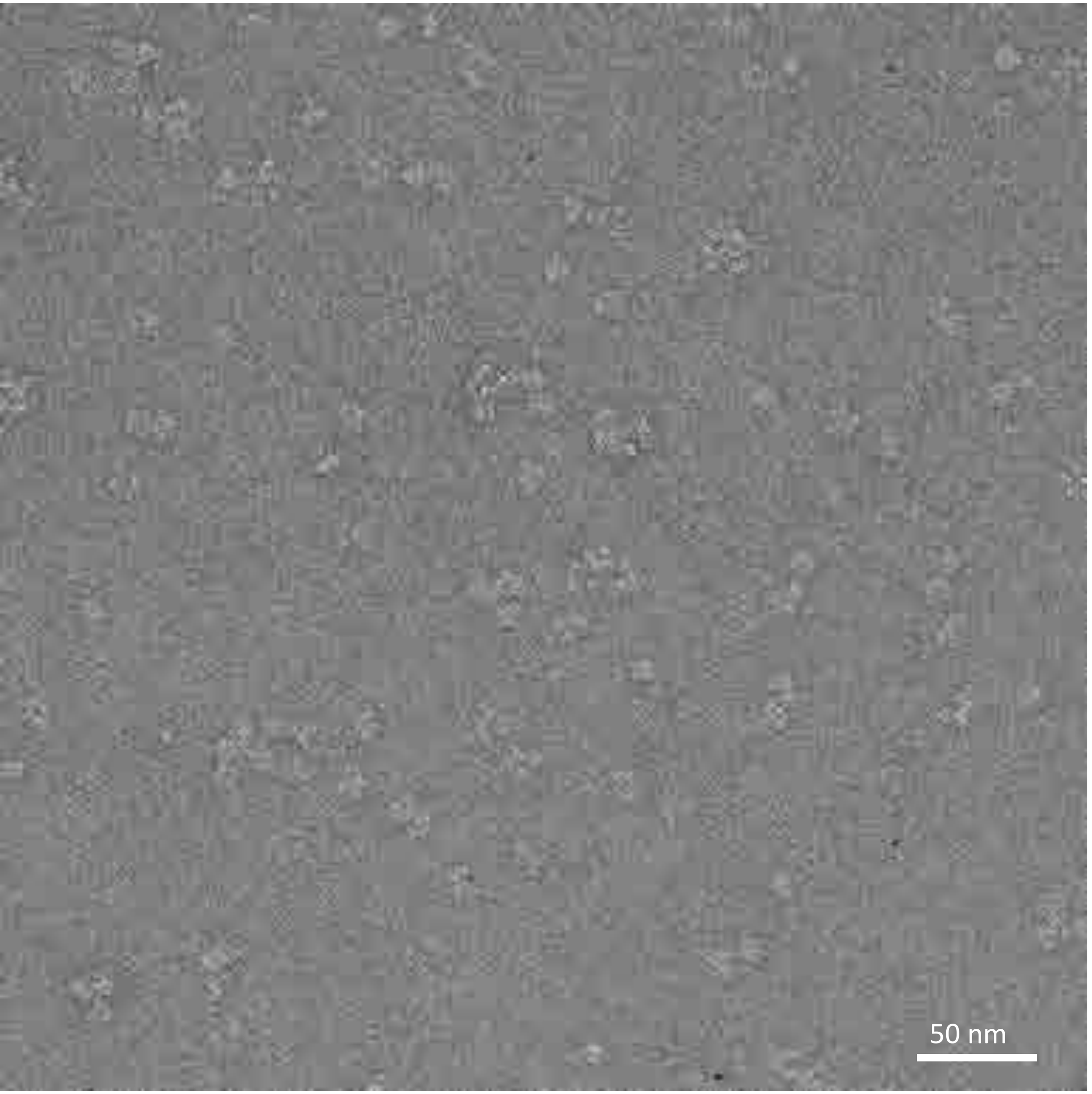


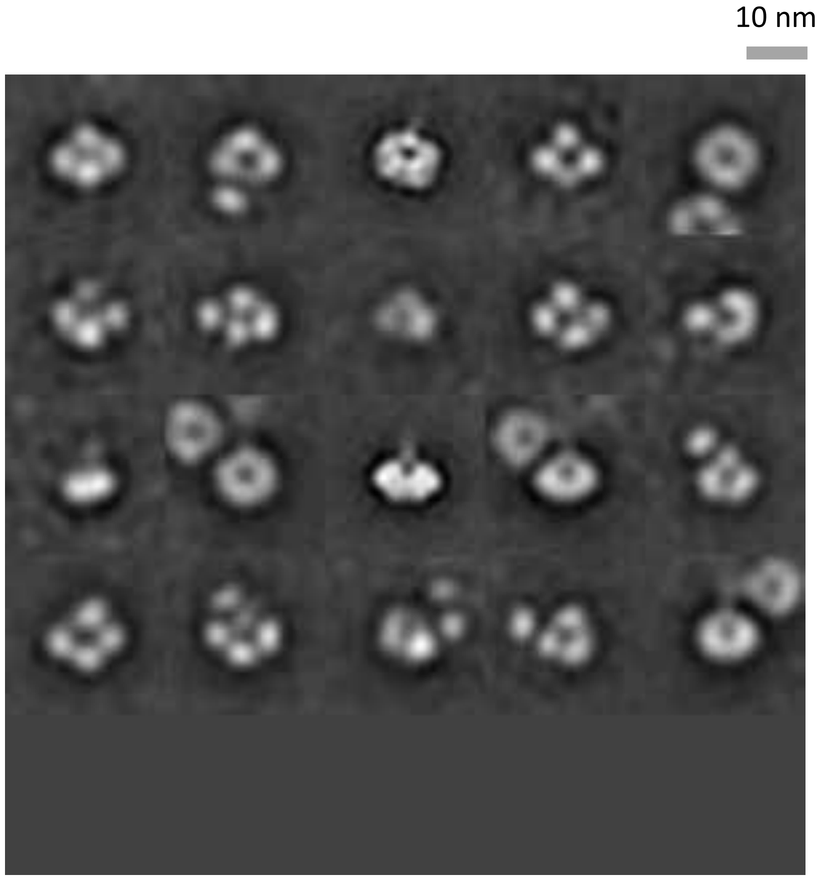

# N8-JD13-WT

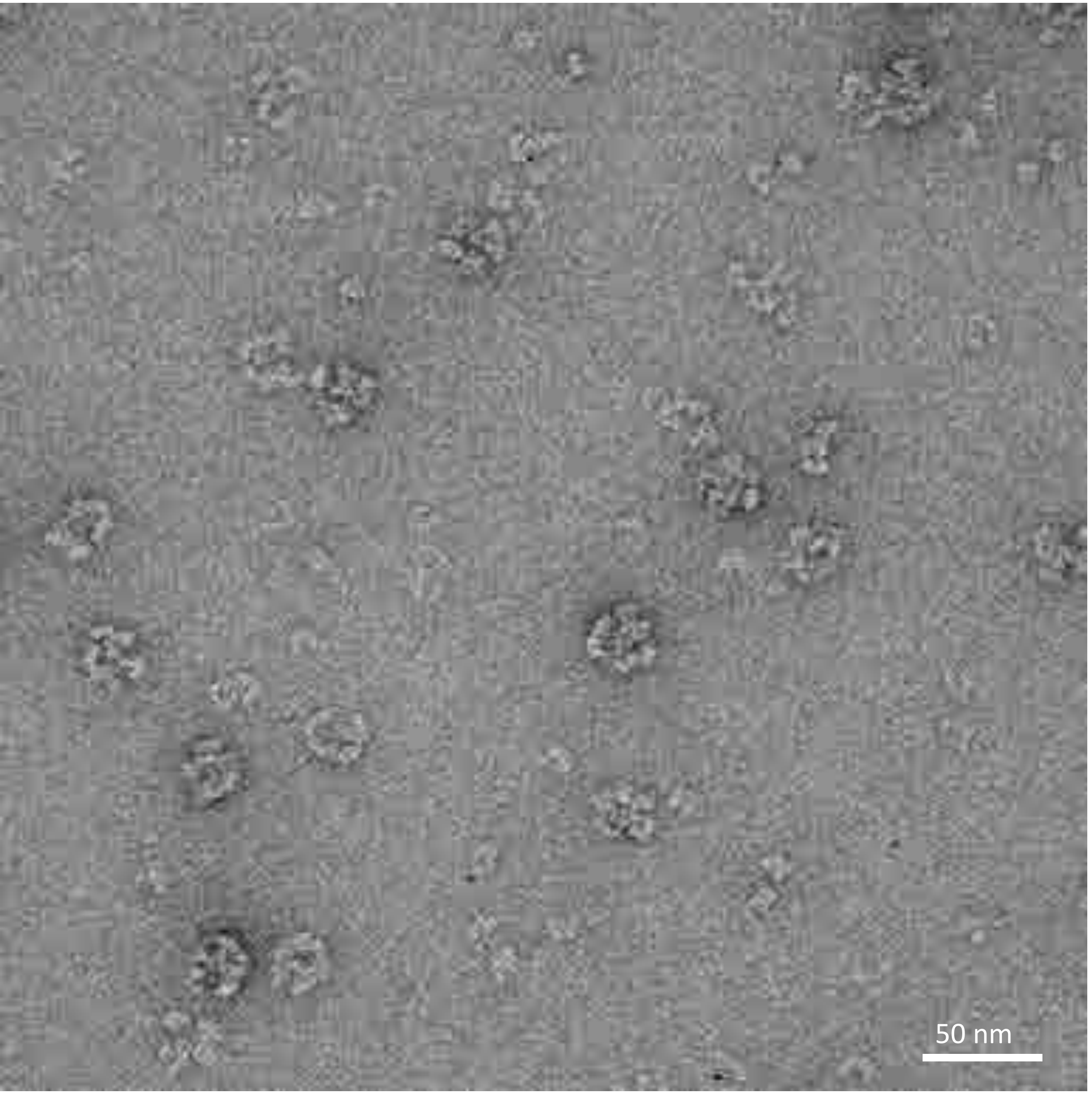


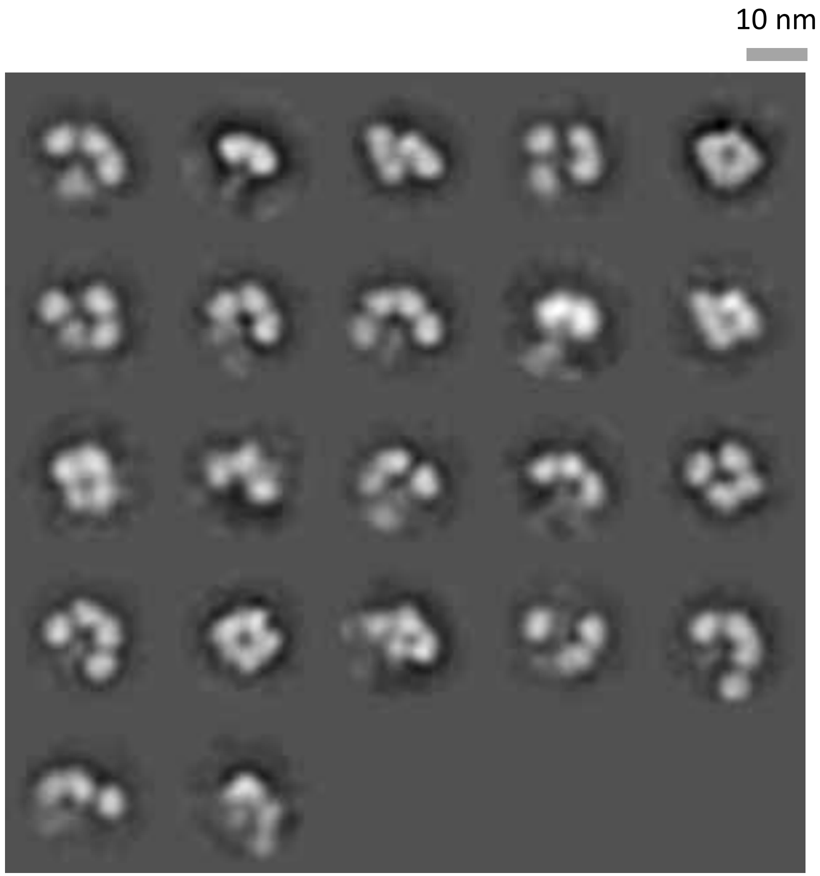

# N9-AN13-WT

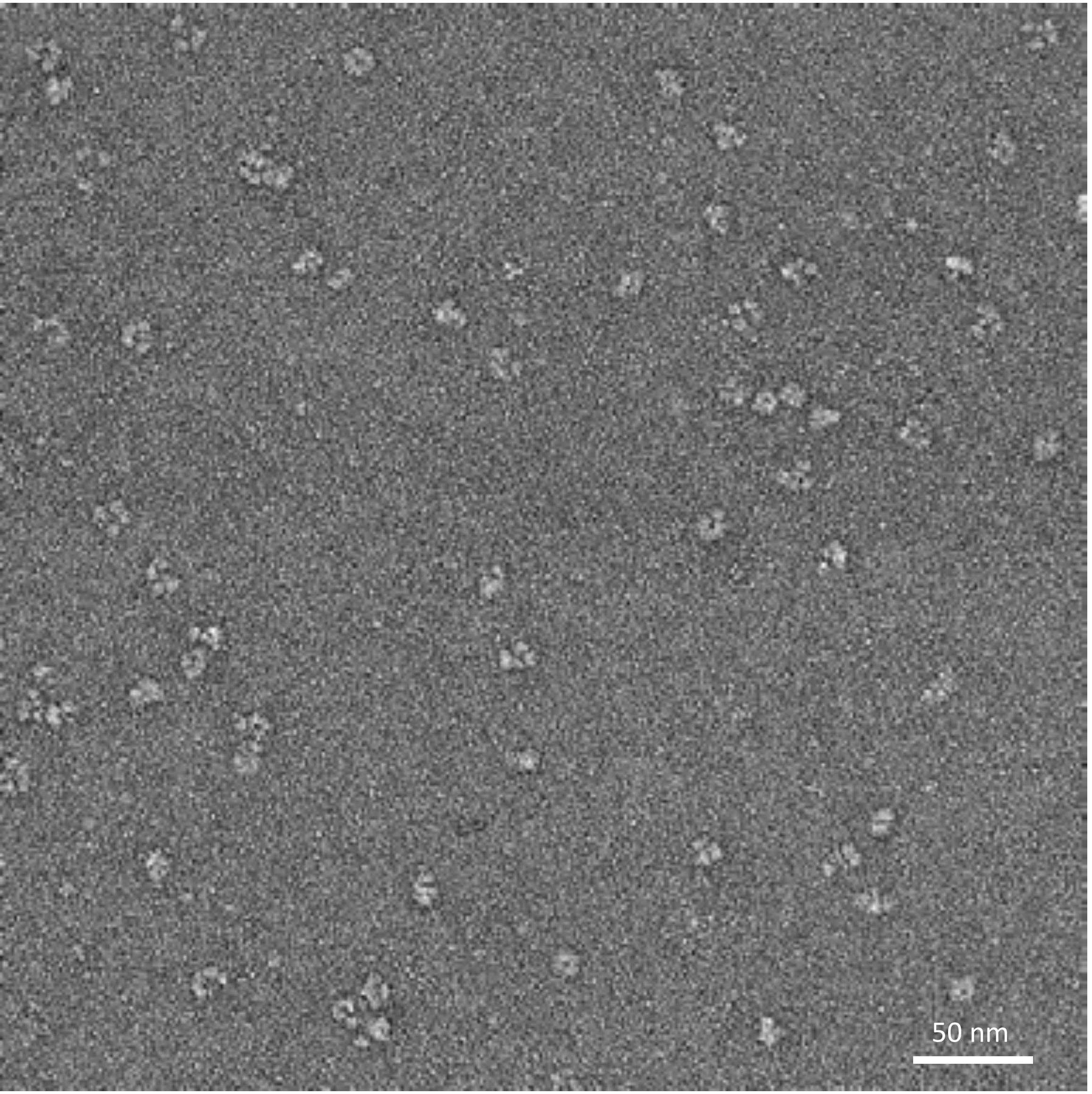


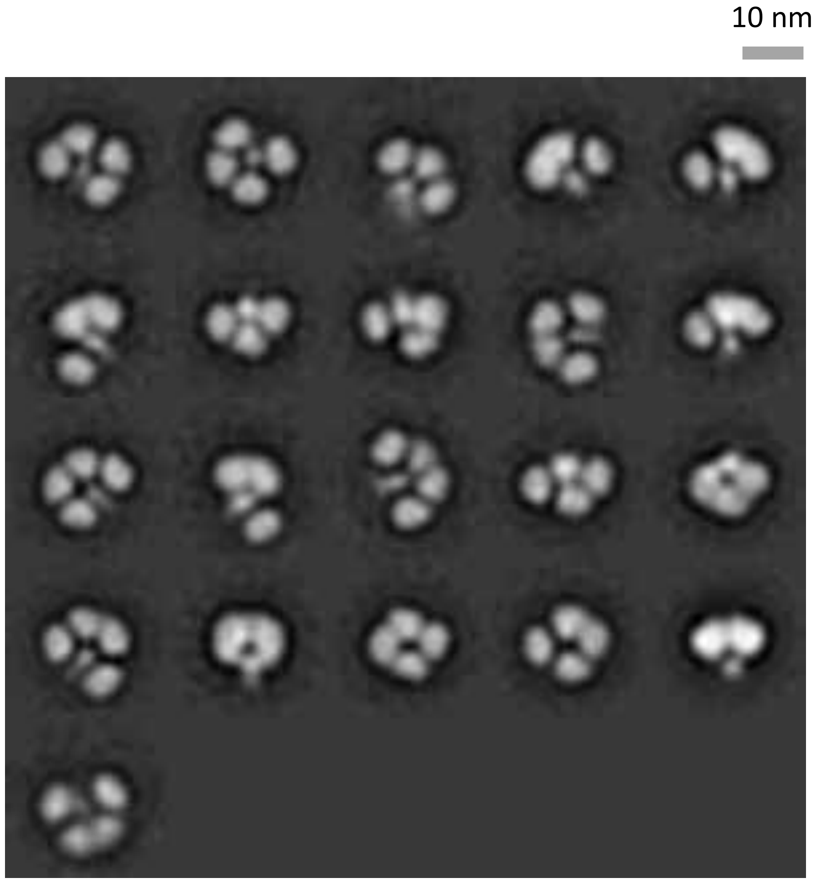

# N2-WI05-WT

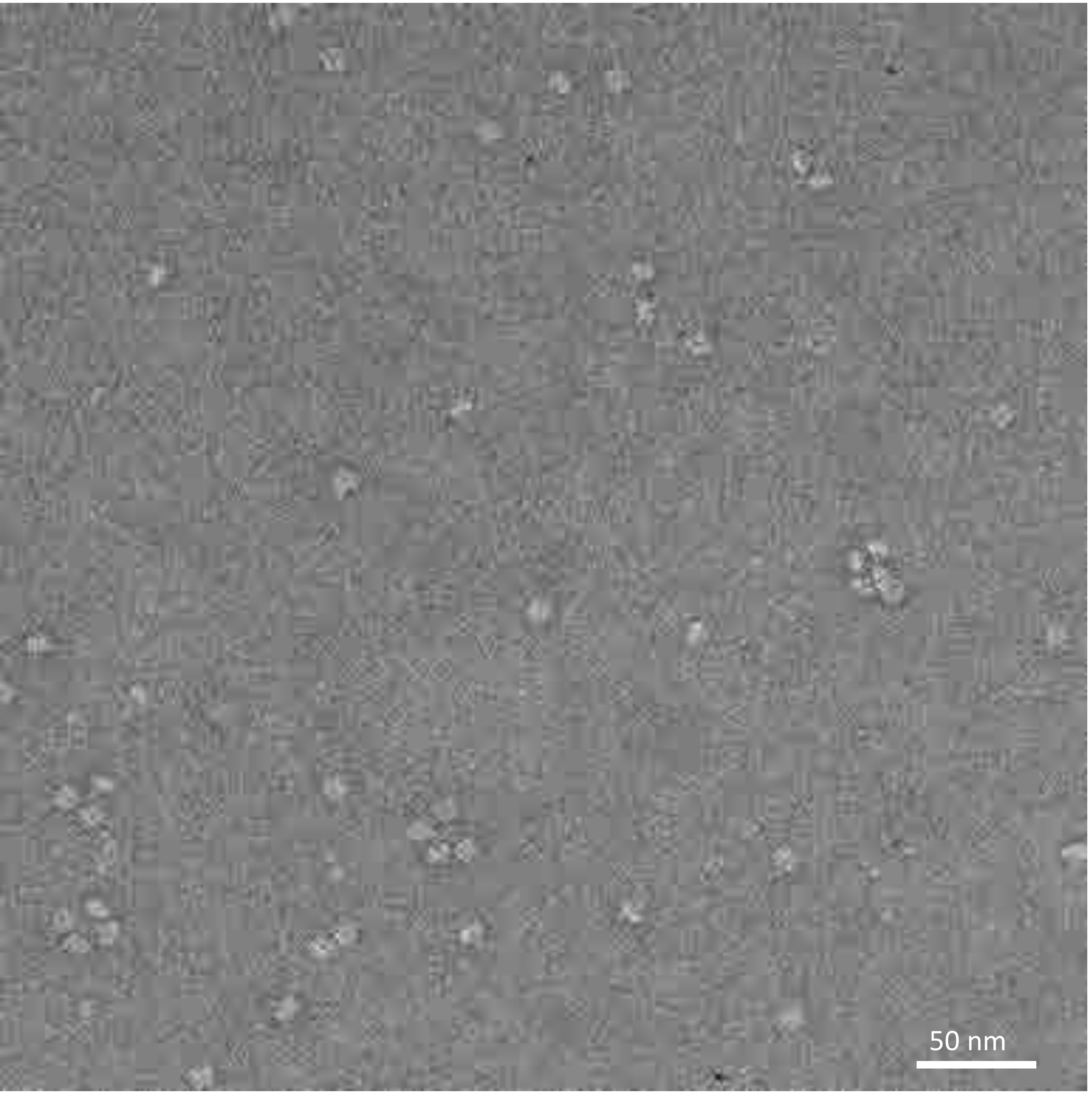


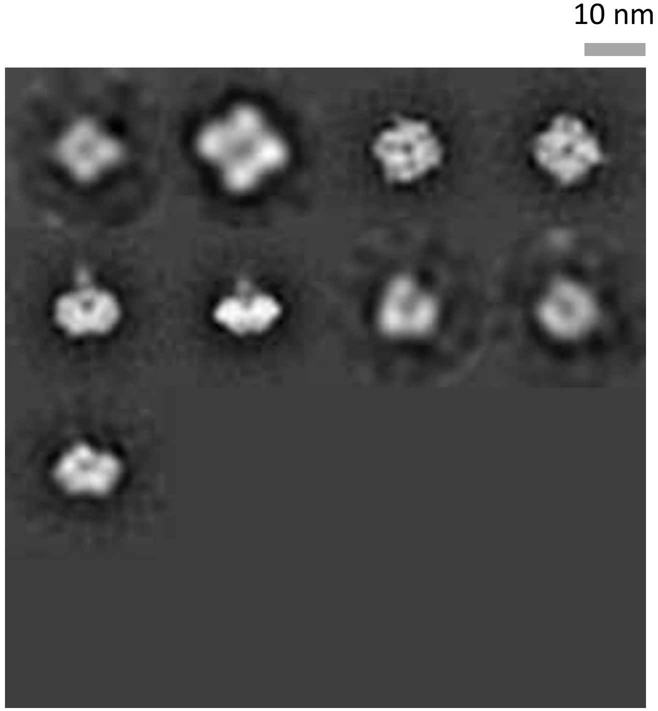

# N2-IN11-WT

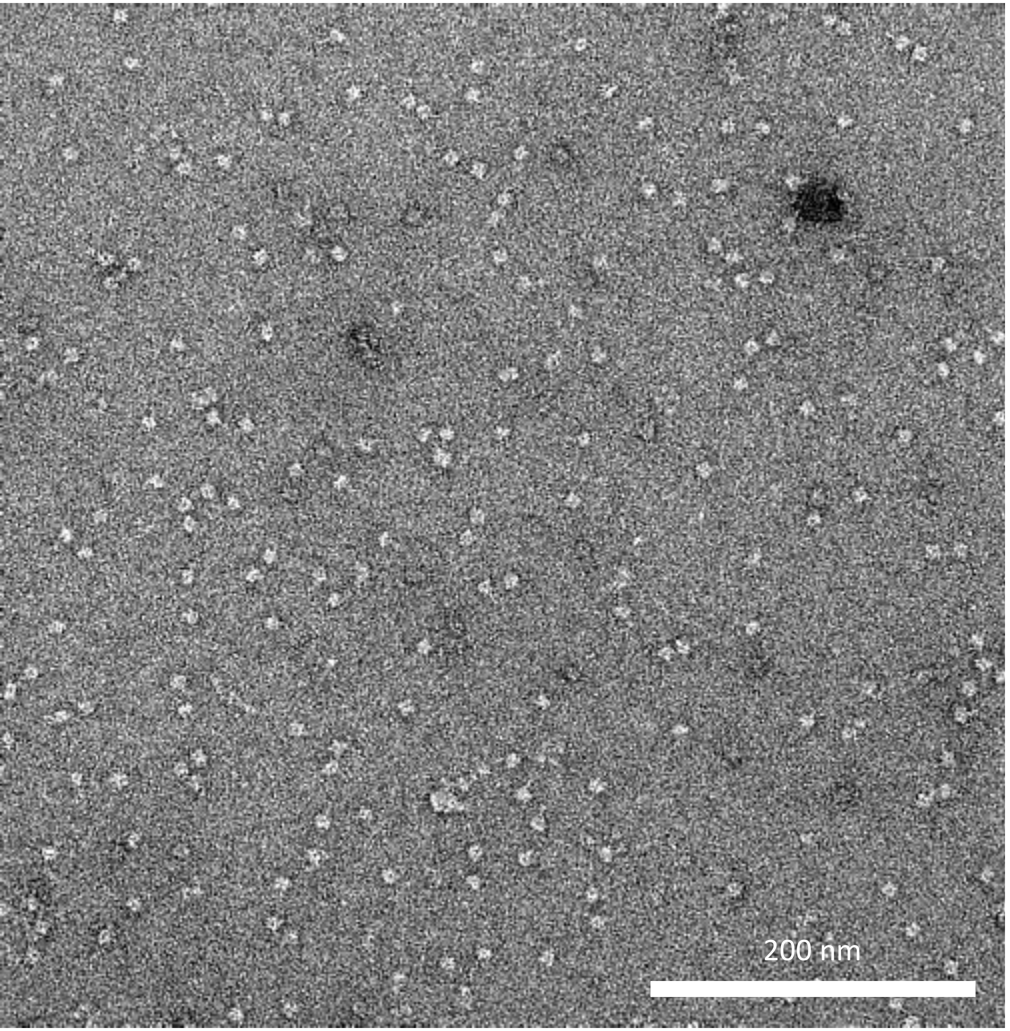


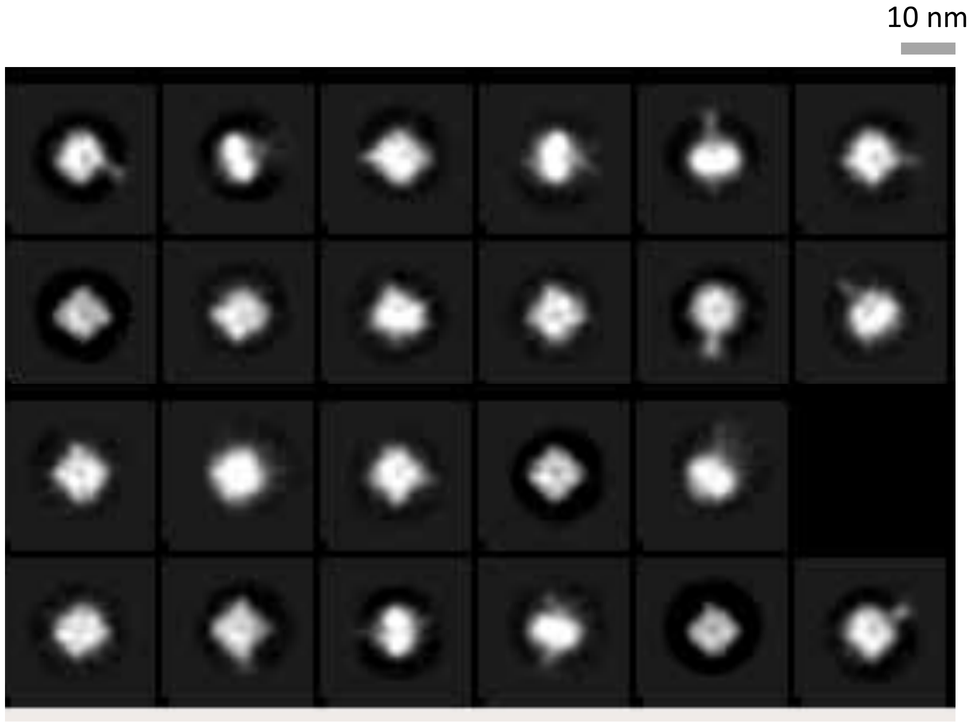

# N3-MI06-WT

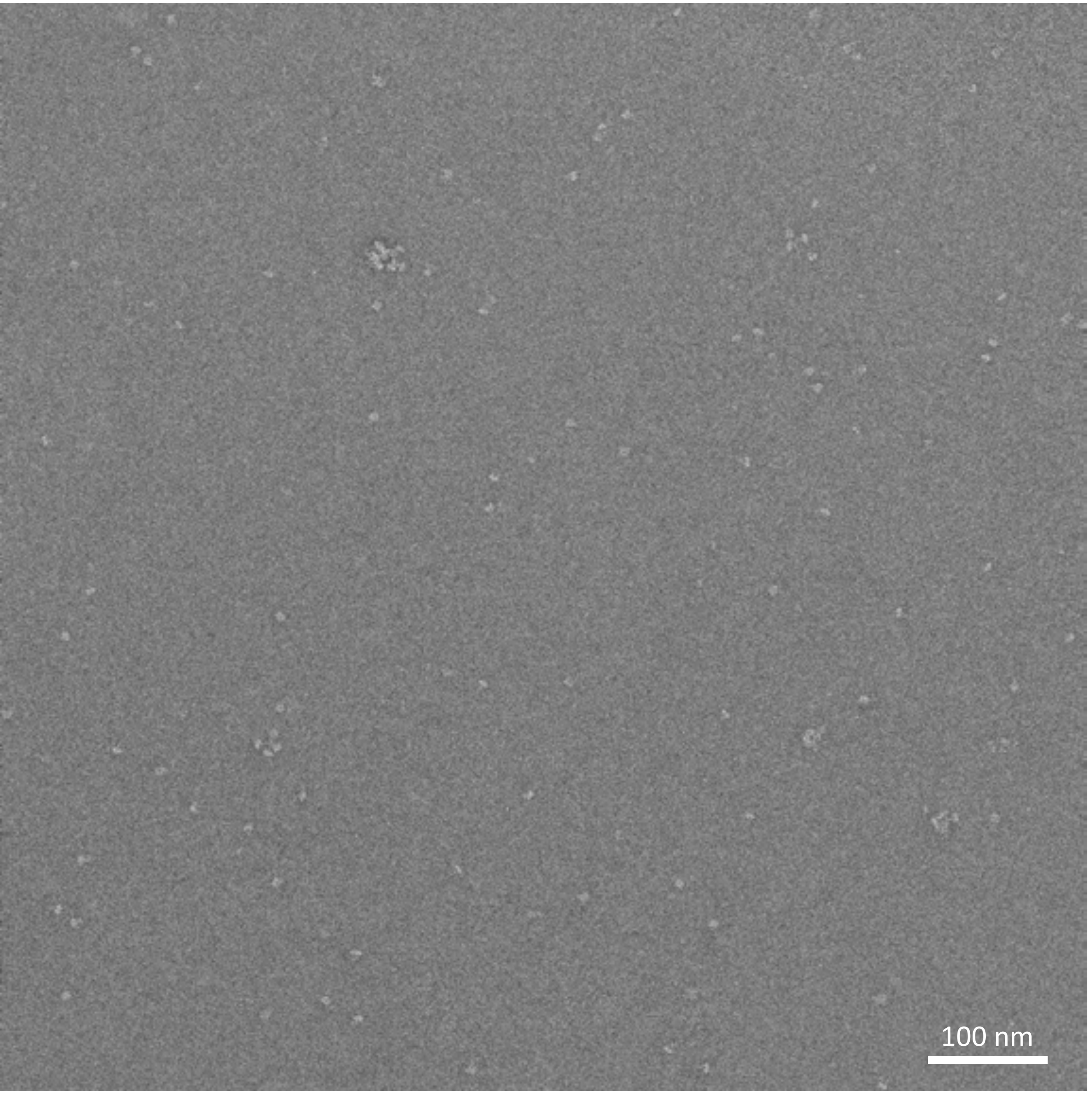


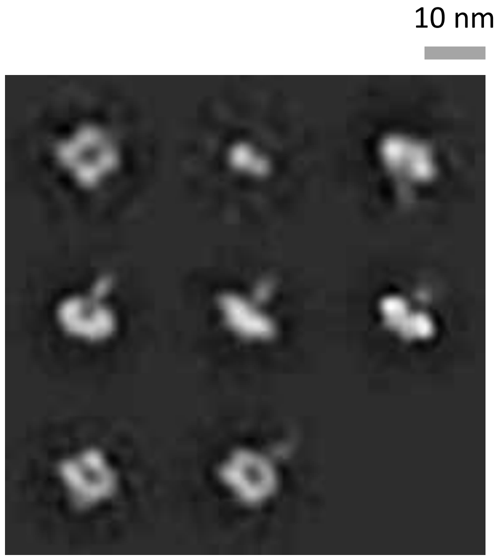

# N5-DB16-WT

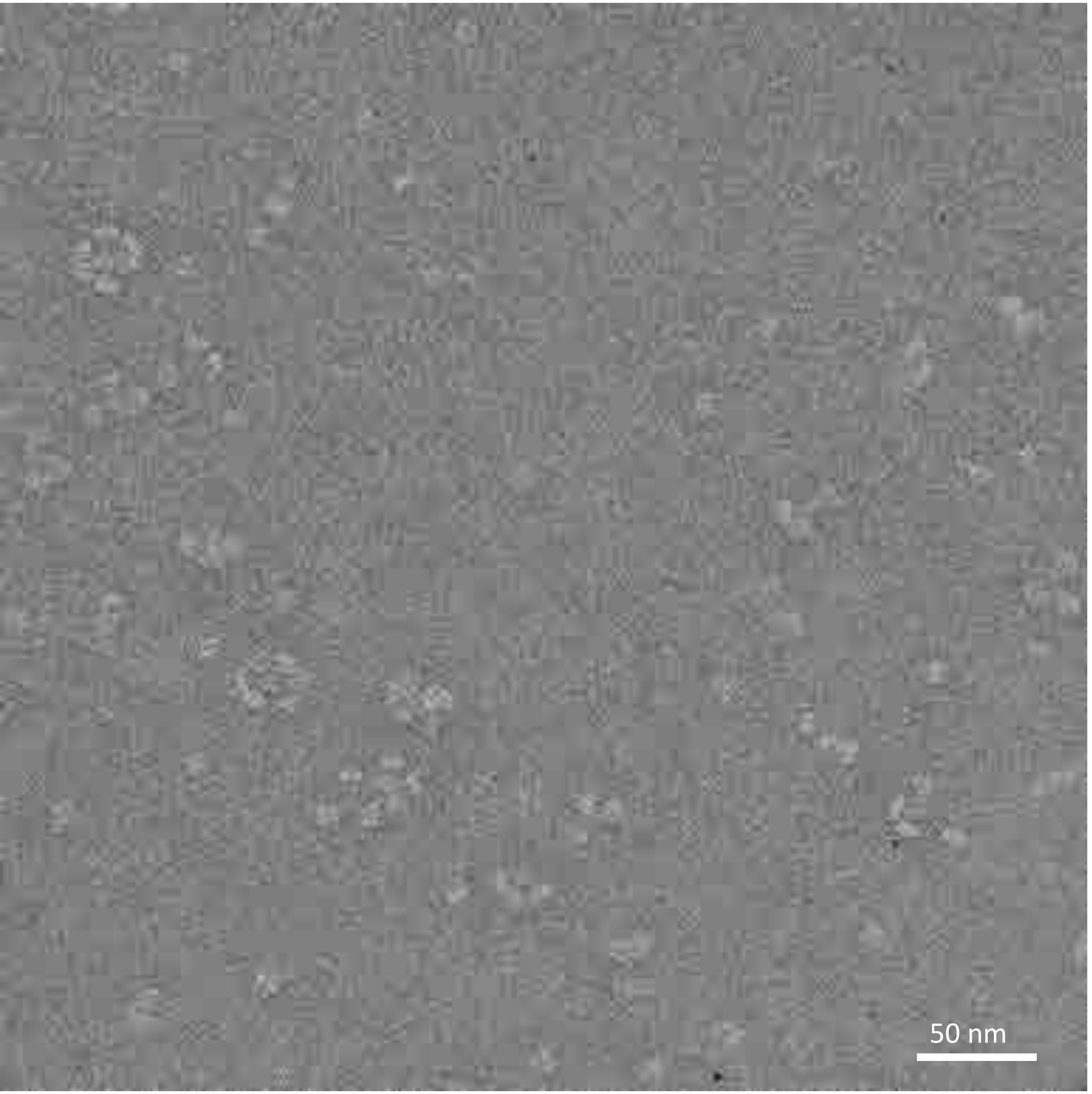


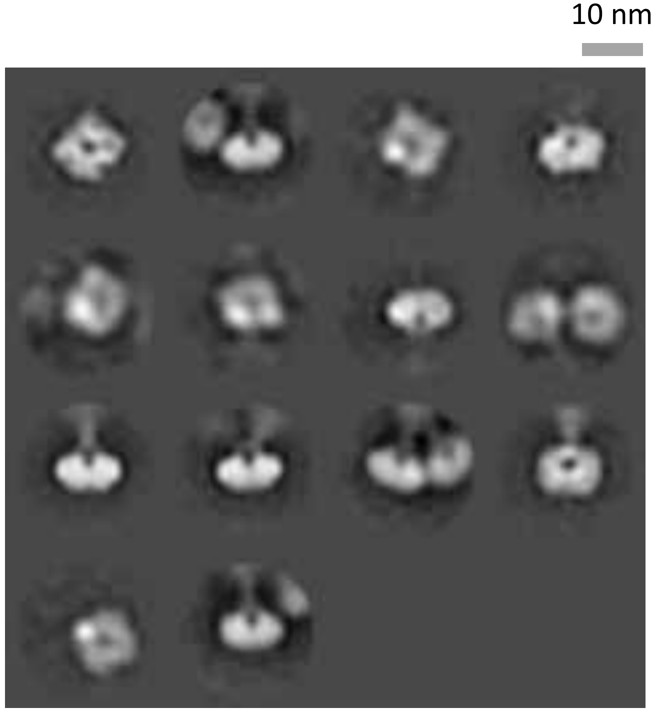

# B-CO17-WT

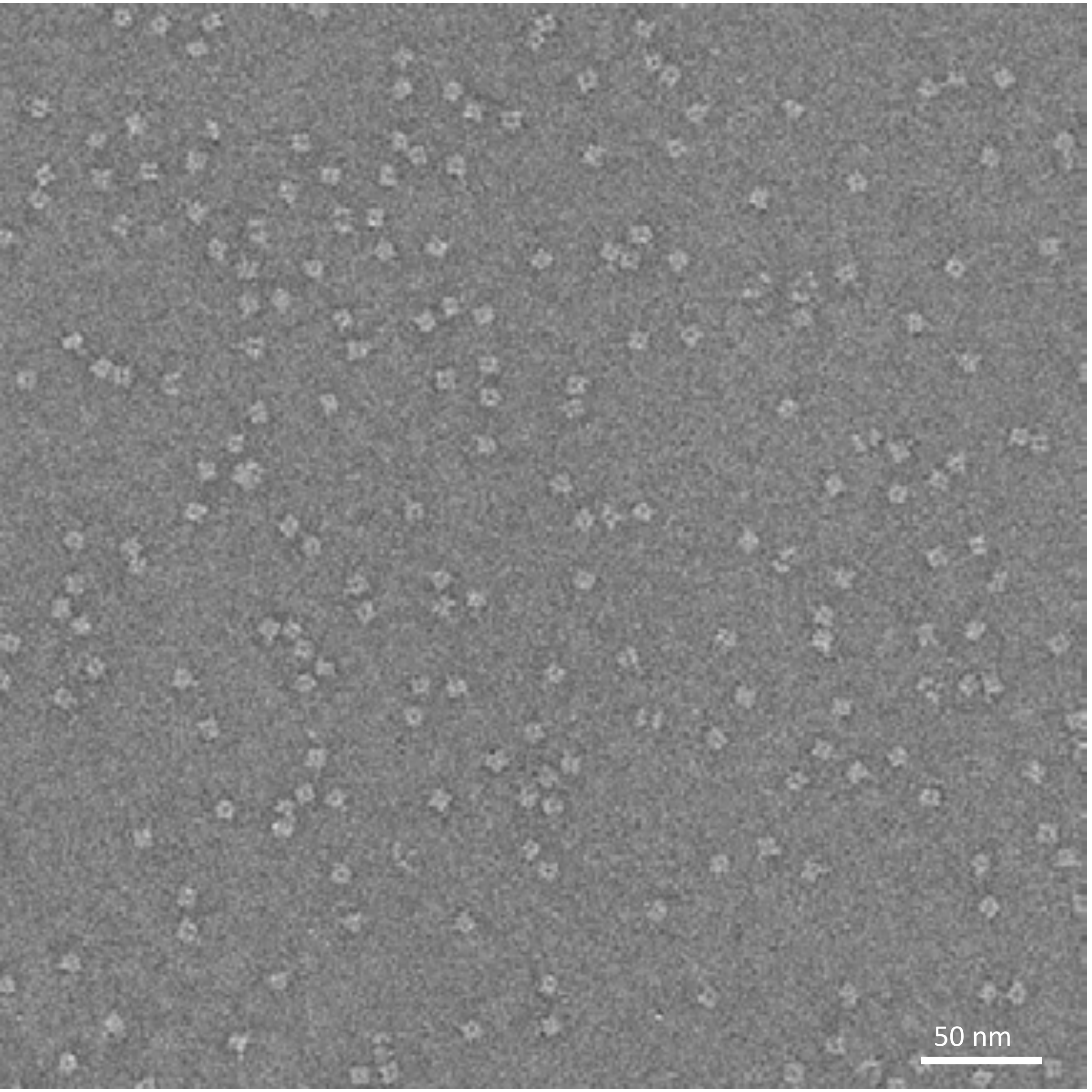


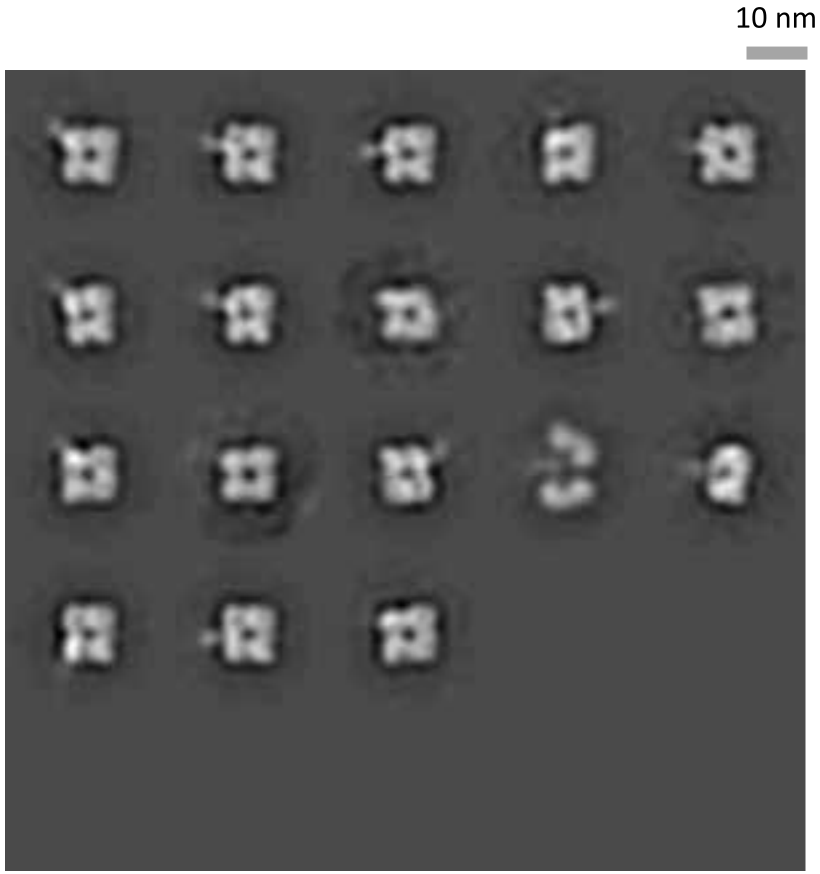

# B-PH13-WT

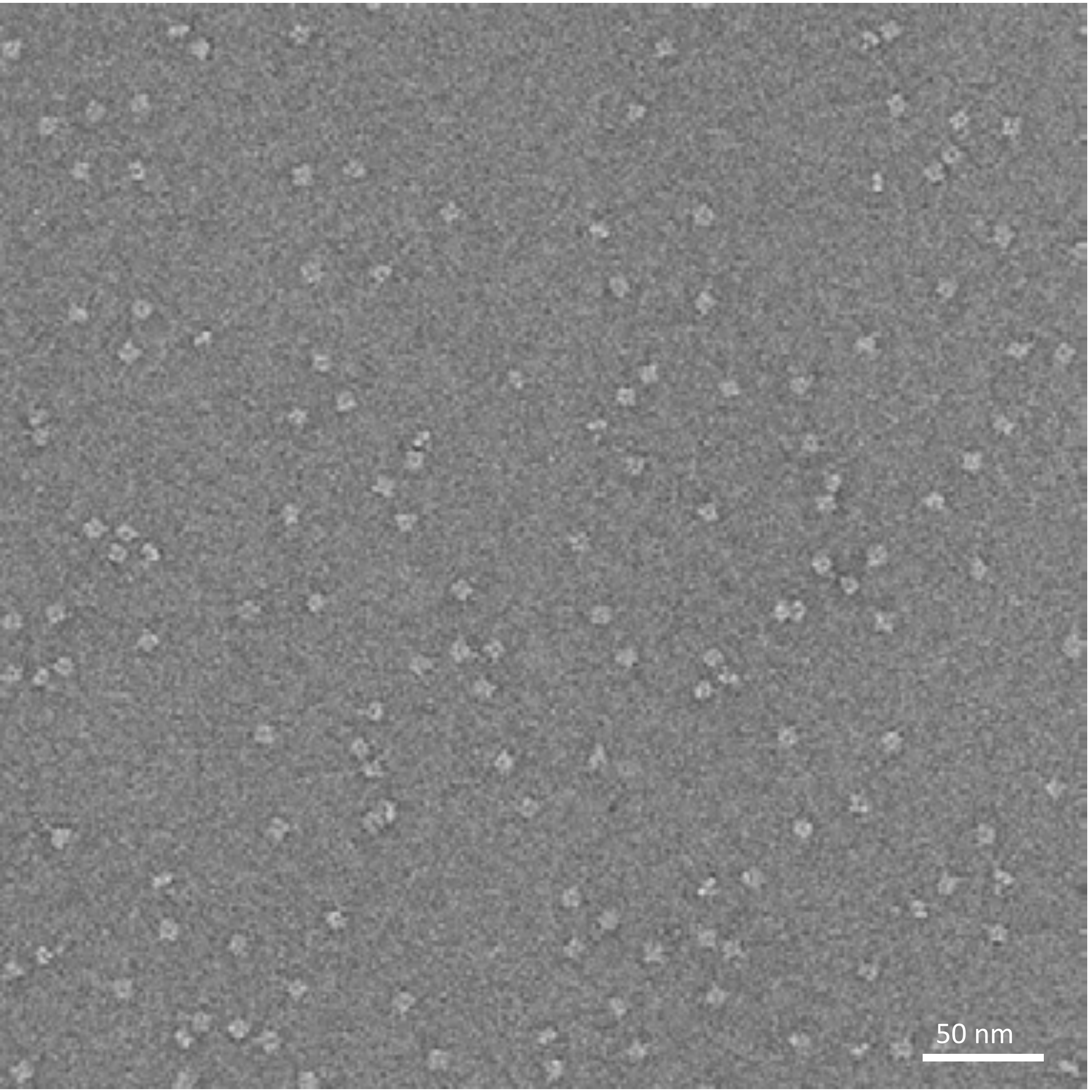


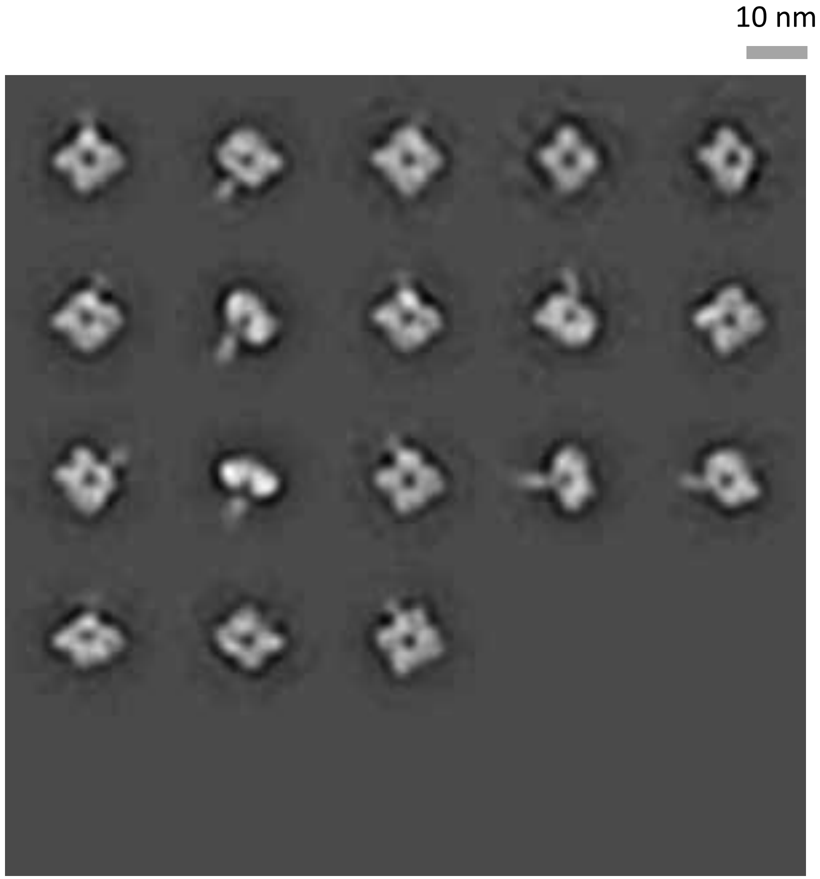

# N9-AN13-Y169aH

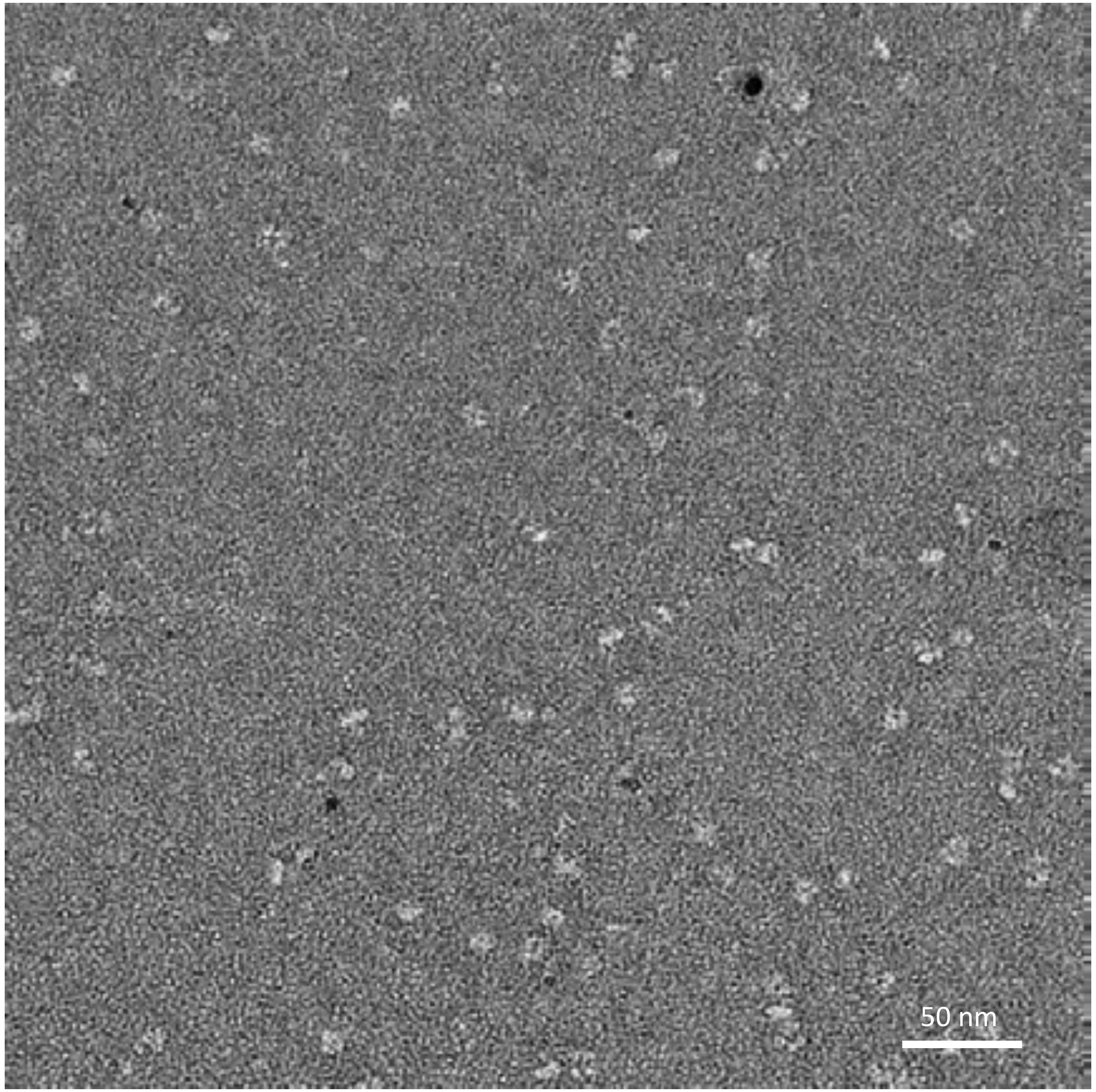


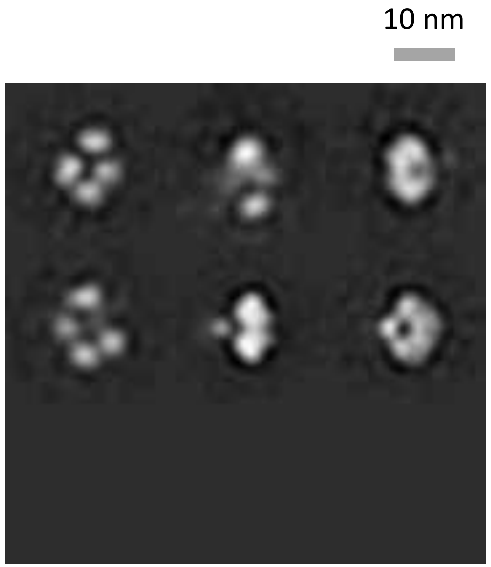

### N2-WI05-WT, obtained from BEI Resources

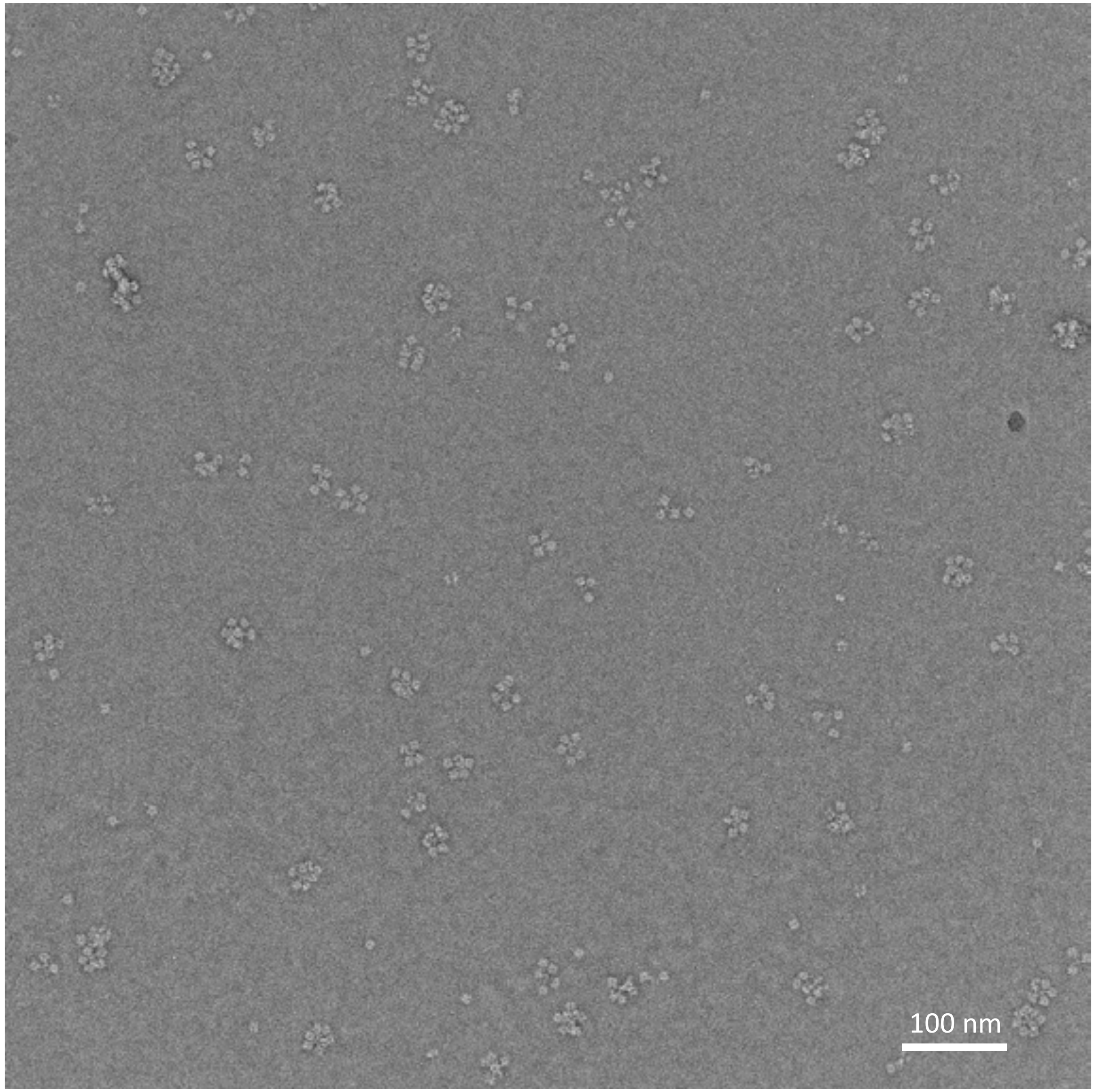


10 nm

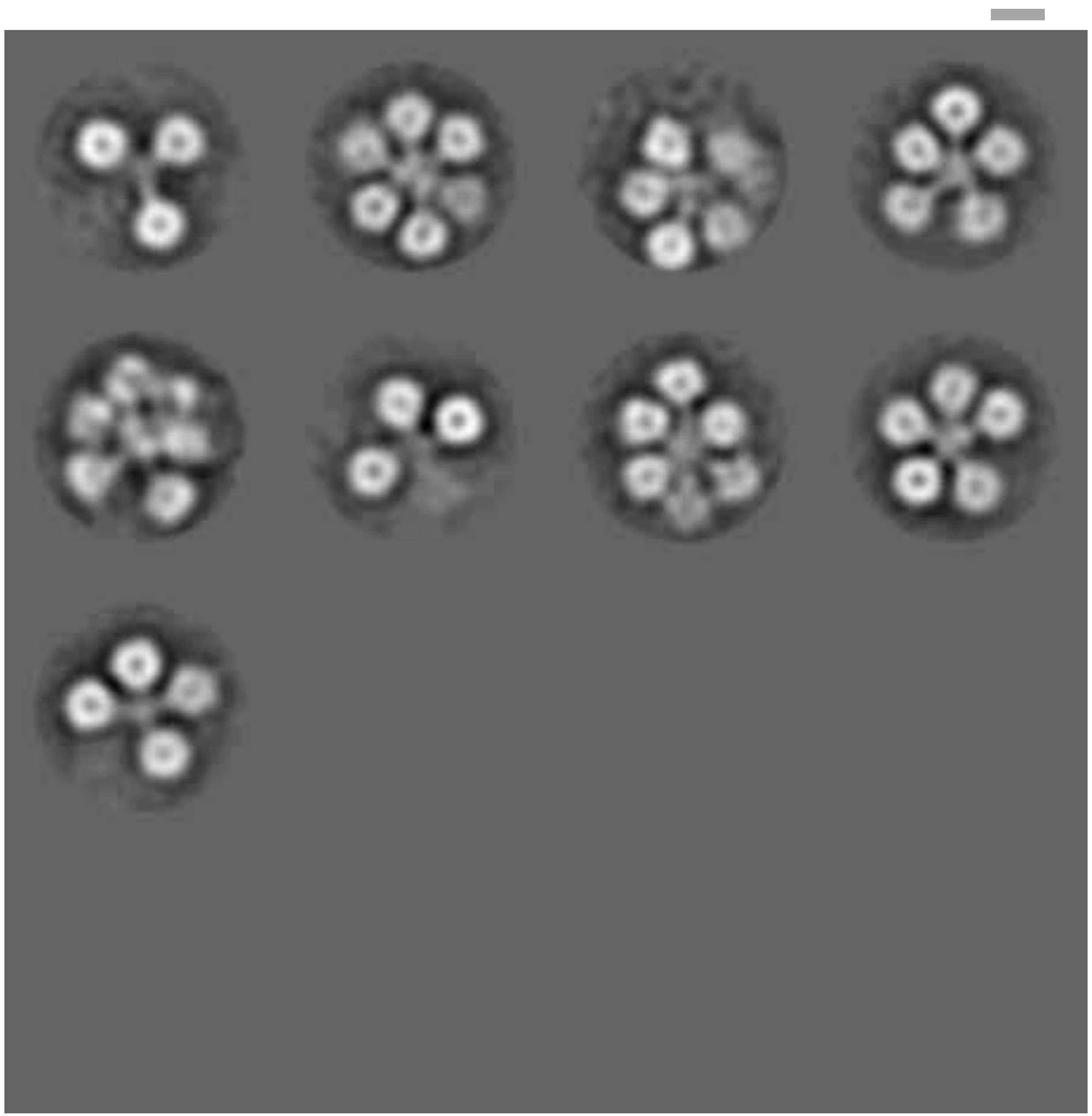

### N1-NC99-WT, obtained from BEI Resources

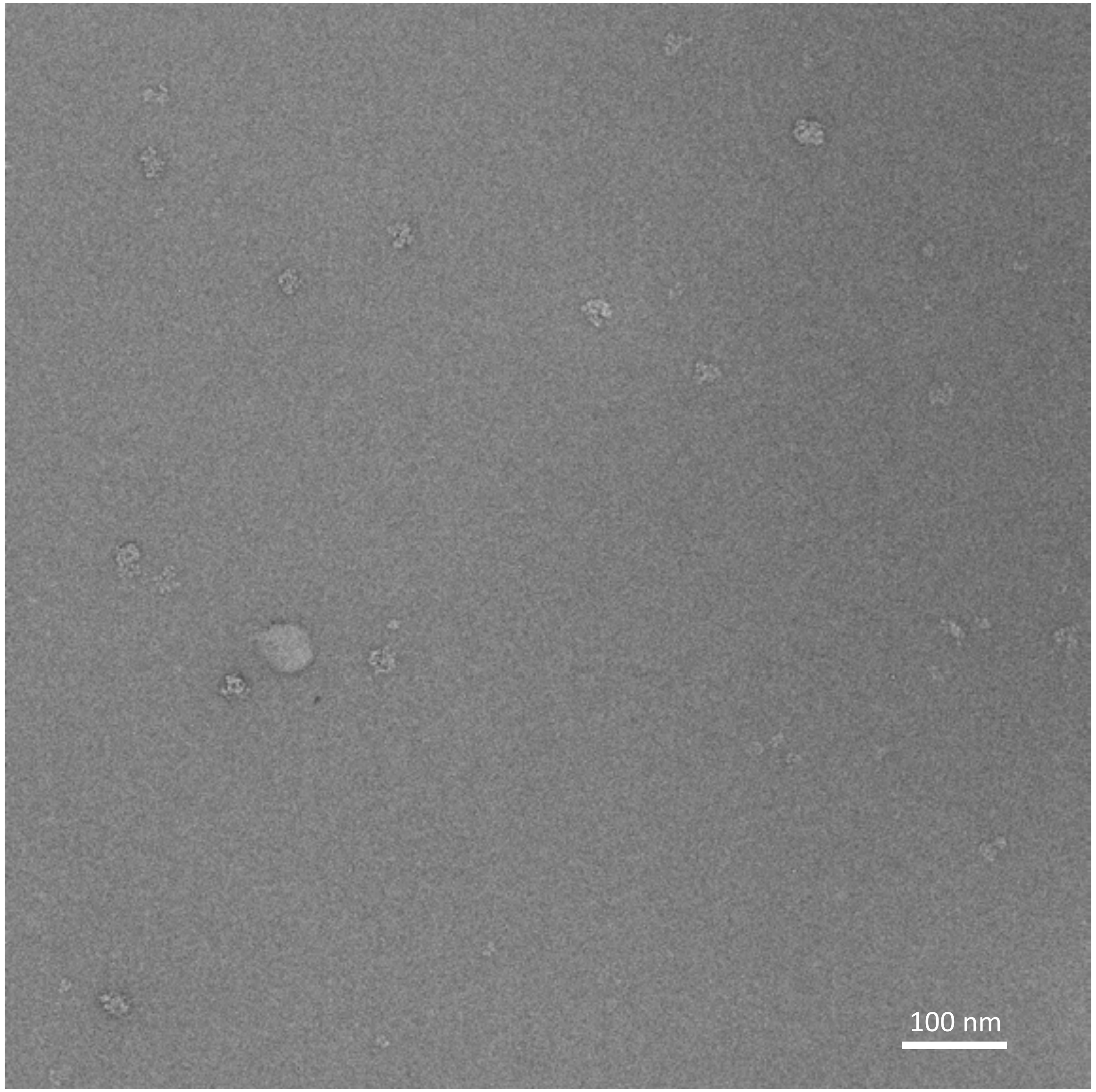


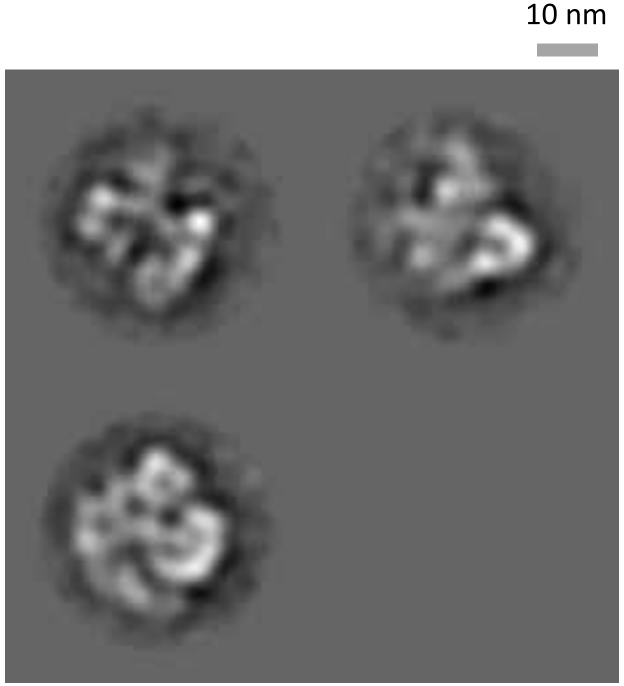

### N1-CA09-WT, obtained from BEI Resources

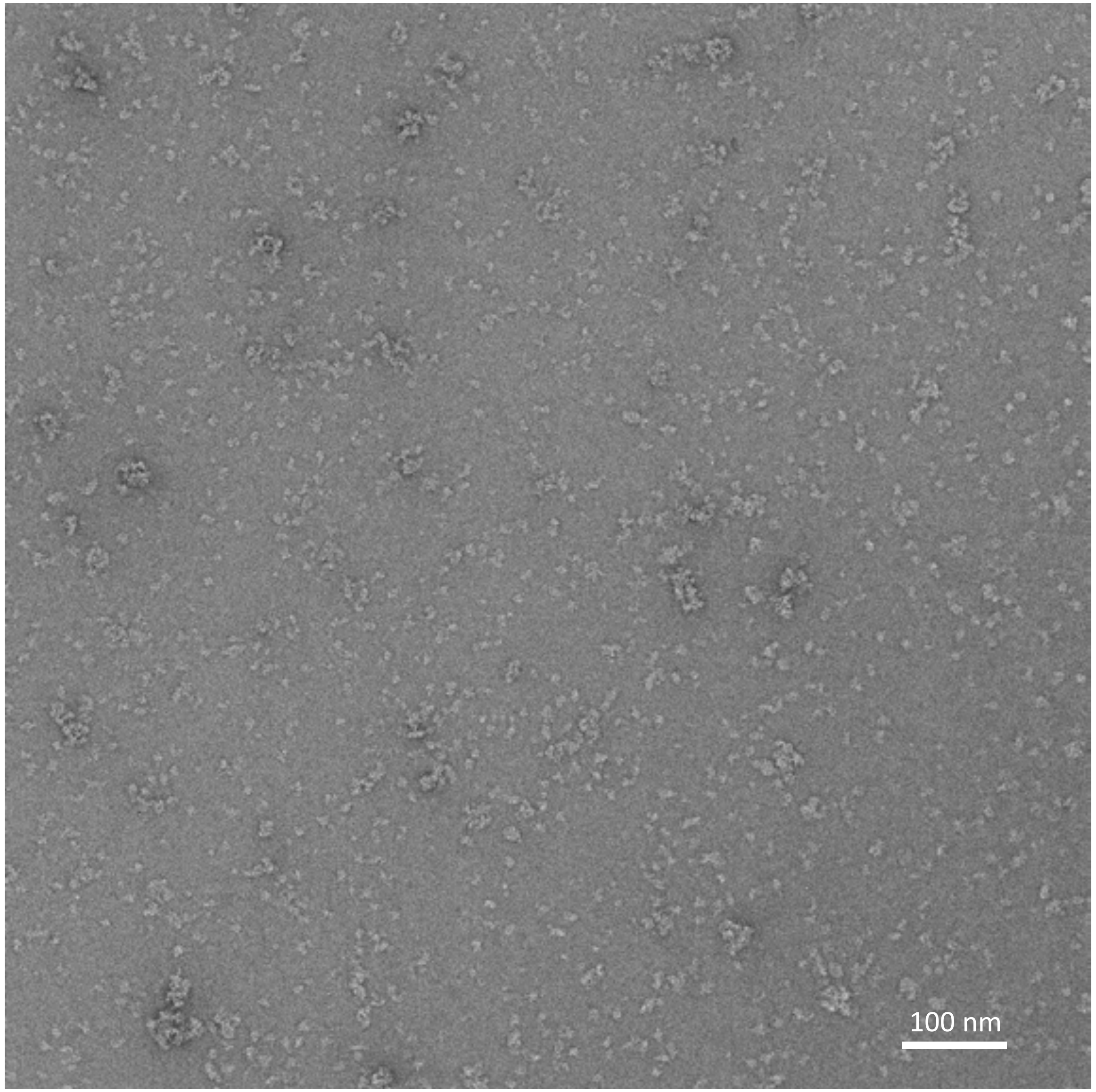


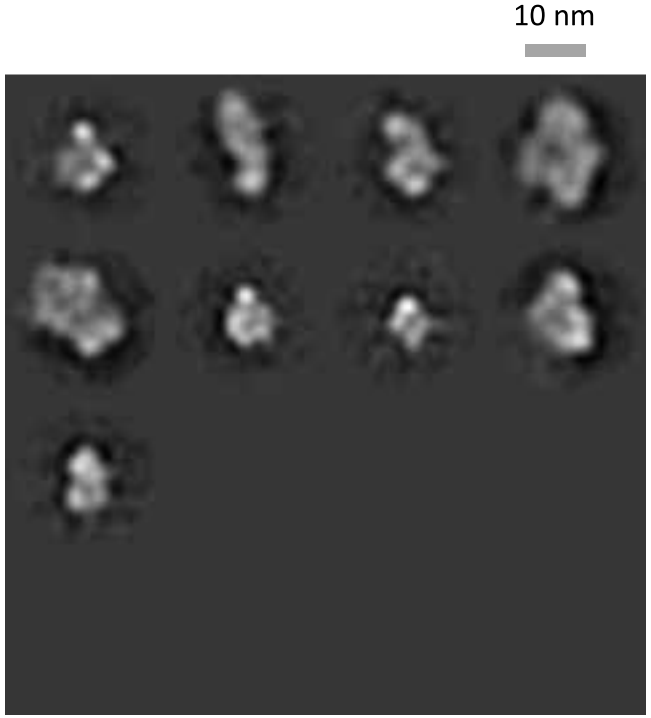

### N1-CA09-WT with 1mM $CaCl_2$

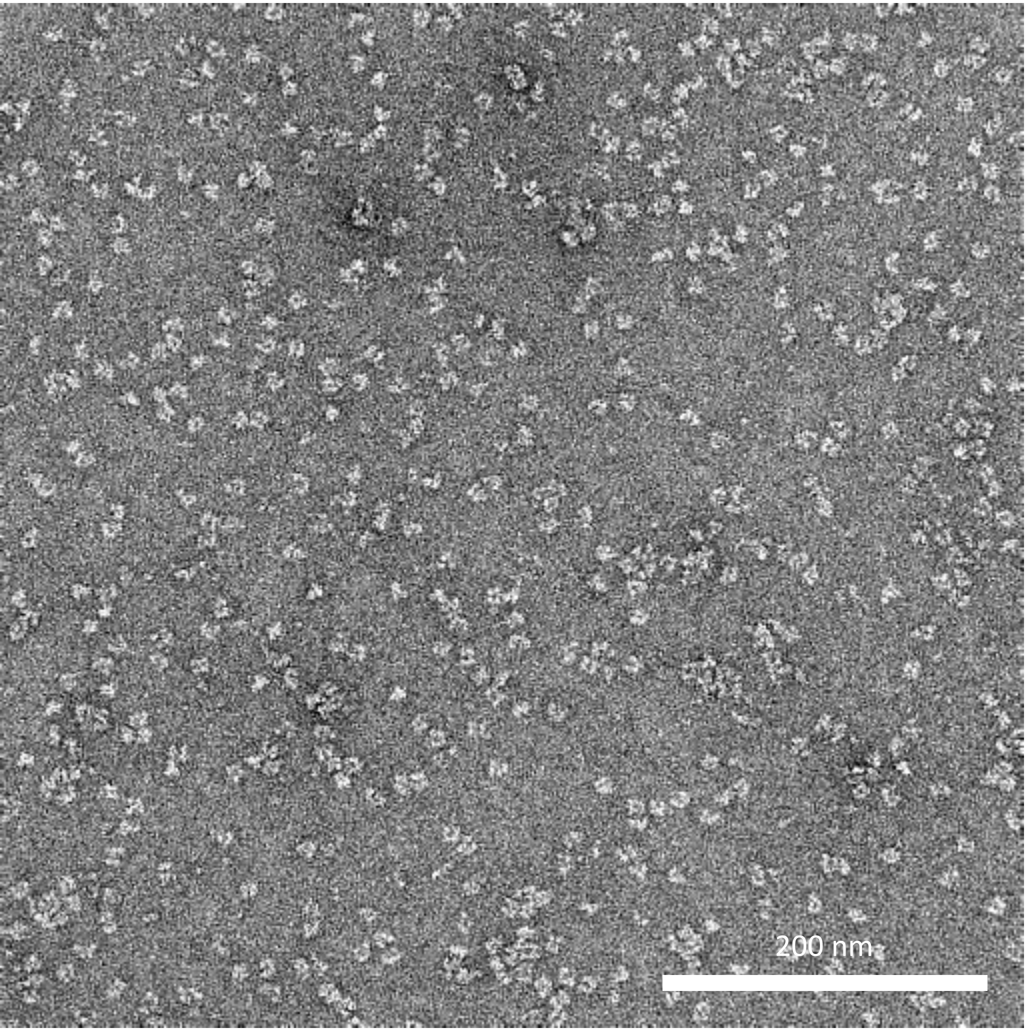


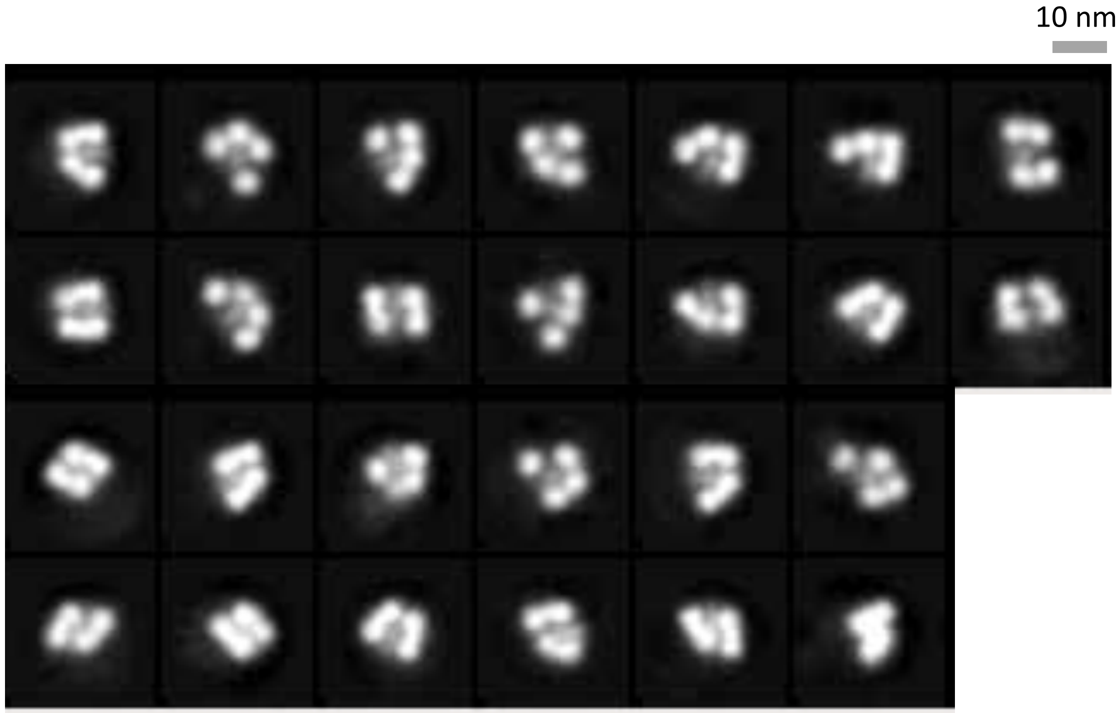

### N1-CA09-sNAp-94

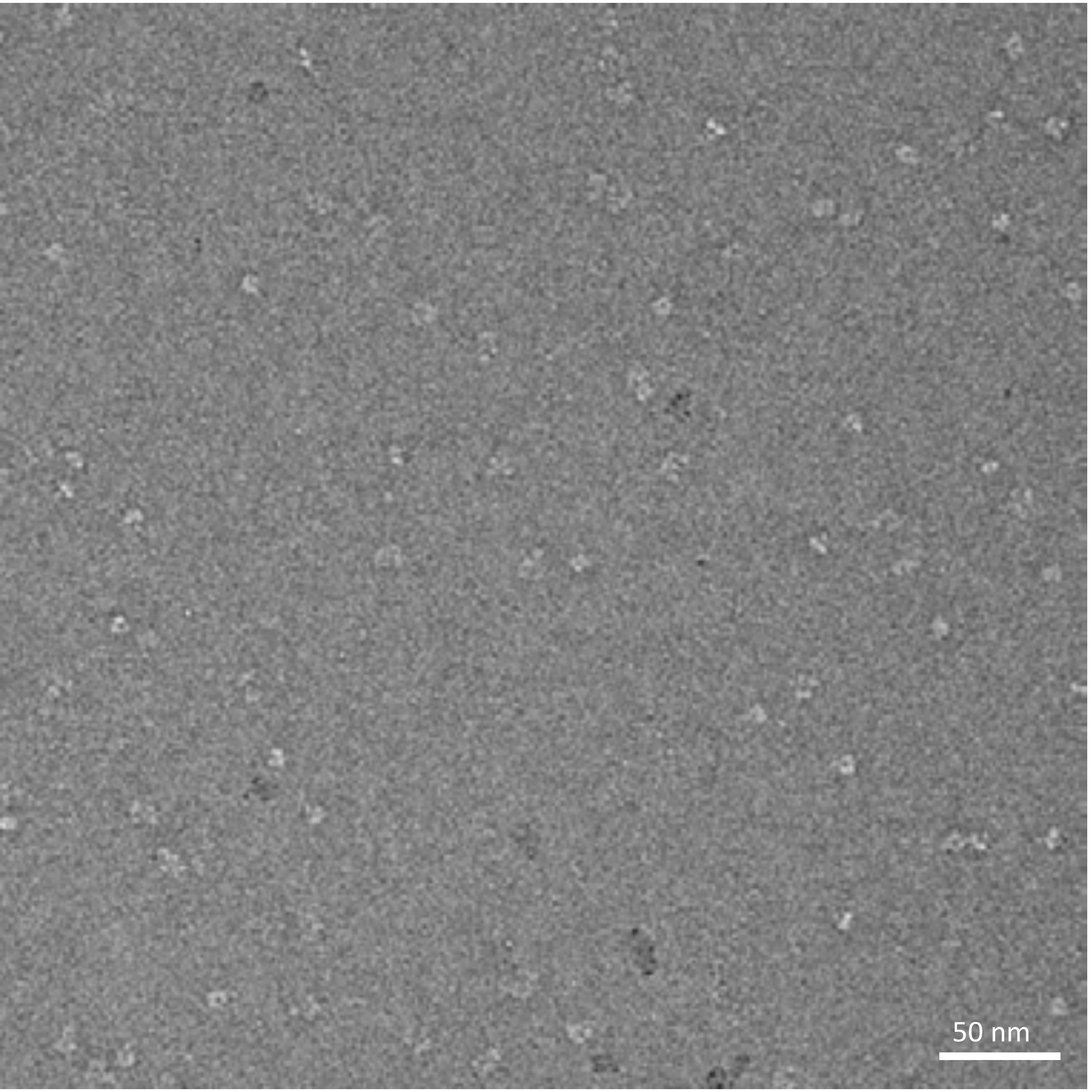


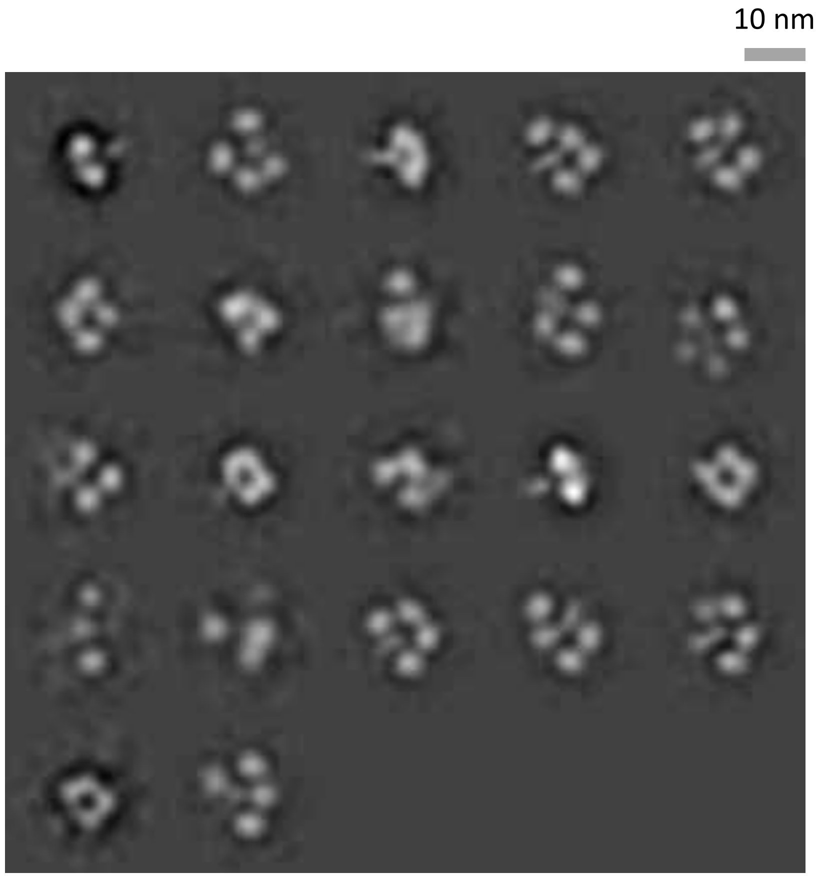

### N1-CA09-sNAp-114

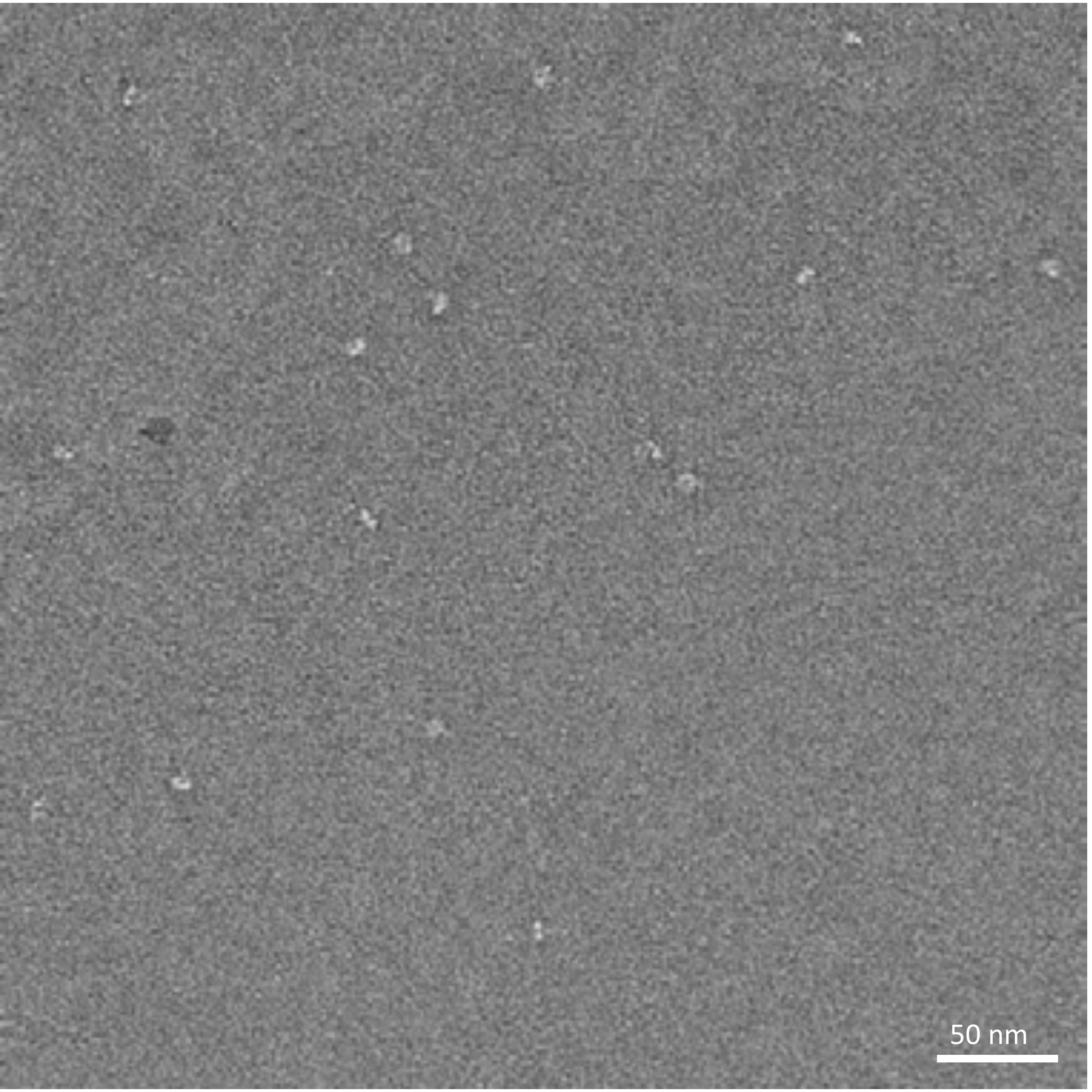


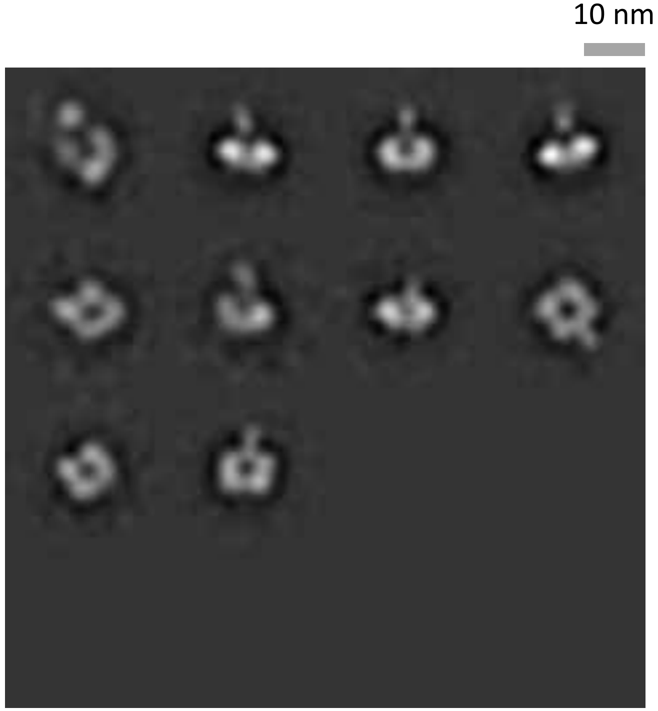

### N1-CA09-sNAp-130

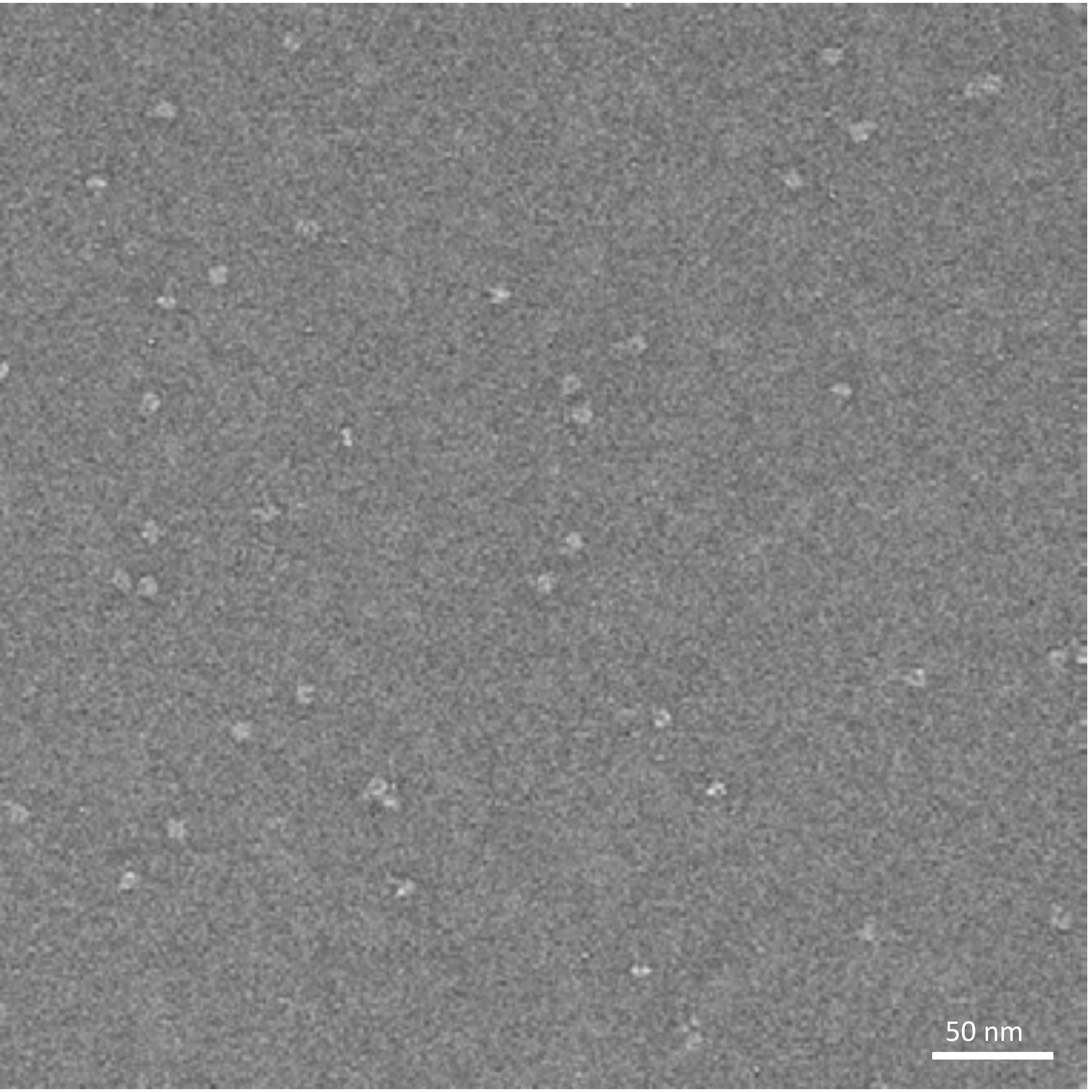


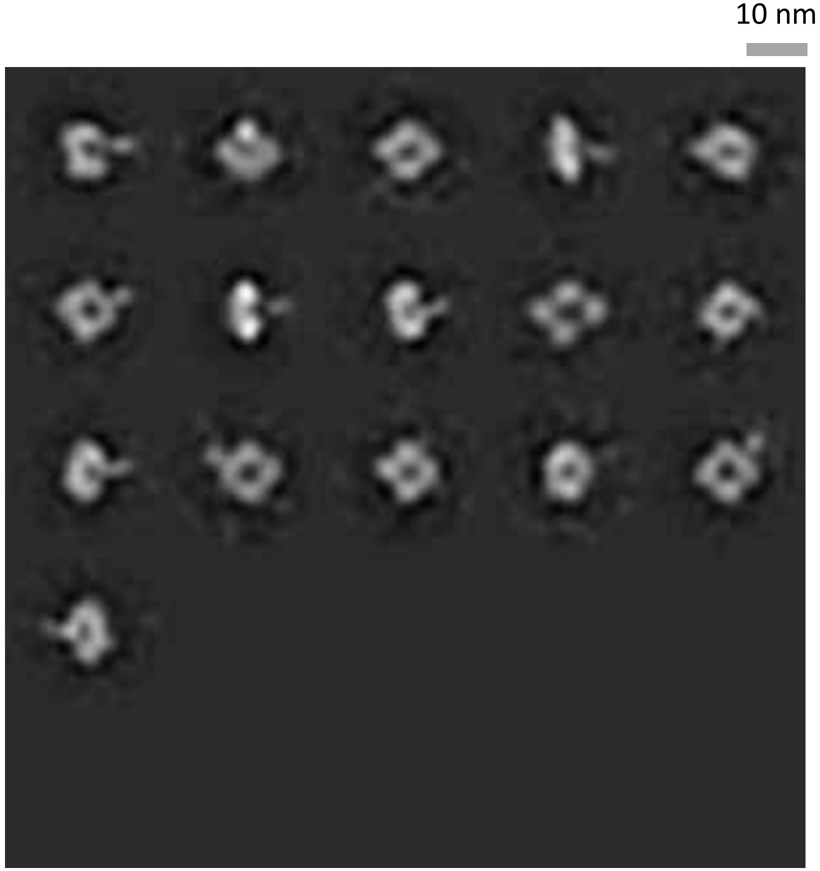

### N1-CA09-sNAp-155

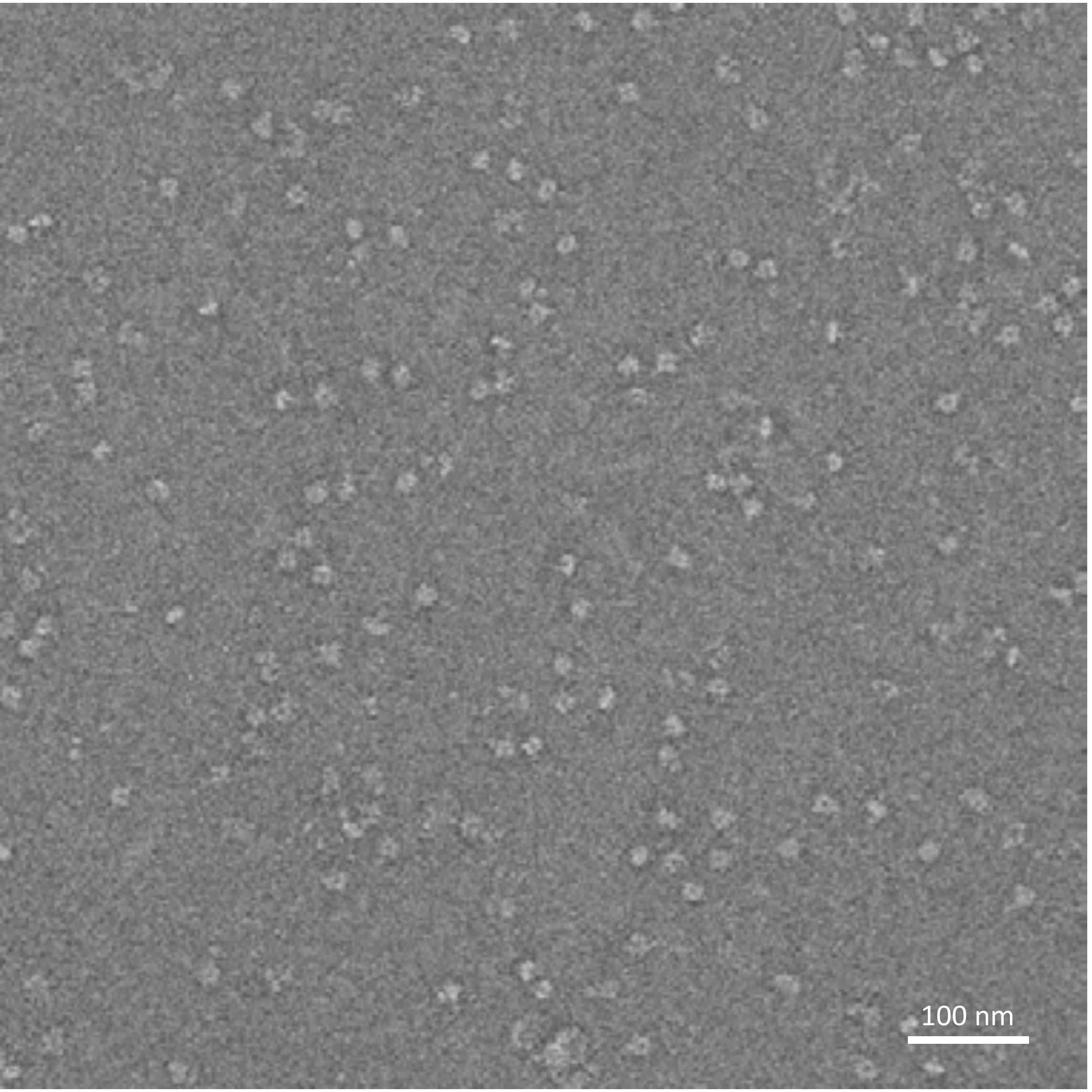


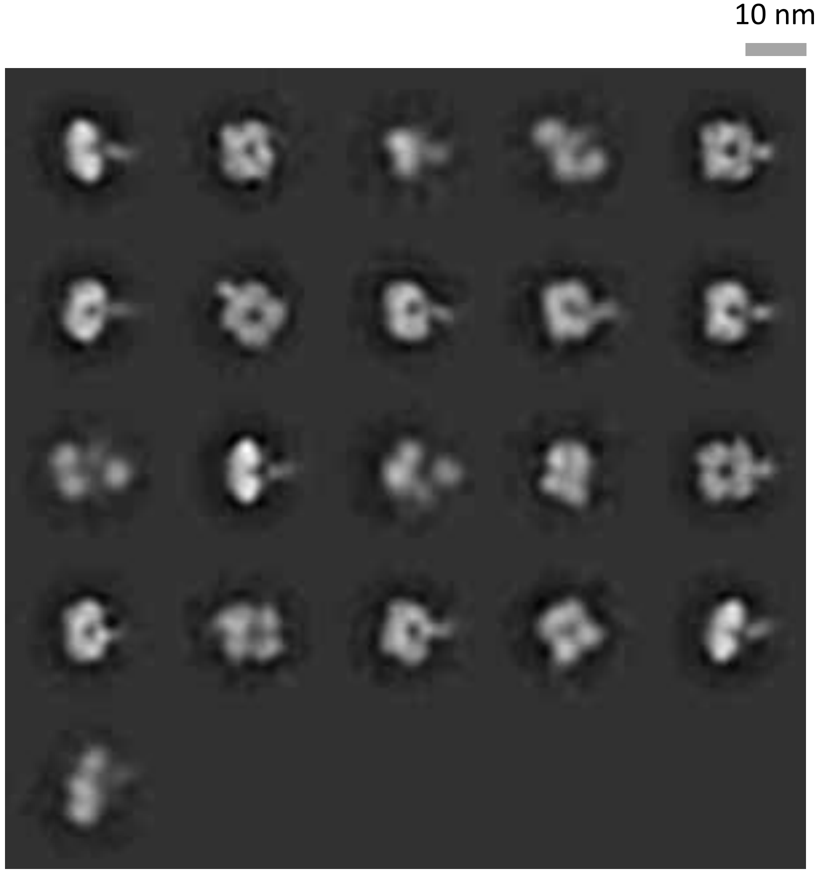

### N1-CA09-sNAp-155, time course study - Day 0

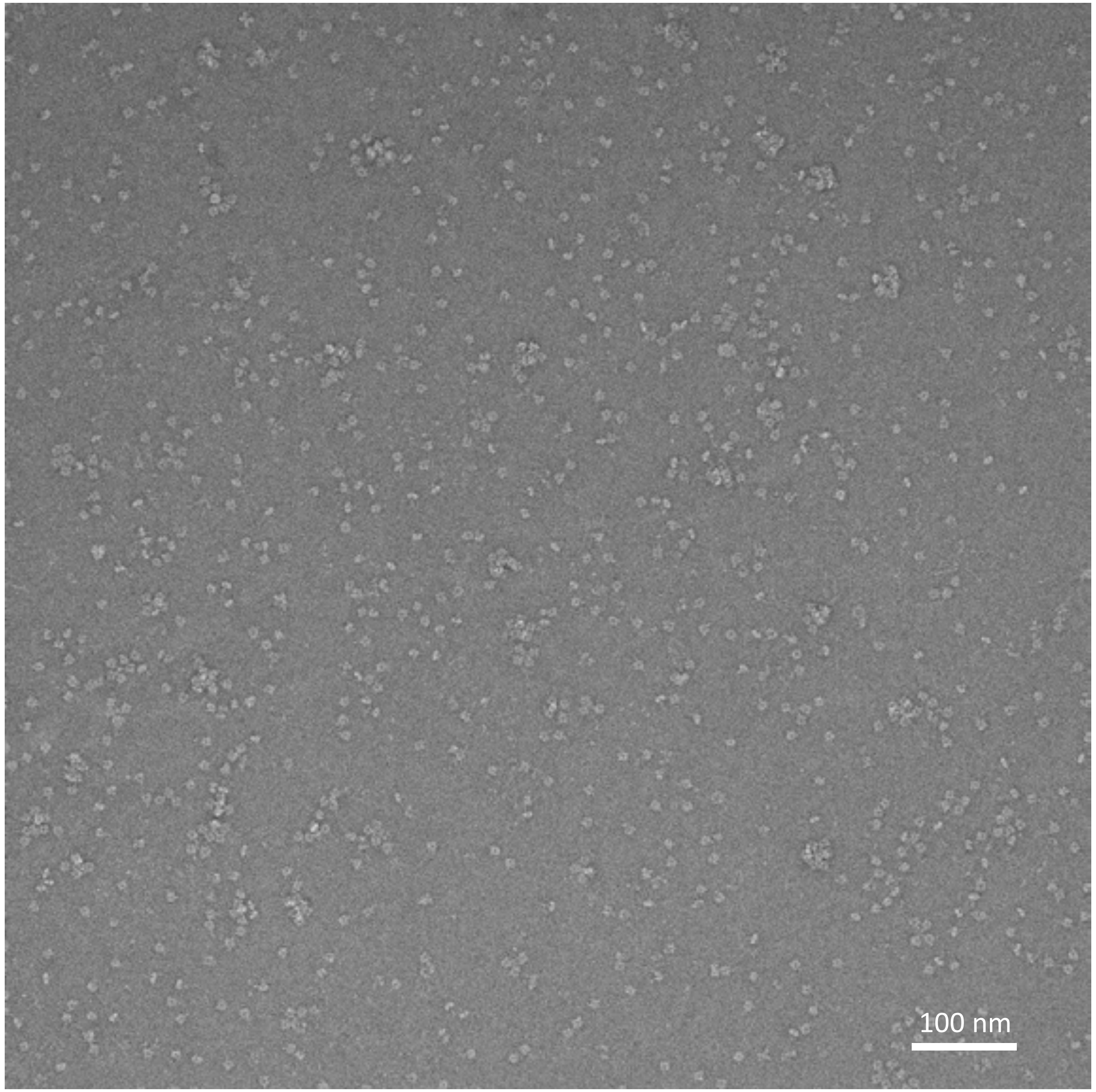


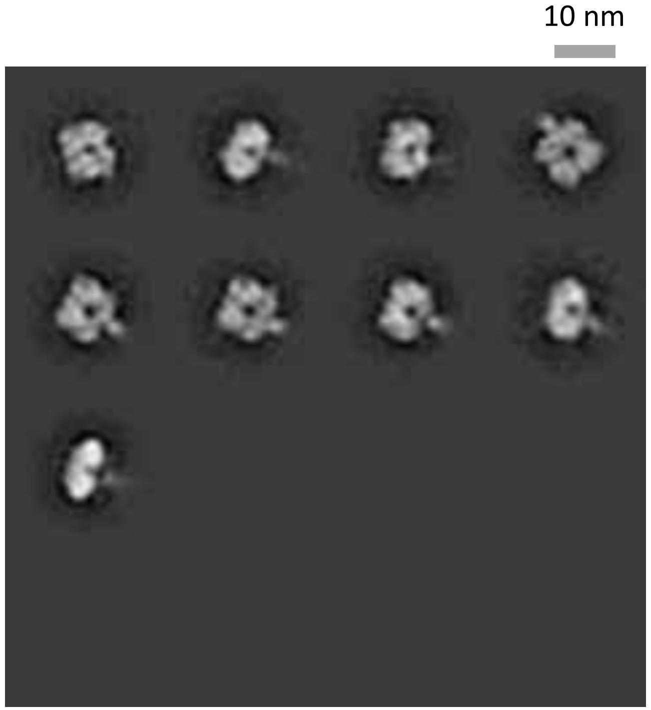

### N1-CA09-sNAp-155, time course study - Day 6

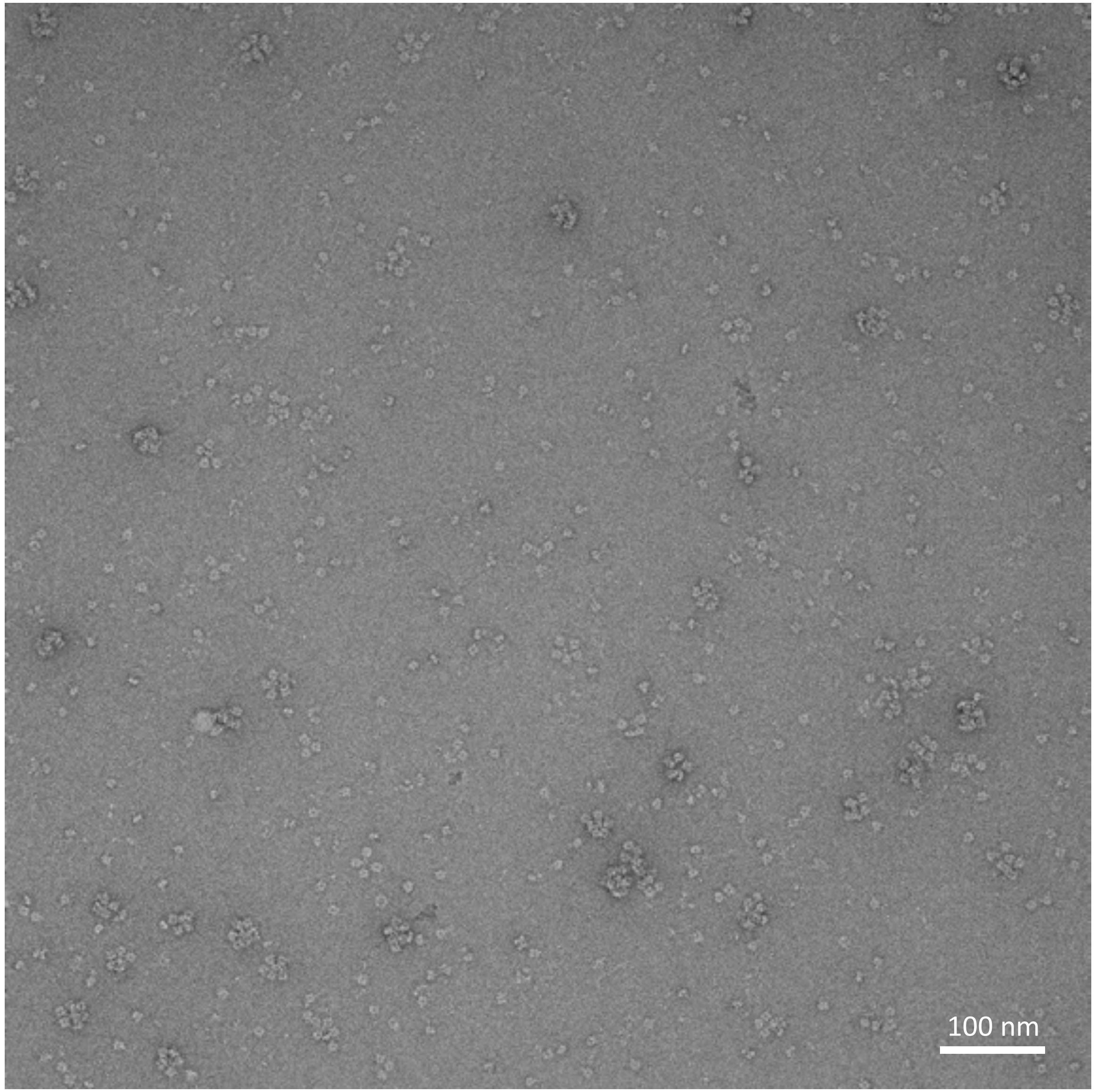


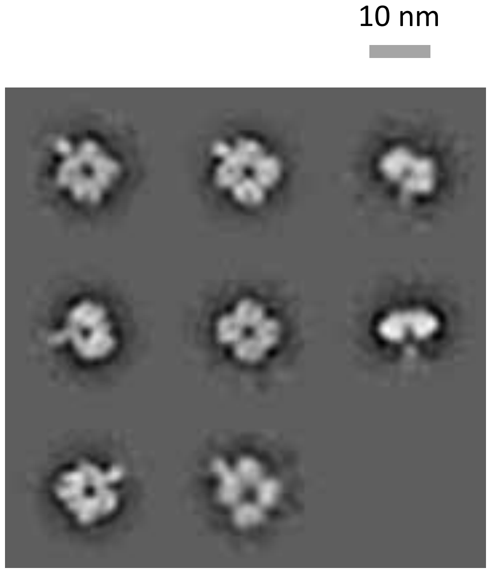

### N1-CA09-sNAp-155, time course study - Day 10

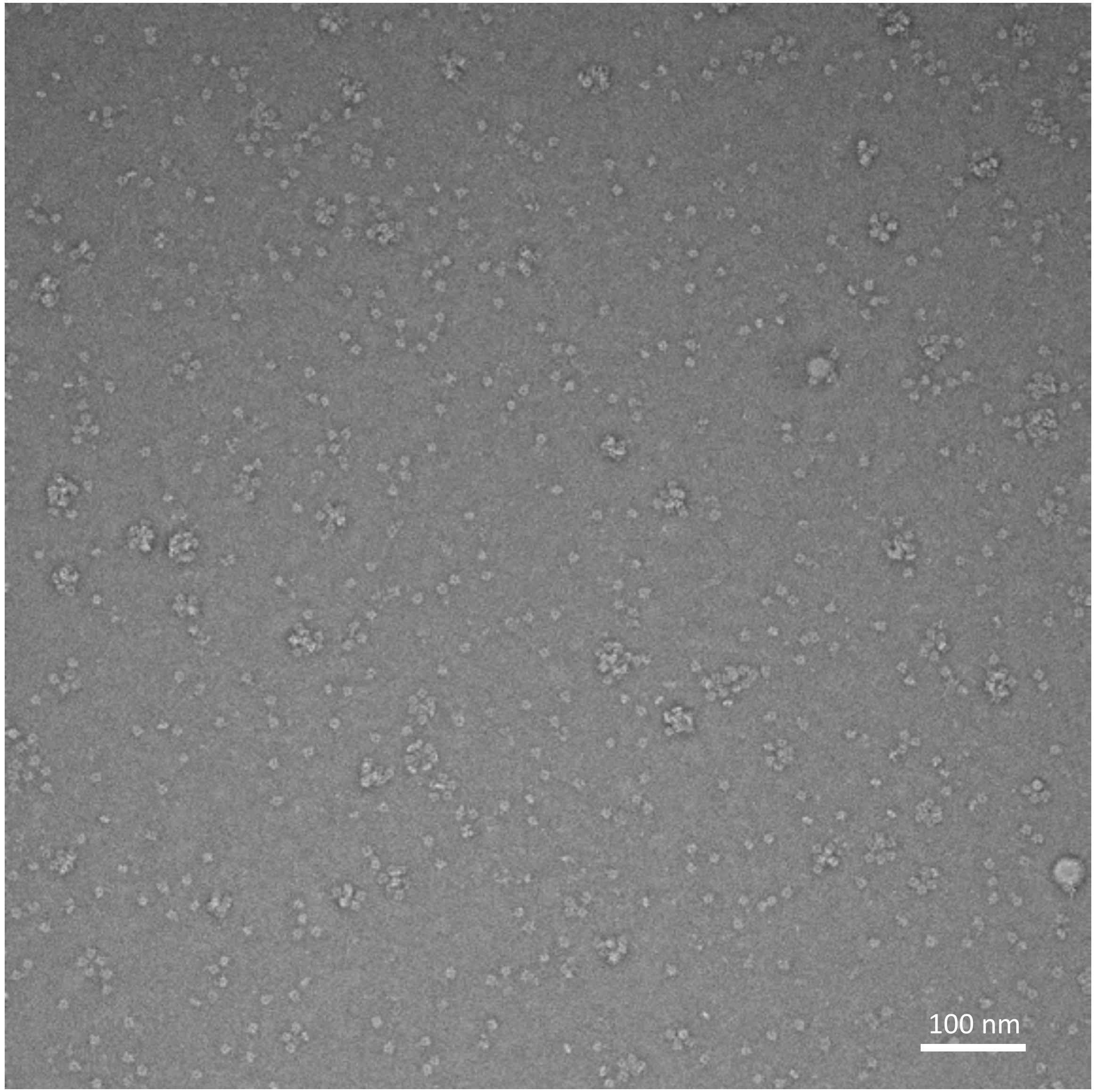


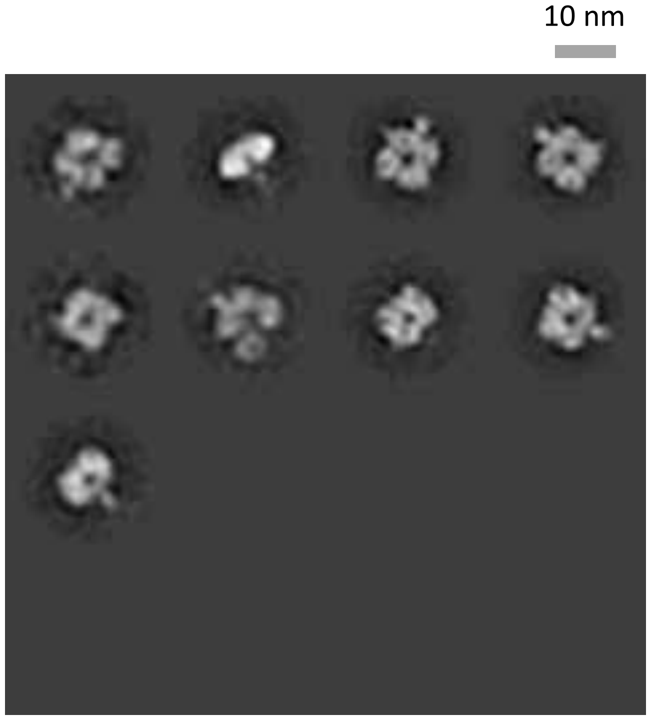

### N1-CA09-sNAp-155, time course study - Day 15

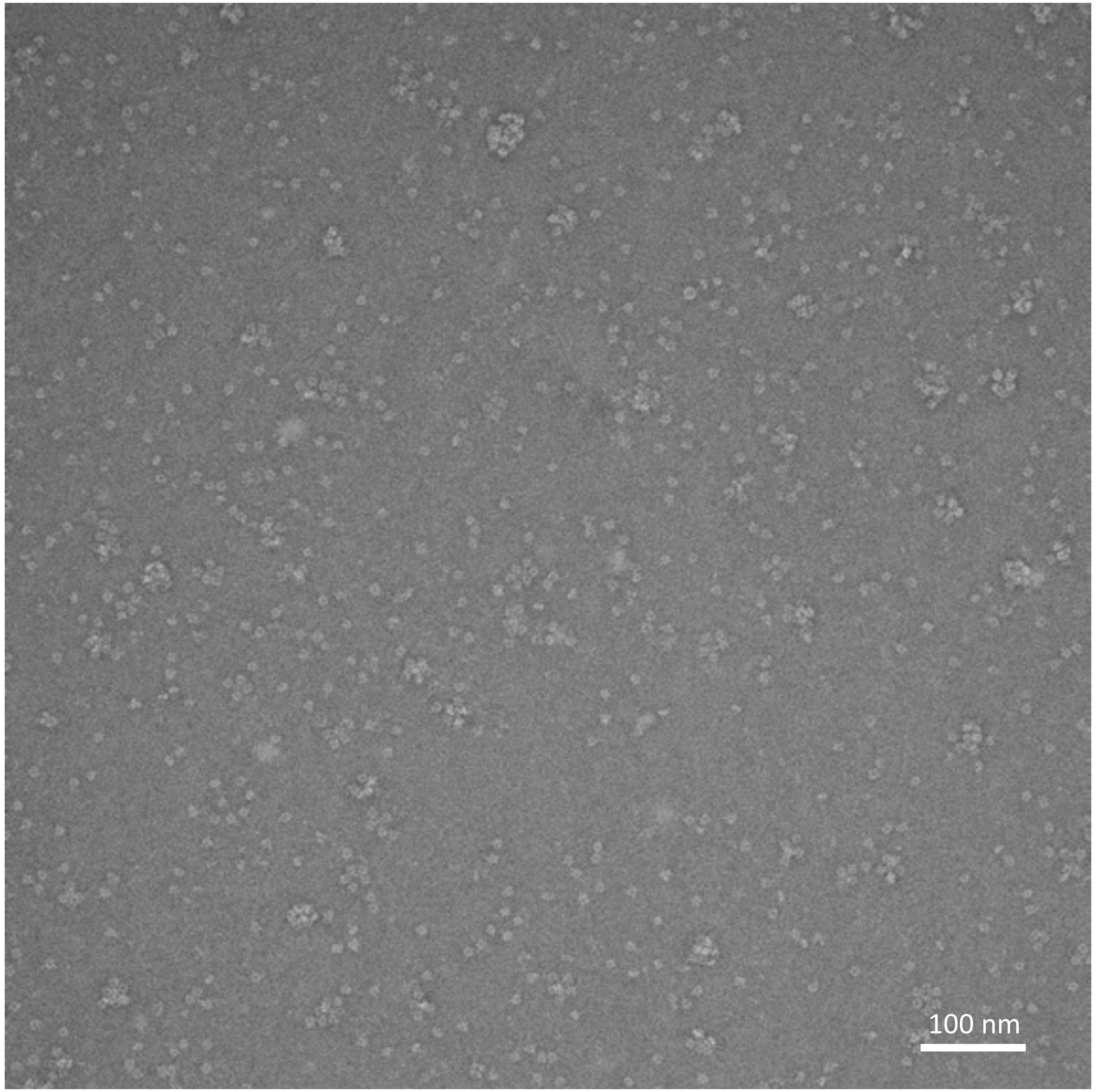


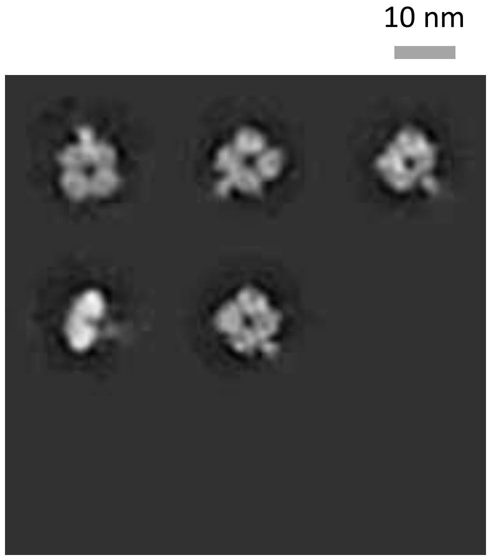

### N1-CA09-sNAp-131

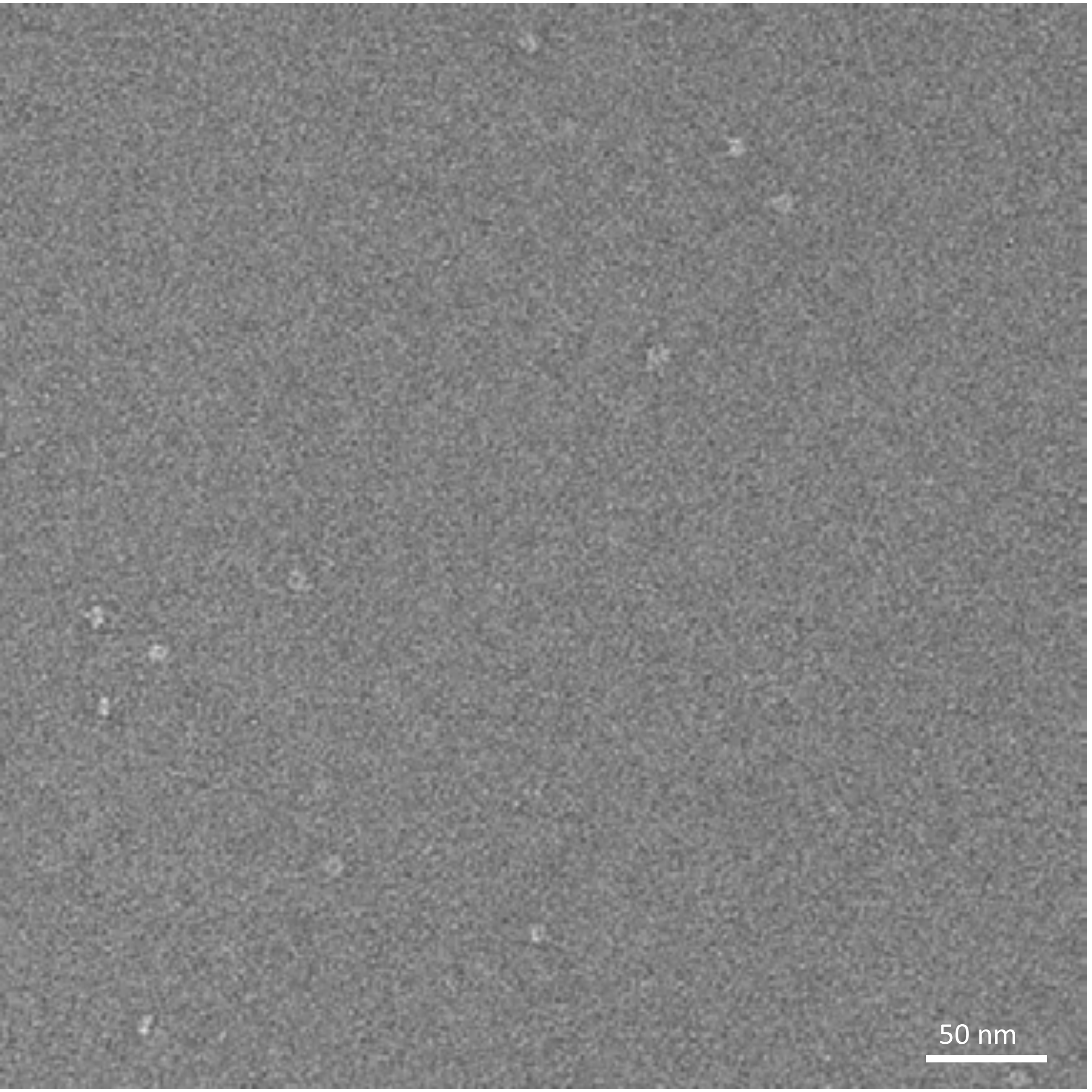


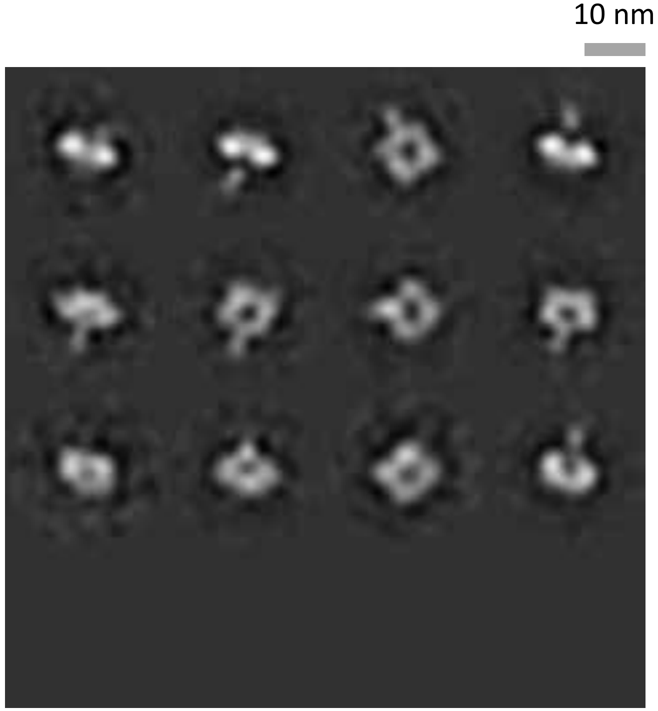

### N1-CA09-sNAp-134

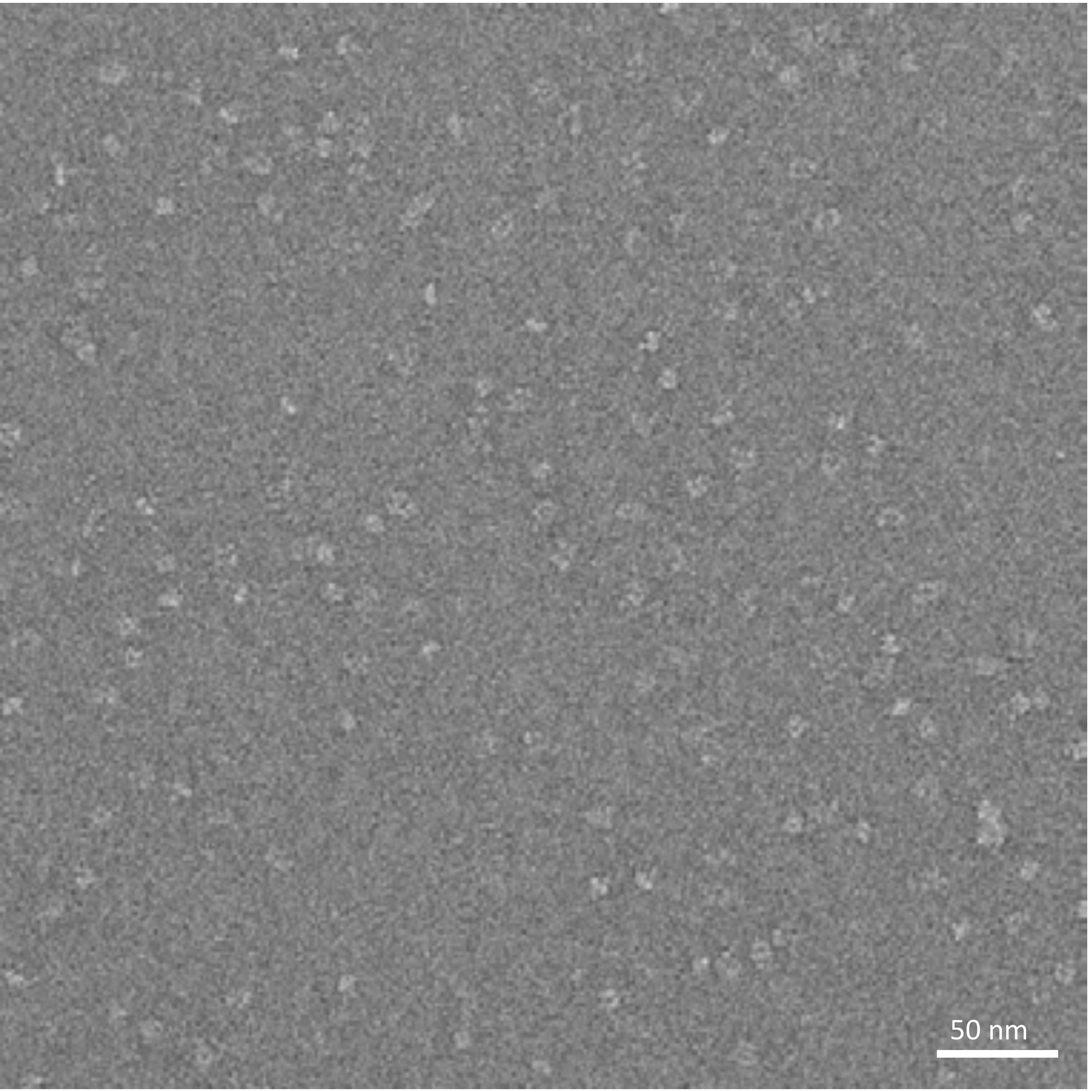


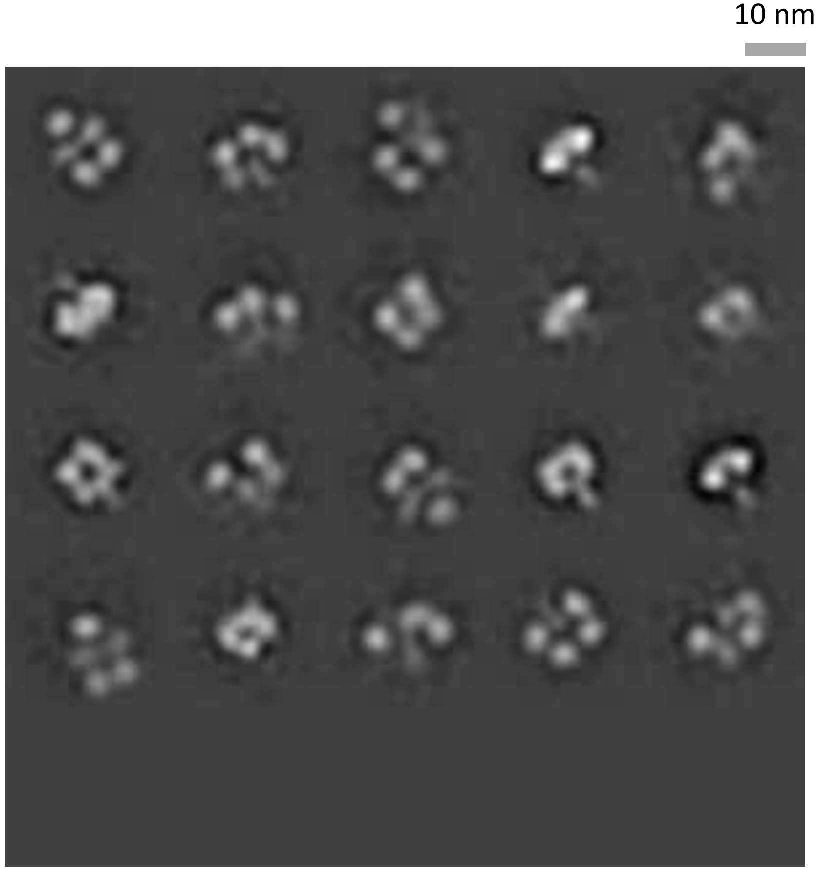

### N8-JD13-sNAp-282

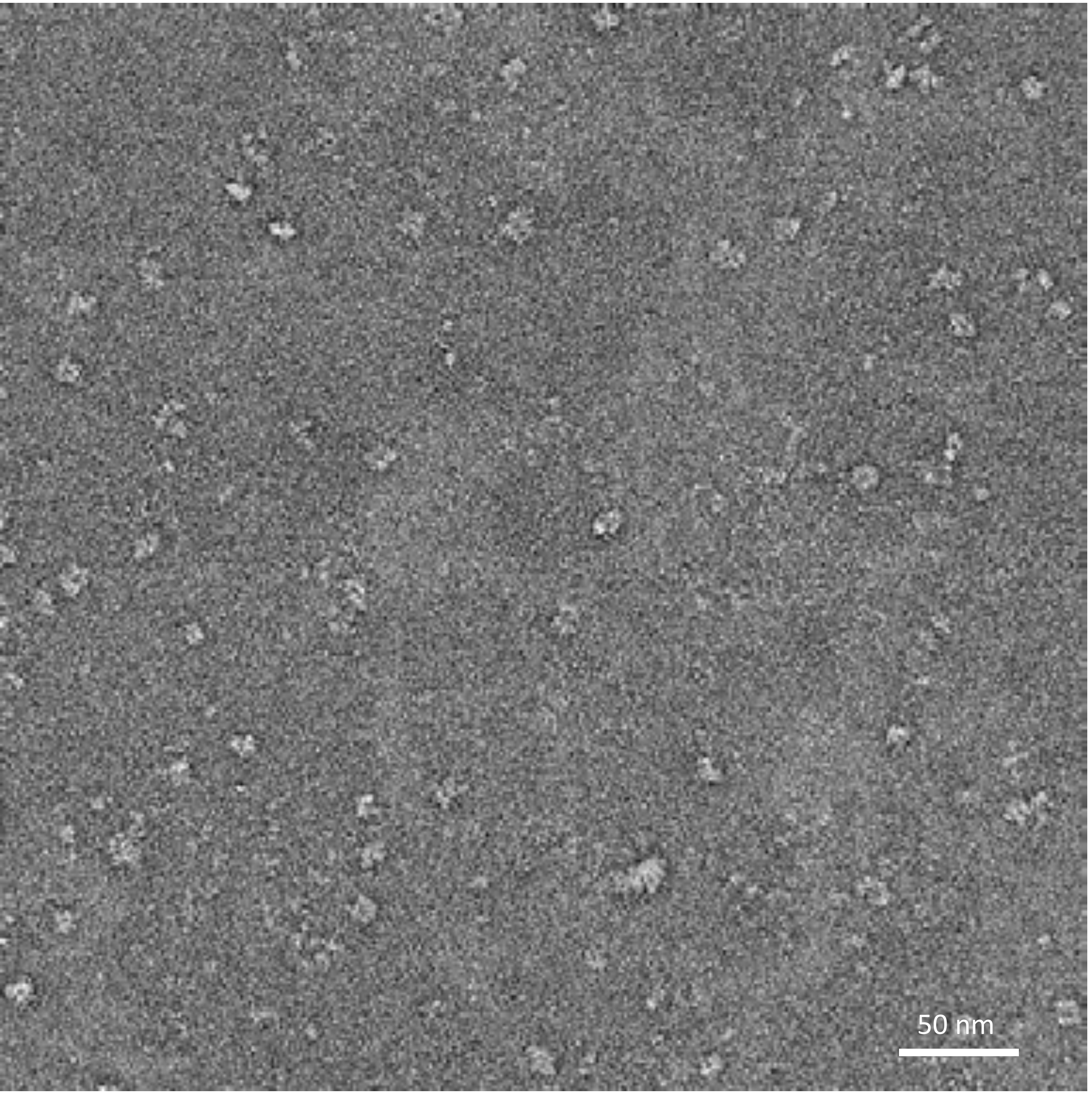


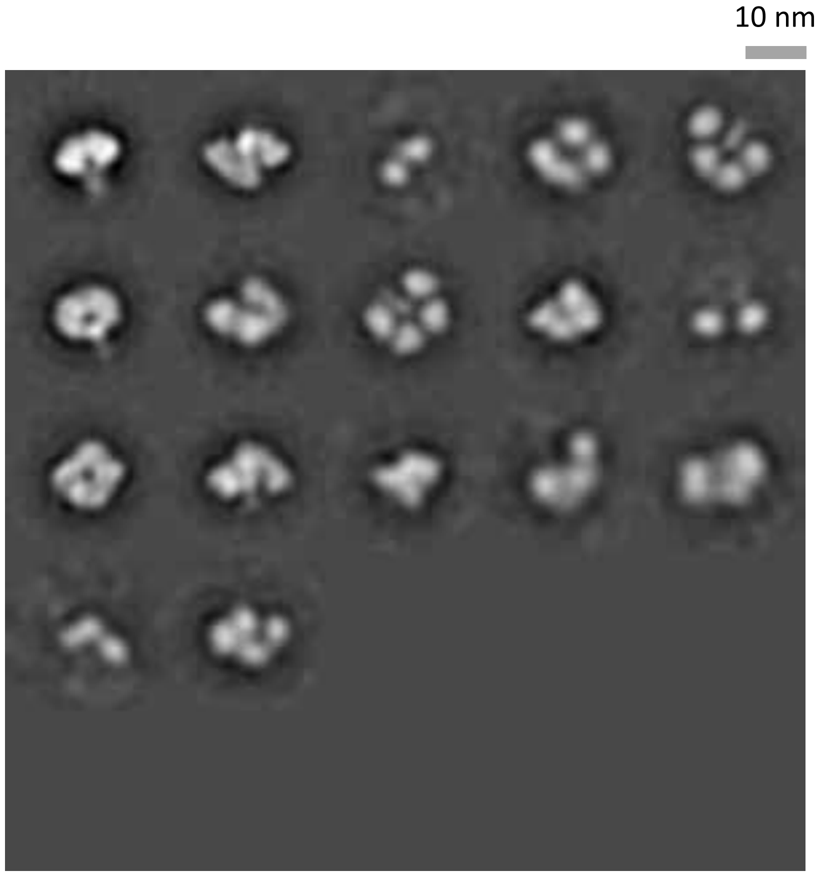

### N8-JD13-sNAp-285

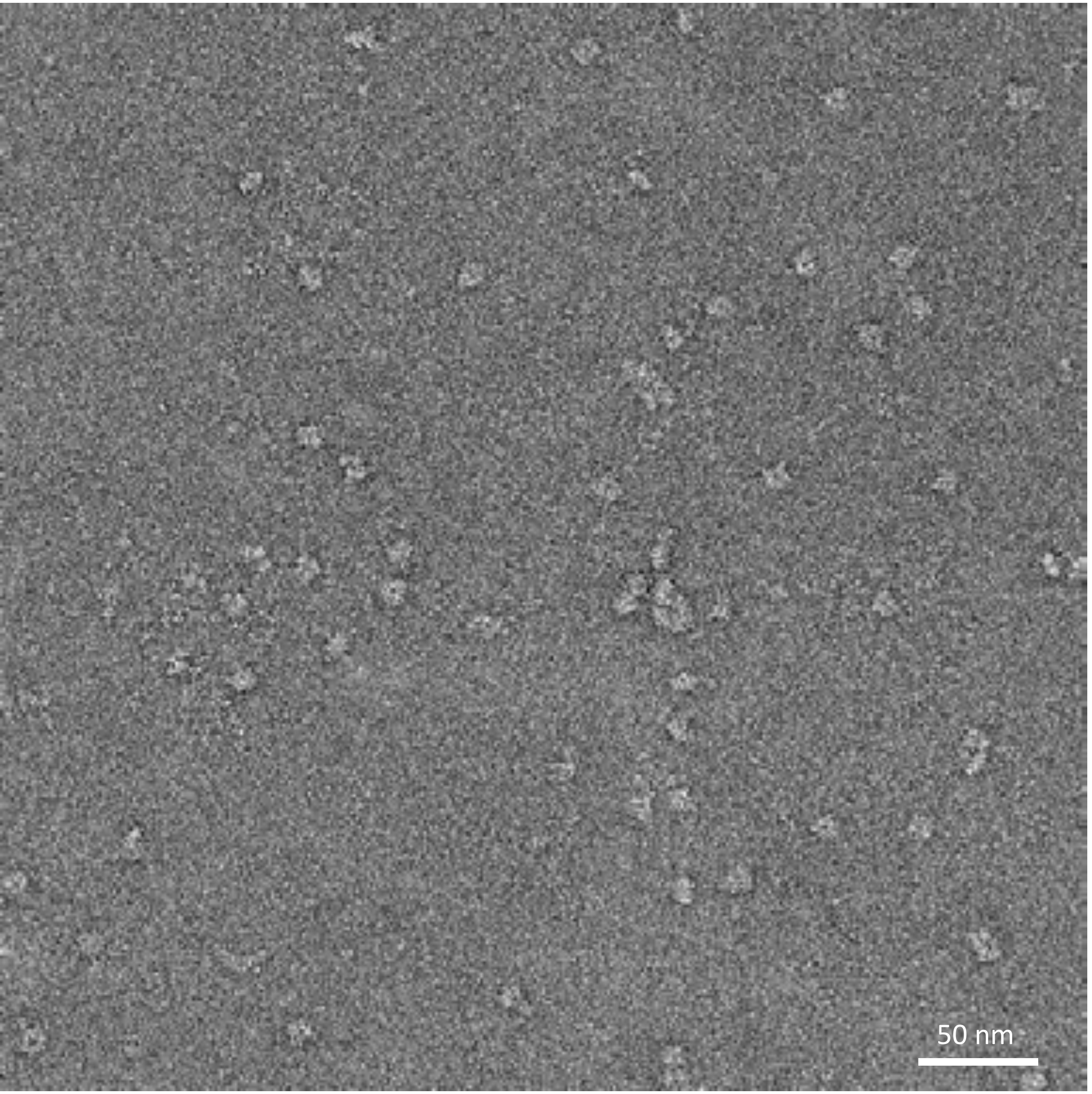


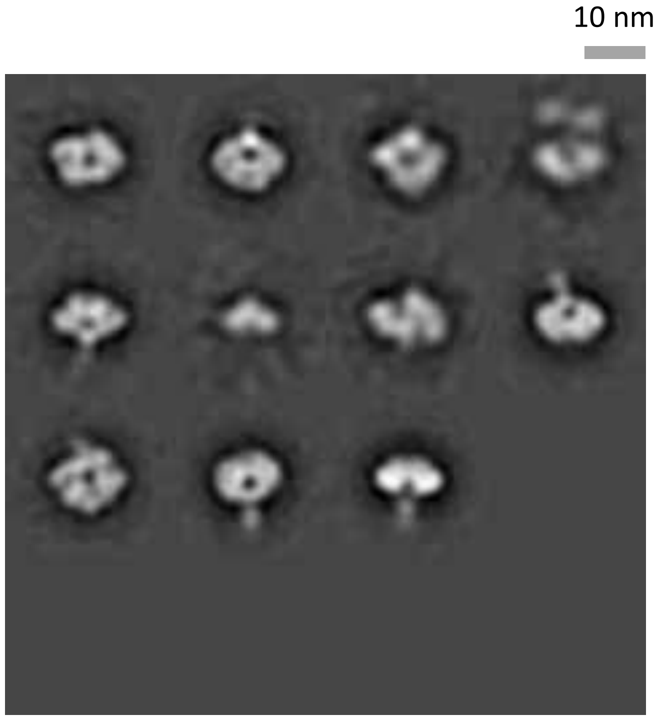

### N8-JD13-sNAp-285, time course study - Day 0

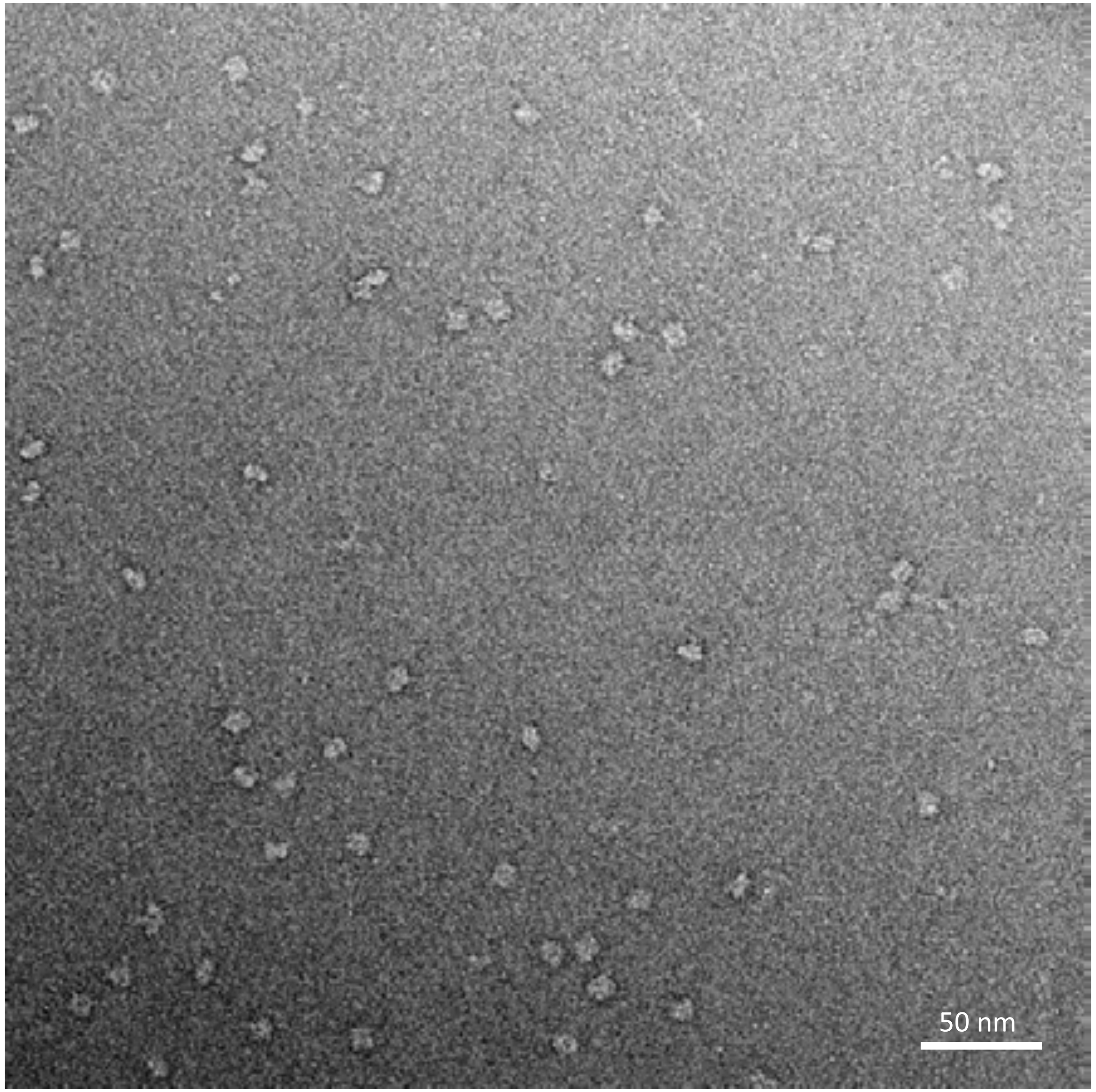


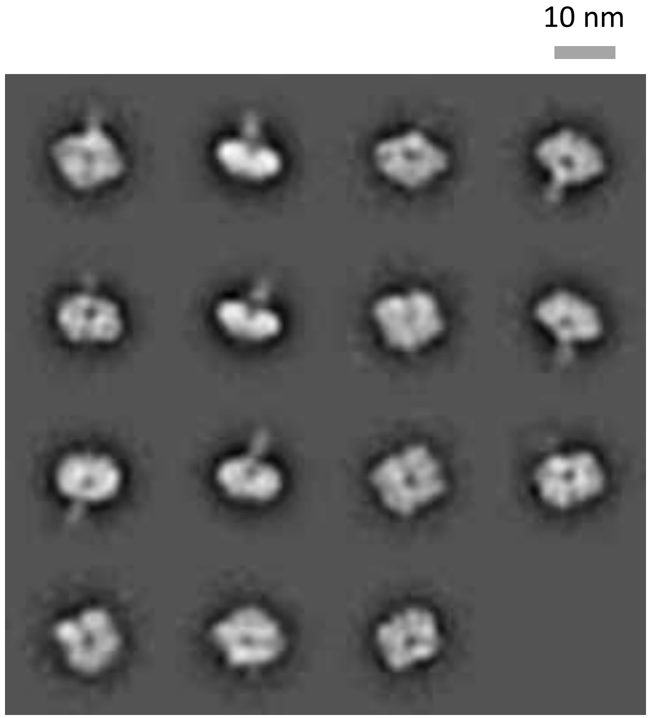

### N8-JD13-sNAp-285, time course study - Day 6

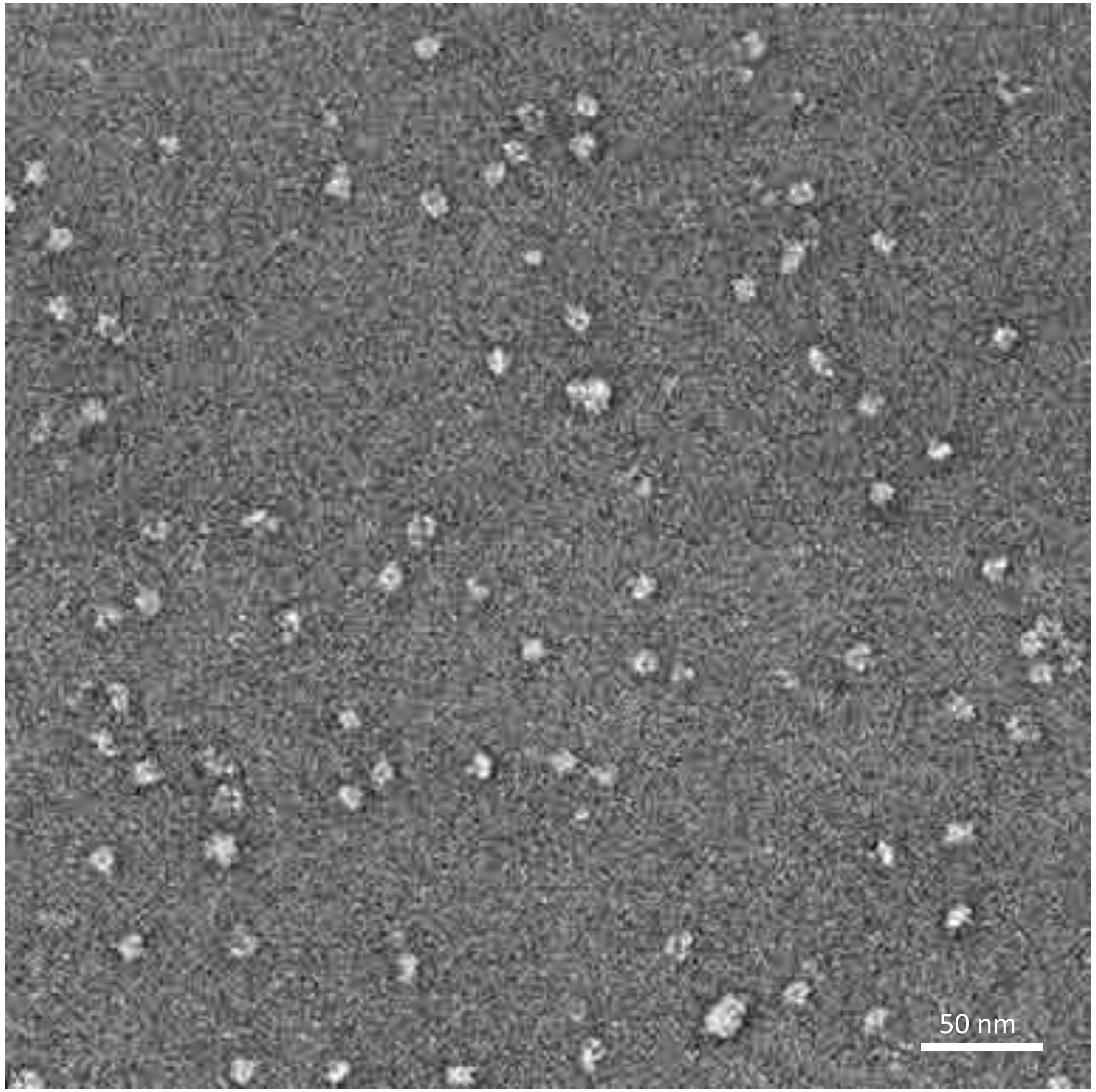


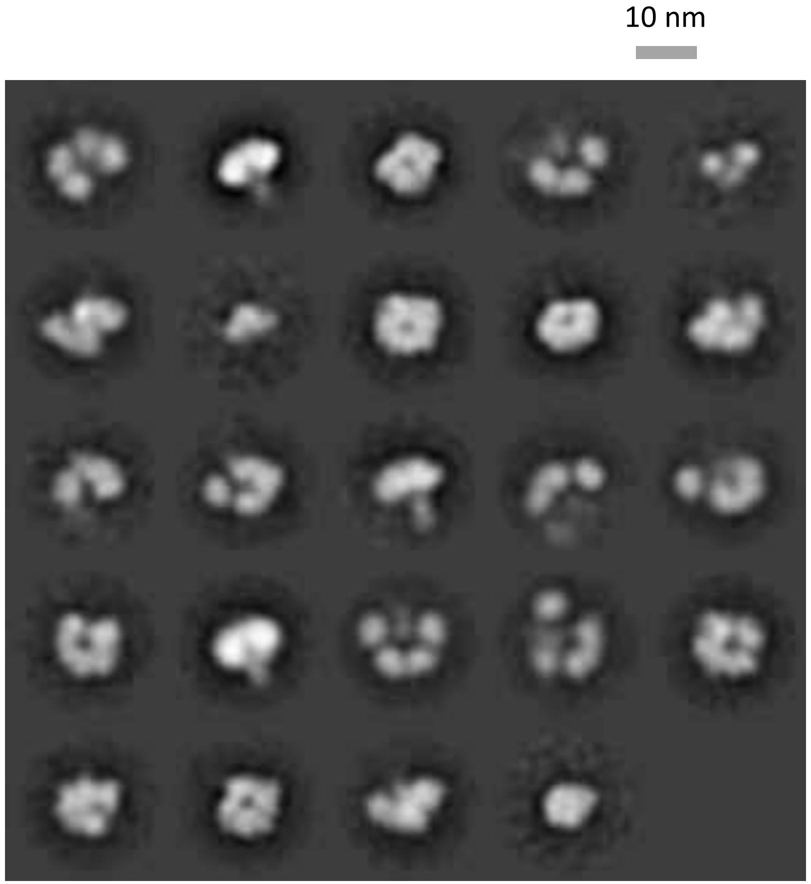

### N2-WI05-desNAp-156

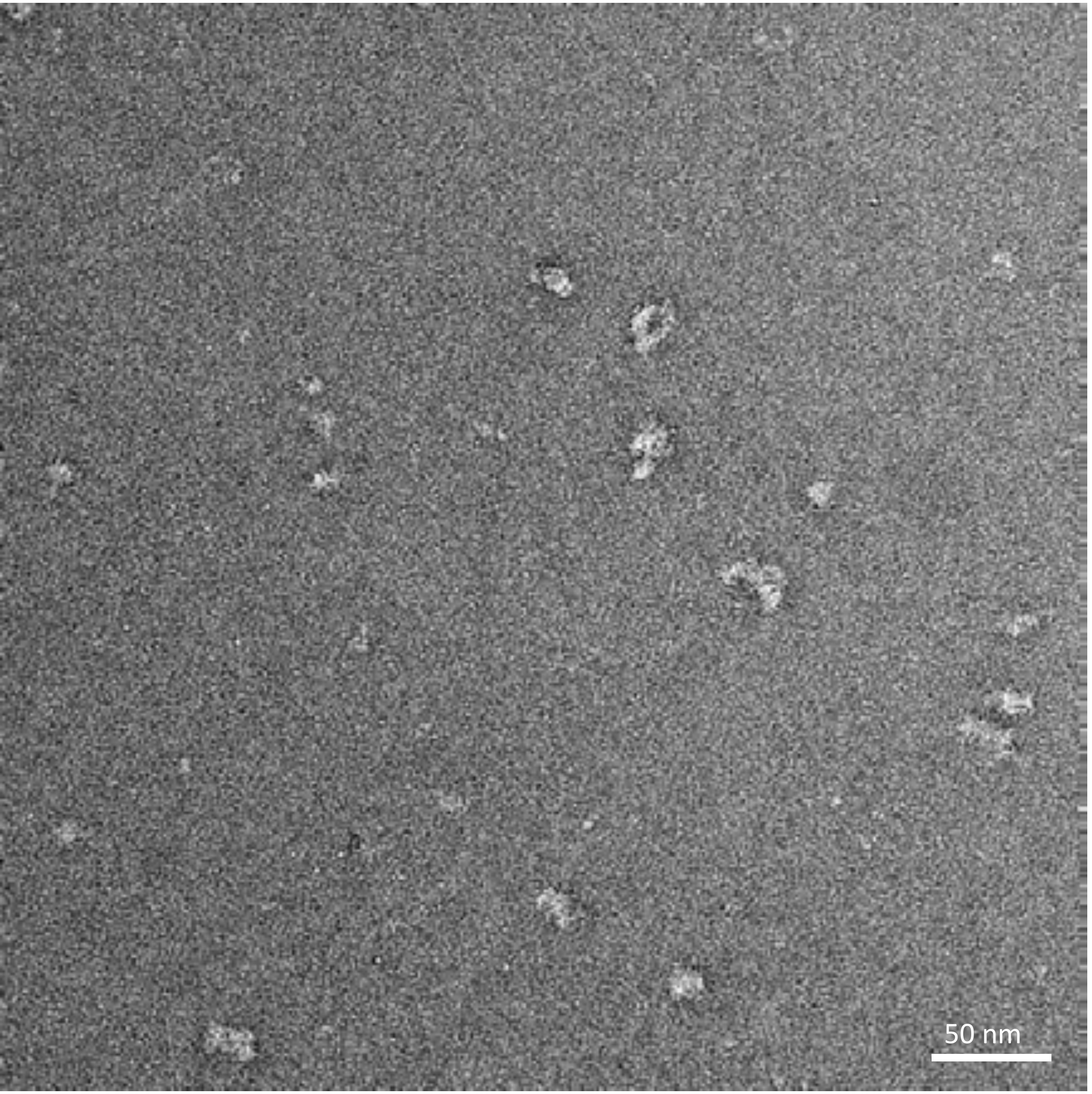


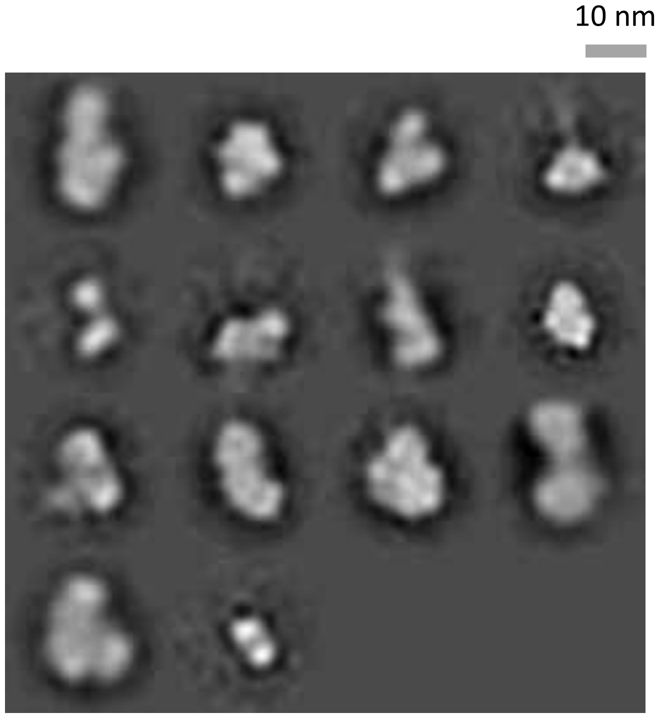

### N2-WI05-desNAp-157

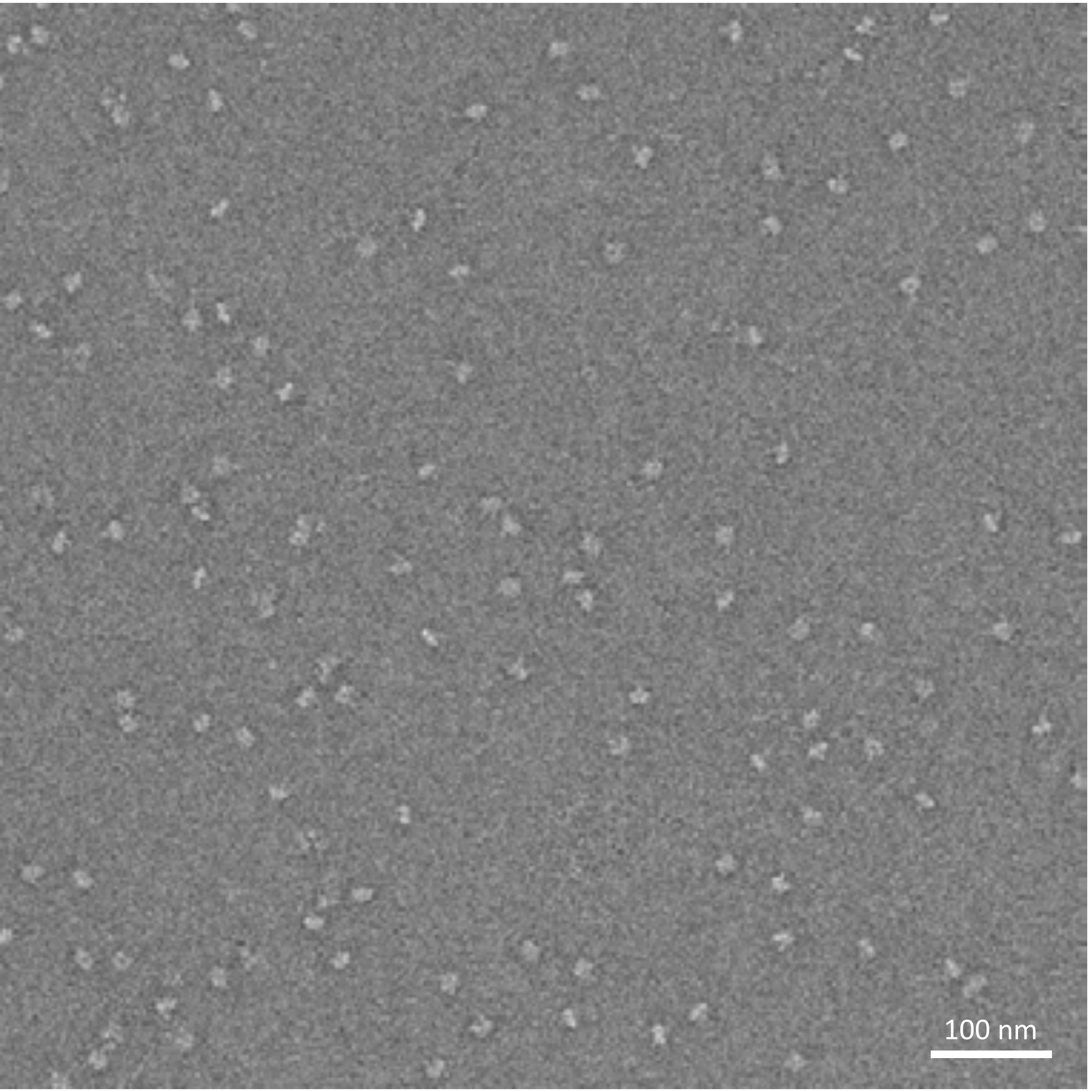


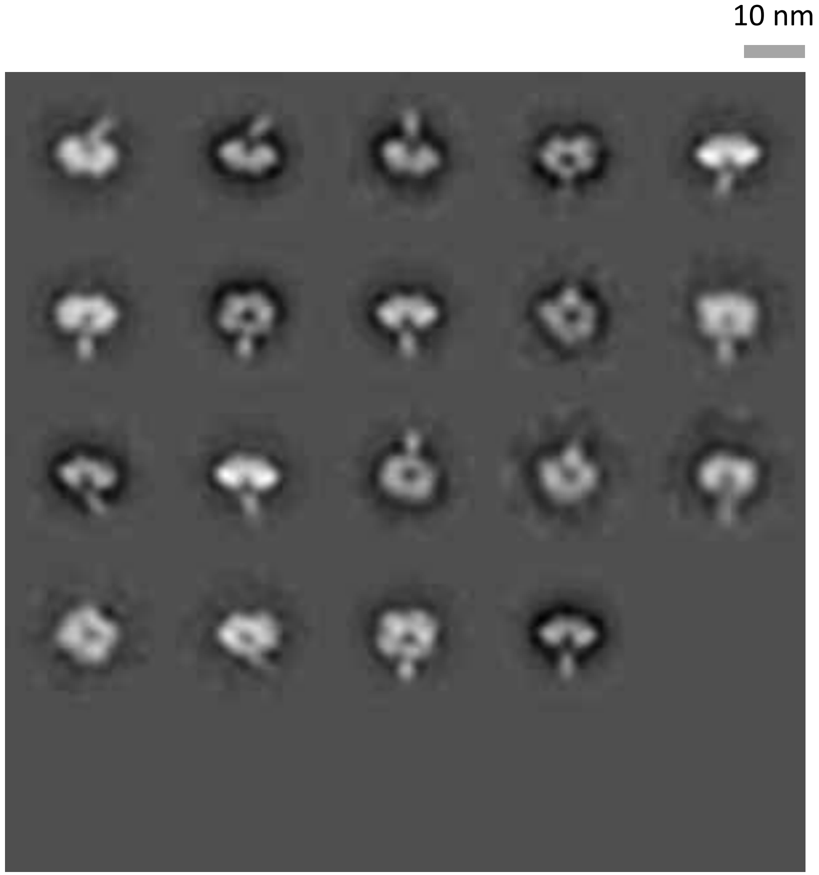

### N2-WI05-desNAp-158

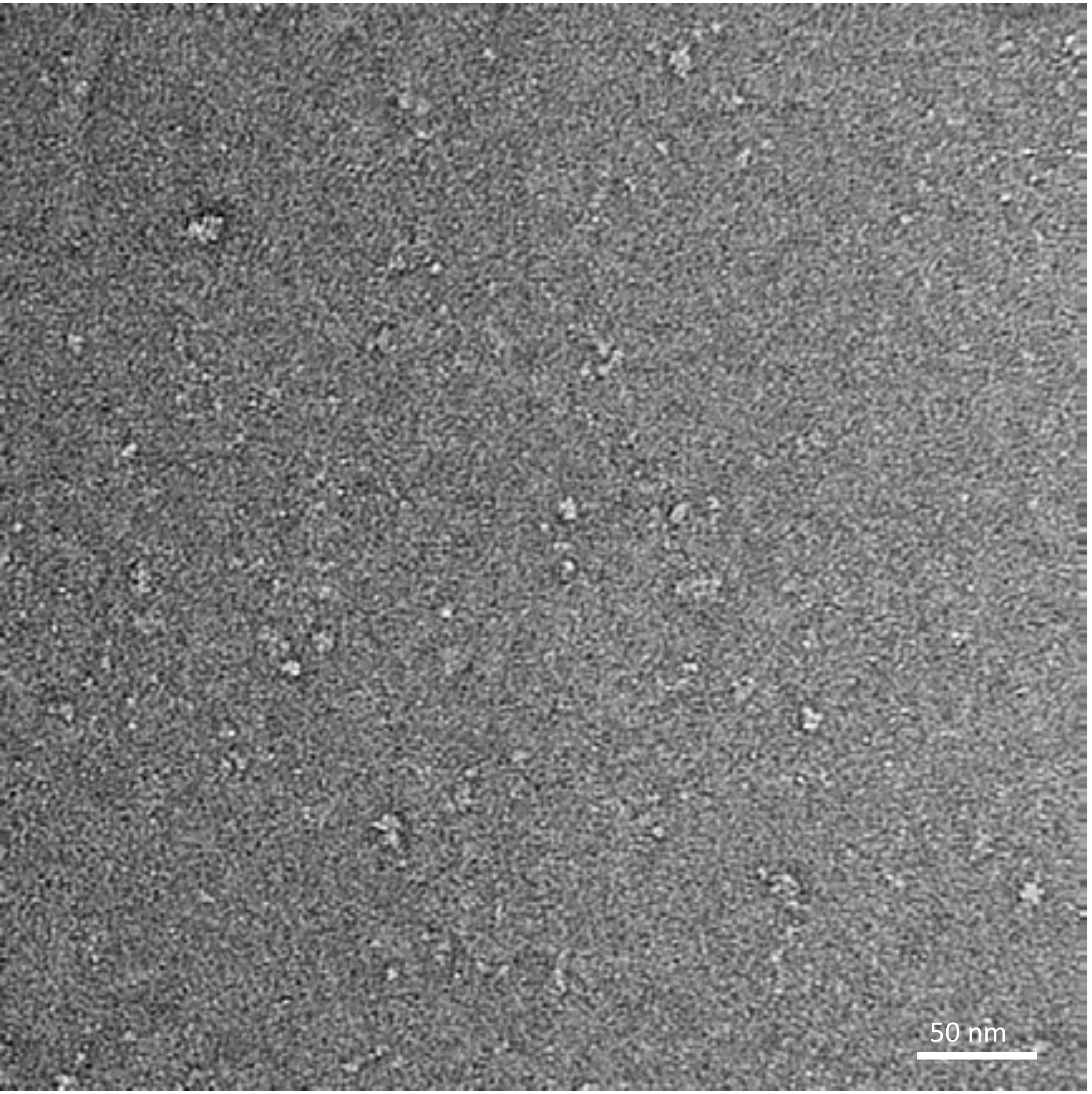


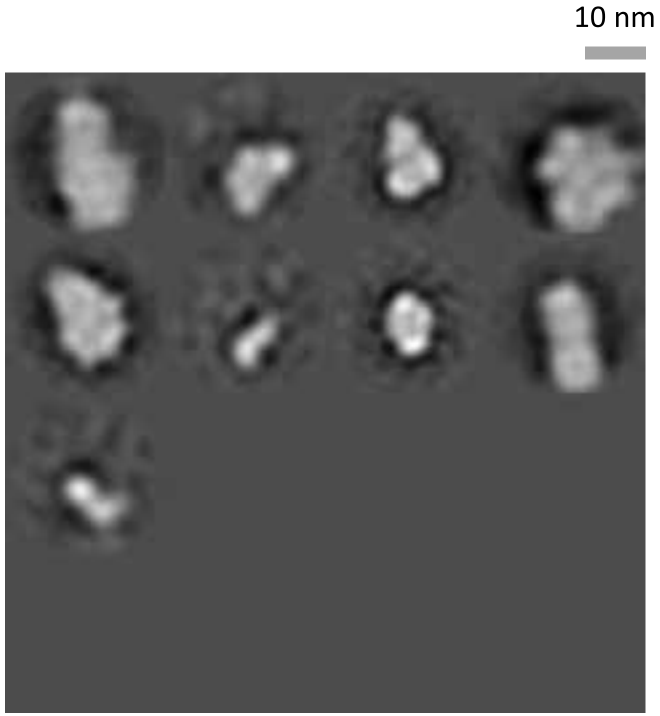

### N2-WI05-desNAp-249

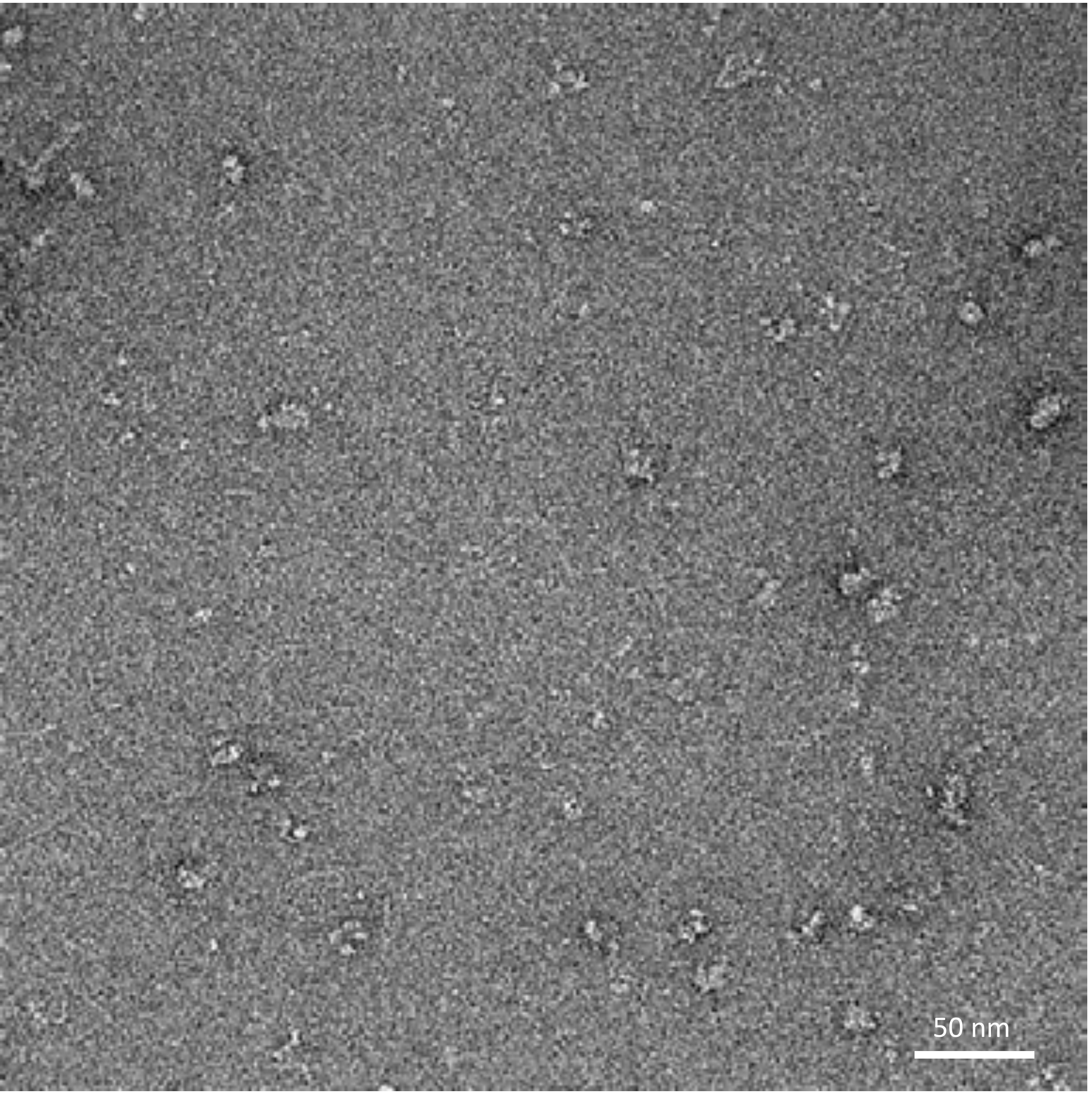


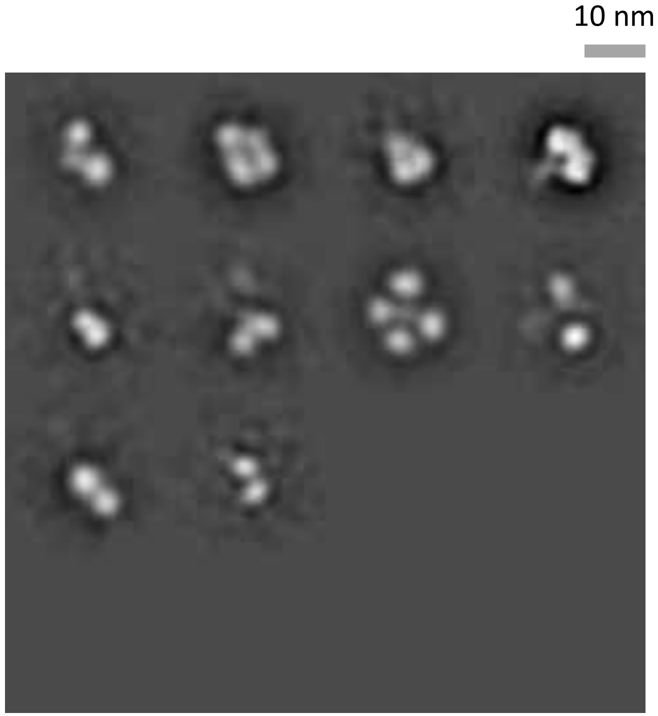

### N2-WI05-desNAp-255

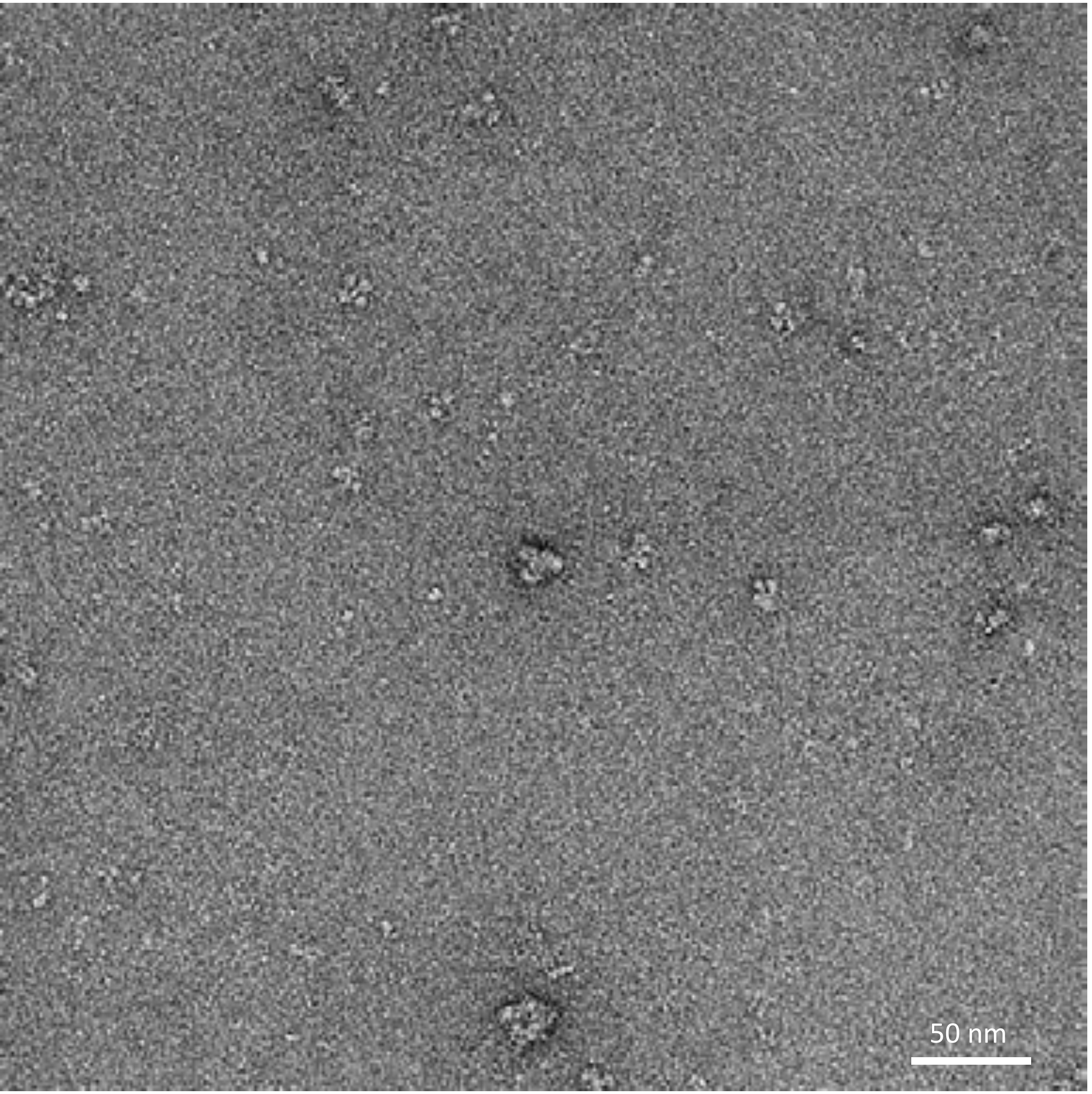


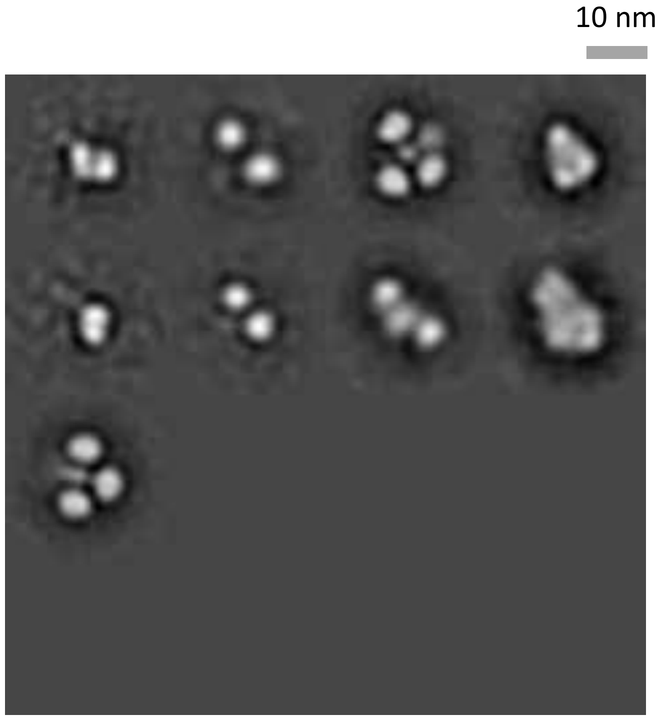

### N1-MI15-sNAp-155

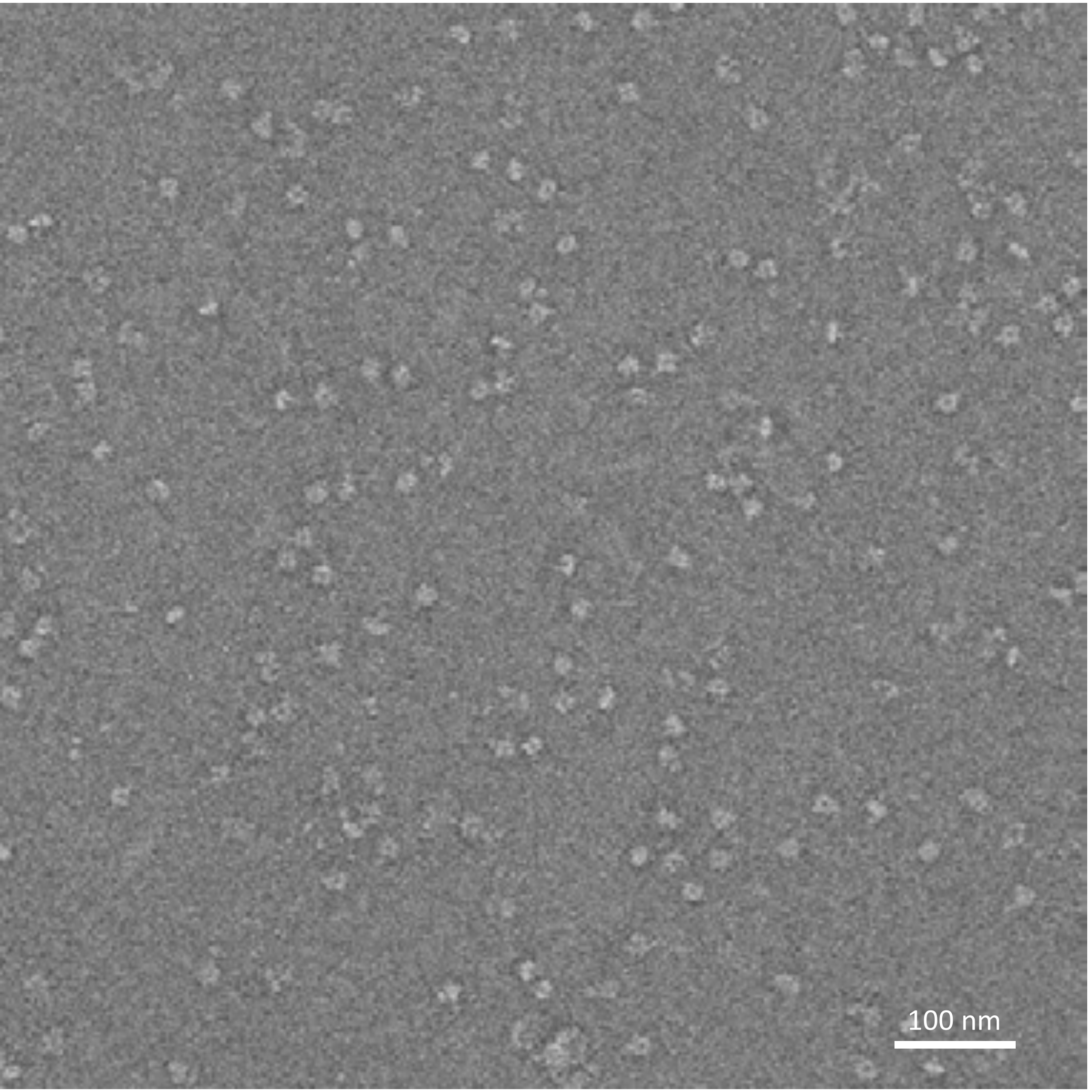


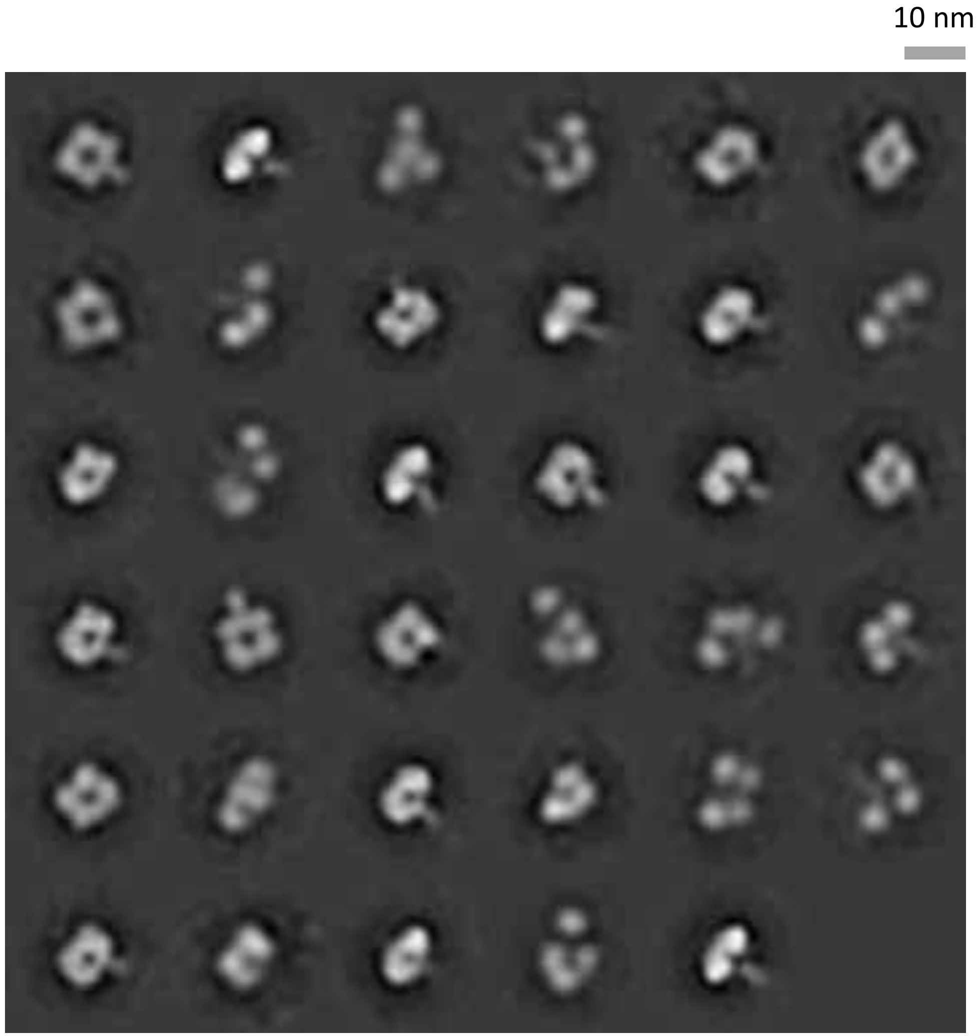

### N1-MI15-sNAp-174

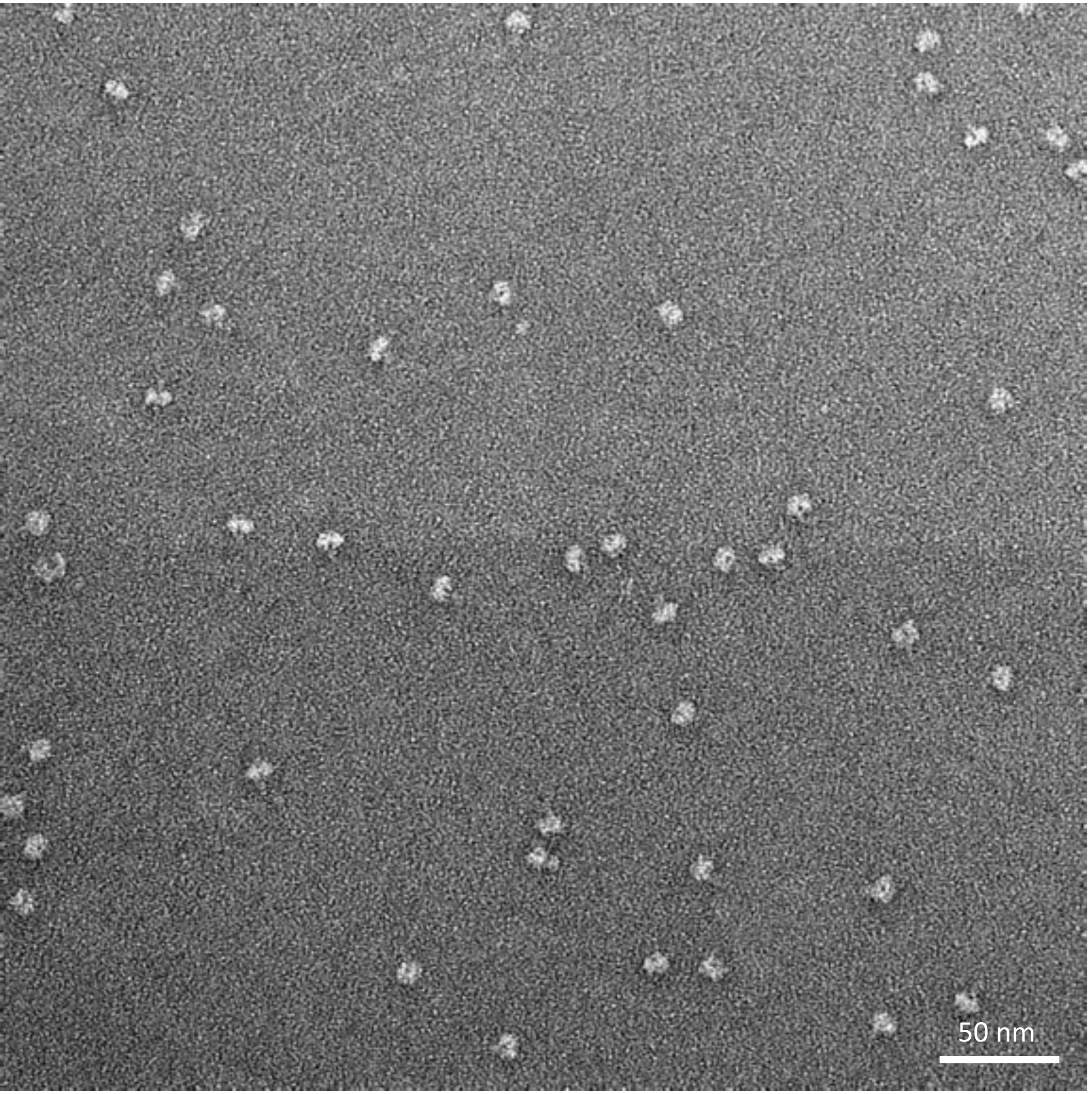


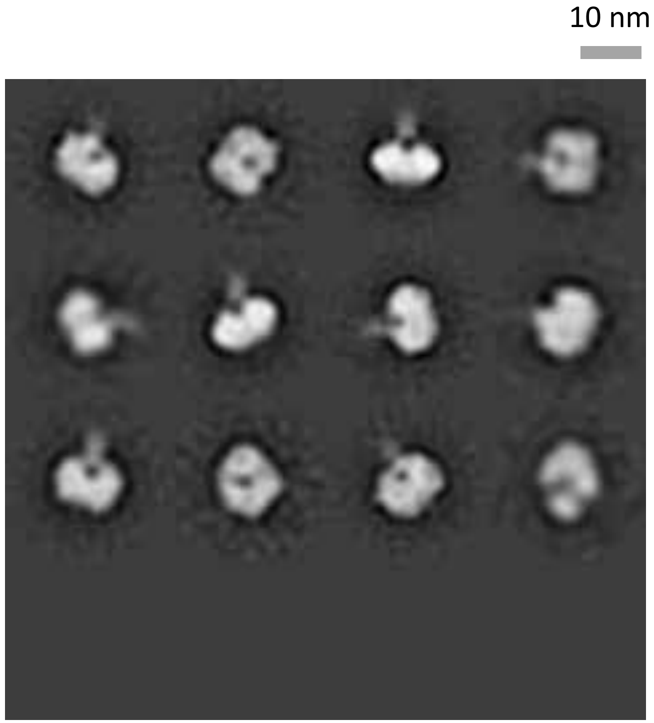

### N1-MI15-sNAp-174, time course study - Day 0

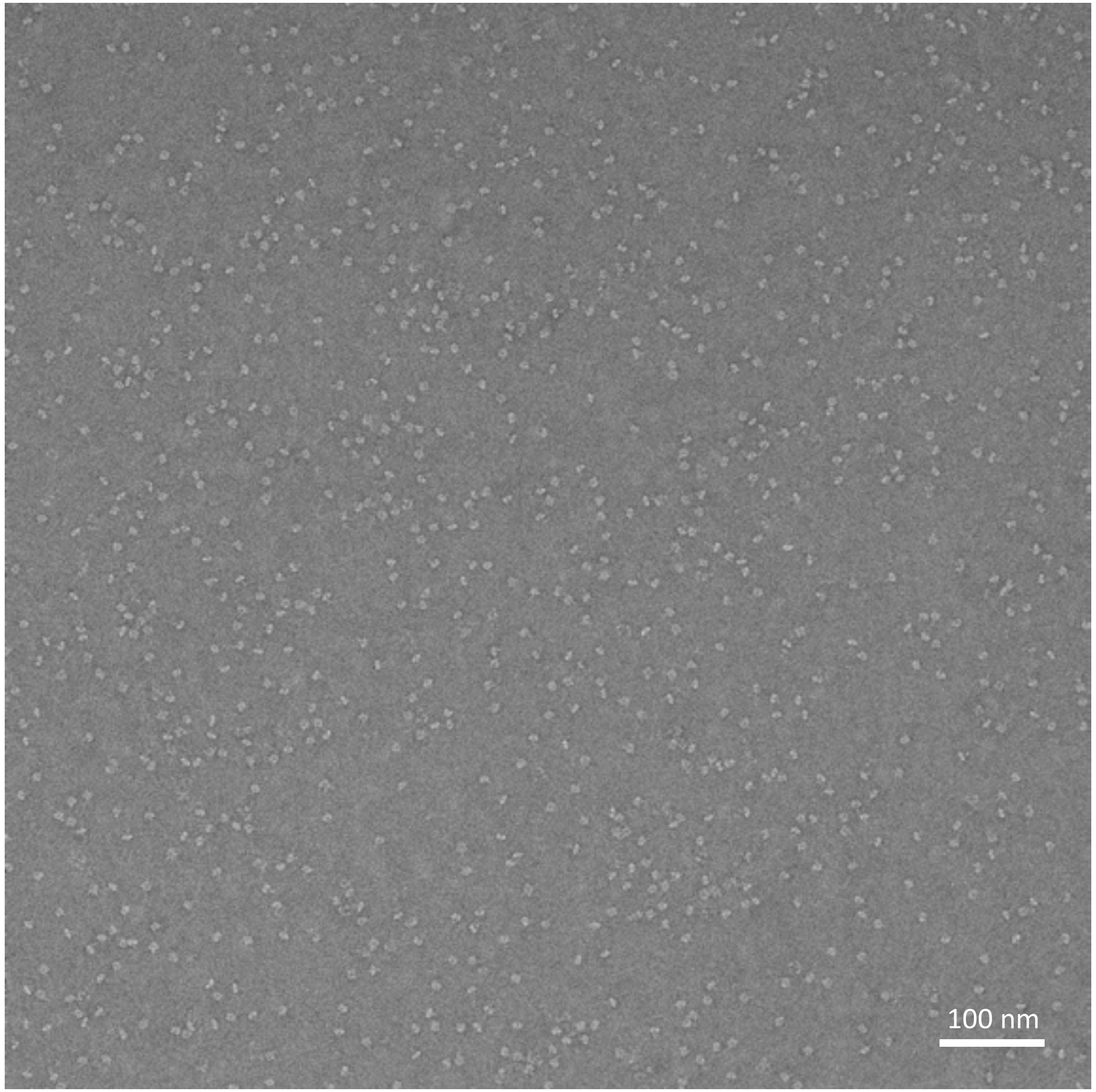


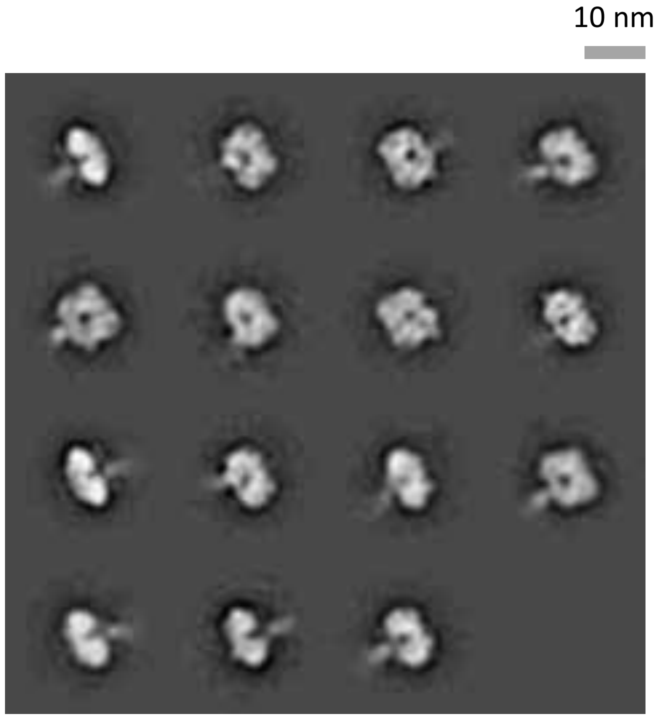

### N1-MI15-sNAp-174, time course study - Day 6

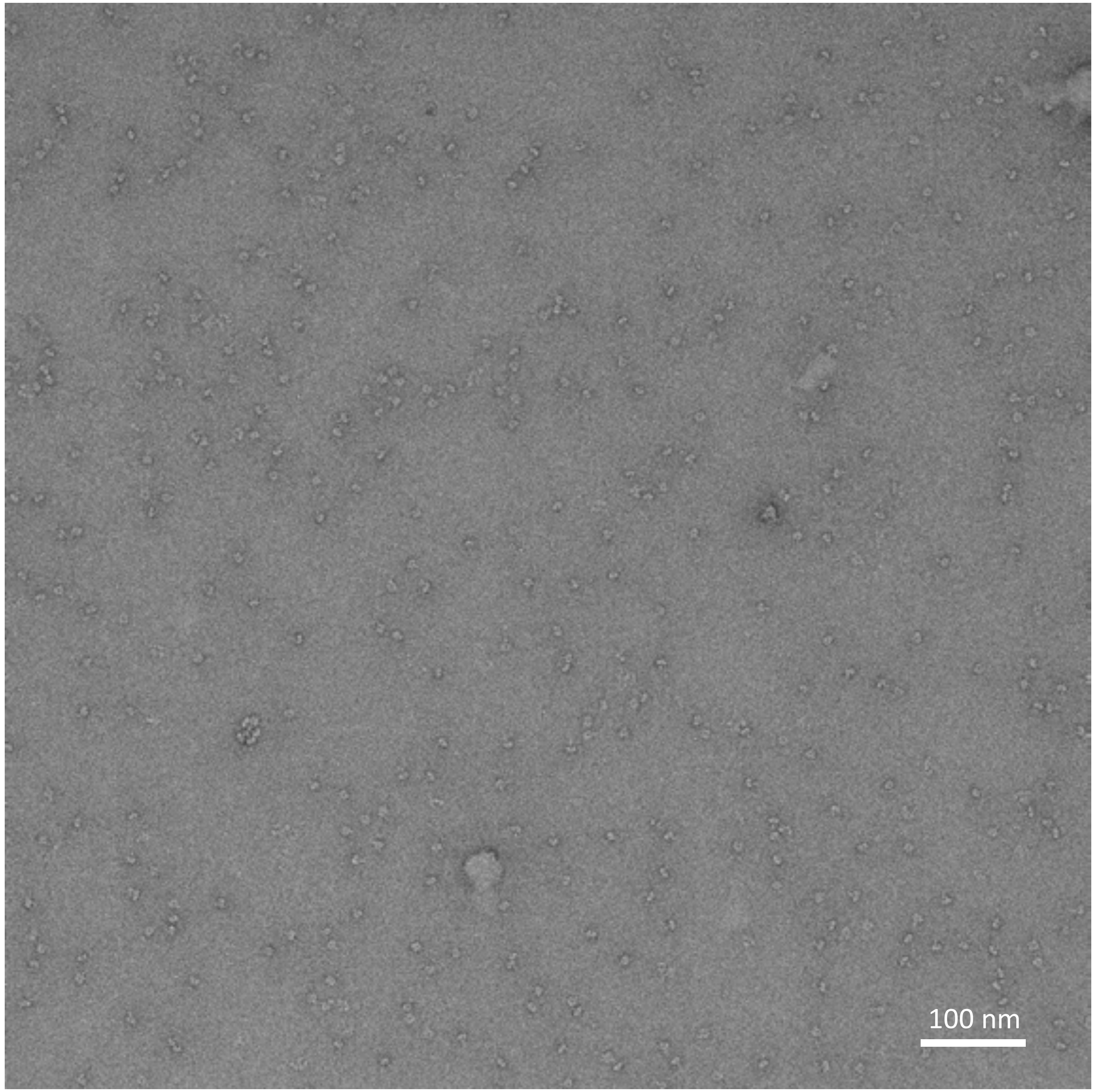


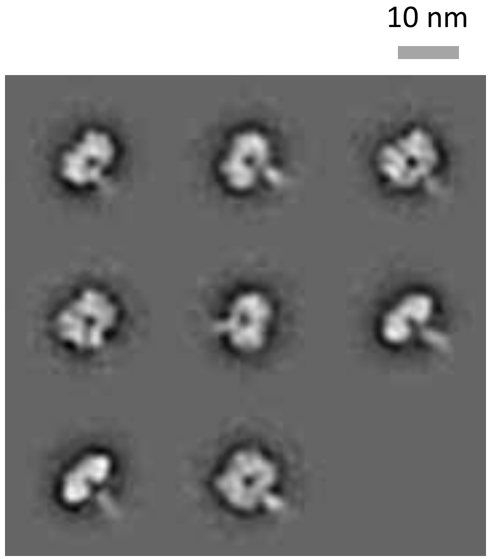

### N1-MI15-sNAp-174, time course study - Day 10

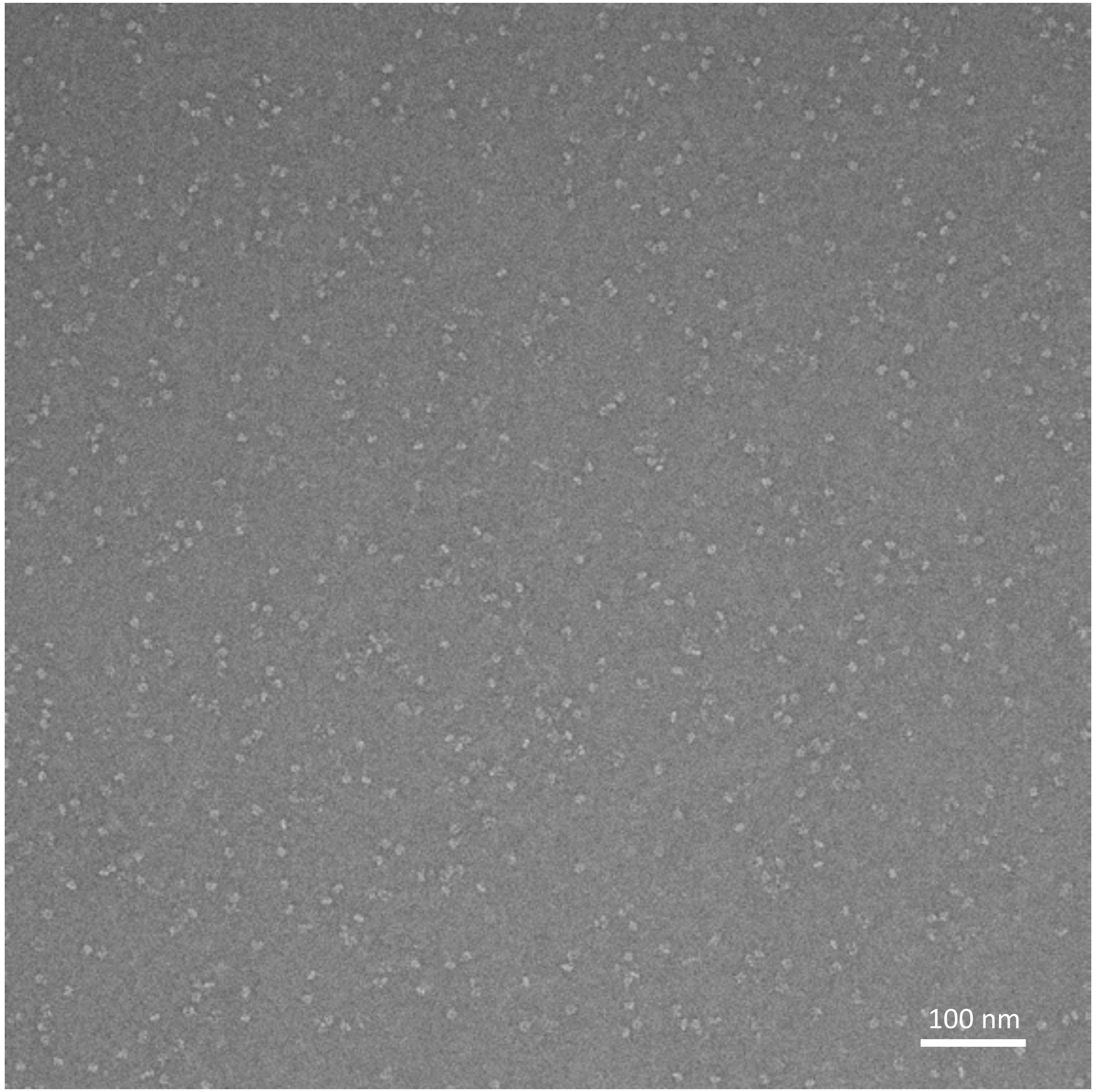


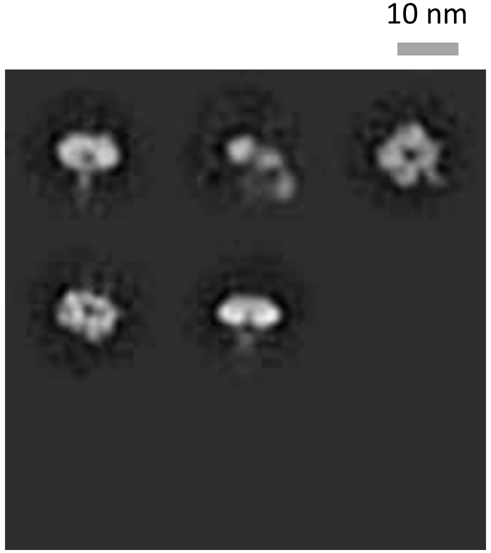

### N1-MI15-sNAp-174, time course study - Day 15

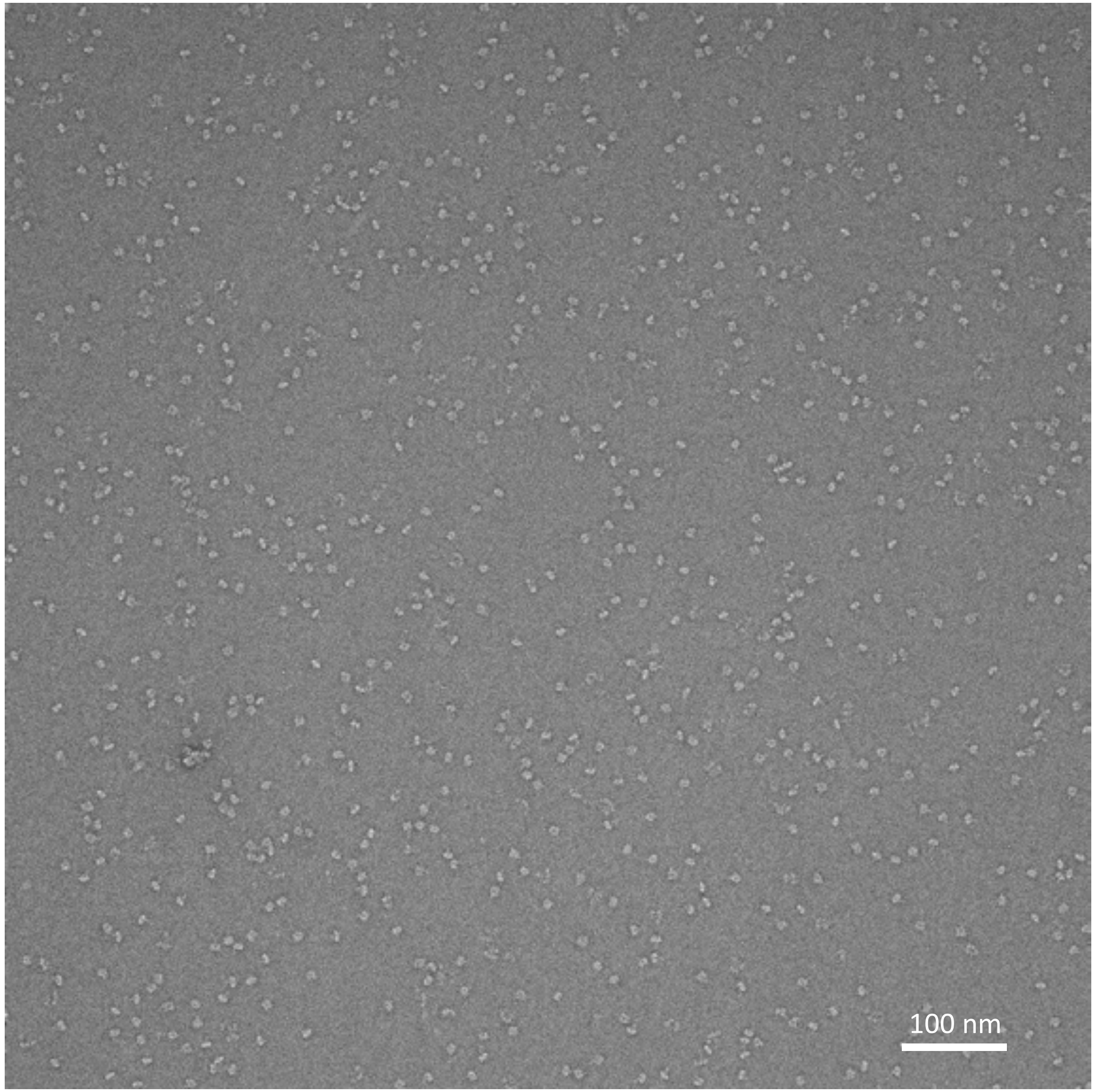


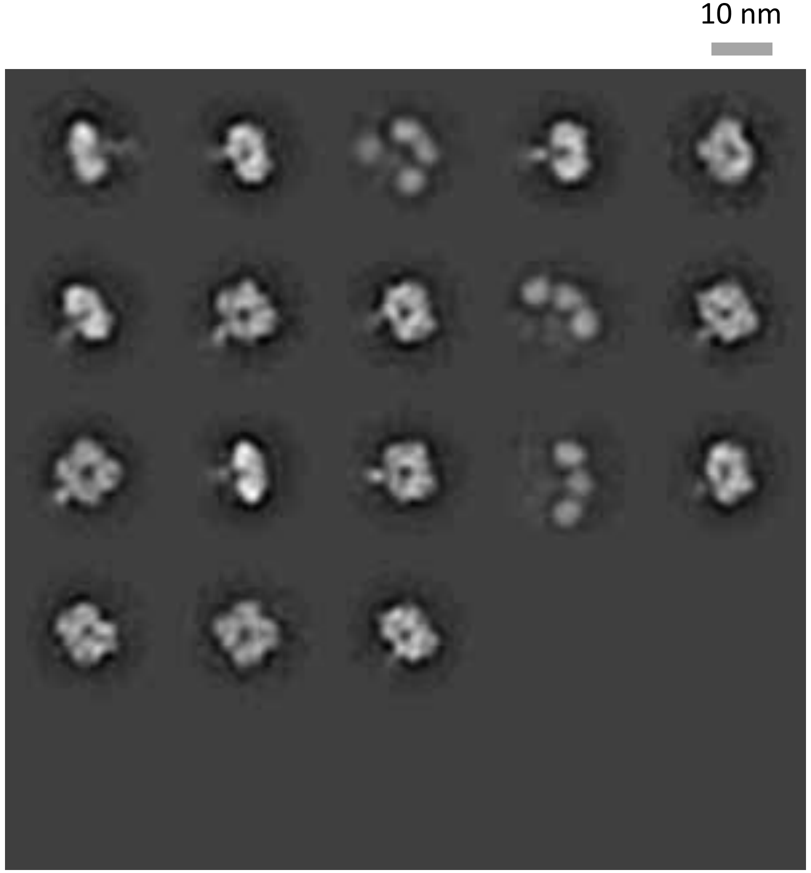

### N1-MI15-sNAp-165

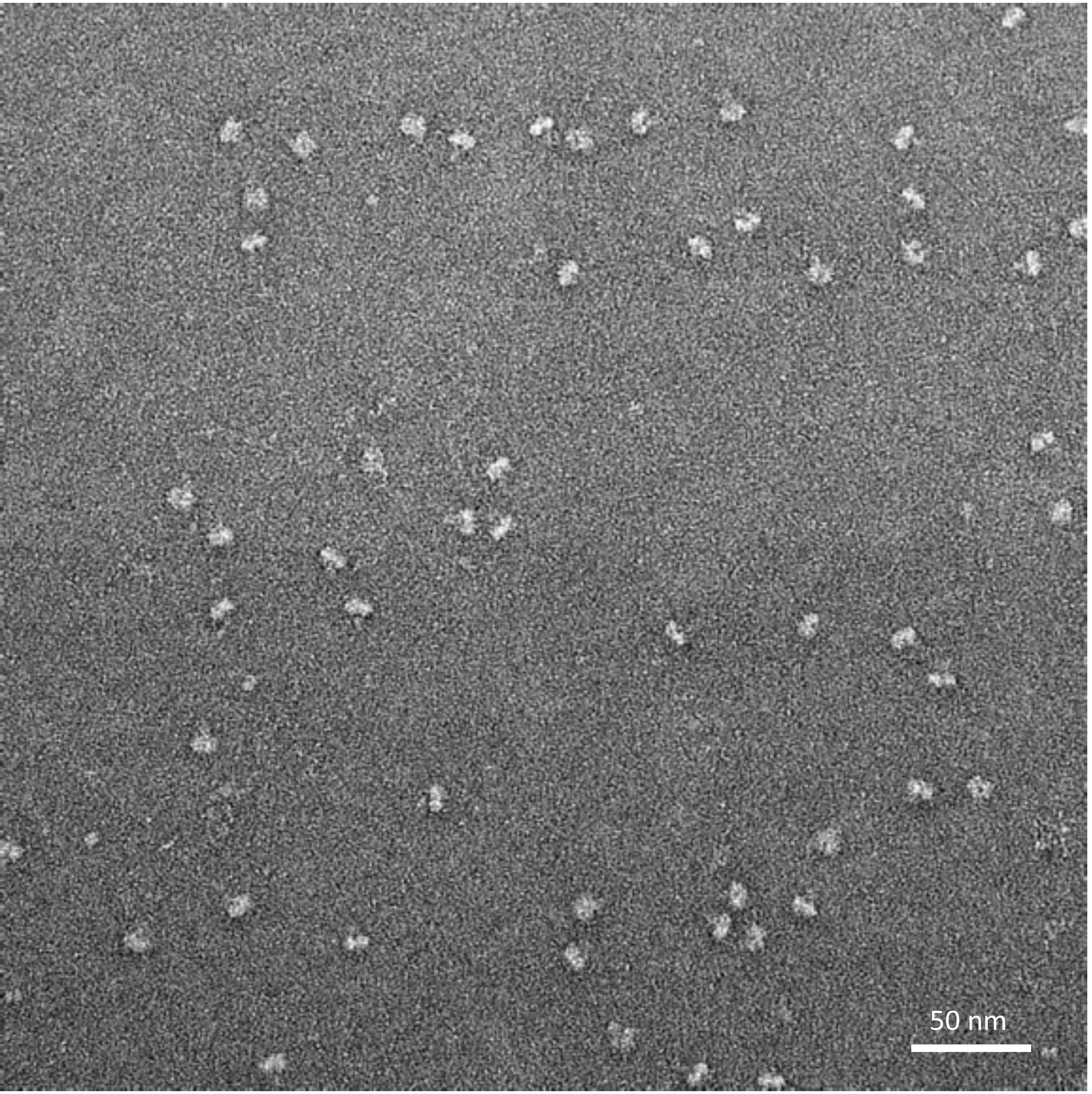


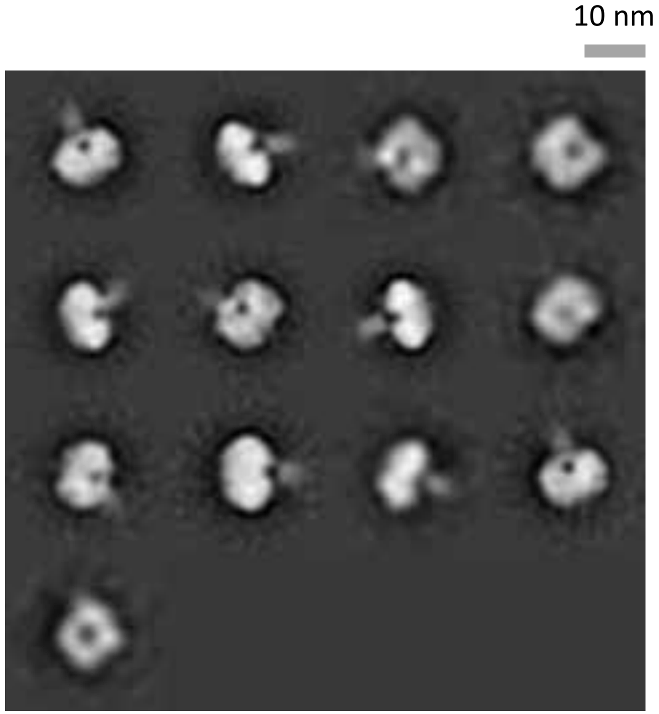

### N1-MI15-sNAp-176

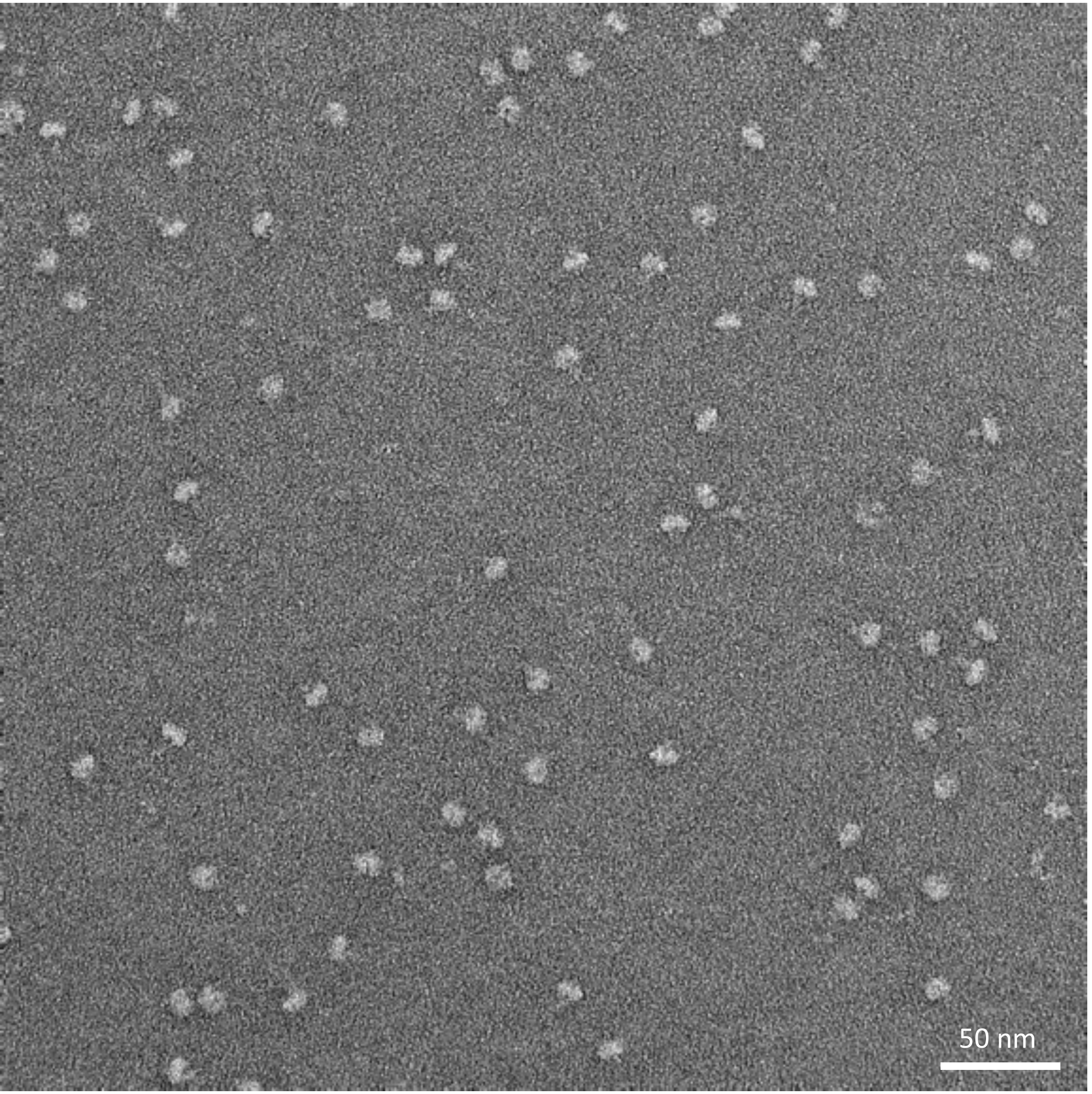


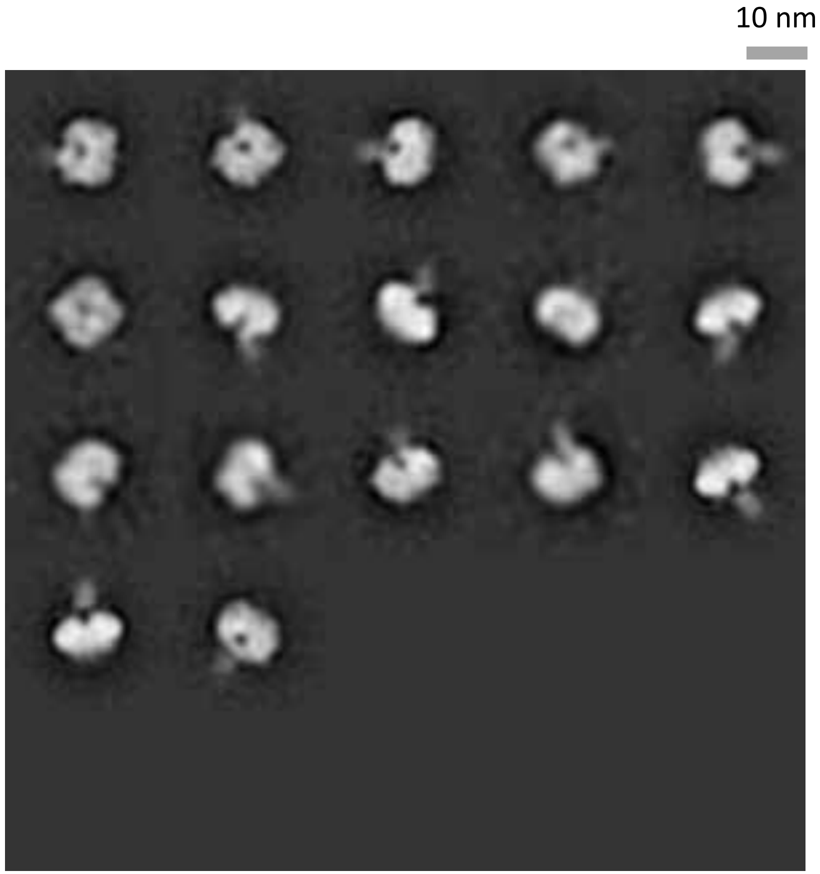

### N1-MI15-sNAp-183

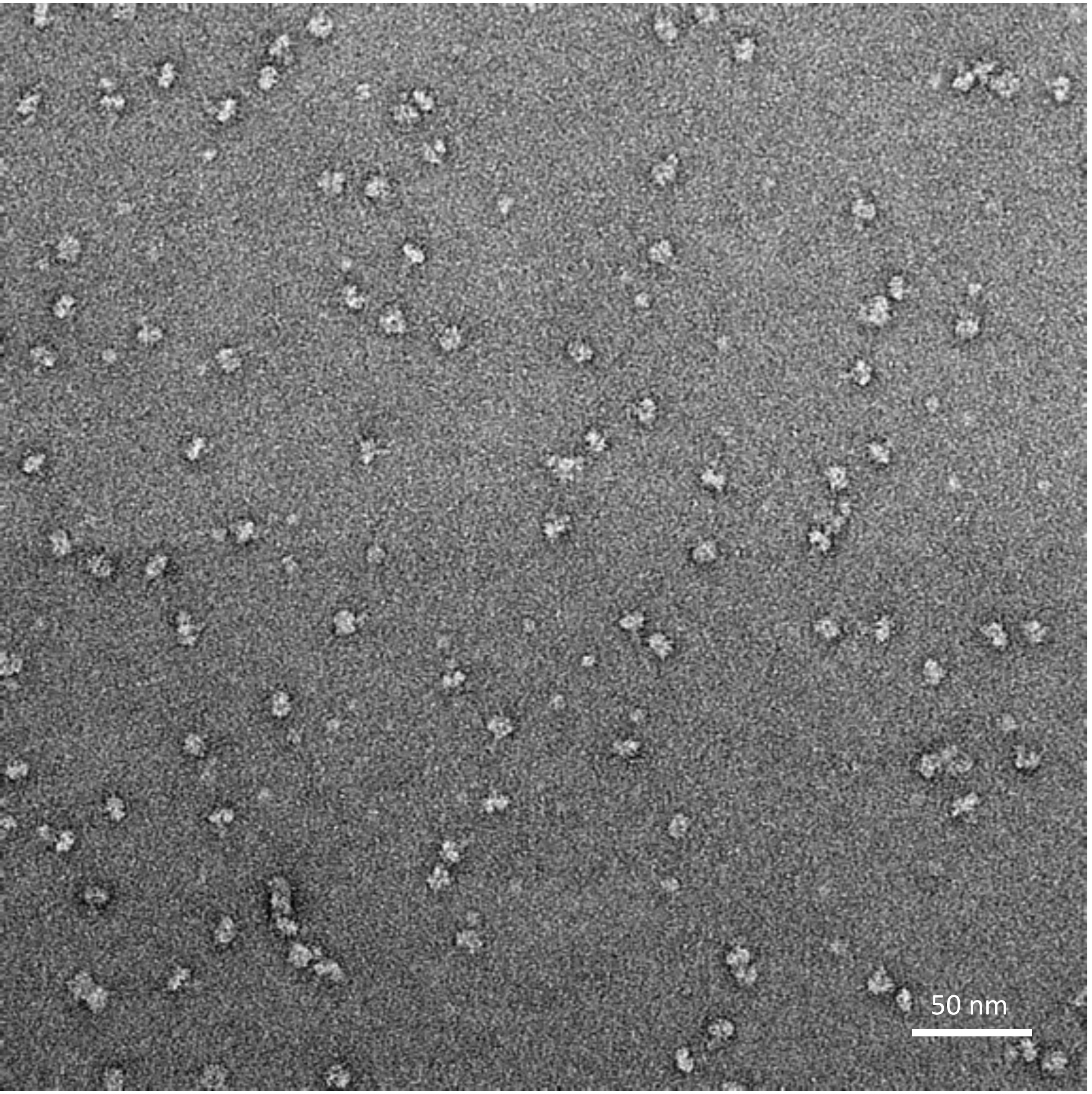


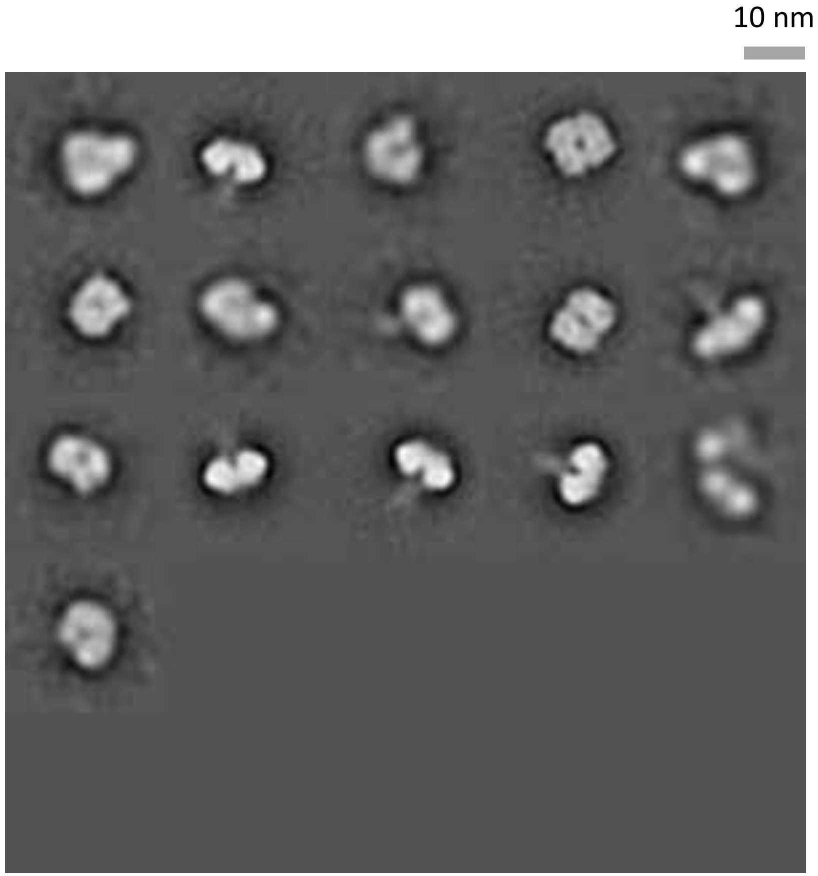

### N1-VN04-sNAp-155

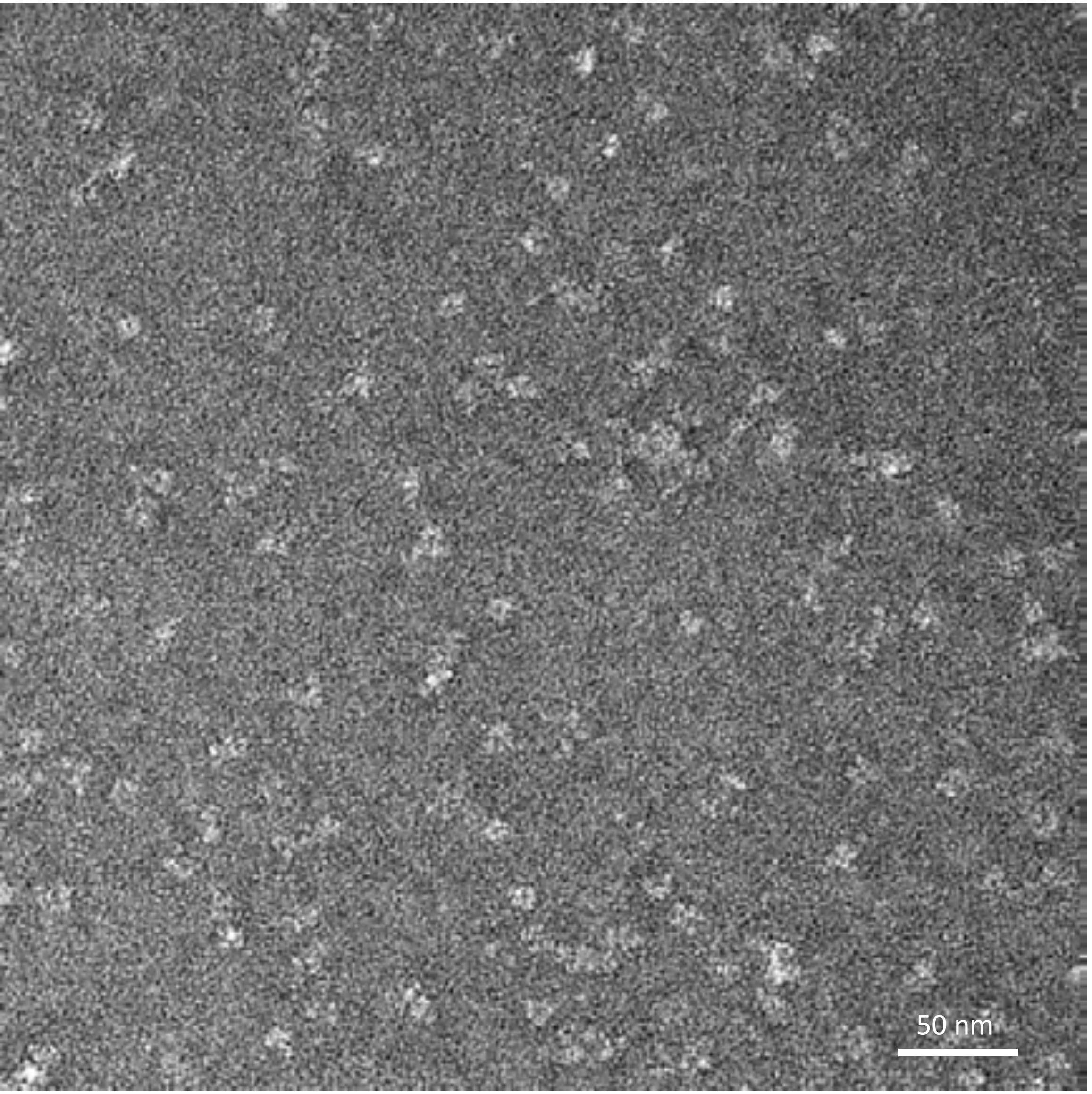


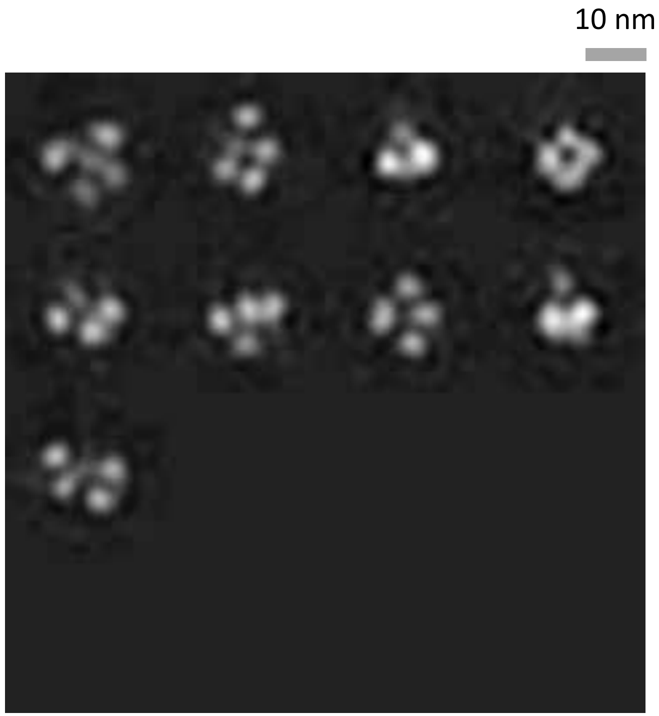

### N1-VN04-sNAp-354

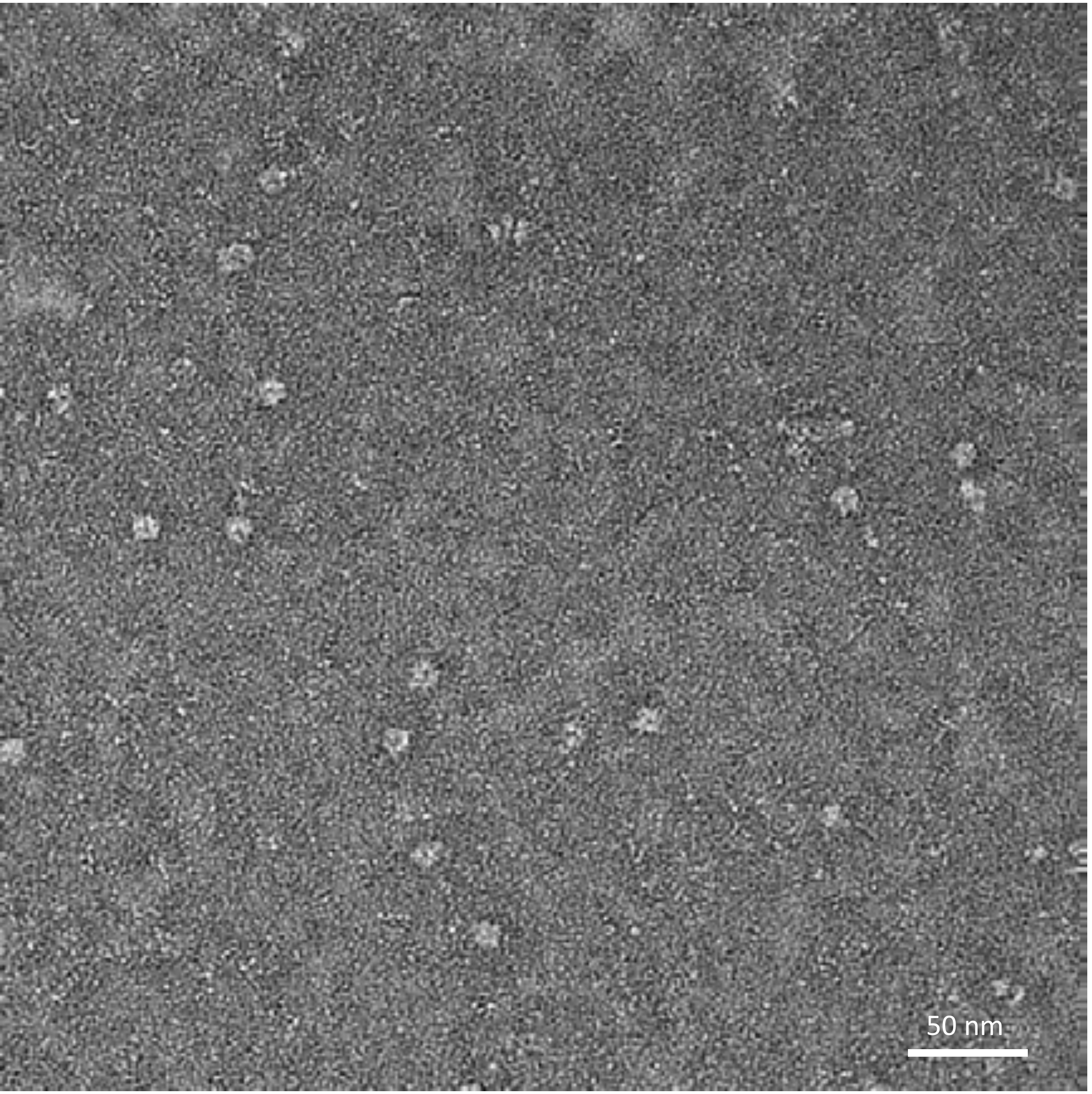


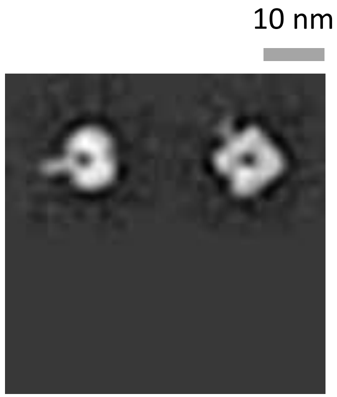

### N1-WSN33-sNAp-155

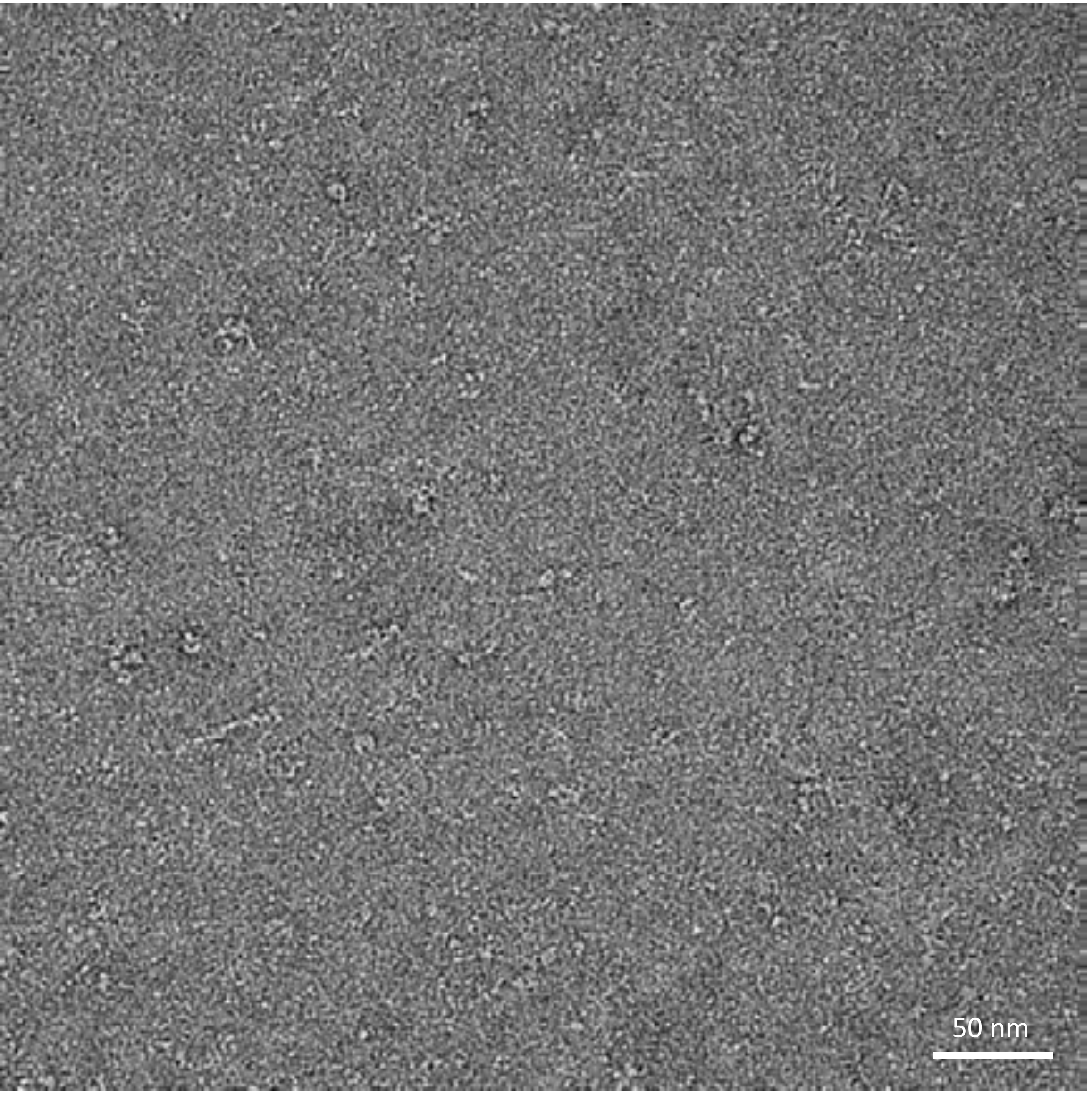


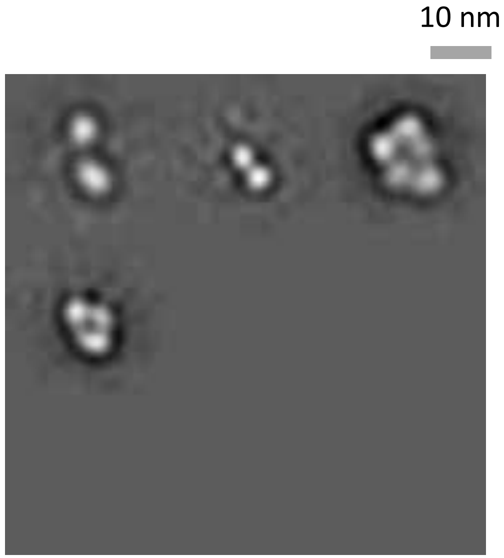

### N1-WSN33-sNAp-366

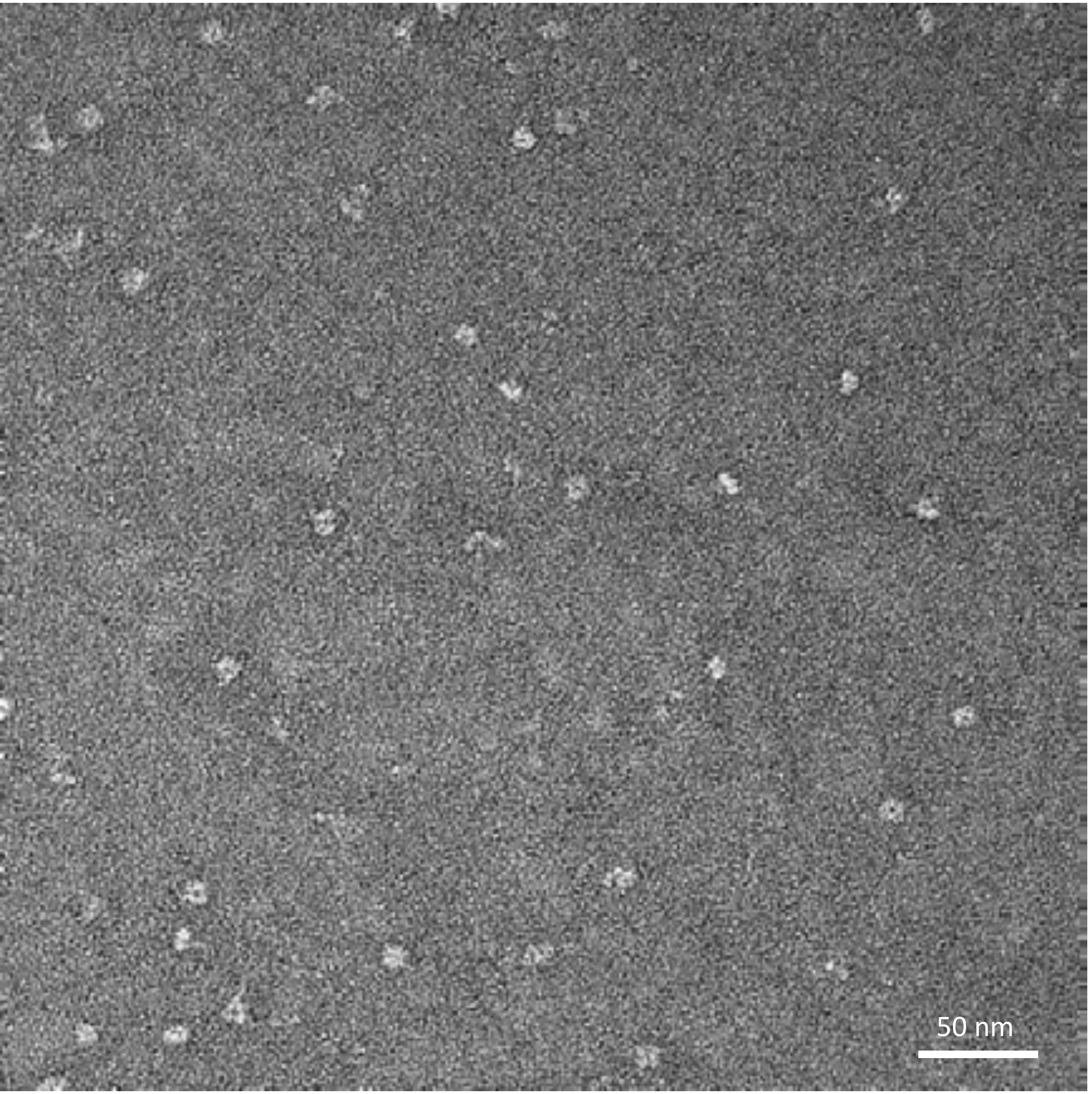


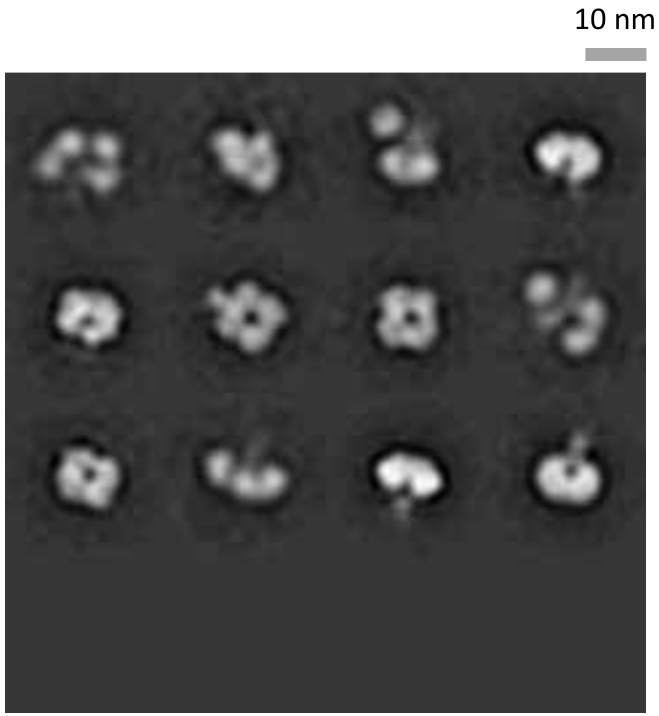

### N1-WSN33-sNAp-367

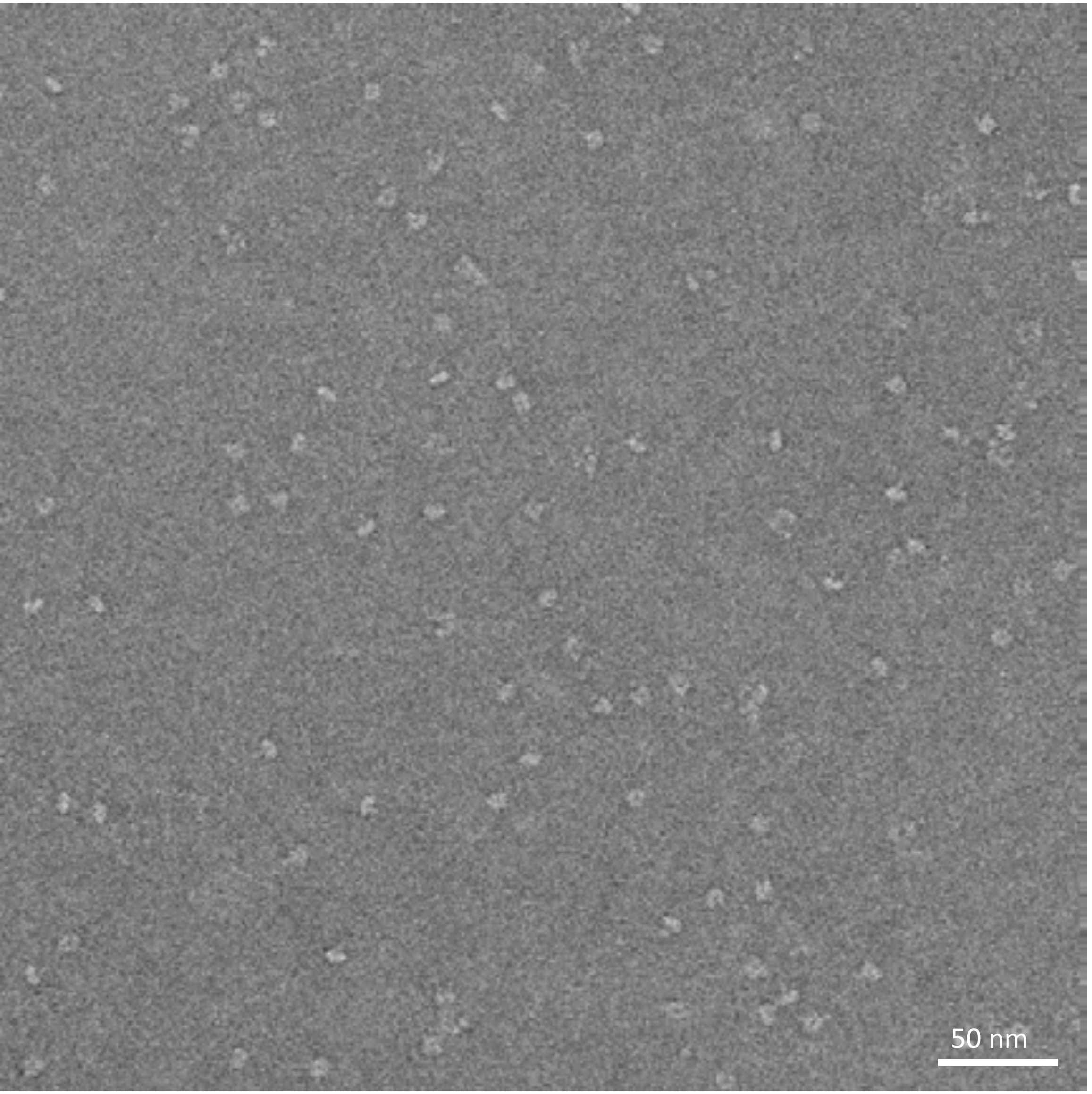


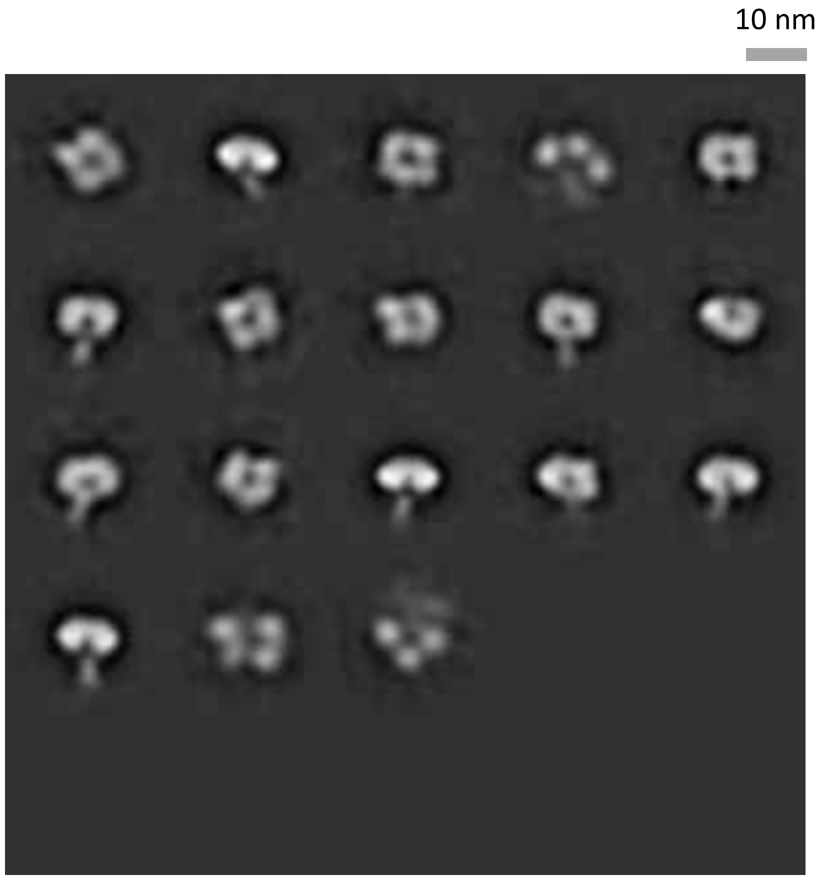

### N1-WSN33-sNAp-375

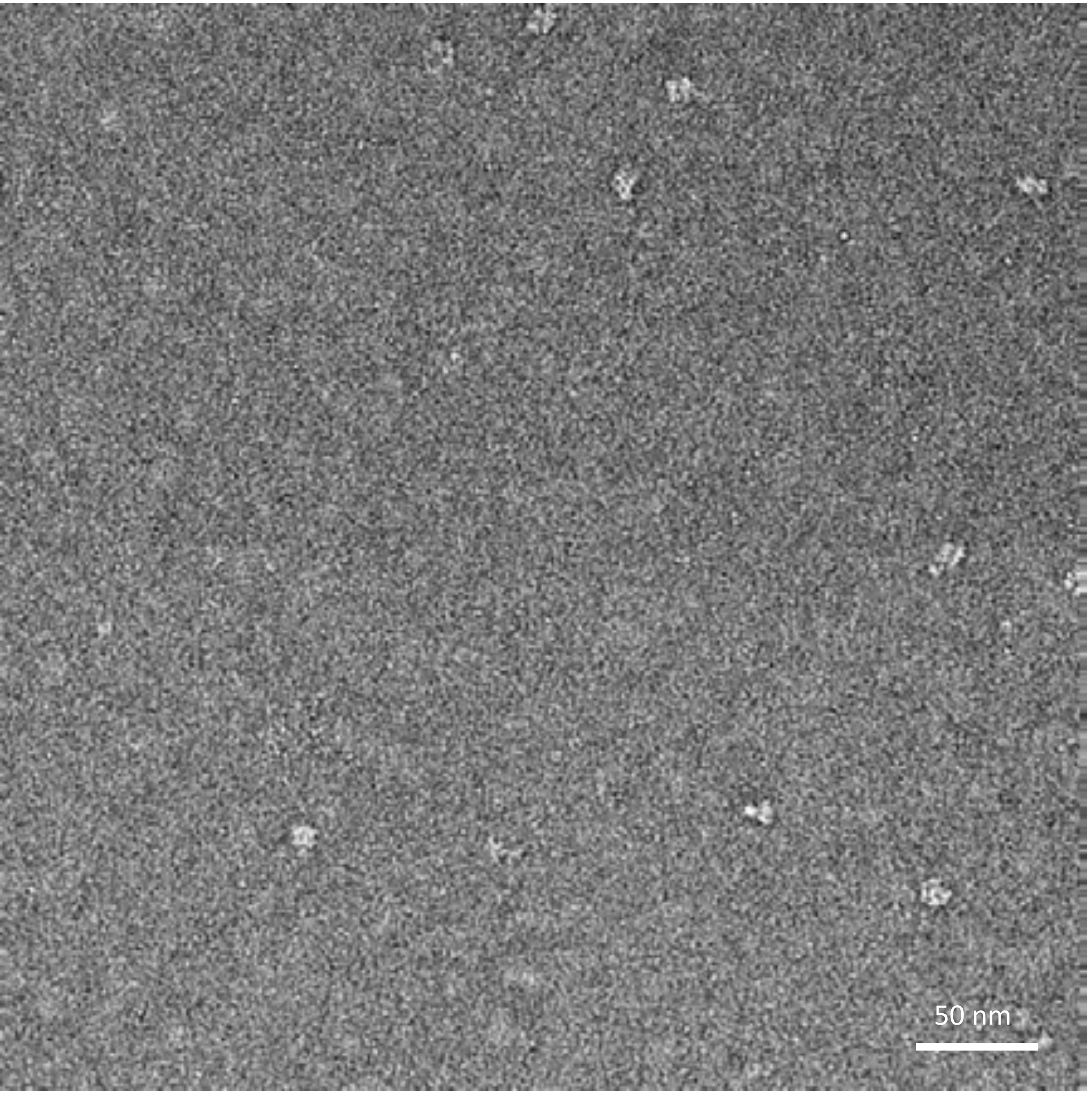


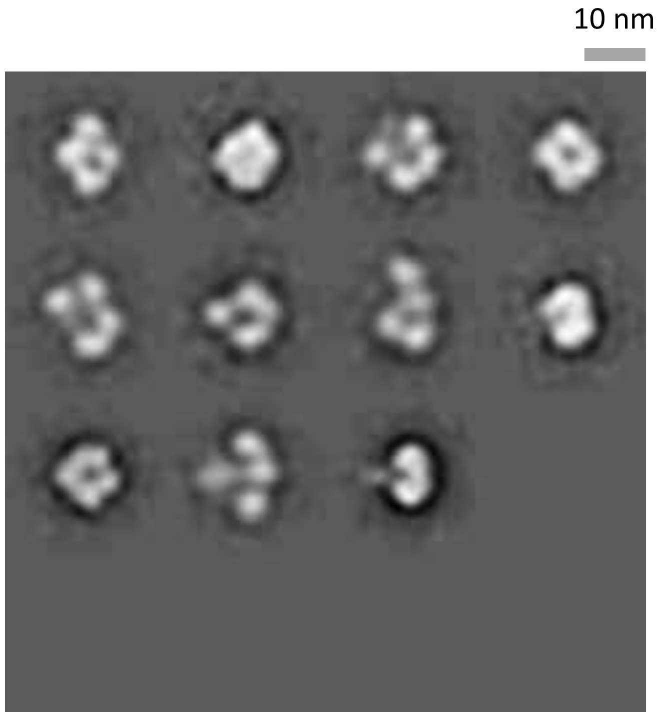

### N1-WSN33-sNAp-378

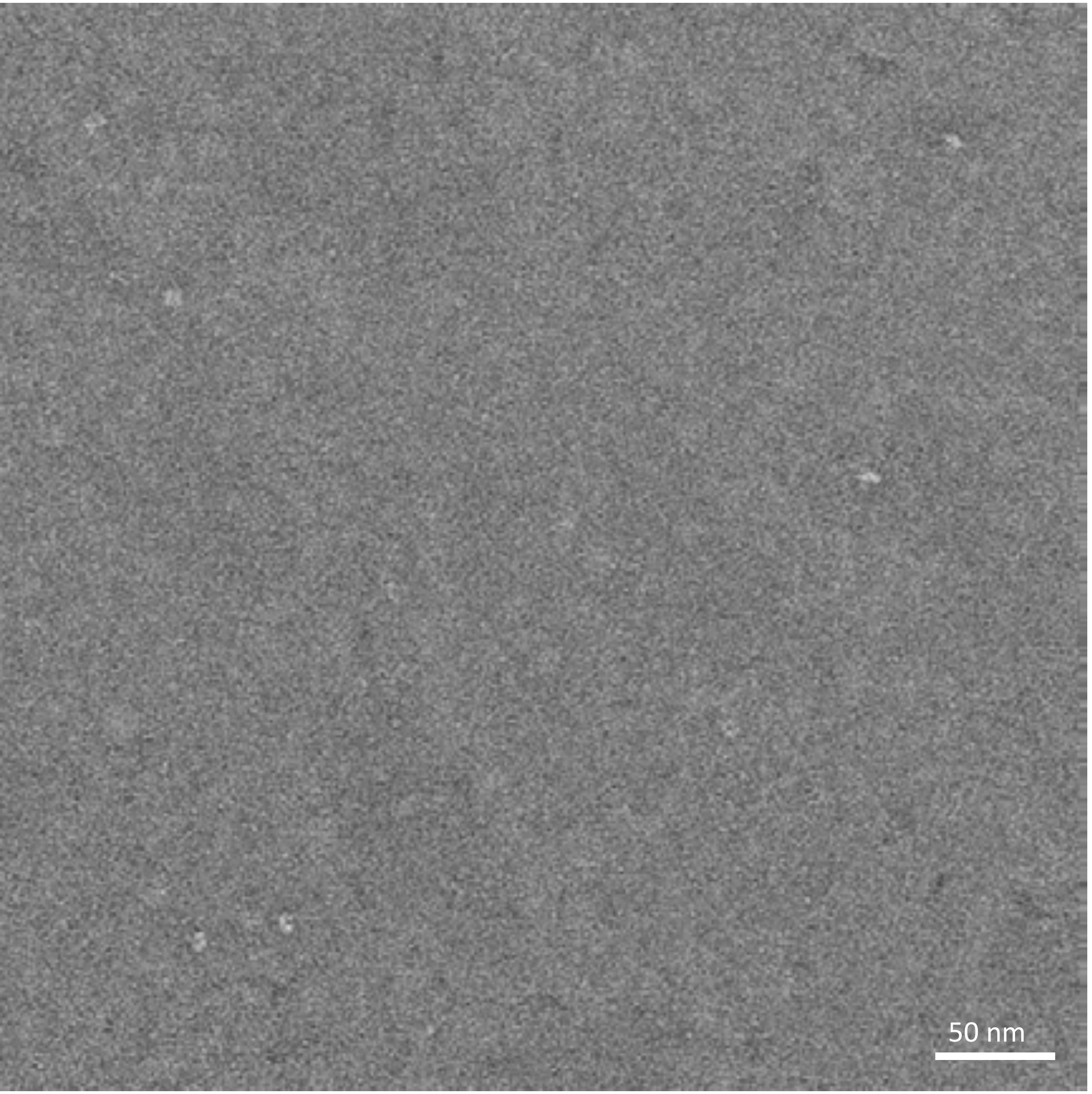


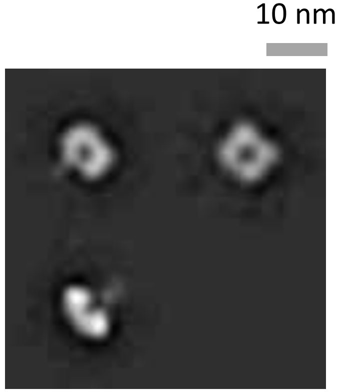

### N1-CA09-sNAp-155-T466A

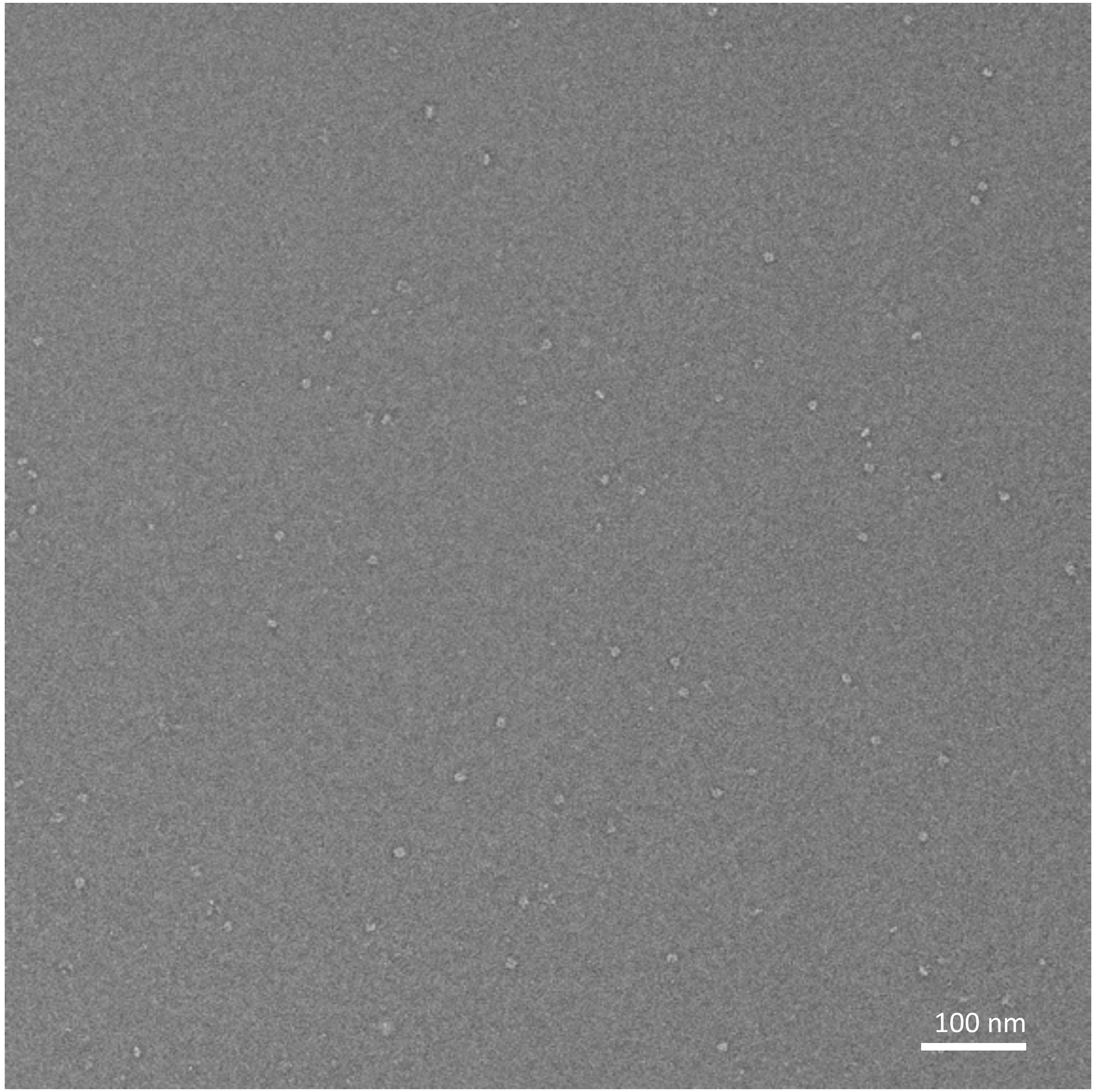


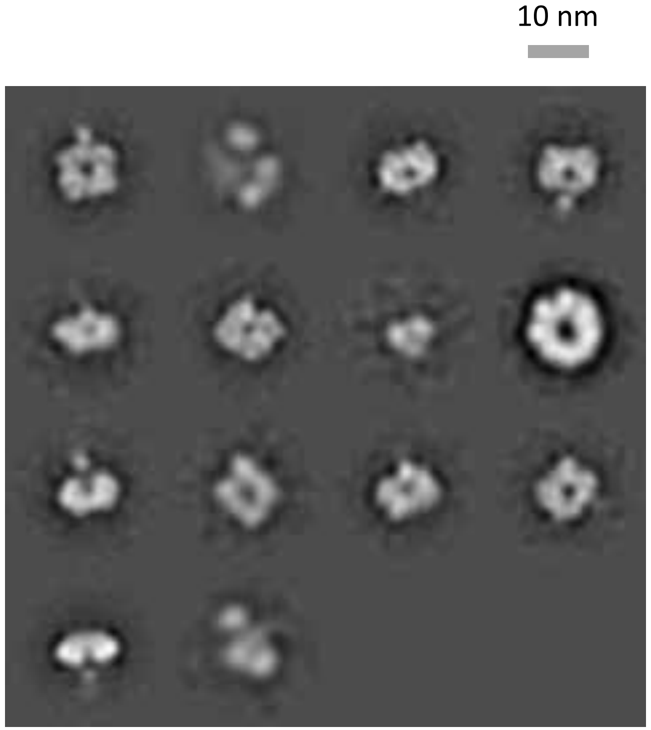
