## Supplemental Item 3 for "Structure-based design of stabilized recombinant influenza neuraminidase tetramers"

```
<ROSETTASCRIPTS>
	<SCOREFXNS>
		<ScoreFunction name="sfx_clean" weights="beta"
symmetric="1" />
	</SCOREFXNS>

	<TASKOPERATIONS>
		// Read resfile
		<ReadResfile name="designed_identities" filename="%
%resfile%%" />
	</TASKOPERATIONS>

	<RESIDUE_SELECTORS>
		<Task name="resfile_muts" designable="true"
task_operations="designed_identities" /> // selects mutations from
resfile
			<Not name="not_resfile"
selector="resfile_muts" /> // everything else that isn't mutated

		<Neighborhood name="mutant_neighbor"
selector="resfile_muts" distance="5" include_focus_in_subset="false" /
> // only operate on other residues that are 5A from mutation
		<Or name="resfile_or_nbors"
selectors="resfile_muts,mutant_neighbor" />
			<Not name="not_res_or_nbors"
selector="resfile_or_nbors" />

	</RESIDUE_SELECTORS>

	<TASKOPERATIONS>
		<IncludeCurrent name="ic" />	//includes current
rotamers
		<LimitAromaChi2 name="limitaro" chi2max="110"
chi2min="70" />  // disallow extreme aromatic rotamers
		<RestrictToRepacking name="repack_only" />
	// for minimize/repack
		<ExtraRotamersGeneric name="ex1_ex2" ex1="1"
ex2="1" />		// use ex1 ex2 rotamers

		<OperateOnResidueSubset name="repack_neighbor"
selector="mutant_neighbor" > // neighbors to mutations are only
allowed to repack
			<RestrictToRepackingRLT/> </
OperateOnResidueSubset>

		<OperateOnResidueSubset
name="lock_not_resfile_or_nbors" selector="not_res_or_nbors" > //
everything that isn't a mutation or neighbor cannot move
			<PreventRepackingRLT/> </
OperateOnResidueSubset>
```

```
        </TASKOPERATIONS>

        <MOVERS>
                <Symmetrizer name="gen_docked_config" symm_file="%
%symfile%%" />
                <SymPackRotamersMover name="design_from_resfile"
scorefxn="sfx_clean"
task_operations="designed_identities,repack_neighbor,lock_not_resfile_
or_nbors,limitaro,ic,ex1_ex2" /> // Adds mutation and repacks the
mutation and neighbors
                <TaskAwareSymMinMover name="rb_min_hard"
scorefxn="sfx_clean" bb="0" chi="1" rb="0"
task_operations="repack_neighbor,lock_not_resfile_or_nbors" /> //
minimizes mutations and neighbors
                <TaskAwareSymMinMover name="rb_min_hard_bb"
scorefxn="sfx_clean" bb="1" chi="1" rb="0"
task_operations="repack_neighbor,lock_not_resfile_or_nbors" />
                <SymPackRotamersMover name="repack_hard"
scorefxn="sfx_clean"
task_operations="repack_only,repack_neighbor,lock_not_resfile_or_nbors
,limitaro,ic,ex1_ex2" /> // repacks mutation and neighbors only

        </MOVERS>

        <FILTERS>
               <SaveResfileToDisk name="save_resfile"
task_operations="designed_identities" designable_only="0"
resfile_prefix="%%outpath%%" resfile_suffix="" resfile_name=""
resfile_general_property="NATRO" selected_resis_property=""
renumber_pdb="0" /> // save a resfile with mutations

        </FILTERS>

        <PROTOCOLS>
                // generate symmetric configuration
                <Add mover_name="gen_docked_config" />

                // design and minimization
                <Add mover_name="design_from_resfile" />
                <Add mover_name="rb_min_hard_bb" />
                <Add mover_name="design_from_resfile" />
                <Add mover_name="rb_min_hard_bb" />

                // save resfile
                <Add filter_name="save_resfile" />

        </PROTOCOLS>

</ROSETTASCRIPTS>
```
